# Declining food availability and habitat shifts drive community responses to marine hypoxia

**DOI:** 10.1101/2023.04.14.536810

**Authors:** E Duskey, M Casini, KE Limburg, A Gårdmark

**Author notes:** Corresponding author. E­mail. **Open Research statement:** All data and parameters necessary to replicate this modeling study are available at https://github.com/epduskey/hypoxiaSSM. The code contained therein is a novel, extensive modification of the R package mizer, available at https://github.com/sizespectrum/mizer. Data and code will be archived in Zenodo upon publication.

## Abstract

Worsening marine hypoxia has had severe negative consequences for fish communities across the globe. While individual­ and population­level impacts of deoxygenation have been identified, it is unknown how they interact to drive changes in food webs. To address this, we incorporated several major impacts of hypoxia, including declines in benthic resources, habitat shifts, increasing mortality, and changes to rates of feeding, assimilation, and reproductive efficiency, into an existing size spectrum food web modeling framework. We used this structure to ask the following questions: which of these direct effects are most critical to capturing population and community dynamics in a representative hypoxic system, how do they interact to result in community responses to deoxygenation, and what are the potential consequences of these effects in the context of accelerating deoxygenation? We tested the effect of different combinations of oxygen­dependent processes, driven by observed oxygen levels, on the food web model’s ability to explain time series of observed somatic growth, diets, biomass, and fishery yields of commercially relevant species in the Baltic Sea. Model results suggest that food availability is most critical to capturing observed dynamics. There is also some evidence for oxygen­dependent habitat use and physological rates as drivers of observed dynamics. Deoxygenation results in declining growth both of benthic and benthopelagic fish species, as the latter are unable to compensate for the loss of benthic resources by consuming more pelagic fish and resources. Analysis of scenarios of ideal, declining, and degraded oxygen conditions show that deoxygenation results in a decline in somatic growth of predators, an altered habitat occupancy resulting in changing species interactions, and a shift in energy flow to benthopelagic predators from benthic to pelagic resources. This may have important implications for management as oxygen declines or improves.

## Introduction

Marine hypoxia occurs when dissolved oxygen declines to levels that are sub­optimal for aerobic life. Hypoxia may occur naturally in some systems, but it has increased worldwide in both severity and frequency due to anthropogenic nutrient enrichment (Breitburg et al. 2018), warming­induced increases in oxygen consumption (Brewer and Peltzer 2016) and stratification (Diaz and Breitburg 2009), and decreases in oxygen solubility with rising ocean temperatures, particularly in the epipelagic (Schmidtko et al. 2017). There are a wide variety of negative impacts of hypoxia on fishes, ranging from altered functioning of cells to community reorganization (Pollock et al. 2007). This presents a challenge to fish species and their prey and hinders society’s ability to use ocean resources sustainably (United Nations 2022).

The effects of deoxygenation are documented at several levels of biological organization. For example, at the population level, we might observe changes in reproductive potential (Wang et al. 2016) and habitat use (Orio et al. 2019; Zhang et al. 2014). Changes in the chemical environment may promote the production of toxicants, resulting in increased mortality (Diaz and Breitburg 2009). At the community level, differential responses among species or functional groups can result in spatial reorganization (Chu and Gale 2017), changes in species composition (Slater et al. 2020), and adaptive foraging resulting in significant changes in food web structure and energy flow (Breitburg et al. 1999; Breitburg et al. 2003; Pollock et al. 2007). Observations at these higher levels of organization arise from behavioral and physiological responses of individuals.

There are several compensatory mechanisms that individual fish employ in order to mitigate the effects of hypoxia, including an increase in ventilation and gill perfusion, increased hemoglobin and oxygen extraction, decreases in activity (Farrell and Richards 2009), as well as behaviors such as aquatic surface respiration (ASR; Dwyer et al. 2014), and changes in swimming activity (Domenici et al. 2013) and habitat occupancy (Eby et al. 2005; Stramma et al. 2012; Casini et al. 2019). Indirect effects on fish may also be common. For example, benthic invertebrates may significantly alter behavior and ultimately be subject to increased mortality under hypoxia (Riedel et al. 2014). This may lead to changes in the diets of predatory fish that feed on them (Powers et al. 2005). Overall, there is a good body of evidence linking low oxygen concentration to individual responses of fish, less evidence indicating how these result in population­level responses (Bergman et al. 2019), and relatively little evidence describing how these in turn im­ pact community­level responses (Rose et al. 2009). Of the latter, investigations of the impacts of hypoxia on predator­prey interactions suggest that differential sensitivity of each species to declining oxygen can result in changes in spatial overlap, predator activity, and prey vulnerability (Breitburg et al. 1997; Chu and Gale 2017). However, it is unknown which of these mechanisms, if any, are most critical and how they interact to drive the observed, higher­level dynamics.

Integrative modeling may help to resolve some of these unknowns by drawing direct connections between the effects of environmental drivers on individuals and higher­level outcomes (Pol­ lock et al. 2007). Hypoxia operates on several physiological and behavioral pathways of fish simultaneously. Its impacts are dependent on body size (Ekau et al. 2010), and are often caused by changes in feeding (Wang et al. 2009). Hypoxia may thus affect individuals, cause emergent effects on populations, and impact the overall community structure by changing competitive dominance and feeding relationships. Therefore, a size structured food web model offers an appropriate framework with which to evaluate the relative importance of major individual­level pathways suggested by the literature as potential drivers of observed population and community dynamics in hypoxic habitats (Koenigstein et al. 2016).

In this study, we developed an oxygen­dependent version of a size spectrum food web model in order to ask the following questions: (1) Which individual­level impacts of deoxygenation are most critical to capturing observed population and community dynamics? (2) How do they interact to result in community­level responses to deoxygenation? and (3) What are the potential consequences of these effects in the context of accelerating deoxygenation? We used the Baltic Sea as a case study and fitted several parameters to a simplified fish food web to explore how environmental hypoxia can affect community composition and dynamics, as well as characteristics of the commercial fisheries yield. By testing the ability of several individual­level pathways – including food availability, vertical habitat occupancy, mortality, and physiological rates of feeding, reproduction, and assimilation – to describe observations at the population and community scale, we identified potential physiological and behavioral drivers of the deleterious effects of hypoxia on marine food webs.

## Methods

### Size Spectrum Food Web Model

We modified the size­structured food web modeling framework **mizer** (Scott et al. 2014) in order to represent fish community dynamics in response to declines in benthic oxygen levels. In food web models of this sort, maximum body size serves as the master trait of each species, determining somatic growth through feeding, reproduction through maturation size and growth, and mortality through size­dependent predation and natural senescence (Peters 1986; Blanchard et al. 2017). For our model, we built a general structure (found here) composed of a benthopelagic predator, a benthic predator, two species of pelagic forage fish, as well as these species’ basal resources in separate benthic and pelagic habitats. This system reflects a community with two competing benthic predators, one of which is more resilient to hypoxia than the other, two competing pelagic prey fish species, and benthic and pelagic invertebrate fauna on which they feed.

The fundamental and theoretical foundation of these models is described in Andersen et al. (2016). They are generally defined by three major ecological processes, all of which scale with body size: somatic growth, reproduction, and mortality. The dynamic size spectrum of each species and basal resource is determined by a flow of energy, or equivalent mass, via trophic relationships, as well as mortality. Conservation of mass is guided by the McKendrick­von Foerster equation (Silvert and Platt 1978):

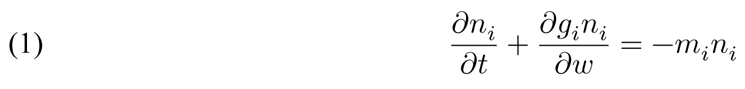

where *n_i_*, *g_i_*, and *m_i_* describe time­dependent population size, somatic growth, and mortality as a function of body size for each species *i*, respectively. Recruitment of individuals of species *i* at a minimal size *w_0_* is described by the boundary condition:

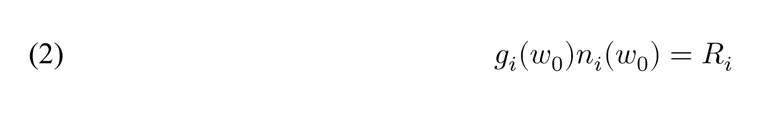

Losses of individual biomass construed as metabolic activity, which also scales with body size (see Brown et al. 2004), and assimilation efficiency, determine the conversion of ingested mass into somatic and reproductive growth. Each individual of each species is recruited to the population at weight *w_0_* and grows by consuming a combination of basal resources and other fish. Growth is also determined in part by species­specific von Bertalanffy growth parameters (Essington et al. 2001), here estimated from data. The basal resources are governed by a simple semi­ chemostat function, as well as by the degree of predation. Recruitment is limited by density de­ pendent processes, typically by the inclusion of a Beverton­Holt or Ricker type stock­recruitment relationship. Feeding is guided by size­ and species­preference, determined by both a distribution describing the preferred predator:prey mass ratio (PPMR) and a square matrix describing horizontal spatial overlap of each species with all others. A maturity ogive guides the allocation of acquired energy to somatic growth versus reproduction throughout an individual’s life. Mortality occurs either through predation, unspecified natural processes, or fishing, the latter of which is determined by user­defined fishing selectivity, effort, and catchability. We estimated each of these parameters with data arising from the Central Baltic Sea (Appendix S2; Figures S1–S5 and Tables S1 and S2 for parameter estimates and data sources). Given this model structure’s focus on individual physiological processes and how these govern community dynamics, it is appropriate for the inclusion of documented effects of hypoxia on feeding, habitat occupancy, metabolism, and mortality (see Figure 1).

**Figure 1.**
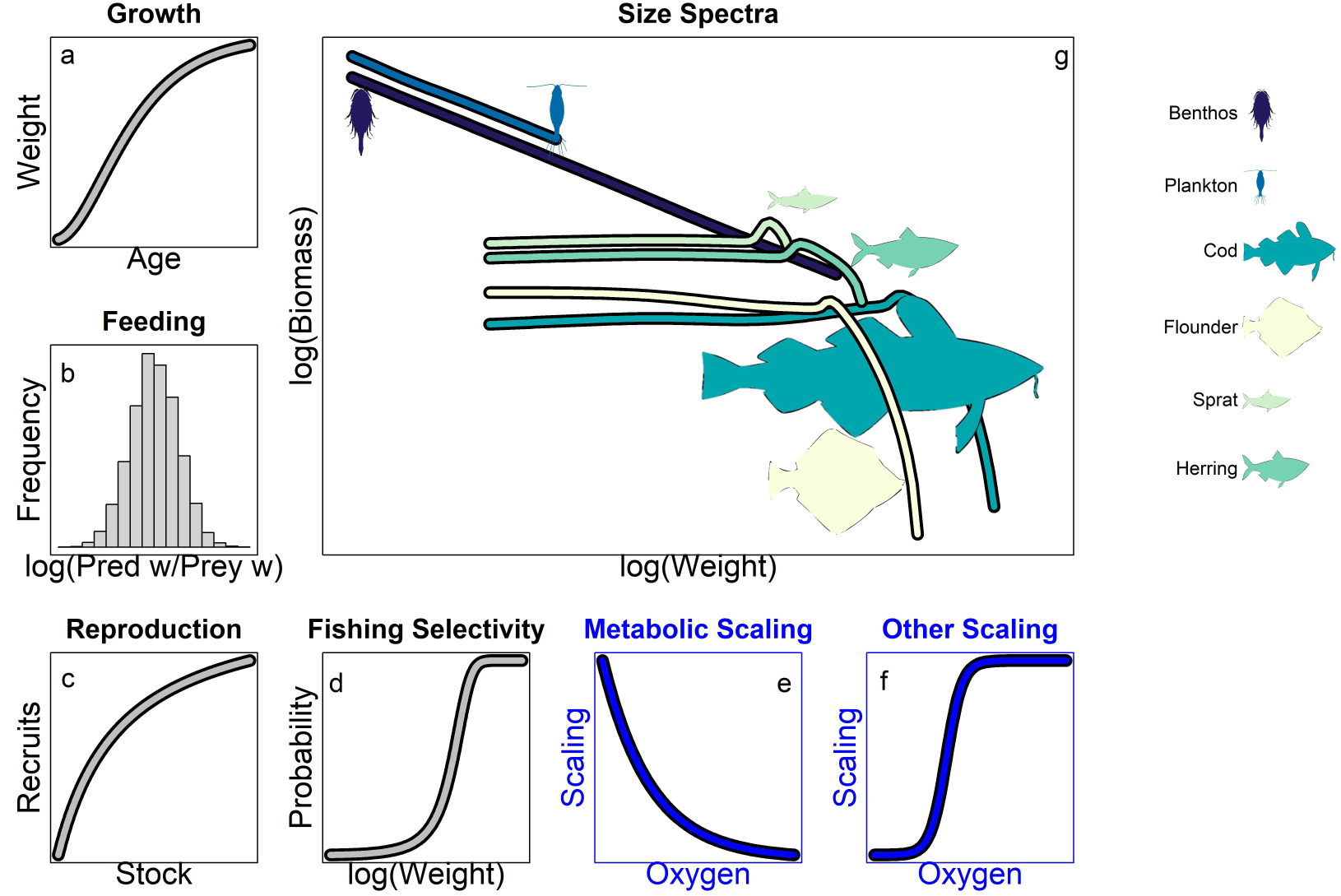
A conceptual illustration of key components and outcomes of the oxygen­dependent size spectrum food web model. Quantitative descriptions of food­ and size­dependent growth (a), size­based feeding preference (b), density dependent reproduction (c), fisheries size selectivity (d), and size­dependent maturation determine the central processes of growth, reproduction, and mortality. In this model, metabolism depends on oxygen with a negative exponential function (e), whereas other physiological processes scale with oxygen according to a logistic function. Together, these processes produce an emergent number and mass of individuals in the population (size spectra) of each of the interacting fish species (g).

### Oxygen dependence

In general, we included deleterious effects of hypoxia in our model by increasing costs and decreasing vital physiological and ecological rates with bounded and continuous functions of oxygen decline (Appendix S1). Though the hypoxia threshold is often defined as an oxygen concentration of 2 mg⋅L^­1^ (Hrycik et al. 2017), a range of concentrations above this threshold are known to have sub­lethal, yet critical effects on the ecology of fishes (Kramer 1987). These effects may or may not vary by body size (Hrycik et al. 2017; Pan et al. 2016), but they do vary by species (Nilsson and Östlund­Nilsson 2008). In a size spectrum food web framework, each individual may be affected to a greater or lesser degree relative to its competitors. Therefore, within these models, hypoxia has the potential to change competitive and predatory interactions, and the direction and magnitude is dependent on several characteristics of the species involved. This flexibility, combined with a focus on individuals throughout ontogeny and ultimately leading to dynamics at a scale at which they are observed, allows us to ask specific questions about the relevance of individual processes at the community scale.

More specifically, we altered the baseline structure of **mizer** (Scott et al. 2014) by adding dynamic independent variables for benthic and pelagic oxygen, given here in units of mL⋅L^­1^. We used the concept of critical oxygen level, or P_crit_, to describe individuals’ relative sensitivity to estimated oxygen levels. The P_crit_ of an individual is the oxygen level below which standard metabolism can no longer be maintained and begins to decline with ambient levels (Ultsch and Regan 2019). That is, it is the level at which oxyregulators (i.e. organisms which maintain constant oxygen consumption) become oxyconformers (i.e. organisms whose oxygen consumption varies with environmental conditions) due to the lack of adequate oxygen to cover basal metabolic costs (Rogers et al. 2016). This value depends not only on species identity (Farrell and Richards 2009), but also on salinity, temperature, and body weight (Rogers et al. 2016). While we kept salinity and temperature constant, maximum weight is one of the defining attribute of individuals in **mizer**, and therefore both behavioral and physiological responses to oxygen may change throughout ontogeny.

We used a P_crit_ database (Rogers et al. 2021) to estimate P_crit_ as a function of temperature, salinity, body size, and resting metabolic rate (RMR), as in Rogers et al. (2016) (Appendix S4; see Figure S7). We scaled the following rates and values as a logistic function of each species’ oxygen exposure, where applicable: benthic resource carrying capacity, occupancy in the benthic habitat, maximum consumption rate, fish search rate for prey, assimilation efficiency, and fish egg survival. The general equation is:

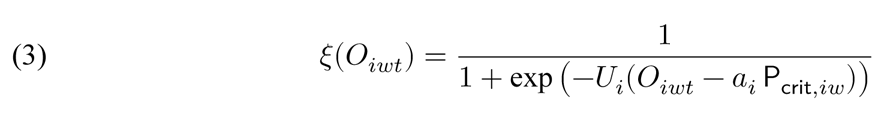

where *O_iwt_* is mean oxygen concentration experienced by species *i* of weight *w* at time *t*, and *U_i_* captures the sensitivity of species *i* to low oxygen, with lower values representing more gradual declines at higher oxygen levels. The product *a_i_* P_crit,*iw*_ represents the oxygen level at which rates of an individual of species *i* at weight *w* have fallen to 50% of their maximum. This scaling function is very similar to that used by Luo et al. (2001), and reflects a simple and flexible method by which to include individual responses to oxygen. All the rates that we scaled according to equation 3 represent activities in excess of standard metabolic rate (SMR). SMR as represented by oxygen consumption can be approximated by P_crit_ (Ekau et al. 2010), and thus treating P_crit_ as a shifting inflection point may be appropriate. We estimated one set of parameters for benthic resource carrying capacity, another for benthic occupancy, and a third set for maximum consumption rate, fish search rate for prey, assimilation efficiency, and egg survival. Note that inherent in this structure is the assumption that all oxygen­dependent physiological processes de­ cline with oxygen at the same rate. That is, the prioritization of activity, feeding, assimilation efficiency, etc. of fishes of each species does not change as hypoxia develops, aside from variation in P_crit_ due to body size and species, as indicated in equation 3. This is a strong assumption, but it significantly reduces the number of parameters that must be estimated, and serves as a simple starting point for analyses of emergent community responses to deoxygenation.

In addition to the rates listed above, we also scaled metabolism and natural mortality (i.e. mortality not due to fishing or, in this structure, predation) with exposure to hypoxic waters. Metabolism in the **mizer** framework is treated as a cost, based on the assumption that costs for maintenance scale proportionally to metabolic rate. This is because energy is assumed to flow through the system as mass equivalents, with somatic growth arising as a balance between gains via consumption and losses via respiration. We therefore scaled it with a simple negative expo­ nential function:

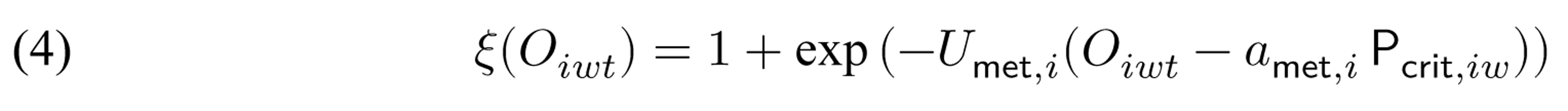

Here, values of *U*_met,*i*_ and *a*_met,*i*_ play a similar role as their counterparts in equation 3, only *a*_met,*i*_ P_crit,*iw*_ instead represents the oxygen level at which metabolic costs have doubled. Additional natural mortality due to exposure to hypoxic waters is expressed as a hazard function (Appendix S4). Both this and occupancy as a function of dynamic variables are new additions to the **mizer** framework, and reflect the flexibility built into the code provided (Scott et al. 2014).

### Study System

We studied emergent responses to hypoxia in the relatively species poor food web of the Central Baltic Sea, which here is mainly focused on commercially important fish species in the open waters. This is an important case study, given the severity and extent of deoxygenation in the region (Carstensen et al. 2014). Our model of this system includes benthopelagic cod (*Gadus morhua*), benthic flounders (*Platichthys flesus* and *Platichthys solemdali*), and the pelagic clupeids sprat (*Sprattus sprattus*) and herring (*Clupea harengus*), as well as their benthic and pelagic invertebrate prey. This region is particularly prone to episodes of deoxygenation, with a water residence time of roughly 25­30 years, and a relatively stable vertical salinity gradient that iso­lates bottom waters from oxygenated surface waters (Snoeijs­Leijonmalm and Andrén 2017). Periodic major Baltic inflows (MBIs) rejuvenate oxygen concentrations in these waters, but with decadal variability and with a lower frequency in the last three decades (Mohrholz 2018). Though recent nutrient loading reduction has led to improving conditions, extensive eutrophication and a legacy of recycling nutrients continue to plague the Baltic Sea (McCrackin et al. 2018; Andersen et al. 2017). As a result, hypoxic areas have expanded exponentially since the mid­ 1990s, reducing the potential habitat for benthic fish such as cod (Limburg et al. 2011; Casini et al. 2016; Casini et al. 2021). Fish otolith microchemistry has also confirmed individual cod exposure to low oxygen, with negative effects on growth and body condition (Limburg and Casini 2018; Limburg and Casini 2019). For the Baltic Sea, there are extensive fisheries data arising from scientific surveys and commercial data collection (ICES 2020), including mean weight as well as time series of spawning stock biomass (SSB, the combined weight of all reproductively active individuals in the population), and fisheries yield (for the derivation of observation data, see Appendix S2). For this system, we estimated oxygen trends from data provided by the Swedish Meteorological and Hydrological Institute (SMHI; Appendix S3; Figure S6 and Table S3). The availability of these data makes it ideal for studying the mechanisms that determine the impacts of deoxygenation on fish communities.

### Oxygen­dependent processes explaining observed population trends

We grouped the oxygen­dependent processes into four general categories: benthic resource carrying capacity (B), cod benthic occupancy (O), mortality (M), and physiology (P) (Appendix S1). The physiological component (P) includes both increases in metabolic costs and declines in maximum consumption rate, search or clearance rate, assimilation efficiency, and reproductive efficiency. We fit all 16 possible combinations of these groups, including both the full model and a model with no scaling, to observations of spawning stock biomass (SSB), commercial yield,and somatic growth of the four fish species during a calibration period (1991–2000; Appendix S5). These models were then used to project forward during the years 2001–2019. We chose the best model based on minimum weighted error during this projection period only, as out­of­ sample model selection is generally more robust (Cooke et al. 2014). We also performed a sensitivity analysis (Appendix S6) to determine the robustness of our results. This procedure can provide insight into which pathways by which hypoxia affects individual fishes that are most critical to an accurate representation of the observed community dynamics.

### Scenario analyses

We used the best model from the procedure described above (and detailed in Appendix S5) in order to test community responses under various oxygen scenarios. We ran the calibrated models during the projection period (2001–2019) with all individual components of the best model in order to determine how they interact to produce the responses in each scenario. These scenarios tracked predicted SSB, yield, and fish weight (in grams) up to maximum observed age in data sources (15 years for cod, 26 for flounder, 16 for sprat, 17 for herring), as well as energy or equivalent mass flow through the food web. The latter values are calculated as a weighted average of diets for each fish species across body size classes. Weights are given by the biomass in each body size class.

For each of these models, we ran three scenarios at benthic oxygen levels spanning the range of observed values throughout the time series. Mean bottom oxygen concentration reached a peak value of roughly 3 mL⋅L^­1^ during a period of inflow in the late 1980’s and early 1990’s and fell to 1 mL⋅L^­1^ in the 2000’s and beyond (Appendix S4; Figure S6). Therefore, we chose values of 3, 2, and 1 mL⋅L^­1^ representing more ideal, deteriorating, and degraded conditions. We then compared somatic growth, SSB, yield, and energy flow among the scenarios to determine how communities respond to declining oxygen.

### Implementation

We estimated all parameters, built all models, and analyzed and plotted all output in R (R Core Team 2021). All code and parameters necessary to reproduce our results are available on GitHub (found here).

## Results

### Oxygen dependence of observed dynamics

Model calibration results suggest there are interactions among the pathways by which deoxygenation affects individual fish. No single process nor a combination of processes provided a superior representation of somatic growth, SSB, and yield observed in the four fish species (cod, flounder, sprat, and herring) during the projection period (2001–2019; see Figures S9 and S8– S19 and Tables S4 and S5). When we weighted errors during the projection period (2001–2019) across outputs to reflect a high confidence in growth data, a moderate confidence in SSB estimates, and a low confidence in yield estimates, the BOM model (i.e. the model with oxygen dependent benthic resource carrying capacity (B), cod benthic occupancy (O), and direct mortality (M) included) performed best (Appendix S5; Figure S11). However, note that direct mortality due to hypoxia (M) does not vary at all with time for both cod and flounder (i.e. *b_i_* = 0 for all models; see equation S.3). Including this component in the BOM model merely adjusts the baseline mortality of these two species. Only benthic food availability (B) and cod occupancy in the benthic habitat (O) vary with oxygen. That is, the interaction of bottom water oxygen on cod “decision” to occupy benthic habitats, combined with hypoxia­driven declines in benthic prey, appears best at explaining observed SSB, yield, and growth of our focal fish species in the Baltic Sea food web.

In general, the BOM model was able to track the observed annual variation in somatic growth of cod throughout the projection period, particularly during the first eight years of life where data are concentrated (Figure 2; see also Figure S15 in Appendix S7). At more advanced ages, the model was prone to over­estimation (Appendix S7; Figure S18), but there are significantly fewer data for these ages, and therefore less certainty regarding actual mean size­at­age. Among­model variation of sizes­at­age is particularly notable at older ages (Figure 2). Growth patterns were relatively static from year to year for other species (see Figures S18 and S19). Flounder scaling parameters were estimated by fitting to a mean predicted von Bertanlanffy growth trajectory across all years, in contrast to the other species for which we had annual estimates for quite a large subset of years. On average, the model was also able to capture the relatively static growth of sprat and herring (see Figures S18 and S19), in addition to the declines in growth of cod.

**Figure 2.**
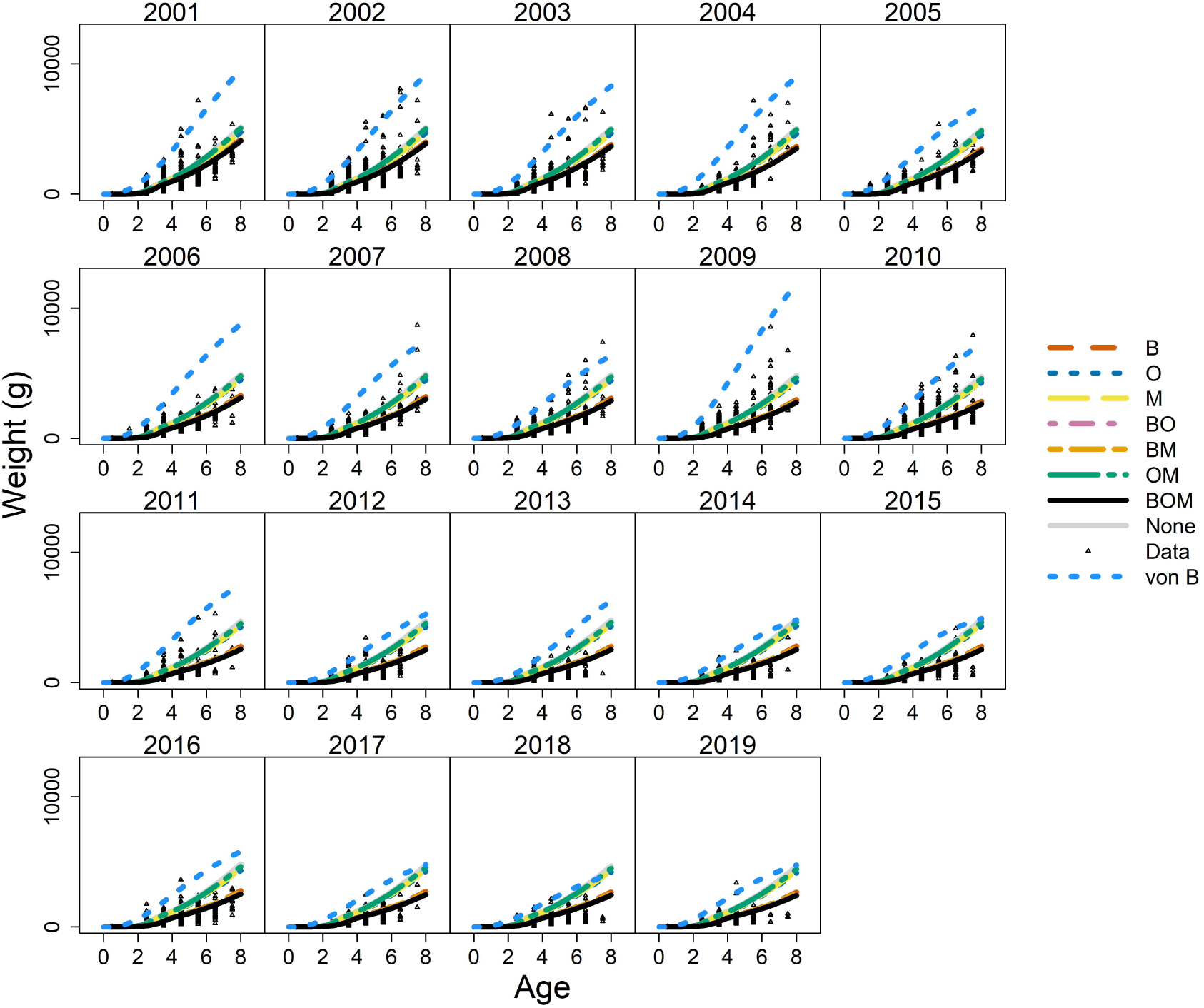
Projections of body growth for cod during the projection period (2001–2019) arising from the best model (BOM i.e. oxygen dependence of benthic food, cod occupancy in the benthic habitat, and mortality), all possible combinations of its components, as well as the model with no oxygen dependence (None). Data are included as small, black triangles. Dotted, light blue lines are von Bertalanffy growth estimates from each year. Note that those lines either with or without oxygen dependence of benthic food availability (B), typically fall nearly beneath or above one another.

While there is variability, most models produced reasonable fits to trends of SSB and yield during the projection period (Figure 3; see also Figure S16 in Appendix S7). Projections of yield were often over­ or under­estimated, but the model did capture the general patterns observed for each species, although less so for flounder (Figure 4; see also Figure S17 in Appendix S7). Differences among all of our models in fits to SSB and yield during the projection period were relatively small (Figures 3 and 4); only those models including scaling of physiological rates (P) without including cod occupancy (O) produced egregious errors (Figures S10 and S11). Therefore, our methods may be less able to distinguish among drivers of observed trends at the population level compared to those at the individual level.

**Figure 3.**
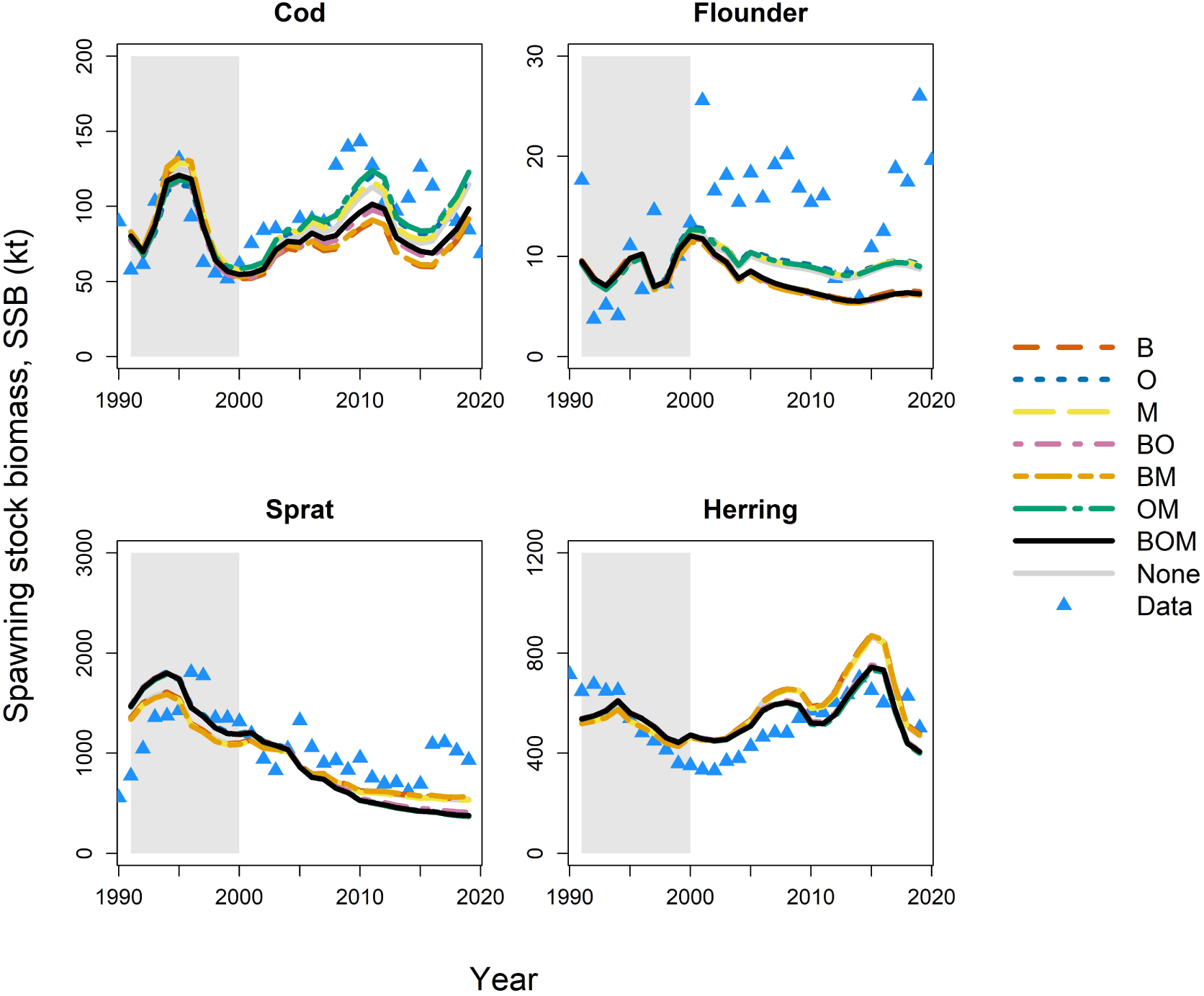
Projections of biomass of mature size classes (SSB) throughout the calibration period (1991–2000; shaded region) and the projection period (2001–2019). Model abbreviations as in Figure 2. Blue triangles are observations (as estimated from stock assessments; see Methods in main text).

**Figure 4.**
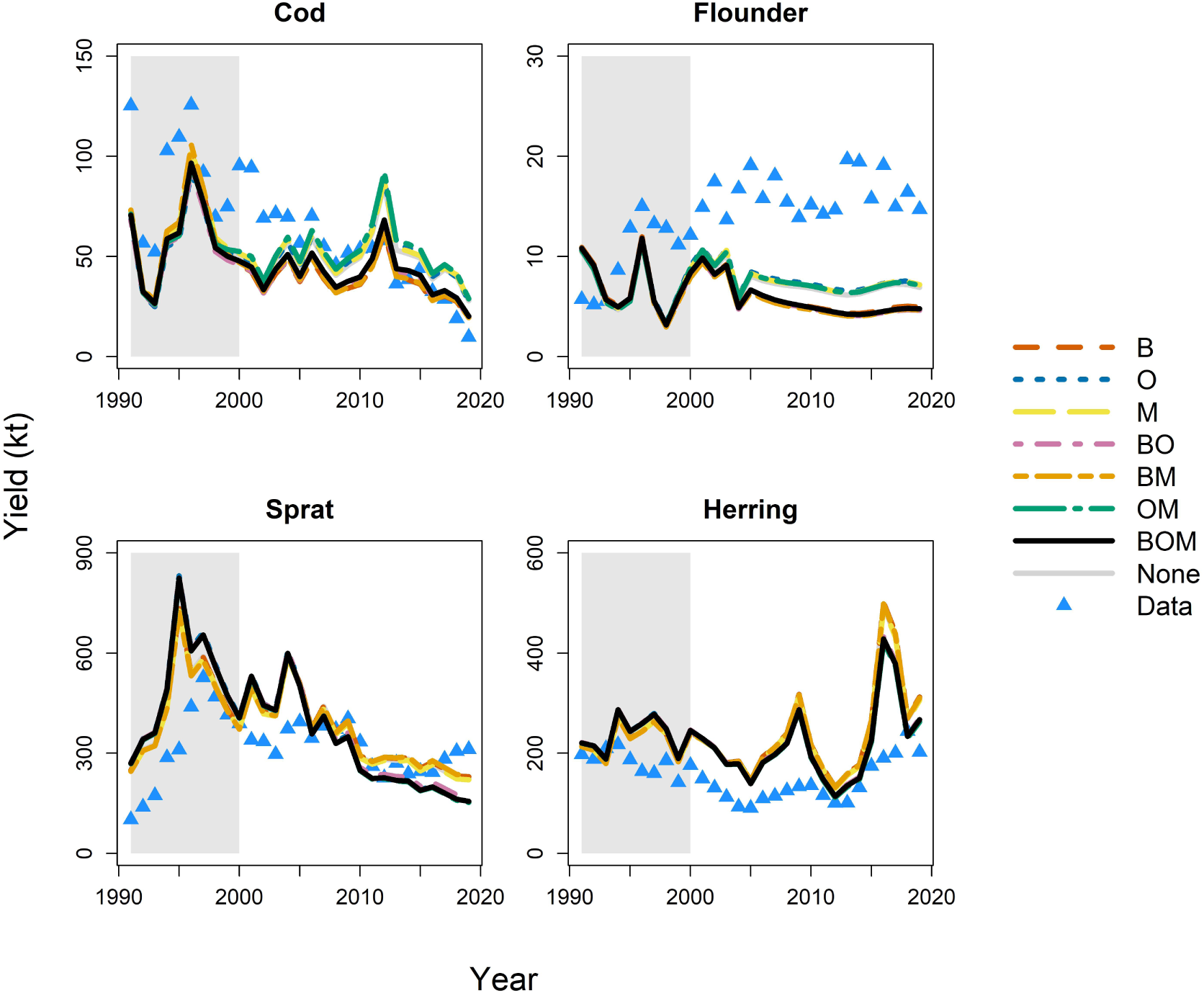
Projections of yield at observed fishing mortality, throughout the calibration period (1991–2000; shaded region) and the projection period (2001–2019). Model abbreviations as in Figure 2. Symbols as in Figure 3.

Results of the sensitivity analysis (Appendix S6) indicate that the best model according to the procedure described above and detailed in Appendix S5 may change depending upon the starting values for scaling parameters. This is most certainly due in part to the the inter­dependency of all model components and output. While the order of the best models does change somewhat from run to run, two things remain consistent. First, models having the lowest relative weighted error always include oxygen dependence of benthic food resources (Appendix S6; Figure S13); they may or may not also include oxygen dependence of occupancy and physiological rates. Second, models with the greatest error always include those with oxygen dependence of physiological rates without oxygen­dependent cod occupancy in the benthic habitat (Appendix S6; Figure S13). That is, benthic food availability and habitat use play a primary role in structuring the results.

### Emergent responses to oxygen scenarios

Comparing the outcome of oxygen scenarios in the best model (BOM) to its components suggests that changing cod occupancy (O) is a compensatory response to declining benthic food availability (B; Figure 5). These results also show that the constant mortality rate has a relatively minor effect on the results. Including oxygen dependence of benthic resource carrying capacity alone (B model) produced unrealistically severe declines in SSB and yield of cod across oxygen scenarios, as well as relatively extreme declines in somatic growth (Figure 5). Those models including both B and O, by contrast, produce less severe, but still significant, declines in individual and population level metrics. The models without oxygen dependence of cod benthic occupancy (O) show no variability of sprat and herring SSB and yield with oxygen (Figure 5). They also tend to produce much greater estimates of biomass and yield of sprat and herring (see Figure S16). This indicates that cod intensify feeding on sprat and herring when relocating to the pelagic habitat, as expected (Figures 6 and 7; see also Table S6). Declines in cod growth, SSB, and yield in the BOM model with declining oxygen (Figure 5) show that moving towards the pelagic habitat does not compensate entirely for the decline in benthic food.

**Figure 5.**
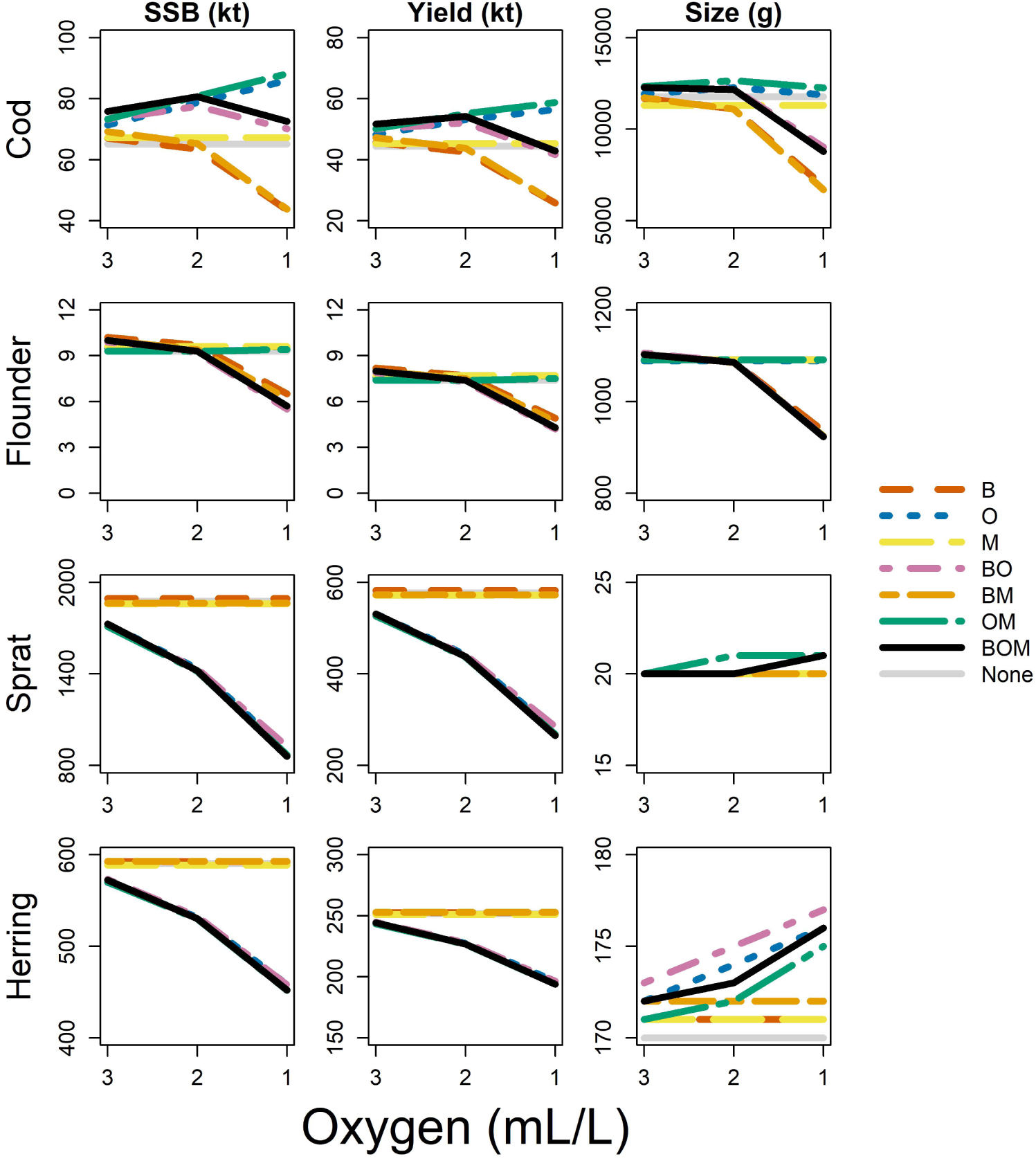
Modeled spawning stock biomass (SSB; kt), yield (kt) at mean fishing mortality ob­ served during the calibration period (1991–2000), and size (g) of individuals at maximum ob­ served age for all species (cod, 15; flounder, 26; sprat, 16; herring, 13) under three oxygen sce­ narios applied in the best model (BOM) and all possible combinations of its components. Model abbreviations as in Figure 2. We also included model results with no oxygen­dependence (None; for which results were identical across oxygen scenarios).

**Figure 6.**
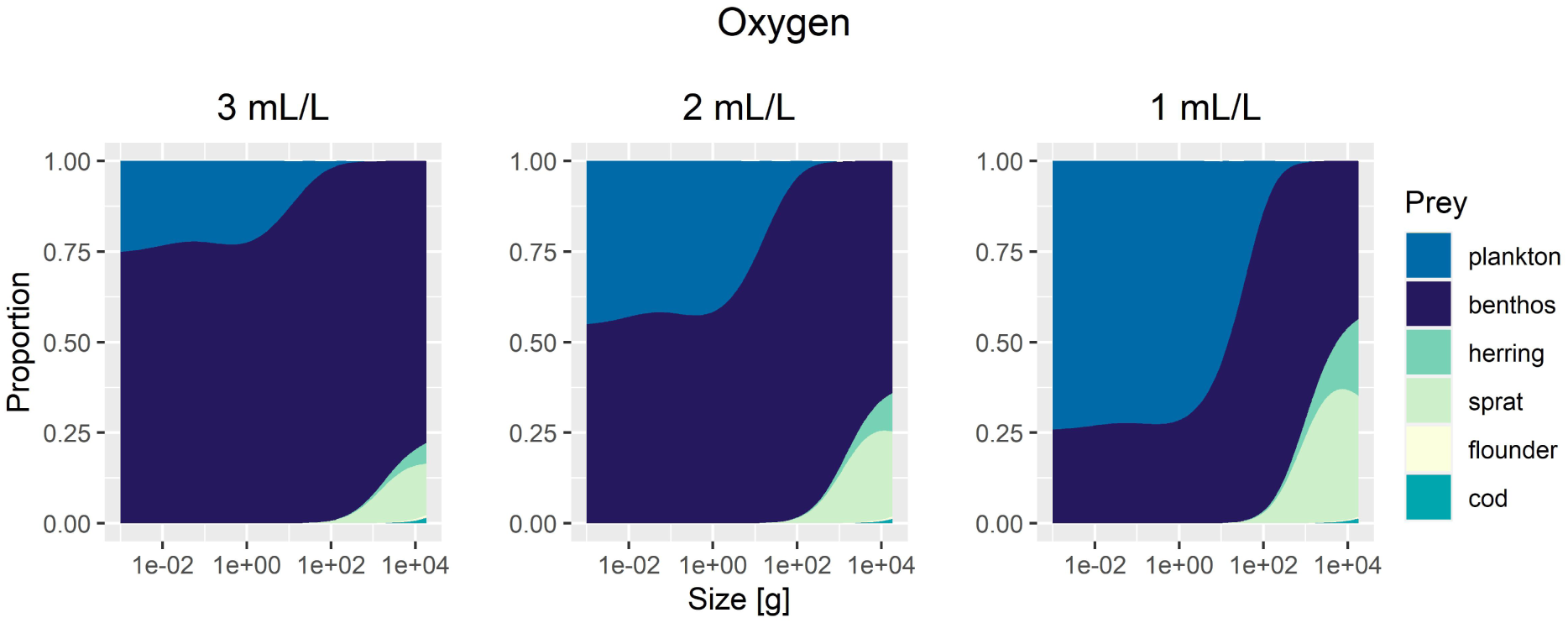
Changes in proportion of each prey item consumed by cod across body sizes as oxygen­ dependent benthic food availability (B) and cod benthic occupancy (O) change with declines in ambient oxygen concentration. The BOM model is used, but note that mortality (M) does not vary with oxygen level.

**Figure 7.**
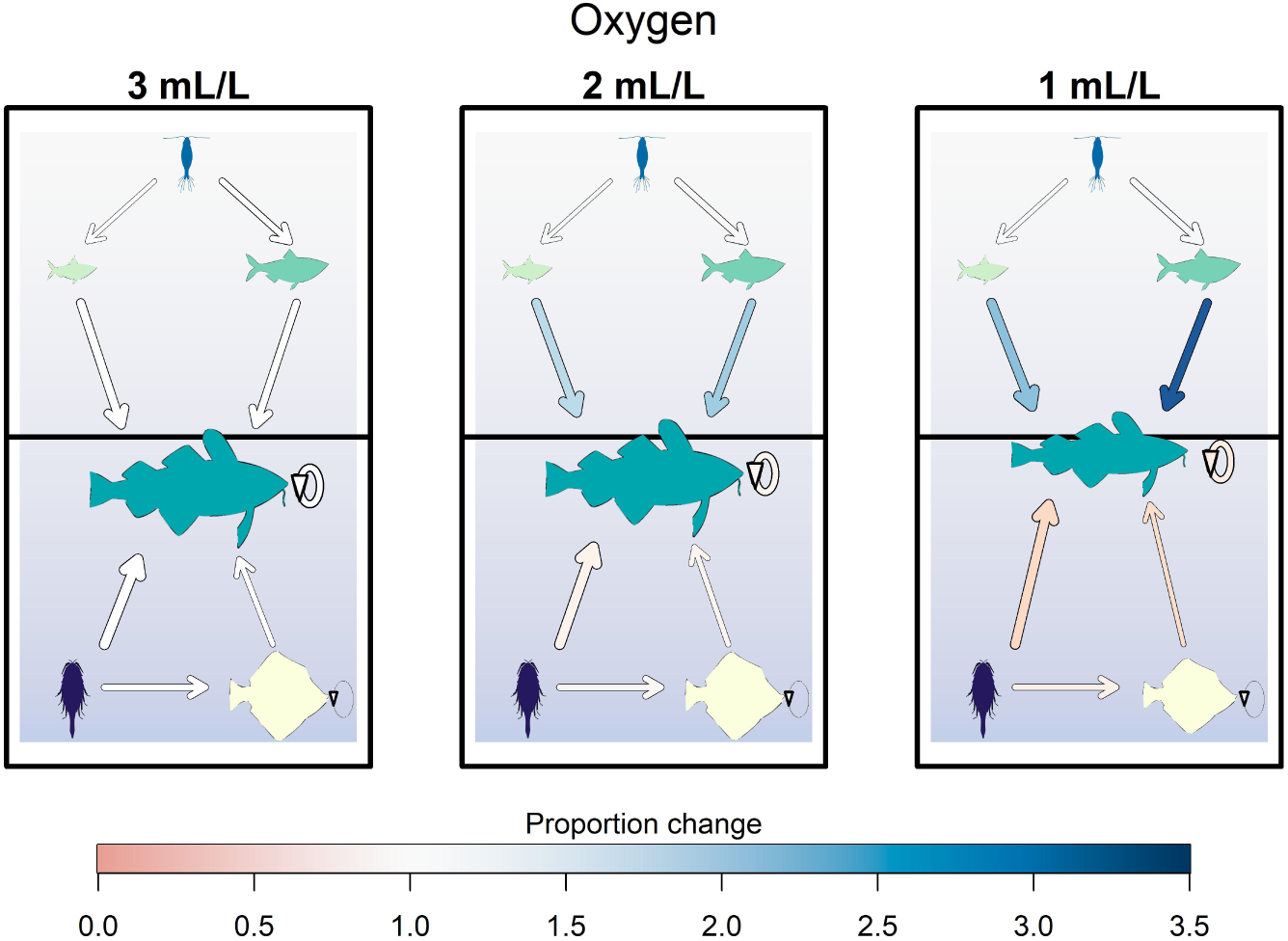
Changes in energy flow among community members as oxygen­dependent benthic food availability (B) and cod occupancy of the benthic habitat (O) change with declines in am­ bient oxygen concentration. The BOM model is used, but note that mortality (M) does not vary with oxygen level. The position of the cod relative to the center black line is calculated from their occupancy of the benthic versus the pelagic habitat, while length and height of the fish drawings reflect declines in body length and weight (g), respectively. The arrows reflect energy flow. Arrow thickness is proportional to the natural log of flow in g⋅year^­1^. Arrow colors represent changes in flow on the linear scale in each scenario relative to those in the 3 mL⋅L^­1^ scenario. On this scale, red shades represent declines, while blues represent increases.

Scenario results also suggest that deoxygenation changes the nature of competitive interactions within and between species (Figure 5). Cod experience greater declines in somatic growth with declining food availability (B model) compared to flounder. This also occurs with both declining food availability and changing cod occupancy (BO model; Figures 5 and 7). When only cod benthic occupancy depends upon oxygen (O model), flounder SSB and yield increase very slightly as cod moves into the pelagic to escape hypoxia, having been released from moderate competition (Figure 5). If benthic food is impacted (B model), then flounder experience marked declines in SSB and yield. While declines in SSB and yield of cod and flounder are of similar magnitude in the B model, the relative decline in body size of cod is more severe than for flounder (Figure 5). This suggests that flounder are the superior competitor within this environment. By contrast, when both benthic food and cod habitat occupancy are oxygen­dependent (BO and BOM model), cod experience only moderate declines in SSB and yield compared to flounder (Figure 5). Low food availability limits flounder in our model (Table S5), but unlike for cod, the limitation decreases flounder carrying capacity and does not affect growth as much. Given the paucity of reliable information on the Baltic flounder populations, it is generally unknown how they are faring in the Central Baltic Sea as a whole. There is relatively more information for sprat and herring, and here results show that SSB and yield of both sprat and herring decline (Figures 5 and 7). As there are no direct effects of benthic deoxygenation on these pelagic species, this is caused by intensifying predation as cod avoid hypoxic bottom waters. Declines are more severe for sprat, as herring can largely outgrow their vulnerability to cod predation (Figure 6). This releases the pelagic fishes from competition somewhat, resulting in further growth (Figure 5). We stress that changes in sprat and herring growth are very minor, as their competition for pelagic food sources is weak in the model. This is primarily caused by setting pelagic resource carrying capacity high in order to reflect observed cod diet, as well as pelagic fish SSB (see Appendix S5). Nonetheless, model results show that the availability of benthic resources and changing vertical habitat use of cod reorder interactions both in the benthic and the pelagic habitats.

Finally, declining benthic food availability and increasing use of the pelagic habitat shifts energy flow to cod away from benthic and toward pelagic prey (Figures 6 and 7). As cod move up in the water column in response to low oxygen, feeding on flounder and benthic resources declines overall, while feeding on sprat and herring increases (Figure 7). Notably, these pelagic fish increase from 25% to over 50% of the diet of large cod when oxygen levels decline from 3 to 1 mL⋅L^­1^, with the remainder of their diet consisting of benthic resources. As benthic resources decline with deoxygenation, small cod feed increasingly on plankton, as they are too small to feed significantly on the larger fish prey in the pelagic habitat (Figure 7).

## Discussion

Marine hypoxia can have severe and complex consequences for fish and fisheries (Rose et al. 2019). Results from our modeling study suggest food availability in deoxygenated habitats, as well as the use of oxygenated refugia, may be primary drivers of emergent patterns in fish populations and communities. As the severity of hypoxia increases, community size structure shifts among benthic and benthopelagic predators to favor smaller individuals, and consumption patterns may change. This may be driven both by changes in size structure, and by changes in habitat use. A mismatch in prey availability relative to the needs of predators could limit the ability of changing habitat use to compensate for declines in somatic growth of benthopelagic species. All of these impacts result in a change in the flow of energy among species in the food web. These insights into potential mechanisms by which the effects of hypoxia on individuals may scale to affect whole communities can provide valuable context for future studies and for the management of fish populations.

We have attempted to answer three questions on drivers of food web responses to deoxygenation with our analysis: first, which individual­level effects of marine deoxygenation are critical to capturing higher level dynamics in hypoxic systems? Our analysis of alternative oxygen­ dependent pathways in a size­structured food web model suggests that a decrease in the availability of benthic prey is essential for explaining observed responses (spawning stock biomass or SSB, fisheries yield, and somatic growth) to deoxygenation. Model results support a decline in the use of benthic habitats and subsequent changes in predatory fish feeding from benthic to pelagic prey may also be critical. Our sensitivity analysis suggests, however, that changes in habitat use may not be as critical as declines in benthic food resources, and that oxygen dependence of physiological rates may also be important (Appendix S6; Figure S13). Observations in the Baltic Sea do suggest a decline in benthic food availability (Karlson et al. 2020; Neuenfeldt et al. 2020). *Saduria entomon*, a major food source for Baltic cod, is limited by oxygen concentrations below about 1.5 mL⋅L^­1^ (Haahtela 1990). Declining benthic food availability (B) remains in all of the best models for the projection period whether or not they also include oxygen dependence of cod benthic occupancy and physiological rates (Appendix S5; Figure S11). Accounting for feeding interactions and their dependence on oxygen conditions thus appears to be essential for understanding emergent food web dynamics to marine deoxygenation.

Second, we asked: how do critical individual­level impacts of deoxygenation interact with one another to result in community­level responses? Examining the ability of all models to explain emergent dynamics in spawning stock biomass (SSB), yield, and fish body growth in the four main commercially exploited fish stocks in the Central Baltic Sea reveals that interactions between mechanisms are important. The inclusion of some level of oxygen­driven change in feeding, whether through a decline in the biomass of benthic invertebrates (B) and in cod occupancy of benthic habitats (O), through a decline in feeding activity, assimilation efficiency, and egg survival (P), or all three of these, all appear in superior models (Appendix S5; Figure S11). The former two mechanisms are supported by observations of a decline in cod feeding on benthos (Neuenfeldt et al. 2020) and in benthic occupancy of cod (Casini et al. 2019). The latter is supported by observations of reduced appetite in hypoxic conditions (Pichavant et al. 2001; Chabot and Claireaux 2008), as well as reduced assimilation efficiency (Wang et al. 2009) and egg survival (Köster et al. 2005) of cod. Of note is that model results suggest both of these mechanisms (B and P) likely operate alongside altered cod occupancy of the benthic habitat (O) for best model performance (Appendix S5; Figure S11), though the implications differ. For models with physiological oxygen dependence (P), this implies that, given observed dynamics, there must be an opportunity for cod to escape from degraded environments. For the best model (BOM), there must be an alternative food source (i.e. access to pelagic prey fish). The importance of dynamic occupancy of the benthic habitat in representing cod population dynamics and somatic growth is supported by the literature. Shifts in the relative use of benthic habitat in cod have been observed in the Baltic Sea (Casini et al. 2019), in addition to shifts in cod diets towards clupeids (Pachur and Horbowy 2013). Furthermore, it has been observed that diets of cod are driven at least in part by differences in vertical overlap caused by the dynamics of salinity and oxygen in bottom waters (Neuenfeldt and Beyer 2006; Pachur and Horbowy 2013). Though changing habitat use may serve as a compensatory mechanism, it is clearly unable to compensate fully for the loss of benthic habitat and benthic prey in our case study. Given this body of evidence, in addition to our own results, accounting both for the negative effects of deoxygenation as well as potential compensatory mechanisms allows for a more complete understanding of both direct and indirect effects of deoxygenation that emerge in communities of interacting species in connected habitats.

Lastly we asked: how might we expect declining oxygen levels to affect fish population and community metrics? Scenario results suggest that, as oxygen declines, most population metrics may also be expected to decline. That said, at moderate oxygen levels (2 mL⋅L^­1^), cod SSB and yield experienced modest increases on the order of 4­5% relative to more ideal oxygen conditions (3 mL⋅L^­1^; Figure 5). This is due to the expanded access to an additional food source (clupeids). Recall that the model assumes constant areal overlap of fish species, and oxygen as the only factor motivating cod movement between habitats. In reality, the horizontal overlap of cod and clupeid prey fish has decreased in the past three decades, with cod concentrated in the southern Baltic Sea and clupeids in the northern areas (Casini et al. 2011; Casini et al. 2016). Note that, while our model tracks declines in cod benthic occupancy during the calibration period, it fails to account for more recent increases (Appendix S5; Figure S9). Therefore, this projected increase in overlap and moderate increase in SSB may be an artefact of model assumptions. However, at minimum observed oxygen levels (1 mL⋅L^­1^), both SSB and yield of cod decline, with the latter declining more severely. This is due to stunted growth. Empirical evidence links reduced growth of cod to increasing exposure to hypoxia (Limburg and Casini 2018; Casini et al. 2021). While selectivity is constant across scenarios, the yield becomes dominated by smaller individuals as oxygen declines. All other species experience declines in SSB and yield across deoxygenation scenarios. The response in body growth depends on species. Growth of pelagic clupeids is largely insensitive to benthic deoxygenation relative to the declines in growth of the benthic species. Both cod and flounder experience declines in growth, with that of the former being more severe. Herring and sprat do experience very slight increases in growth as cod rise in the water column and prey upon the sprat and the herring, more so the former than the latter. Changes in sprat and herring are particularly interesting, given that their responses to deoxygenation are driven entirely by interactions with cod as they move through the water column. Thus, all species experience changes at the population level across scenarios, though it varies greatly by species. Indirect effects arise, particularly with reference to clupeids, as food web interactions change in response to declining oxygen.

Energy flow through the food web also changes in response to marine deoxygenation, primarily as benthopelagic cod move vertically in the water column. This mediates indirect effects of hypoxia. Our model results suggest that size and species composition of prey are altered as cod escape hypoxic conditions. As benthic food declines, the proportion of pelagic food in the diet of small cod increases, shifting the overall size distribution of cod prey towards smaller items in the early years of life. Similar shifts in size and species composition of prey sources of cod are observed in the Baltic Sea (Pihl 1994; Haase et al. 2020; Neuenfeldt et al. 2020). Pelagic fishes become more important in the diet of cod as they grow (Griffiths et al. 2017). According to our results, oxygen declines may intensify this shift. Overall, the consequences of the interaction between declining benthic food and changing habitat use is a reorganization of energy flow away from benthic and towards pelagic prey among benthopelagic species in the food web, and dramatic changes in the community size structure. Most notable, this includes a sharp decline in large predators. It is unlikely that pelagic forage fish are otherwise unaffected by benthic oxygen, as they may use deeper habitats as feeding grounds (Möllmann et al. 2004; Ludsin et al. 2009), or refugia from warming waters, thus altering food web structure (Tunney et al. 2014). That said, it is clear that strong indirect effects can arise in systems driven by deoxygenation. Community species that are concentrated within oxygenated refugia may suffer from increased predation, while species escaping deoxygenated habitats may suffer from a mismatch in dietary needs relative to availability.

We made several simplifying assumptions in our model which explicitly defy observations. For example, we used a static representation of size at maturation during the calibration period for all species, whereas evidence suggests that size at maturation has declined for Baltic cod (Vainikka et al. 2009; Köster et al. 2017). We also assumed a constant temperature. There is strong evolutionary pressure on both thermal and hypoxic limits for fishes (Deutsch et al. 2020). Sustained increases in both temperature, and in the frequency and severity of hypoxic events, may thus result in rapid evolutionary changes. In general, we would expect there to be strong interactions between declining oxygen and warming, as temperature affects both oxygen supply and demand (Roman et al. 2019). It is known that these two stressors can act synergistically (Ekström et al. 2021), though the system­specific response will depend on species identity, species interactions, and community plasticity (McBryan et al. 2013). Which mechanisms may drive this interaction is generally unknown, and currently unexplored in our model structure. We also ignored individual variability in the model outside of differences in species and maximum body size. Neither did we account for sex. In reality, differences amongst individuals, including those dependent upon sex, may cause variation in both space and time. For example, some fish with higher metabolic rates tend to be more active and more willing to take risks in hypoxic environments (Killen et al. 2012). Depending on the outcome, and given that metabolic rate is an heritable trait (Maciak and Konarzewski 2010), spatiotemporal variability could potentially be driven by changes in selective forces that are in turn driving the distribution of metabolic rates among the population. Our conclusions may only hold if the variation and mechanisms described above and elsewhere are negligible. However, even if hypoxia acts through vulnerability to increasing temperature, incidence of parasitism and disease, or other drivers, our model is general enough to capture declines in community metrics due to implicit drivers. One may include individual­level impacts of alternative drivers both in addition to, and in place of, the drivers included here to determine whether they are important.

Do our conclusions hold if we broaden our perspective to other systems? In the Gulf of Mexico, pelagic prey fishes move into alternative habitats where prey is scarce and the risk of predation is high in response to changes in habitat quality wrought by hypoxia and warming (Zhang et al. 2014). In Lake Erie, demersal Yellow Perch (*Perca flavescens*) move either horizontally or vertically in response to hypoxic bottom waters (Roberts et al. 2009). Though they continue to dive into degraded waters for their benthic prey (Behrens et al. 2012), diet composition still shifts towards pelagic sources (Roberts et al. 2009). Differences in susceptibility to hypoxia among species in our simplified Baltic Sea food web serve also to reorganize energy flow. This has been observed for example in the Neuse River Estuary in North Carolina, where the strength of the escape response and therefore changes in predation pressure vary by species (Bell and Eggleston 2005). Looking at lower trophic levels in the Chesapeake Bay, escape response of benthic invertebrates can significantly alter the flow of energy from the sediment to demersal predators (Pihl et al. 1992). Differential responses across species in other community assemblages will likely result in different responses, though even species resistant to hypoxia may still be observed to exhibit similar responses (e.g. diet shifts in Atlantic croaker; Mohan and Walther 2016; Steube et al. 2021) Therefore, there seems to be broad support for our conclusions across systems.

Overall, our study suggests that declining benthic food availability and escape of demersal predators towards pelagic habitats are important pathways by which persistent and severe hypoxia affects community dynamics. These ecological consequences can help to inform forecasts of biomass, yield, and growth of component species as hypoxia stagnates, worsens, or, more optimistically, as environmental conditions improve. Failing to account for the potential impacts of deoxygenation may lead to erroneous stock assessments and poor management advice (Rose et al. 2019). For example, ignoring changes in natural mortality caused by increased predation by displaced benthopelagic predators on pelagic prey fishes could introduce significant bias into biomass estimates and therefore into recommended quotas (Clark 1999). Resultant changes in community size structure, and uncertainty therein introduced by developing stressors like declining oxygen and rising temperatures, are necessary components of projections used in the decision­making process (Reum et al. 2020). While our approach is only a first step towards partitioning observed fish community dynamics among environmental impacts, it may serve as a useful baseline with which to rigorously evaluate various precautionary approaches to fishing in communities burdened by hypoxic waters.

## Conflict of Interest Statement

The authors declare there are no conflicts of interest.

## Supporting information

LaTeX files

## Acknowledgements

We are grateful to Gustav Delius and Ken Andersen for discussions regarding proper use of **mizer**. We would also like to thank Max Lindmark for his guidance on using **mizer** to investigate environmental effects on food webs. The entire SPECTRE reading and discussion group encouraged our use and understanding of size spectrum models more generally. This work was supported by funding for Project Breathless (NSF OCE1923965; to KL, MC and AG). Additional funding was provided by the Swedish Research Council Formas (grant no. 2018­00775 to MC).

---

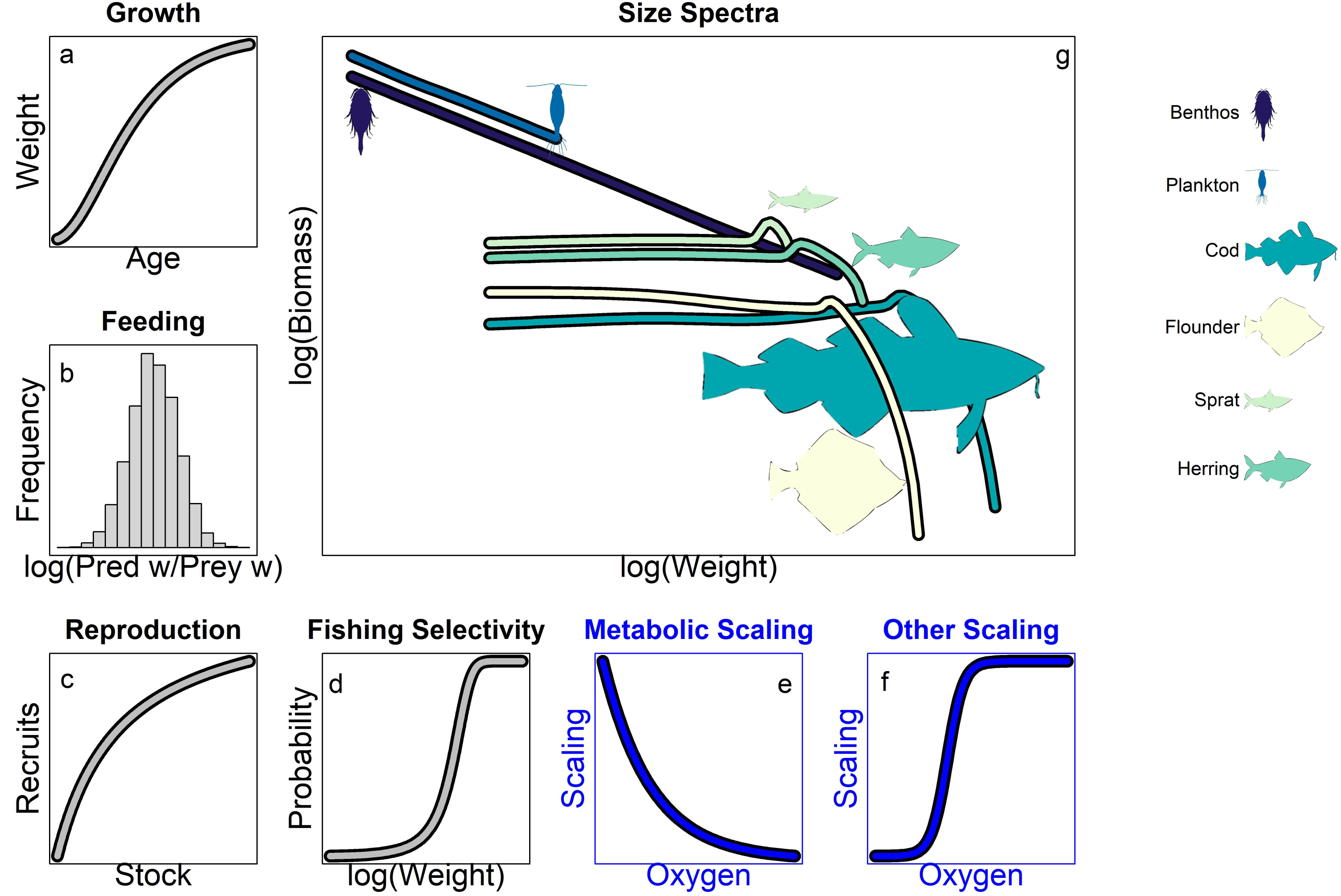

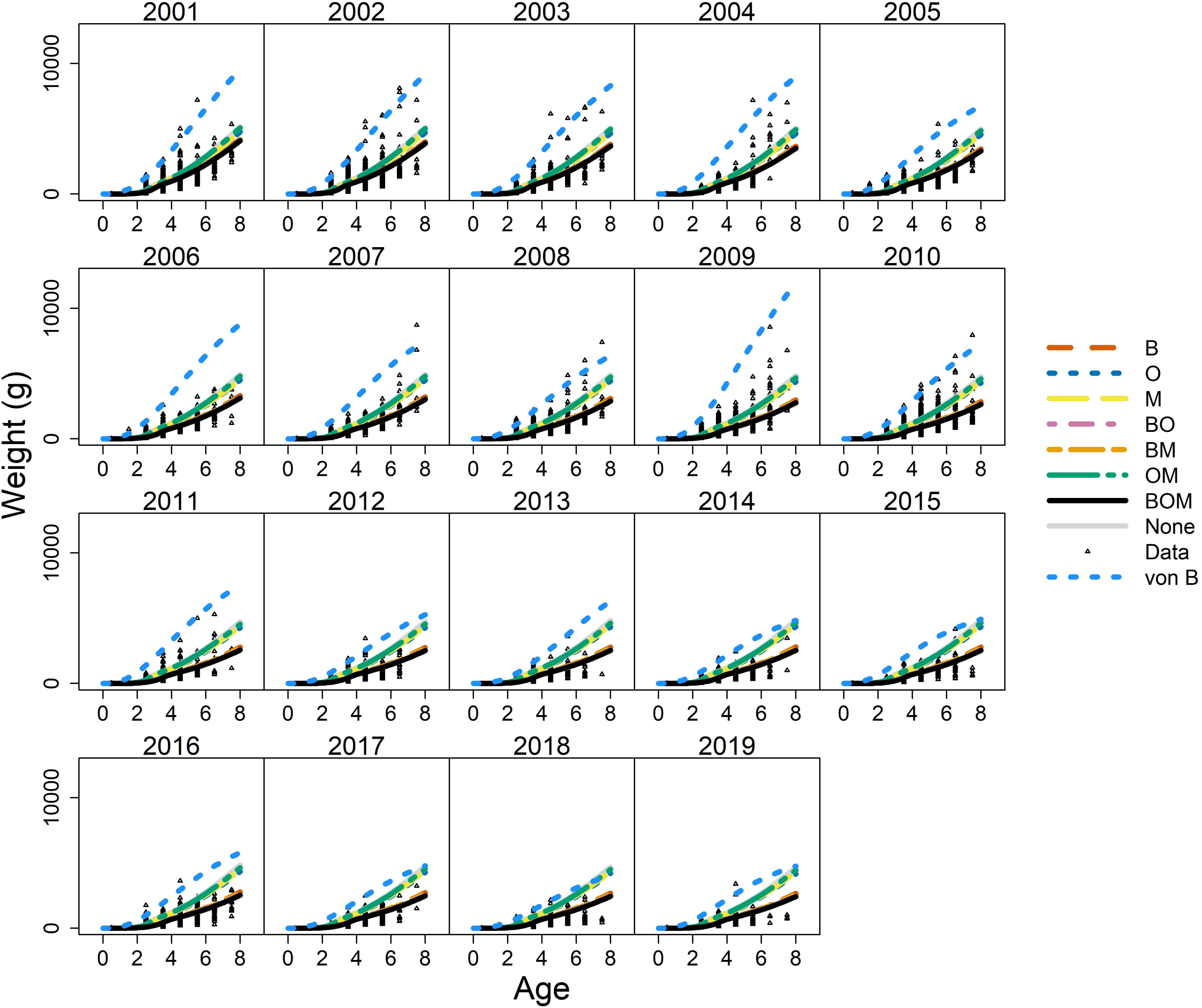

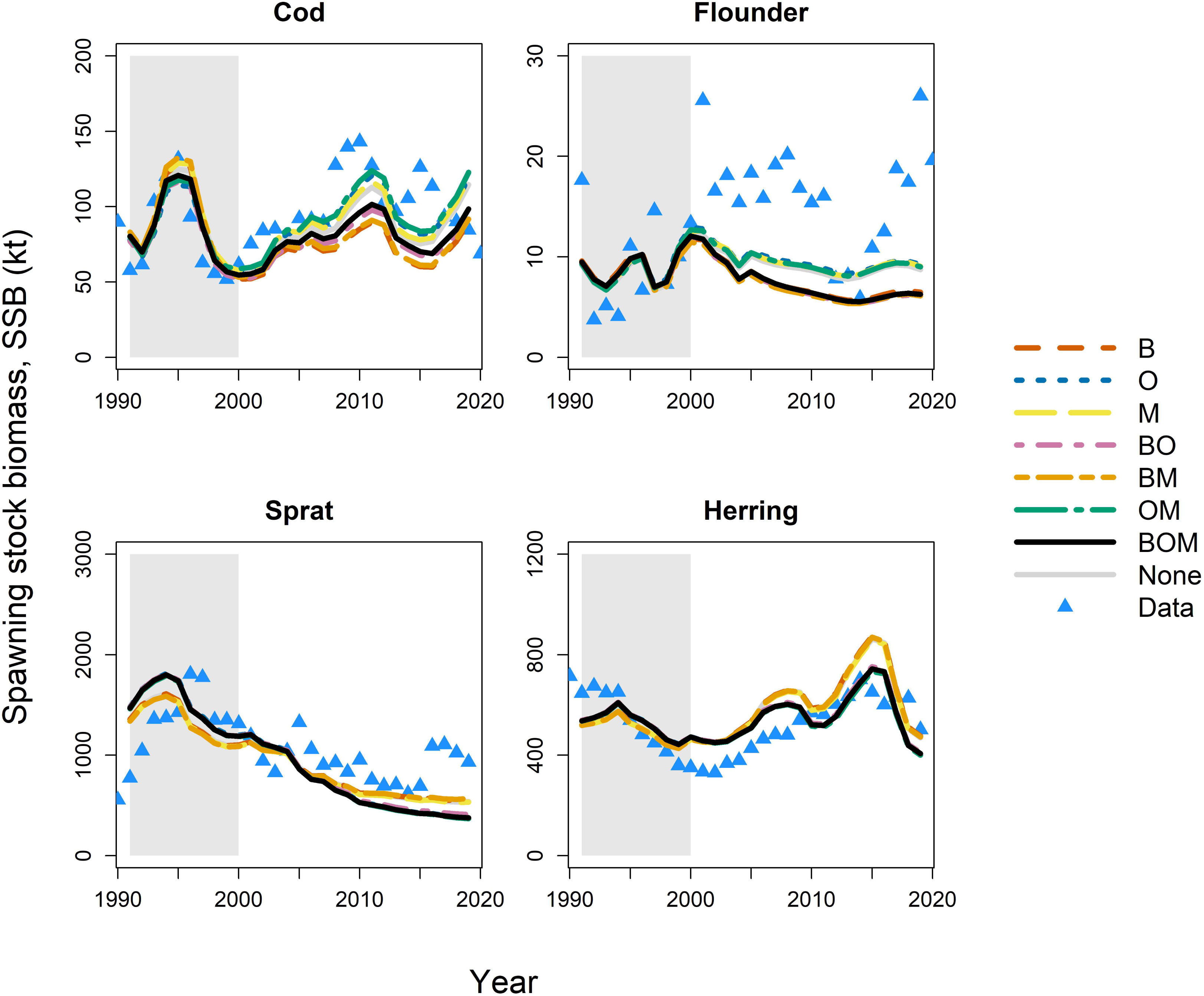

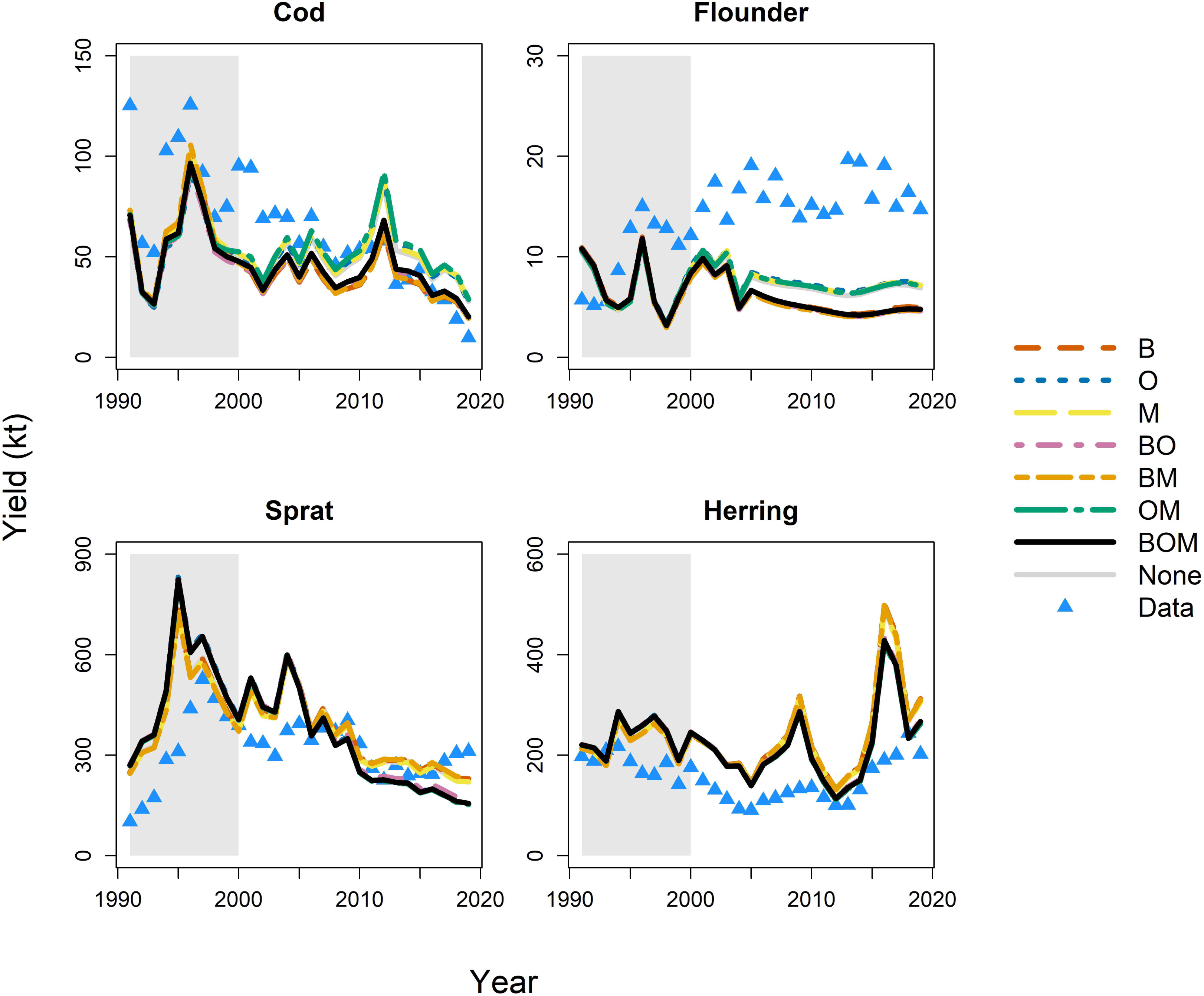

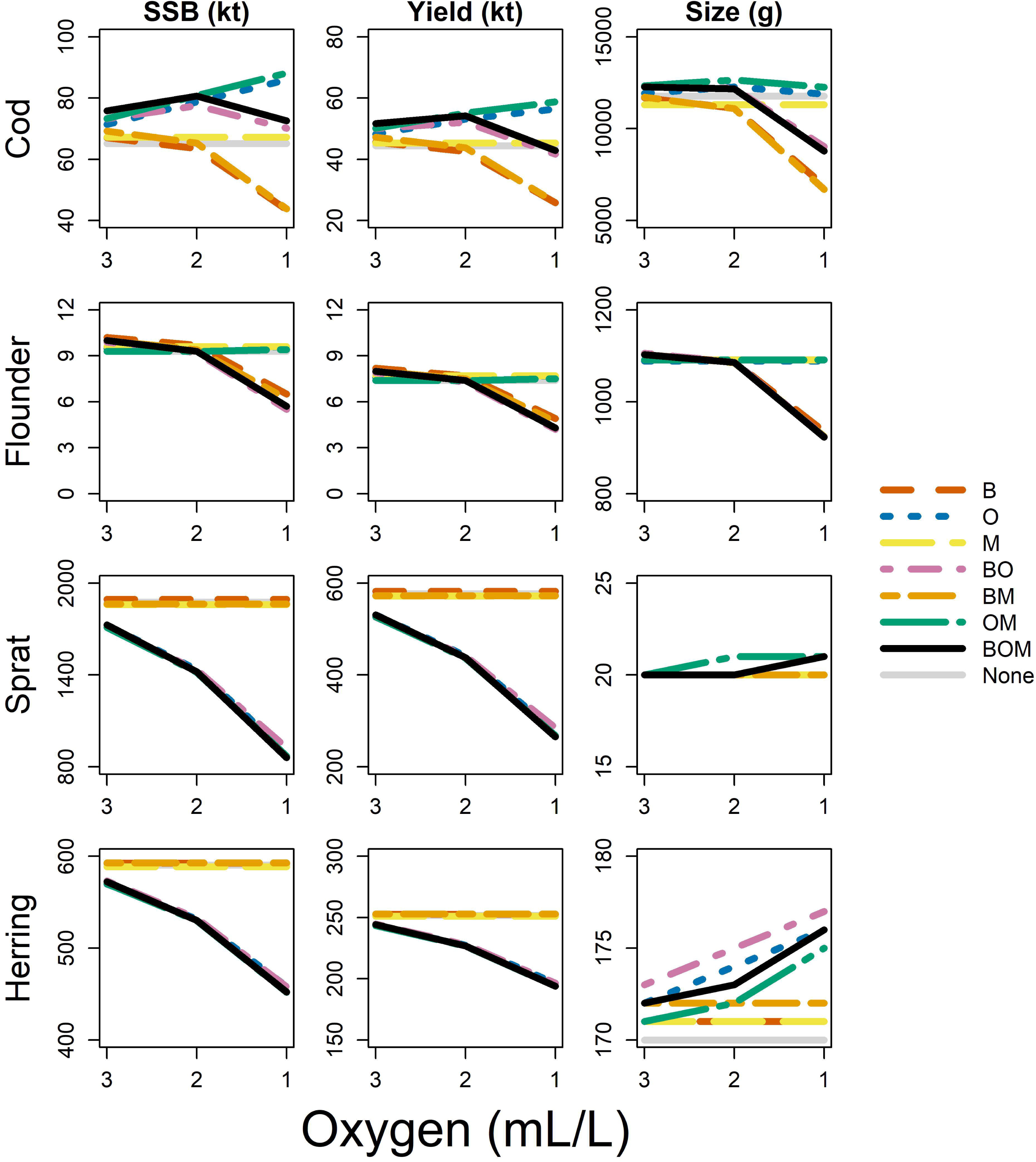

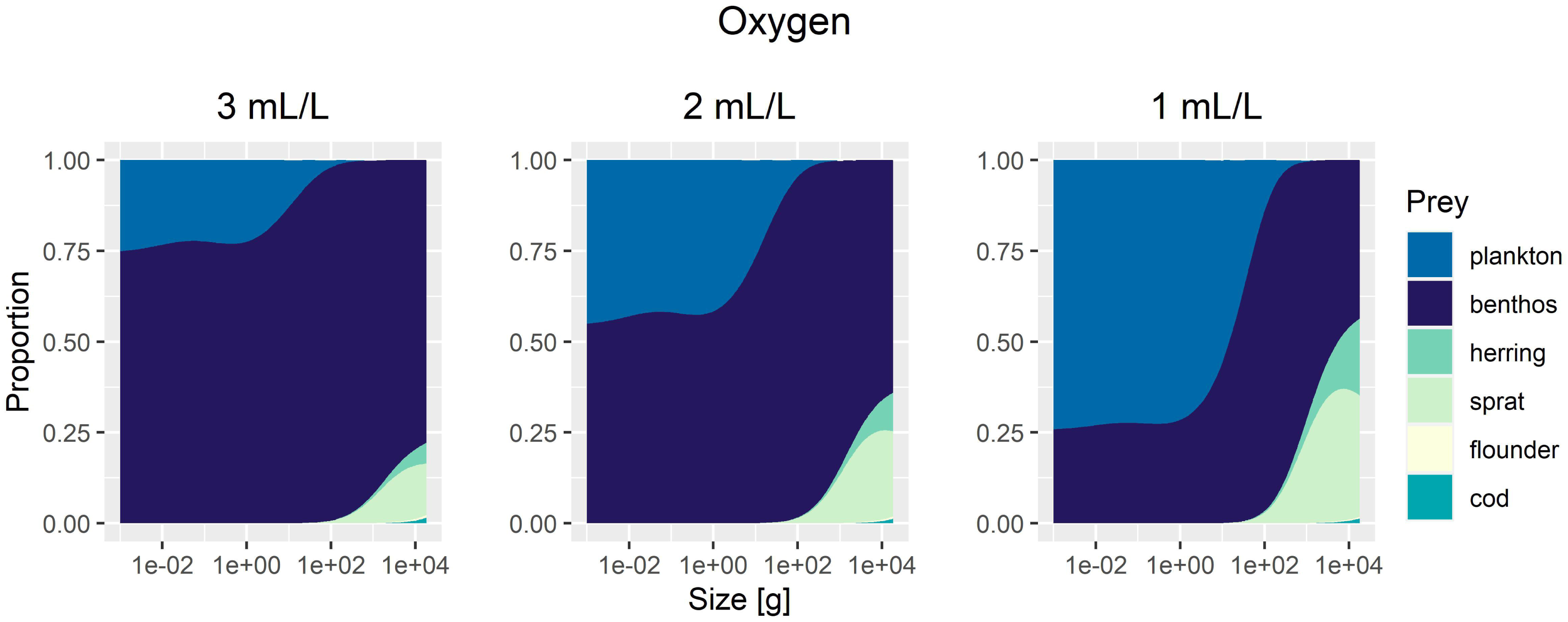

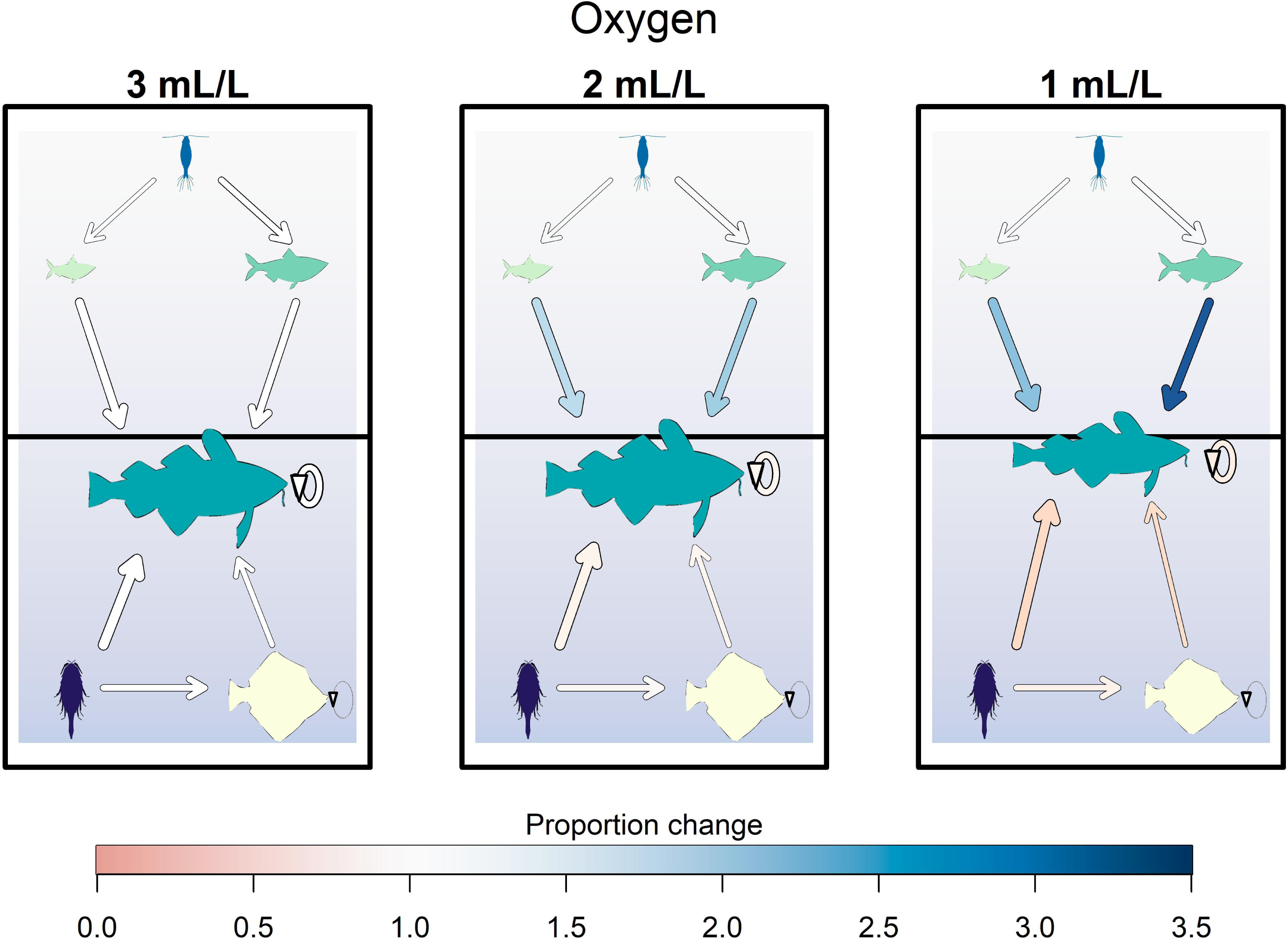

## Notes

https://github.com/epduskey/hypoxiaSSM

### Competing Interest Statement

The authors have declared no competing interest.

## References

1. Andersen, J. H., Carstensen, J., Conley, D. J., Dromph, K., Fleming­Lehtinen, V., Gustafsson, B. G., Josefson, A. B., Norkko, A., Villnäs, A., and Murray, C. (2017). “Long­term temporal and spatial trends in eutrophication status of the Baltic Sea”. In: Biological Reviews 92.1, pp. 135–149. DOI: 10.1111/brv.12221.

2. Andersen, K. H., Jacobsen, N. S., and Farnsworth, K. D. (2016). “The theoretical foundations for size spectrum models of fish communities”. In: Canadian Journal of Fisheries and Aquatic Sciences 73.4, pp. 575–588. DOI: 10.1139/cjfas-2015-0230.

3. Behrens, J. W., Axelsson, M., Neuenfeldt, S., and Seth, H. (2012). “Effects of Hypoxic Exposure During Feeding on SDA and Postprandial Cardiovascular Physiology in the Atlantic cod, Gadus morhua”. In: PLoS ONE 7.9, e46227. DOI: 10.1371/journal.pone.0046227.

4. Bell, G. and Eggleston, D. (2005). “Species­specific avoidance responses by blue crabs and fish to chronic and episodic hypoxia”. In: Marine Biology 146.4, pp. 761–770. DOI: 10.1007/s00227-004-1483-7.

5. Bergman, J. N., Bennett, J. R., Binley, A. D., Cooke, S. J., Fyson, V., Hlina, B. L., Reid, C. H., Vala, M. A., and Madliger, C. L. (2019). “Scaling from individual physiological measures to population­level demographic change: case studies and future directions for conservation management”. In: Biological Conservation 238, p. 108242. DOI: 10.1016/j.biocon.2019.108242.

6. Blanchard, J. L., Heneghan, R. F., Everett, J. D., Trebilco, R., and Richardson, A. J. (2017). “From Bacteria to Whales: using Functional Size Spectra to Model Marine Ecosystems”. In: Trends in Ecology & Evolution 32.3, pp. 174–186. DOI: 10.1016/j.tree.2016.12.003.

7. Breitburg, D., Levin, L. A., Oschlies, A., Grégoire, M., Chavez, F. P., Conley, D. J., Garçon, V., Gilbert, D., Gutiérrez, D., Isensee, K., Jacinto, G. S., Limburg, K. E., Montes, I., Naqvi, S., Pitcher, G. C., Rabalais, N. N., Roman, M. R., Rose, K. A., Seibel, B. A., Telszewski, M., Yasuhara, M., and Zhang, J. (2018). “Declining oxygen in the global ocean and coastal waters”. In: Science 359.6371, eaam7240. DOI: 10.1126/science.aam7240.

8. Breitburg, D. L., Adamack, A., Rose, K. A., Kolesar, S. E., Decker, B., Purcell, J. E., Keister, J. E., and Cowan, J. H. (2003). “The pattern and influence of low dissolved oxygen in the Patuxent River, a seasonally hypoxic estuary”. In: Estuaries 26.2, pp. 280–297. DOI: 10.1007/BF02695967.

9. Breitburg, D. L., Loher, T., Pacey, C. A., and Gerstein, A. (1997). “VARYING EFFECTS OF LOW DISSOLVED OXYGEN ON TROPHIC INTERACTIONS IN AN ESTUARINE FOOD WEB”. In: Ecological Monographs 67.4, pp. 489–507. DOI: 10.1890/0012-9615(1997)067[0489:VEOLDO]2.0.CO;2.

10. Breitburg, D. L., Rose, K. A., and Cowan Jr, J. H. (1999). “Linking water quality to larval survival: predation mortality of fish larvae in an oxygen­stratified water column”. In: Marine Ecology Progress Series 178, pp. 39–54. DOI: 10.3354/meps178039.

11. Brewer, P. G. and Peltzer, E. T. (2016). “Ocean chemistry, ocean warming, and emerging hypoxia: Commentary”. In: Journal of Geophysical Research: Oceans 121.5, pp. 3659–3667. DOI: 10.1002/2016JC011651.

12. Brown, J. H., Gillooly, J. F., Allen, A. P., Savage, V. M., and West, G. B. (2004). “TOWARD A METABOLIC THEORY OF ECOLOGY”. In: Ecology 85.7, pp. 1771–1789. DOI: 10.1890/03-9000.

13. Carstensen, J., Andersen, J. H., Gustafsson, B. G., and Conley, D. J. (2014). “Deoxygenation of the Baltic Sea during the last century”. In: Proceedings of the National Academy of Sciences 111.15, pp. 5628–5633. DOI: 10.1073/pnas.1323156111.

14. Casini, M., Hansson, M., Orio, A., and Limburg, K. (2021). “Changes in population depth distribution and oxygen stratification are involved in the current low condition of the eastern Baltic Sea cod (Gadus morhua)”. In: Biogeosciences 18.4, pp. 1321–1331. DOI: 10.5194/bg-18-1321-2021.

15. Casini, M., Käll, F., Hansson, M., Plikshs, M., Baranova, T., Karlsson, O., Lundström, K., Neuenfeldt, S., Gårdmark, A., and Hjelm, J. (2016). “Hypoxic areas, density­dependence and food limitation drive the body condition of a heavily exploited marine fish predator”. In: Royal Society open science 3.160416. DOI: 10.1098/rsos.160416.

16. Casini, M., Kornilovs, G., Cardinale, M., Möllmann, C., Grygiel, W., Jonsson, P., Raid, T., Flinkman, J., and Feldman, V. (2011). “Spatial and temporal density dependence regulates the condition of central Baltic Sea clupeids: compelling evidence using an extensive international acoustic survey”. In: Population Ecology 53.4, pp. 511–523.

17. Casini, M., Tian, H., Hansson, M., Grygiel, W., Strods, G., Statkus, R., Sepp, E., Gröhsler, T., Orio, A., and Larson, N. (2019). “Spatio­temporal dynamics and behavioural ecology of a “demersal” fish population as detected using research survey pelagic trawl catches: the Eastern Baltic Sea cod (Gadus morhua)”. In: ICES Journal of Marine Science 76.6, pp. 1591–1600. DOI: 10.1093/icesjms/fsz016.

18. Chabot, D. and Claireaux, G. (2008). “Environmental hypoxia as a metabolic constraint on fish: the case of Atlantic cod, Gadus morhua”. In: Marine Pollution Bulletin 57.6, pp. 287–294. DOI: 10.1016/j.marpolbul.2008.04.001.

19. Chu, J. W. and Gale, K. S. (2017). “Ecophysiological limits to aerobic metabolism in hypoxia determine epibenthic distributions and energy sequestration in the northeast Pacific ocean”. In: Limnology and Oceanography 62.1, pp. 59–74. DOI: 10.1002/lno.10370.

20. Clark, W. G. (1999). “Effects of an erroneous natural mortality rate on a simple age­structured stock assessment”. In: Canadian Journal of Fisheries and Aquatic Sciences 56.10, pp. 1721–1731. DOI: 10.1139/f99-085.

21. Cooke, R. M., Wittmann, M. E., Lodge, D. M., Rothlisberger, J. D., Rutherford, E. S., Zhang, H., and Mason, D. M. (2014). “Out­of­Sample Validation for Structured Expert Judgment of Asian Carp Establishment in Lake Erie”. In: Integrated Environmental Assessment and Management 10.4, pp. 522–528. DOI: 10.1002/ieam.1559.

22. Deutsch, C., Penn, J. L., and Seibel, B. (2020). “Metabolic trait diversity shapes marine biogeography”. In: Nature 585, pp. 557–562. DOI: 10.1038/s41586-020-2721-y.

23. Diaz, R. J. and Breitburg, D. L. (2009). “The Hypoxic Environment”. In: Fish Physiology. Vol. 27. Elsevier, pp. 1–23. DOI: 10.1016/S1546-5098(08)00001-0.

24. Domenici, P., Herbert, N., Lefrançois, C., Steffensen, J., and McKenzie, D. (2013). “The Effect of Hypoxia on Fish Swimming Performance and Behaviour”. In: Swimming Physiology of Fish. Ed. by A. P. Palstra and J. V. Planas. Berlin, Heidelberg: Springer, pp. 129–159. DOI: 10.1007/978-3-642-31049-2_6.

25. Dwyer, G. K., Stoffels, R. J., and Pridmore, P. A. (2014). “Morphology, metabolism and behaviour: responses of three fishes with different lifestyles to acute hypoxia”. In: Freshwater Biology 59.4, pp. 819–831. DOI: 10.1111/fwb.12306.

26. Eby, L. A., Crowder, L. B., McClellan, C. M., Peterson, C. H., and Powers, M. J. (2005). “Habitat degradation from intermittent hypoxia: impacts on demersal fishes”. In: Marine Ecology Progress Series 291, pp. 249–262. DOI: 10.3354/meps291249.

27. Ekau, W., Auel, H., Pörtner, H.­O., and Gilbert, D. (2010). “Impacts of hypoxia on the structure and processes in pelagic communities (zooplankton, macro­invertebrates and fish)”. In: Biogeosciences 7.5, pp. 1669–1699. DOI: 10.5194/bg-7-1669-2010.

28. Ekström, A., Sundell, E., Morgenroth, D., McArley, T., Gårdmark, A., Huss, M., and Sandblom, E. (2021). “Cardiorespiratory adjustments to chronic environmental warming improve hypoxia tolerance in European perch (Perca fluviatilis)”. In: Journal of Experimental Biology 224.6. jeb241554. DOI: 10.1242/jeb.241554.

29. Essington, T. E., Kitchell, J. F., and Walters, C. J. (2001). “The von Bertalanffy growth function, bioenergetics, and the consumption rates of fish”. In: Canadian Journal of Fisheries and Aquatic Sciences 58.11, pp. 2129–2138. DOI: 10.1139/f01-151.

30. Farrell, A. P. and Richards, J. G. (2009). “Defining Hypoxia: An Integrative Synthesis of the Responses of Fish to Hypoxia”. In: Fish Physiology. Vol. 27. Elsevier, pp. 487–503. DOI: 10.1016/S1546-5098(08)00011-3.

31. Griffiths, J. R., Kadin, M., Nascimento, F. J., Tamelander, T., Törnroos, A., Bonaglia, S., Bonsdorff, E., Brüchert, V., Gårdmark, A., Järnström, M., Jonne Kotta MARTIN Lindegren, M. C. N., Norkko, A., Olsson, J., Weigel, B., Žydelis, R., Blenckner, T., Niiranen, S., and Winder, M. (2017). “The importance of benthic–pelagic coupling for marine ecosystem functioning in a changing world”. In: Global change biology 23.6, pp. 2179–2196. DOI: 10.1111/gcb.13642.

32. Haahtela, I. (1990). “What do Baltic studies tell us about the isopod Saduria entomon (L.)?” In: Annales Zoologici Fennici. Vol. 27. 3. JSTOR, pp. 269–278. URL: https://www.jstor.org/stable/23736048.

33. Haase, K., Orio, A., Pawlak, J., Pachur, M., and Casini, M. (2020). “Diet of dominant demersal fish species in the Baltic Sea: Is flounder stealing benthic food from cod?” In: Marine Ecology Progress Series 645, pp. 159–170. DOI: 10.3354/meps13360.

34. Hrycik, A. R., Almeida, L. Z., and Höök, T. O. (2017). “Sub­lethal effects on fish provide insight into a biologically­relevant threshold of hypoxia”. In: Oikos 126.3, pp. 307–317. DOI: 10.1111/oik.03678.

35. ICES (2020). Baltic Fisheries Assessment Working Group (WGBFAS). Tech. rep. 45. ICES Scientific Reports, p. 643. DOI: 10.17895/ices.pub.6024.

36. Karlson, A. M., Gorokhova, E., Gårdmark, A., Pekcan­Hekim, Z., Casini, M., Albertsson, J., Sundelin, B., Karlsson, O., and Bergström, L. (2020). “Linking consumer physiological status to food­web structure and prey food value in the Baltic Sea”. In: Ambio 49.2, pp. 391–406. DOI: 10.1007/s13280-019-01201-1.

37. Killen, S. S., Marras, S., Ryan, M. R., Domenici, P., and McKenzie, D. J. (2012). “A relationship between metabolic rate and risk­taking behaviour is revealed during hypoxia in juvenile European sea bass”. In: Functional Ecology 26.1, pp. 134–143. DOI: 10.1111/j.1365-2435.2011.01920.x.

38. Koenigstein, S., Mark, F. C., Gößling­Reisemann, S., Reuter, H., and Poertner, H.­O. (2016). “Modelling climate change impacts on marine fish populations: process­based integration of ocean warming, acidification and other environmental drivers”. In: Fish and Fisheries 17.4, pp. 972–1004. DOI: 10.1111/faf.12155.

39. Köster, F. W., Huwer, B., Hinrichsen, H.­H., Neumann, V., Makarchouk, A., Eero, M., Dewitz, B. V., Hüssy, K., Tomkiewicz, J., Margonski, P., Temming, A., Hermann, J.­P., Oesterwind, D., Dierking, J., Kotterba, P., and Plikshs, M. (2017). “Eastern Baltic cod recruitment revisited—dynamics and impacting factors”. In: ICES Journal of Marine Science 74.1, pp. 3–19. DOI: 10.1093/icesjms/fsw172.

40. Köster, F. W., Möllmann, C., Hinrichsen, H.­H., Wieland, K., Tomkiewicz, J., Kraus, G., Voss, R., Makarchouk, A., MacKenzie, B. R., St. John, M. A., Schnack, D., Rohlf, N., Linkowski, T., and Beyer, J. E. (2005). “Baltic cod recruitment–the impact of climate variability on key processes”. In: ICES Journal of Marine Science 62.7, pp. 1408–1425. DOI: 10.1016/j.icesjms.2005.05.004.

41. Kramer, D. L. (1987). “Dissolved oxygen and fish behavior”. In: Environmental Biology of Fishes 18.2, pp. 81–92. DOI: 10.1007/BF00002597.

42. Limburg, K. E. and Casini, M. (2018). “Effect of Marine Hypoxia on Baltic Sea Cod Gadus morhua: evidence From Otolith Chemical Proxies”. In: Frontiers in Marine Science 5.482. DOI: 10.3389/fmars.2018.00482.

43. — (2019). “Otolith chemistry indicates recent worsened Baltic cod condition is linked to hypoxia exposure”. In: Biology Letters 15.12, p. 20190352. DOI: 10.1098/rsbl.2019.0352.

44. Limburg, K. E., Olson, C., Walther, Y., Dale, D., Slomp, C. P., and Høie, H. (2011). “Tracking Baltic hypoxia and cod migration over millennia with natural tags”. In: Proceedings of the National Academy of Sciences 108.22, E177–E182. DOI: 10.1073/pnas.1100684108.

45. Ludsin, S. A., Zhang, X., Brandt, S. B., Roman, M. R., Boicourt, W. C., Mason, D. M., and Costantini, M. (2009). “Hypoxia­avoidance by planktivorous fish in Chesapeake Bay: implications for food web interactions and fish recruitment”. In: Journal of Experimental Marine Biology and Ecology 381, S121–S131. DOI: 10.1016/j.jembe.2009.07.016.

46. Luo, J., Hartman, K. J., Brandt, S. B., Cerco, C. F., and Rippetoe, T. H. (2001). “A spatially­explicit approach for estimating carrying capacity: an application for the Atlantic menhaden (Brevoortia tyrannus) in Chesapeake Bay”. In: Estuaries 24.4, pp. 545–556. DOI: 10.2307/1353256.

47. Maciak, S. and Konarzewski, M. (2010). “Repeatability of standard metabolic rate (SMR) in a small fish, the spined loach (Cobitis taenia)”. In: Comparative Biochemistry and Physiology Part A: Molecular & Integrative Physiology 157.2, pp. 136–141. DOI: 10.1016/j.cbpa.2010.05.017.

48. McBryan, T., Anttila, K., Healy, T., and Schulte, P. (2013). “Responses to Temperature and Hypoxia as Interacting Stressors in Fish: implications for Adaptation to Environmental Change”. In: Integrative and Comparative Biology 53.4, pp. 648–659. DOI: 10.1093/icb/ict066.

49. McCrackin, M. L., Muller­Karulis, B., Gustafsson, B. G., Howarth, R. W., Humborg, C., Svanbäck, A., and Swaney, D. P. (2018). “A Century of Legacy Phosphorus Dynamics in a Large Drainage Basin”. In: Global Biogeochemical Cycles 32.7, pp. 1107–1122. DOI: 10.1029/2018GB005914.

50. Mohan, J. and Walther, B. (2016). “Out of breath and hungry: natural tags reveal trophic resilience of Atlantic croaker to hypoxia exposure”. In: Marine Ecology Progress Series 560, pp. 207–221. DOI: 10.3354/meps11934.

51. Mohrholz, V. (2018). “Major Baltic Inflow Statistics–Revised”. In: Frontiers in Marine Science 5, p. 384. DOI: 10.3389/fmars.2018.00384.

52. Möllmann, C., Kornilovs, G., Fetter, M., and Köster, F. (2004). “Feeding ecology of central Baltic Sea herring and sprat”. In: Journal of Fish Biology 65.6, pp. 1563–1581. DOI: 10.1111/j.0022-1112.2004.00566.x.

53. Neuenfeldt, S., Bartolino, V., Orio, A., Andersen, K. H., Andersen, N. G., Niiranen, S., Bergström, U., Ustups, D., Kulatska, N., and Casini, M. (2020). “Feeding and growth of Atlantic cod (Gadus morhua L.) in the eastern Baltic Sea under environmental change”. In: ICES Journal of Marine Science 77.2, pp. 624–632. DOI: 10.1093/icesjms/fsz248.

54. Neuenfeldt, S. and Beyer, J. E. (2006). “Environmentally driven predator–prey overlaps determine the aggregate diet of the cod Gadus morhua in the Baltic Sea”. In: Marine Ecology Progress Series 310, pp. 151–163. DOI: 10.3354/meps310151.

55. Nilsson, G. E. and Östlund­Nilsson, S. (2008). “Does size matter for hypoxia tolerance in fish?” In: Biological Reviews 83.2, pp. 173–189. DOI: 10.1111/j.1469-185X.2008.00038.x.

56. Orio, A., Bergström, U., Florin, A.­B., Lehmann, A., Šics, I., and Casini, M. (2019). “Spatial contraction of demersal fish populations in a large marine ecosystem”. In: Journal of Biogeography 46.3, pp. 633–645. DOI: 10.1111/jbi.13510.

57. Pachur, M. E. and Horbowy, J. (2013). “FOOD COMPOSITION AND PREY SELECTION OF COD, GADUS MORHUA (ACTINOPTERYGII: GADIFORMES: GADIDAE), IN THE SOUTHERN BALTIC SEA”. In: Acta Ichthyologica et Piscatoria 43.2, pp. 109–118. DOI: 10.3750/AIP2013.43.2.03.

58. Pan, Y., Ern, R., and Esbaugh, A. (2016). “Hypoxia tolerance decreases with body size in red drum Sciaenops ocellatus”. In: Journal of Fish Biology 89.2, pp. 1488–1493. DOI: 10.1111/jfb.13035.

59. Peters, R. H. (1986). The ecological implications of body size. Vol. 2. Cambridge University Press.

60. Pichavant, K., Person­Le­Ruyet, J., Bayon, N. L., Severe, A., Roux, A. L., and Boeuf, G. (2001). “Comparative effects of long­term hypoxia on growth, feeding and oxygen consumption in juvenile turbot and European sea bass”. In: Journal of Fish Biology 59.4, pp. 875–883. DOI: 10.1111/j.1095-8649.2001.tb00158.x.

61. Pihl, L. (1994). “Changes in the Diet of Demersal Fish due to Eutrophication­Induced Hypoxia in the Kattegat, Sweden”. In: Canadian Journal of Fisheries and Aquatic Sciences 51.2 pp. 321–336. DOI: 10.1139/f94-033.

62. Pihl, L., Baden, S. P., Diaz, R. J., and Schaffner, L. C. (1992). “Hypoxia­induced structural changes in the diet of bottom­feeding fish and Crustacea”. In: Marine Biology 112.3, pp. 349–361. DOI: 10.1007/BF00356279.

63. Pollock, M., Clarke, L., and Dubé, M. (2007). “The effects of hypoxia on fishes: from ecological relevance to physiological effects”. In: Environmental Reviews 15.NA, pp. 1–14.

64. Powers, S. P., Peterson, C. H., Christian, R. R., Sullivan, E., Powers, M. J., Bishop, M. J., and Buzzelli, C. P. (2005). “Effects of eutrophication on bottom habitat and prey resources of demersal fishes”. In: Marine Ecology Progress Series 302, pp. 233–243. DOI: 10.3354/meps302233.

65. R Core Team (2021). R: A Language and Environment for Statistical Computing. v. 4.2.2. R Foundation for Statistical Computing. Vienna, Austria. URL: https://www.R-project.org/.

66. Reum, J. C., Blanchard, J. L., Holsman, K. K., Aydin, K., Hollowed, A. B., Hermann, A. J., Cheng, W., Faig, A., Haynie, A. C., and Punt, A. E. (2020). “Ensemble Projections of Future Climate Change Impacts on the Eastern Bering Sea Food Web Using a Multispecies Size Spectrum Model”. In: Frontiers in Marine Science 7.124. DOI: 10.3389/fmars.2020.00124.

67. Riedel, B., Pados, T., Pretterebner, K., Schiemer, L., Steckbauer, A., Haselmair, A., Zuschin, M., and Stachowitsch, M. (2014). “Effect of hypoxia and anoxia on invertebrate behaviour: ecological perspectives from species to community level”. In: Biogeosciences 11.6, pp. 1491–1518. DOI: 10.5194/bg-11-1491-2014.

68. Roberts, J. J., Höök, T. O., Ludsin, S. A., Pothoven, S. A., Vanderploeg, H. A., and Brandt, S. B. (2009). “Effects of hypolimnetic hypoxia on foraging and distributions of Lake Erie yellow perch”. In: Journal of Experimental Marine Biology and Ecology 381. Ecological Impacts of Hypoxia on Living Resources, S132–S142. DOI: 10.1016/j.jembe.2009.07.017.

69. Rogers, N. J., Urbina, M. A., Reardon, E. E., McKenzie, D. J., Birchenough, S. N., and Wilson, R. W. (2021). Database of critical oxygen level (P_crit_) in freshwater and marine fishes 1974–2015. Version 1. Cefas, UK. DOI: 10.14466/CefasDataHub.121.

70. Rogers, N. J., Urbina, M. A., Reardon, E. E., McKenzie, D. J., and Wilson, R. W. (2016). “A new analysis of hypoxia tolerance in fishes using a database of critical oxygen level (P_crit_)”. In: Conservation Physiology 4.1. cow012. DOI: 10.1093/conphys/cow012.

71. Roman, M. R., Brandt, S. B., Houde, E. D., and Pierson, J. J. (2019). “Interactive Effects of Hypoxia and Temperature on Coastal Pelagic Zooplankton and Fish”. In: Frontiers in Marine Science 6, p. 139. DOI: 10.3389/fmars.2019.00139.

72. Rose, K. A., Adamack, A. T., Murphy, C. A., Sable, S. E., Kolesar, S. E., Craig, J. K., Breitburg, D. L., Thomas, P., Brouwer, M. H., Cerco, C. F., and Diamond, S. (2009). “Does hypoxia have population­level effects on coastal fish? Musings from the virtual world”. In: Journal of Experimental Marine Biology and Ecology 381, S188–S203. DOI: 10.1016/j.jembe.2009.07.022.

73. Rose, K. A., Gutiérrez, D., Breitburg, D., Conley, D., Craig, K. J., Froehlich, H. E., Jeyabaskaran, R., Kripa, V., Mbaye, B. C., Mohamed, K., Padua, S., and Prema, D. (2019). “Impacts of ocean deoxygenation on fisheries”. In: Schmidtko, S., Stramma, L., and Visbeck, M. (2017). “Decline in global oceanic oxygen content during the past five decades”. In: Nature 542.7641, pp. 335–339. DOI: 10.1038/nature21399.

74. Scott, F., Blanchard, J. L., and Andersen, K. H. (2014). “mizer: an R package for multispecies, trait­based and community size spectrum ecological modelling”. In: Methods in Ecology and Evolution 5.10, pp. 1121–1125. DOI: 10.1111/2041-210X.12256.

75. Silvert, W. and Platt, T. (1978). “Energy flux in the pelagic ecosystem: a time­dependent equation”. In: Limnology and Oceanography 23.4, pp. 813–816. DOI: 10.4319/lo.1978.23.4.0813.

76. Slater, W. L., Pierson, J. J., Decker, M. B., Houde, E. D., Lozano, C., and Seuberling, J. (2020). “Fewer Copepods, Fewer Anchovies, and More Jellyfish: How Does Hypoxia Impact the Chesapeake Bay Zooplankton Community?” In: Diversity 12.1, p. 35. DOI: 10.3390/d12010035.

77. Snoeijs­Leijonmalm, P. and Andrén, E. (2017). “Why is the Baltic Sea so special to live in?” In: Biological oceanography of the Baltic Sea. Ed. by P. Snoeijs­Leijonmalm, H. Schubert, and T. Radziejewska. Springer, pp. 23–84. DOI: 10.1007/978-94-007-0668-2_2.

78. Steube, T. R., Altenritter, M. E., and Walther, B. D. (2021). “Distributive stress: individually variable responses to hypoxia expand trophic niches in fish”. In: Ecology 102.6, e03356. DOI: 10.1002/ecy.3356.

79. Stramma, L., Prince, E. D., Schmidtko, S., Luo, J., Hoolihan, J. P., Visbeck, M., Wallace, D. W., Brandt, P., and Körtzinger, A. (2012). “Expansion of oxygen minimum zones may reduce available habitat for tropical pelagic fishes”. In: Nature Climate Change 2, pp. 33–37. DOI: 10.1038/nclimate1304.

80. Tunney, T. D., McCann, K. S., Lester, N. P., and Shuter, B. J. (2014). “Effects of differential habitat warming on complex communities”. In: Proceedings of the National Academy of Sciences 111.22, pp. 8077–8082. DOI: 10.1073/pnas.1319618111.

81. Ultsch, G. R. and Regan, M. D. (2019). “The utility and determination of P_crit_ in fishes”. In: Journal of Experimental Biology 222.22. jeb203646. DOI: 10.1242/jeb.203646.

82. United Nations (2022). Department of Economic and Social Affairs, Sustainable Development. URL: https://sdgs.un.org/goals.

83. Vainikka, A., Gårdmark, A., Bland, B., and Hjelm, J. (2009). “Two­and three­dimensional maturation reaction norms for the eastern Baltic cod, Gadus morhua”. In: ICES Journal of Marine Science 66.2, pp. 248–257. DOI: 10.1093/icesjms/fsn199.

84. Wang, S. Y., Lau, K., Lai, K.­P., Zhang, J.­W., Tse, A. C.­K., Li, J.­W., Tong, Y., Chan, T.­F., Wong, C. K.­C., Chiu, J. M.­Y., Sze­Tsai Wong, A., Yuen­Chong Kong, R., and Shiu­Sun Wu, R. (2016). “Hypoxia causes transgenerational impairments in reproduction of fish”. In: Nature communications 7.12114, pp. 1–9. DOI: 10.1038/ncomms12114.

85. Wang, T., Lefevre, S., Huong, D. T. T., Cong, N. van, and Bayley, M. (2009). “The Effects of Hypoxia on Growth and Digestion”. In: Fish Physiology. Vol. 27. Elsevier, pp. 361–396. DOI: 10.1016/S1546-5098(08)00008-3.

86. Zhang, H., Mason, D. M., Stow, C. A., Adamack, A. T., Brandt, S. B., Zhang, X., Kimmel, D. G., Roman, M. R., Boicourt, W. C., and Ludsin, S. A. (2014). “Effects of hypoxia on habitat quality of pelagic planktivorous fishes in the northern Gulf of Mexico”. In: Marine Ecology Progress Series 505, pp. 209–226. DOI: 10.3354/meps10768.

