## Supplementary material for "Declining food availability and habitat shifts drive community responses to marine hypoxia": LaTeX files: main.tex

\documentclass[leqno,12pt,english]{article}
\usepackage[document]{ragged2e}
\usepackage[utf8]{inputenc}
\usepackage[left = 1in, right = 1in, top = 1in, bottom = 1in]{geometry}
\usepackage[displaymath]{lineno}
\usepackage{mathtools}
\usepackage{unicode-math}
\usepackage{bm}
\usepackage{setspace}
\usepackage[style=authoryear,uniquename=false,uniquelist=false,maxcitenames=2,maxbibnames=99,giveninits=true]{biblatex}
\DeclareNameAlias{sortname}{last-first}
\addbibresource{ssm.bib}
\addbibresource{supp.bib}
\usepackage{chemformula}
\usepackage{threeparttable}
\usepackage{caption}
\usepackage{pdflscape}
\usepackage{titling}
\usepackage{authblk}
\usepackage{verbatim}
\usepackage{multirow}
\usepackage{booktabs}
\usepackage{adjustbox}
\usepackage{fontspec}
\setmainfont{Times New Roman}
\usepackage{hyperref}
\hypersetup{
    colorlinks=true,
    linkcolor=blue,
    urlcolor=blue,
    citecolor=blue
}

\newcommand\numberthis{\addtocounter{equation}{1}\tag{\theequation}}

\renewcommand\Affilfont{\footnotesize}
\linenumbers

\begin{document}

\title{Declining food availability and habitat shifts drive community responses to marine hypoxia}
\author[1,2,*]{E Duskey}
\author[1,3]{M Casini}
\author[1,2]{KE Limburg}
\author[1]{A G\r{a}rdmark}
\affil[1]{Department of Aquatic Resources, Swedish University of Agricultural Sciences, Uppsala, Sweden}
\affil[2]{Department of Environmental and Forest Biology, College of Environmental Science and Forestry, State University of New York, Syracuse, NY, USA}
\affil[3]{Department of Biological, Geological and Environmental Sciences, University of Bologna, Bologna, Italy}
\date{}
\begin{large}
    \paragraph{\noindent \large} Journal name: Ecological Applications
    \paragraph{\noindent} Manuscript type: Article
\end{large}
{\let\newpage\relax\maketitle}
\paragraph{Open Research statement:} All data and parameters necessary to replicate this modeling study are available at \href{https://github.com/epduskey/hypoxiaSSM}{https://github.com/epduskey/hypoxiaSSM}.  The code contained therein is a novel, extensive modification of the R package mizer, available at \href{https://github.com/sizespectrum/mizer}{https://github.com/sizespectrum/mizer}.  Data and code will be archived in Zenodo upon publication.
\paragraph{Keywords:} Baltic Sea; environmental hypoxia; fish community dynamics; food web models; ocean deoxygenation; size spectrum models

\doublespacing
\clearpage

\begin{abstract}
    Worsening marine hypoxia has had severe negative consequences for fish communities across the globe.  While individual- and population-level impacts of deoxygenation have been identified, it is unknown how they interact to drive changes in food webs.  To address this, we incorporated several major impacts of hypoxia, including declines in benthic resources, habitat shifts, increasing mortality, and changes to rates of feeding, assimilation, and reproductive efficiency, into an existing size spectrum food web modeling framework.  We used this structure to ask the following questions: which of these direct effects are most critical to capturing population and community dynamics in a representative hypoxic system, how do they interact to result in community responses to deoxygenation, and what are the potential consequences of these effects in the context of accelerating deoxygenation?  We tested the effect of different combinations of oxygen-dependent processes, driven by observed oxygen levels, on the food web model's ability to explain time series of observed somatic growth, diets, biomass, and fishery yields of commercially relevant species in the Baltic Sea.  Model results suggest that food availability is most critical to capturing observed dynamics.  There is also some evidence for oxygen-dependent habitat use and physological rates as drivers of observed dynamics.  Deoxygenation results in declining growth both of benthic and benthopelagic fish species, as the latter are unable to compensate for the loss of benthic resources by consuming more pelagic fish and resources.  Analysis of scenarios of ideal, declining, and degraded oxygen conditions show that deoxygenation results in a decline in somatic growth of predators, an altered habitat occupancy resulting in changing species interactions, and a shift in energy flow to benthopelagic predators from benthic to pelagic resources.  This may have important implications for management as oxygen declines or improves.  
\end{abstract}

\clearpage

\begin{refsection}

\section*{Introduction}

Marine hypoxia occurs when dissolved oxygen declines to levels that are sub-optimal for aerobic life.  Hypoxia may occur naturally in some systems, but it has increased worldwide in both severity and frequency due to anthropogenic nutrient enrichment \parencite{breitburg2018declining}, warming-induced increases in oxygen consumption \parencite{brewer2016ocean} and stratification \parencite{diaz2009hypoxic}, and decreases in oxygen solubility with rising ocean temperatures, particularly in the epipelagic \parencite{schmidtko2017decline}.  There are a wide variety of negative impacts of hypoxia on fishes, ranging from altered functioning of cells to community reorganization \parencite{pollock2007effects}.  This presents a challenge to fish species and their prey and hinders society's ability to use ocean resources sustainably \parencite{sdg}.
\paragraph{} The effects of deoxygenation are documented at several levels of biological organization.  For example, at the population level, we might observe changes in reproductive potential \parencite{wang2016hypoxia} and habitat use \parencite{orio2019spatial,zhang2014effects}.  Changes in the chemical environment may promote the production of toxicants, resulting in increased mortality \parencite{diaz2009hypoxic}.  At the community level, differential responses among species or functional groups can result in spatial reorganization \parencite{chu2017ecophysiological}, changes in species composition \parencite{slater2020fewer}, and adaptive foraging resulting in significant changes in food web structure and energy flow \parencite{breitburg1999linking,breitburg2003pattern,pollock2007effects}.  Observations at these higher levels of organization arise from behavioral and physiological responses of individuals.  There are several compensatory mechanisms that individual fish employ in order to mitigate the effects of hypoxia, including an increase in ventilation and gill perfusion, increased hemoglobin and oxygen extraction, decreases in activity \parencite{farrell2009defining}, as well as behaviors such as aquatic surface respiration \parencite[ASR;][]{dwyer2014morphology}, and changes in swimming activity \parencite{domenici2013effect} and habitat occupancy \parencite{eby2005habitat,stramma2012expansion,casini2019spatio}.  Indirect effects on fish may also be common.  For example, benthic invertebrates may significantly alter behavior and ultimately be subject to increased mortality under hypoxia \parencite{riedel2014effect}.  This may lead to changes in the diets of predatory fish that feed on them \parencite{powers2005effects}.  Overall, there is a good body of evidence linking low oxygen concentration to individual responses of fish, less evidence indicating how these result in population-level responses \parencite{bergman2019scaling}, and relatively little evidence describing how these in turn impact community-level responses \parencite{rose2009does}.  Of the latter, investigations of the impacts of hypoxia on predator-prey interactions suggest that differential sensitivity of each species to declining oxygen can result in changes in spatial overlap, predator activity, and prey vulnerability \parencite{breitburg1997varying,chu2017ecophysiological}.  However, it is unknown which of these mechanisms, if any, are most critical and how they interact to drive the observed, higher-level dynamics.  
\paragraph{} Integrative modeling may help to resolve some of these unknowns by drawing direct connections between the effects of environmental drivers on individuals and higher-level outcomes \parencite{pollock2007effects}.  Hypoxia operates on several physiological and behavioral pathways of fish simultaneously.  Its impacts are dependent on body size \parencite{ekau2010impacts}, and are often caused by changes in feeding \parencite{wang2009effects}.  Hypoxia may thus affect individuals, cause emergent effects on populations, and impact the overall community structure by changing competitive dominance and feeding relationships.  Therefore, a size structured food web model offers an appropriate framework with which to evaluate the relative importance of major individual-level pathways suggested by the literature as potential drivers of observed population and community dynamics in hypoxic habitats \parencite{koenigstein2016modelling}.
\paragraph{} In this study, we developed an oxygen-dependent version of a size spectrum food web model in order to ask the following questions: (1) Which individual-level impacts of deoxygenation are most critical to capturing observed population and community dynamics?  (2) How do they interact to result in community-level responses to deoxygenation? and (3) What are the potential consequences of these effects in the context of accelerating deoxygenation? We used the Baltic Sea as a case study and fitted several parameters to a simplified fish food web to explore how environmental hypoxia can affect community composition and dynamics, as well as characteristics of the commercial fisheries yield.  By testing the ability of several individual-level pathways -- including food availability, vertical habitat occupancy, mortality, and physiological rates of feeding, reproduction, and assimilation -- to describe observations at the population and community scale, we identified potential physiological and behavioral drivers of the deleterious effects of hypoxia on marine food webs.

\section*{Methods}

\subsection*{Size Spectrum Food Web Model}

We modified the size-structured food web modeling framework \textbf{mizer} \parencite{mizer} in order to represent fish community dynamics in response to declines in benthic oxygen levels.  In food web models of this sort, maximum body size serves as the master trait of each species, determining somatic growth through feeding, reproduction through maturation size and growth, and mortality through size-dependent predation and natural senescence \parencite{peters1986ecological,blanchard2017bacteria}.  For our model, we built a general structure (\href{https://github.com/epduskey/hypoxiaSSM}{found here}) composed of a benthopelagic predator, a benthic predator, two species of pelagic forage fish, as well as these species' basal resources in separate benthic and pelagic habitats.  This system reflects a community with two competing benthic predators, one of which is more resilient to hypoxia than the other, two competing pelagic prey fish species, and benthic and pelagic invertebrate fauna on which they feed.  
\paragraph{} The fundamental and theoretical foundation of these models is described in \textcite{andersen2016theoretical}.  They are generally defined by three major ecological processes, all of which scale with body size: somatic growth, reproduction, and mortality.  The dynamic size spectrum of each species and basal resource is determined by a flow of energy, or equivalent mass, via trophic relationships, as well as mortality.  Conservation of mass is guided by the McKendrick-von Foerster equation \parencite{silvert1978energy}:
    \begin{linenomath*}
    \postdisplaypenalty = 0
    \begin{equation}
        \frac{\partial n_{i}}{\partial t} + \frac{\partial g_{i}n_{i}}{\partial w} = -m_{i}n_{i}
    \end{equation} \label{eq:mckvf}
    \end{linenomath*}
where \(n_{i}\), \(g_{i}\), and \(m_{i}\) describe time-dependent population size, somatic growth, and mortality as a function of body size for each species \(i\), respectively.  Recruitment of individuals of species \(i\) at a minimal size \(w_{0}\) is described by the boundary condition:
    \begin{linenomath*}
    \postdisplaypenalty = 0
    \begin{equation}
        g_{i}(w_{0})n_{i}(w_{0}) = R_{i}
    \end{equation} \label{eq:recruit}
    \end{linenomath*}
Losses of individual biomass construed as metabolic activity, which also scales with body size \parencite[see][]{brown2004toward}, and assimilation efficiency, determine the conversion of ingested mass into somatic and reproductive growth.  Each individual of each species is recruited to the population at weight \(w_{0}\) and grows by consuming a combination of basal resources and other fish.  Growth is also determined in part by species-specific von Bertalanffy growth parameters \parencite{essington2001bertalanffy}, here estimated from data.  The basal resources are governed by a simple semi-chemostat function, as well as by the degree of predation.  Recruitment is limited by density dependent processes, typically by the inclusion of a Beverton-Holt or Ricker type stock-recruitment relationship.  Feeding is guided by size- and species-preference, determined by both a distribution describing the preferred predator:prey mass ratio (PPMR) and a square matrix describing horizontal spatial overlap of each species with all others.  A maturity ogive guides the allocation of acquired energy to somatic growth versus reproduction throughout an individual's life.  Mortality occurs either through predation, unspecified natural processes, or fishing, the latter of which is determined by user-defined fishing selectivity, effort, and catchability.  We estimated each of these parameters with data arising from the Central Baltic Sea (Appendix S2; Figures \ref{fig:all_lengthweight}--\ref{fig:all_ppmr} and Tables \ref{tab:data} and \ref{tab:allparams} for parameter estimates and data sources).  Given this model structure's focus on individual physiological processes and how these govern community dynamics, it is appropriate for the inclusion of documented effects of hypoxia on feeding, habitat occupancy, metabolism, and mortality (see Figure \ref{fig:concept}).

\subsection*{Oxygen dependence}

In general, we included deleterious effects of hypoxia in our model by increasing costs and decreasing vital physiological and ecological rates with bounded and continuous functions of oxygen decline (Appendix S1). Though the hypoxia threshold is often defined as an oxygen concentration of 2 mg\(\cdot\)L\textsuperscript{-1} \parencite{hrycik2017sub}, a range of concentrations above this threshold are known to have sub-lethal, yet critical effects on the ecology of fishes \parencite{kramer1987dissolved}.  These effects may or may not vary by body size \parencite{hrycik2017sub,pan2016hypoxia}, but they do vary by species \parencite{nilsson2008does}.  In a size spectrum food web framework, each individual may be affected to a greater or lesser degree relative to its competitors.  Therefore, within these models, hypoxia has the potential to change competitive and predatory interactions, and the direction and magnitude is dependent on several characteristics of the species involved.  This flexibility, combined with a focus on individuals throughout ontogeny and ultimately leading to dynamics at a scale at which they are observed, allows us to ask specific questions about the relevance of individual processes at the community scale.
\paragraph{}  More specifically, we altered the baseline structure of \textbf{mizer} \parencite{mizer} by adding dynamic independent variables for benthic and pelagic oxygen, given here in units of mL\(\cdot\)L\textsuperscript{-1}.  We used the concept of critical oxygen level, or P\textsubscript{crit}, to describe individuals' relative sensitivity to estimated oxygen levels.  The P\textsubscript{crit} of an individual is the oxygen level below which standard metabolism can no longer be maintained and begins to decline with ambient levels \parencite{ultsch2019utility}.  That is, it is the level at which oxyregulators (i.e. organisms which maintain constant oxygen consumption) become oxyconformers (i.e. organisms whose oxygen consumption varies with environmental conditions) due to the lack of adequate oxygen to cover basal metabolic costs \parencite{rogers2016new}.  This value depends not only on species identity  \parencite{farrell2009defining}, but also on salinity, temperature, and body weight \parencite{rogers2016new}.  While we kept salinity and temperature constant, maximum weight is one of the defining attribute of individuals in \textbf{mizer}, and therefore both behavioral and physiological responses to oxygen may change throughout ontogeny.
\paragraph{} We used a P\textsubscript{crit} database \parencite{pcritdatabase2021} to estimate P\textsubscript{crit} as a function of temperature, salinity, body size, and resting metabolic rate (RMR), as in \textcite{rogers2016new} (Appendix S4; see Figure \ref{fig:pcrit_size}).  We scaled the following rates and values as a logistic function of each species' oxygen exposure, where applicable: benthic resource carrying capacity, occupancy in the benthic habitat, maximum consumption rate, fish search rate for prey, assimilation efficiency, and fish egg survival.  The general equation is:
    \begin{linenomath*}
    \begin{equation} \label{eq:scaling}
        \xi(O_{iwt}) = \frac{1}{1 + \exp{\left(-U_{i}(O_{iwt}-a_{i}\,\textsf{P}_{\textsf{crit},iw})\right)}}
    \end{equation}
    \end{linenomath*}
where \(O_{iwt}\) is mean oxygen concentration experienced by species \(i\) of weight \(w\) at time \(t\), and \(U_{i}\) captures the sensitivity of species \(i\) to low oxygen, with lower values representing more gradual declines at higher oxygen levels.  The product \(a_{i}\,\textsf{P}_{\textsf{crit},iw}\) represents the oxygen level at which rates of an individual of species \(i\) at weight \(w\) have fallen to 50\% of their maximum.  This scaling function is very similar to that used by \textcite{luo2001spatially}, and reflects a simple and flexible method by which to include individual responses to oxygen.  All the rates that we scaled according to equation \ref{eq:scaling} represent activities in excess of standard metabolic rate (SMR).  SMR as represented by oxygen consumption can be approximated by P\textsubscript{crit} \parencite{ekau2010impacts}, and thus treating P\textsubscript{crit} as a shifting inflection point may be appropriate.  We estimated one set of parameters for benthic resource carrying capacity, another for benthic occupancy, and a third set for maximum consumption rate, fish search rate for prey, assimilation efficiency, and egg survival.  Note that inherent in this structure is the assumption that all oxygen-dependent physiological processes decline with oxygen at the same rate.  That is, the prioritization of activity, feeding, assimilation efficiency, etc. of fishes of each species does not change as hypoxia develops, aside from variation in P\textsubscript{crit} due to body size and species, as indicated in equation \ref{eq:scaling}.  This is a strong assumption, but it significantly reduces the number of parameters that must be estimated, and serves as a simple starting point for analyses of emergent community responses to deoxygenation.
\paragraph{} In addition to the rates listed above, we also scaled metabolism and natural mortality (i.e. mortality not due to fishing or, in this structure, predation) with exposure to hypoxic waters.  Metabolism in the \textbf{mizer} framework is treated as a cost, based on the assumption that costs for maintenance scale proportionally to metabolic rate.  This is because energy is assumed to flow through the system as mass equivalents, with somatic growth arising as a balance between gains via consumption and losses via respiration.  We therefore scaled it with a simple negative exponential function:
    \begin{linenomath}
    \begin{equation} \label{eq:met_scaling}
        \xi(O_{iwt}) = 1 + \exp{(-U_{\textsf{met},i}(O_{iwt}-a_{\textsf{met},i}\,\textsf{P}_{\textsf{crit},iw}))}
    \end{equation}
    \end{linenomath}
Here, values of \(U_{\textsf{met},i}\) and \(a_{\textsf{met},i}\) play a similar role as their counterparts in equation \ref{eq:scaling}, only \(a_{\textsf{met},i}\,\textsf{P}_{\textsf{crit},iw}\) instead represents the oxygen level at which metabolic costs have doubled.  Additional natural mortality due to exposure to hypoxic waters is expressed as a hazard function (Appendix S4).  Both this and occupancy as a function of dynamic variables are new additions to the \textbf{mizer} framework, and reflect the flexibility built into the code provided \parencite{mizer}.

\subsection*{Study System}

\paragraph{} We studied emergent responses to hypoxia in the relatively species poor food web of the Central Baltic Sea, which here is mainly focused on commercially important fish species in the open waters.  This is an important case study, given the severity and extent of deoxygenation in the region \parencite{carstensen2014deoxygenation}.  Our model of this system includes benthopelagic cod (\textit{Gadus morhua}), benthic flounders (\textit{Platichthys flesus} and \textit{Platichthys solemdali}), and the pelagic clupeids sprat (\textit{Sprattus sprattus}) and herring (\textit{Clupea harengus}), as well as their benthic and pelagic invertebrate prey.  This region is particularly prone to episodes of deoxygenation, with a water residence time of roughly 25-30 years, and a relatively stable vertical salinity gradient that isolates bottom waters from oxygenated surface waters \parencite{snoeijs2017baltic}.  Periodic major Baltic inflows (MBIs) rejuvenate oxygen concentrations in these waters, but with decadal variability and with a lower frequency in the last three decades \parencite{mohrholz2018major}.  Though recent nutrient loading reduction has led to improving conditions, extensive eutrophication and a legacy of recycling nutrients continue to plague the Baltic Sea \parencite{mccrackin2018,andersen2017long}.  As a result, hypoxic areas have expanded exponentially since the mid-1990s, reducing the potential habitat for benthic fish such as cod \parencite{limburg2011tracking,casini2016hypoxic,casini2021changes}.  Fish otolith microchemistry has also confirmed individual cod exposure to low oxygen, with negative effects on growth and body condition  \parencite{limburg2018effect,limburg2019otolith}.  For the Baltic Sea, there are extensive fisheries data arising from scientific surveys and commercial data collection \parencite{wgbifs2020}, including mean weight as well as time series of spawning stock biomass (SSB, the combined weight of all reproductively active individuals in the population), and fisheries yield (for the derivation of observation data, see Appendix S2).  For this system, we estimated oxygen trends from data provided by the Swedish Meteorological and Hydrological Institute (SMHI; Appendix S3; Figure \ref{fig:oxy_years} and Table \ref{tab:oxygam}).  The availability of these data makes it ideal for studying the mechanisms that determine the impacts of deoxygenation on fish communities.

\subsection*{Oxygen-dependent processes explaining observed population trends}

\paragraph{} We grouped the oxygen-dependent processes into four general categories: benthic resource carrying capacity (B), cod benthic occupancy (O), mortality (M), and physiology (P) (Appendix S1).  The physiological component (P) includes both increases in metabolic costs and declines in maximum consumption rate, search or clearance rate, assimilation efficiency, and reproductive efficiency.  We fit all 16 possible combinations of these groups, including both the full model and a model with no scaling, to observations of spawning stock biomass (SSB), commercial yield, and somatic growth of the four fish species during a calibration period (1991--2000; Appendix S5).  These models were then used to project forward during the years 2001--2019.  We chose the best model based on minimum weighted error during this projection period only, as out-of-sample model selection is generally more robust \parencite{cooke2014out}.  We also performed a sensitivity analysis (Appendix S6) to determine the robustness of our results.  This procedure can provide insight into which pathways by which hypoxia affects individual fishes that are most critical to an accurate representation of the observed community dynamics.

\subsection*{Scenario analyses}

\paragraph{} We used the best model from the procedure described above (and detailed in Appendix S5) in order to test community responses under various oxygen scenarios.  We ran the calibrated models during the projection period (2001--2019) with all individual components of the best model in order to determine how they interact to produce the responses in each scenario.  These scenarios tracked predicted SSB, yield, and fish weight (in grams) up to maximum observed age in data sources (15 years for cod, 26 for flounder, 16 for sprat, 17 for herring), as well as energy or equivalent mass flow through the food web.  The latter values are calculated as a weighted average of diets for each fish species across body size classes.  Weights are given by the biomass in each body size class.
\paragraph{} For each of these models, we ran three scenarios at benthic oxygen levels spanning the range of observed values throughout the time series.  Mean bottom oxygen concentration reached a peak value of roughly 3 mL\(\cdot\)L\textsuperscript{-1} during a period of inflow in the late 1980's and early 1990's and fell to 1 mL\(\cdot\)L\textsuperscript{-1} in the 2000's and beyond (Appendix S4; Figure \ref{fig:oxy_years}).  Therefore, we chose values of 3, 2, and 1 mL\(\cdot\)L\textsuperscript{-1} representing more ideal, deteriorating, and degraded conditions.  We then compared somatic growth, SSB, yield, and energy flow among the scenarios to determine how communities respond to declining oxygen.

\subsection*{Implementation}

We estimated all parameters, built all models, and analyzed and plotted all output in R \parencite{rmanual}.  All code and parameters necessary to reproduce our results are available on GitHub (\href{https://github.com/epduskey/hypoxiaSSM}{found here}).

\section*{Results}

\subsection*{Oxygen dependence of observed dynamics}

Model calibration results suggest there are interactions among the pathways by which deoxygenation affects individual fish.  No single process nor a combination of processes provided a superior representation of somatic growth, SSB, and yield observed in the four fish species (cod, flounder, sprat, and herring) during the projection period (2001--2019; see Figures \ref{fig:occupancy} and \ref{fig:all_scaling}--\ref{fig:sim_growth_eight} and Tables \ref{tab:cal_params} and \ref{tab:all_scaling}).  When we weighted errors during the projection period (2001--2019) across outputs to reflect a high confidence in growth data, a moderate confidence in SSB estimates, and a low confidence in yield estimates, the BOM model (i.e. the model with oxygen dependent benthic resource carrying capacity (B), cod benthic occupancy (O), and direct mortality (M) included) performed best (Appendix S5; Figure {\ref{fig:final_rank_sim}}).  However, note that direct mortality due to hypoxia (M) does not vary at all with time for both cod and flounder (i.e. \(b_{i}=0\) for all models; see equation \ref{eq:hazard}).  Including this component in the BOM model merely adjusts the baseline mortality of these two species.  Only benthic food availability (B) and cod occupancy in the benthic habitat (O) vary with oxygen.  That is, the interaction of bottom water oxygen on cod ``decision'' to occupy benthic habitats, combined with hypoxia-driven declines in benthic prey, appears best at explaining observed SSB, yield, and growth of our focal fish species in the Baltic Sea food web.
\paragraph{} In general, the BOM model was able to track the observed annual variation in somatic growth of cod throughout the projection period, particularly during the first eight years of life where data are concentrated (Figure {\ref{fig:cod_growth_eight}}; see also Figure {\ref{fig:cod_growth_eight_supp}} in Appendix S7).  At more advanced ages, the model was prone to over-estimation (Appendix S7; Figure \ref{fig:sim_growth}), but there are significantly fewer data for these ages, and therefore less certainty regarding actual mean size-at-age.  Among-model variation of sizes-at-age is particularly notable at older ages (Figure \ref{fig:cod_growth_eight}).  Growth patterns were relatively static from year to year for other species (see Figures \ref{fig:sim_growth} and \ref{fig:sim_growth_eight}).  Flounder scaling parameters were estimated by fitting to a mean predicted von Bertanlanffy growth trajectory across all years, in contrast to the other species for which we had annual estimates for quite a large subset of years.  On average, the model was also able to capture the relatively static growth of sprat and herring (see Figures \ref{fig:sim_growth} and \ref{fig:sim_growth_eight}), in addition to the declines in growth of cod.
\paragraph{} While there is variability, most models produced reasonable fits to trends of SSB and yield during the projection period (Figure \ref{fig:sim_ssb}; see also Figure \ref{fig:sim_ssb_supp} in Appendix S7).  Projections of yield were often over- or under-estimated, but the model did capture the general patterns observed for each species, although less so for flounder (Figure \ref{fig:sim_yield}; see also Figure \ref{fig:sim_yield_supp} in Appendix S7).  Differences among all of our models in fits to SSB and yield during the projection period were relatively small (Figures \ref{fig:sim_ssb} and \ref{fig:sim_yield}); only those models including scaling of physiological rates (P) without including cod occupancy (O) produced egregious errors (Figures \ref{fig:sim_error} and \ref{fig:final_rank_sim}).  Therefore, our methods may be less able to distinguish among drivers of observed trends at the population level compared to those at the individual level.
\paragraph{} Results of the sensitivity analysis (Appendix S6) indicate that the best model according to the procedure described above and detailed in Appendix S5 may change depending upon the starting values for scaling parameters.  This is most certainly due in part to the the inter-dependency of all model components and output.  While the order of the best models does change somewhat from run to run, two things remain consistent.  First, models having the lowest relative weighted error always include oxygen dependence of benthic food resources (Appendix S6; Figure {\ref{fig:sens_rank_sim}}); they may or may not also include oxygen dependence of occupancy and physiological rates.  Second, models with the greatest error always include those with oxygen dependence of physiological rates without oxygen-dependent cod occupancy in the benthic habitat (Appendix S6; Figure \ref{fig:sens_rank_sim}).  That is, benthic food availability and habitat use play a primary role in structuring the results.

\subsection*{Emergent responses to oxygen scenarios}

\paragraph{} Comparing the outcome of oxygen scenarios in the best model (BOM) to its components suggests that changing cod occupancy (O) is a compensatory response to declining benthic food availability (B; Figure {\ref{fig:scenarios}}).  These results also show that the constant mortality rate has a relatively minor effect on the results.  Including oxygen dependence of benthic resource carrying capacity alone (B model) produced unrealistically severe declines in SSB and yield of cod across oxygen scenarios, as well as relatively extreme declines in somatic growth (Figure {\ref{fig:scenarios}}).  Those models including both B and O, by contrast, produce less severe, but still significant, declines in individual and population level metrics.  The models without oxygen dependence of cod benthic occupancy (O) show no variability of sprat and herring SSB and yield with oxygen (Figure {\ref{fig:scenarios}}).  They also tend to produce much greater estimates of biomass and yield of sprat and herring (see Figure {\ref{fig:sim_ssb_supp}}).  This indicates that cod intensify feeding on sprat and herring when relocating to the pelagic habitat, as expected (Figures \ref{fig:diet_scenarios} and \ref{fig:flow_scenarios}; see also Table \ref{tab:supp_scenarios}).  Declines in cod growth, SSB, and yield in the BOM model with declining oxygen (Figure \ref{fig:scenarios}) show that moving towards the pelagic habitat does not compensate entirely for the decline in benthic food.
\paragraph{} Scenario results also suggest that deoxygenation changes the nature of competitive interactions within and between species (Figure \ref{fig:scenarios}).  Cod experience greater declines in somatic growth with declining food availability (B model) compared to flounder.  This also occurs with both declining food availability and changing cod occupancy (BO model; Figures \ref{fig:scenarios} and \ref{fig:flow_scenarios}).  When only cod benthic occupancy depends upon oxygen (O model), flounder SSB and yield increase very slightly as cod moves into the pelagic to escape hypoxia, having been released from moderate competition (Figure \ref{fig:scenarios}).  If benthic food is impacted (B model), then flounder experience marked declines in SSB and yield.  While declines in SSB and yield of cod and flounder are of similar magnitude in the B model, the relative decline in body size of cod is more severe than for flounder (Figure \ref{fig:scenarios}).  This suggests that flounder are the superior competitor within this environment.  By contrast, when both benthic food and cod habitat occupancy are oxygen-dependent (BO and BOM model), cod experience only moderate declines in SSB and yield compared to flounder (Figure \ref{fig:scenarios}).  Low food availability limits flounder in our model (Table \ref{tab:all_scaling}), but unlike for cod, the limitation decreases flounder carrying capacity and does not affect growth as much.  Given the paucity of reliable information on the Baltic flounder populations, it is generally unknown how they are faring in the Central Baltic Sea as a whole.  There is relatively more information for sprat and herring, and here results show that SSB and yield of both sprat and herring decline (Figures \ref{fig:scenarios} and \ref{fig:flow_scenarios}).  As there are no direct effects of benthic deoxygenation on these pelagic species, this is caused by intensifying predation as cod avoid hypoxic bottom waters.  Declines are more severe for sprat, as herring can largely outgrow their vulnerability to cod predation (Figure \ref{fig:diet_scenarios}).  This releases the pelagic fishes from competition somewhat, resulting in further growth (Figure \ref{fig:scenarios}).  We stress that changes in sprat and herring growth are very minor, as their competition for pelagic food sources is weak in the model.  This is primarily caused by setting pelagic resource carrying capacity high in order to reflect observed cod diet, as well as pelagic fish SSB (see Appendix S5).  Nonetheless, model results show that the availability of benthic resources and changing vertical habitat use of cod reorder interactions both in the benthic and the pelagic habitats.
\paragraph{} Finally, declining benthic food availability and increasing use of the pelagic habitat shifts energy flow to cod away from benthic and toward pelagic prey (Figures \ref{fig:diet_scenarios} and \ref{fig:flow_scenarios}).  As cod move up in the water column in response to low oxygen, feeding on flounder and benthic resources declines overall, while feeding on sprat and herring increases (Figure \ref{fig:flow_scenarios}).  Notably, these pelagic fish increase from 25\% to over 50\% of the diet of large cod when oxygen levels decline from 3 to 1 mL\(\cdot\)L\textsuperscript{-1}, with the remainder of their diet consisting of benthic resources. As benthic resources decline with deoxygenation, small cod feed increasingly on plankton, as they are too small to feed significantly on the larger fish prey in the pelagic habitat (Figure \ref{fig:flow_scenarios}).

\section*{Discussion}

Marine hypoxia can have severe and complex consequences for fish and fisheries \parencite{rose2019impacts}.  Results from our modeling study suggest food availability in deoxygenated habitats, as well as the use of oxygenated refugia, may be primary drivers of emergent patterns in fish populations and communities.  As the severity of hypoxia increases, community size structure shifts among benthic and benthopelagic predators to favor smaller individuals, and consumption patterns may change.  This may be driven both by changes in size structure, and by changes in habitat use.  A mismatch in prey availability relative to the needs of predators could limit the ability of changing habitat use to compensate for declines in somatic growth of benthopelagic species. All of these impacts result in a change in the flow of energy among species in the food web.  These insights into potential mechanisms by which the effects of hypoxia on individuals may scale to affect whole communities can provide valuable context for future studies and for the management of fish populations.
\paragraph{} We have attempted to answer three questions on drivers of food web responses to deoxygenation with our analysis: first, which individual-level effects of marine deoxygenation are critical to capturing higher level dynamics in hypoxic systems?  Our analysis of alternative oxygen-dependent pathways in a size-structured food web model suggests that a decrease in the availability of benthic prey is essential for explaining observed responses (spawning stock biomass or SSB, fisheries yield, and somatic growth) to deoxygenation.  Model results support a decline in the use of benthic habitats and subsequent changes in predatory fish feeding from benthic to pelagic prey may also be critical.  Our sensitivity analysis suggests, however, that changes in habitat use may not be as critical as declines in benthic food resources, and that oxygen dependence of physiological rates may also be important (Appendix S6; Figure {\ref{fig:sens_rank_sim}}).  Observations in the Baltic Sea do suggest a decline in benthic food availability \parencite{karlson2020linking,neuenfeldt2020feeding}.  \textit{Saduria entomon}, a major food source for Baltic cod, is limited by oxygen concentrations below about 1.5 mL\(\cdot\)L\textsuperscript{-1} \parencite{haahtela1990baltic}.  Declining benthic food availability (B) remains in all of the best models for the projection period whether or not they also include oxygen dependence of cod benthic occupancy and physiological rates (Appendix S5; Figure {\ref{fig:final_rank_sim}}).  Accounting for feeding interactions and their dependence on oxygen conditions thus appears to be essential for understanding emergent food web dynamics to marine deoxygenation.
\paragraph{} Second, we asked: how do critical individual-level impacts of deoxygenation interact with one another to result in community-level responses?  Examining the ability of all models to explain emergent dynamics in spawning stock biomass (SSB), yield, and fish body growth in the four main commercially exploited fish stocks in the Central Baltic Sea reveals that interactions between mechanisms are important.  The inclusion of some level of oxygen-driven change in feeding, whether through a decline in the biomass of benthic invertebrates (B) and in cod occupancy of benthic habitats (O), through a decline in feeding activity, assimilation efficiency, and egg survival (P), or all three of these, all appear in superior models (Appendix S5; Figure {\ref{fig:final_rank_sim}}).  The former two mechanisms are supported by observations of a decline in cod feeding on benthos \parencite{neuenfeldt2020feeding} and in benthic occupancy of cod \parencite{casini2019spatio}.  The latter is supported by observations of reduced appetite in hypoxic conditions \parencite{pichavant2001comparative,chabot2008environmental}, as well as reduced assimilation efficiency \parencite{wang2009effects} and egg survival \parencite{koster2005baltic} of cod.  Of note is that model results suggest both of these mechanisms (B and P) likely operate alongside altered cod occupancy of the benthic habitat (O) for best model performance (Appendix S5; Figure {\ref{fig:final_rank_sim}}), though the implications differ.  For models with physiological oxygen dependence (P), this implies that, given observed dynamics, there must be an opportunity for cod to escape from degraded environments.  For the best model (BOM), there must be an alternative food source (i.e. access to pelagic prey fish).  The importance of dynamic occupancy of the benthic habitat in representing cod population dynamics and somatic growth is supported by the literature.  Shifts in the relative use of benthic habitat in cod have been observed in the Baltic Sea \parencite{casini2019spatio}, in addition to shifts in cod diets towards clupeids \parencite{pachur2013food}.  Furthermore, it has been observed that diets of cod are driven at least in part by differences in vertical overlap caused by the dynamics of salinity and oxygen in bottom waters \parencite{neuenfeldt2006environmentally,pachur2013food}.  Though changing habitat use may serve as a compensatory mechanism, it is clearly unable to compensate fully for the loss of benthic habitat and benthic prey in our case study.  Given this body of evidence, in addition to our own results, accounting both for the negative effects of deoxygenation as well as potential compensatory mechanisms allows for a more complete understanding of both direct and indirect effects of deoxygenation that emerge in communities of interacting species in connected habitats.
\paragraph{} Lastly we asked: how might we expect declining oxygen levels to affect fish population and community metrics?  Scenario results suggest that, as oxygen declines, most population metrics may also be expected to decline.  That said, at moderate oxygen levels (2 mL\(\cdot\)L\textsuperscript{-1}), cod SSB and yield experienced modest increases on the order of 4-5\% relative to more ideal oxygen conditions (3 mL\(\cdot\)L\textsuperscript{-1}; Figure \ref{fig:scenarios}).  This is due to the expanded access to an additional food source (clupeids).  Recall that the model assumes constant areal overlap of fish species, and oxygen as the only factor motivating cod movement between habitats.  In reality, the horizontal overlap of cod and clupeid prey fish has decreased in the past three decades, with cod concentrated in the southern Baltic Sea and clupeids in the northern areas \parencite{casini2011spatial,casini2016hypoxic}.  Note that, while our model tracks declines in cod benthic occupancy during the calibration period, it fails to account for more recent increases (Appendix S5; Figure \ref{fig:occupancy}).  Therefore, this projected increase in overlap and moderate increase in SSB may be an artefact of model assumptions.  However, at minimum observed oxygen levels (1 mL\(\cdot\)L\textsuperscript{-1}), both SSB and yield of cod decline, with the latter declining more severely.  This is due to stunted growth.  Empirical evidence links reduced growth of cod to increasing exposure to hypoxia \parencite{limburg2018effect,casini2021changes}. While selectivity is constant across scenarios, the yield becomes dominated by smaller individuals as oxygen declines.  All other species experience declines in SSB and yield across deoxygenation scenarios.  The response in body growth depends on species.  Growth of pelagic clupeids is largely insensitive to benthic deoxygenation relative to the declines in growth of the benthic species.  Both cod and flounder experience declines in growth, with that of the former being more severe.  Herring and sprat do experience very slight increases in growth as cod rise in the water column and prey upon the sprat and the herring, more so the former than the latter.  Changes in sprat and herring are particularly interesting, given that their responses to deoxygenation are driven entirely by interactions with cod as they move through the water column.  Thus, all species experience changes at the population level across scenarios, though it varies greatly by species.  Indirect effects arise, particularly with reference to clupeids, as food web interactions change in response to declining oxygen.
\paragraph{} Energy flow through the food web also changes in response to marine deoxygenation, primarily as benthopelagic cod move vertically in the water column.  This mediates indirect effects of hypoxia.  Our model results suggest that size and species composition of prey are altered as cod escape hypoxic conditions.  As benthic food declines, the proportion of pelagic food in the diet of small cod increases, shifting the overall size distribution of cod prey towards smaller items in the early years of life.  Similar shifts in size and species composition of prey sources of cod are observed in the Baltic Sea \parencite{pihl1994changes,haase2020diet,neuenfeldt2020feeding}.  Pelagic fishes become more important in the diet of cod as they grow \parencite{griffiths2017importance}.  According to our results, oxygen declines may intensify this shift.  Overall, the consequences of the interaction between declining benthic food and changing habitat use is a reorganization of energy flow away from benthic and towards pelagic prey among benthopelagic species in the food web, and dramatic changes in the community size structure.  Most notable, this includes a sharp decline in large predators.  It is unlikely that pelagic forage fish are otherwise unaffected by benthic oxygen, as they may use deeper habitats as feeding grounds \parencite{mollmann2004feeding,ludsin2009hypoxia}, or refugia from warming waters, thus altering food web structure \parencite{tunney2014effects}.  That said, it is clear that strong indirect effects can arise in systems driven by deoxygenation.  Community species that are concentrated within oxygenated refugia may suffer from increased predation, while species escaping deoxygenated habitats may suffer from a mismatch in dietary needs relative to availability.
\paragraph{} We made several simplifying assumptions in our model which explicitly defy observations.  For example, we used a static representation of size at maturation during the calibration period for all species, whereas evidence suggests that size at maturation has declined for Baltic cod \parencite{vainikka2009two,koster2017eastern}.  We also assumed a constant temperature.  There is strong evolutionary pressure on both thermal and hypoxic limits for fishes \parencite{deutsch2020metabolic}.  Sustained increases in both temperature, and in the frequency and severity of hypoxic events, may thus result in rapid evolutionary changes.  In general, we would expect there to be strong interactions between declining oxygen and warming, as temperature affects both oxygen supply and demand \parencite{roman2019interactive}.  It is known that these two stressors can act synergistically \parencite{ekstrom2021cardiorespiratory}, though the system-specific response will depend on species identity, species interactions, and community plasticity \parencite{mcbryan2013responses}.  Which mechanisms may drive this interaction is generally unknown, and currently unexplored in our model structure.  We also ignored individual variability in the model outside of differences in species and maximum body size.  Neither did we account for sex.  In reality, differences amongst individuals, including those dependent upon sex, may cause variation in both space and time.  For example, some fish with higher metabolic rates tend to be more active and more willing to take risks in hypoxic environments \parencite{killen2012relationship}.  Depending on the outcome, and given that metabolic rate is an heritable trait \parencite{maciak2010repeatability}, spatiotemporal variability could potentially be driven by changes in selective forces that are in turn driving the distribution of metabolic rates among the population.  Our conclusions may only hold if the variation and mechanisms described above and elsewhere are negligible.  However, even if hypoxia acts through vulnerability to increasing temperature, incidence of parasitism and disease, or other drivers, our model is general enough to capture declines in community metrics due to implicit drivers.  One may include individual-level impacts of alternative drivers both in addition to, and in place of, the drivers included here to determine whether they are important.
\paragraph{} Do our conclusions hold if we broaden our perspective to other systems?  In the Gulf of Mexico, pelagic prey fishes move into alternative habitats where prey is scarce and the risk of predation is high in response to changes in habitat quality wrought by hypoxia and warming \parencite{zhang2014effects}.  In Lake Erie, demersal Yellow Perch (\textit{Perca flavescens}) move either horizontally or vertically in response to hypoxic bottom waters \parencite{roberts2009effects}.  Though they continue to dive into degraded waters for their benthic prey \parencite{behrens2012effects}, diet composition still shifts towards pelagic sources \parencite{roberts2009effects}.  Differences in susceptibility to hypoxia among species in our simplified Baltic Sea food web serve also to reorganize energy flow.  This has been observed for example in the Neuse River Estuary in North Carolina, where the strength of the escape response and therefore changes in predation pressure vary by species \parencite{bell2005species}.  Looking at lower trophic levels in the Chesapeake Bay, escape response of benthic invertebrates can significantly alter the flow of energy from the sediment to demersal predators \parencite{pihl1992hypoxia}.  Differential responses across species in other community assemblages will likely result in different responses, though even species resistant to hypoxia may still be observed to exhibit similar responses (e.g. diet shifts in Atlantic croaker; \cite{mohan2016out,steube2021distributive})  Therefore, there seems to be broad support for our conclusions across systems.
\paragraph{} Overall, our study suggests that declining benthic food availability and escape of demersal predators towards pelagic habitats are important pathways by which persistent and severe hypoxia affects community dynamics.  These ecological consequences can help to inform forecasts of biomass, yield, and growth of component species as hypoxia stagnates, worsens, or, more optimistically, as environmental conditions improve.  Failing to account for the potential impacts of deoxygenation may lead to erroneous stock assessments and poor management advice \parencite{rose2019impacts}.  For example, ignoring changes in natural mortality caused by increased predation by displaced benthopelagic predators on pelagic prey fishes could introduce significant bias into biomass estimates and therefore into recommended quotas \parencite{clark1999effects}.  Resultant changes in community size structure, and uncertainty therein introduced by developing stressors like declining oxygen and rising temperatures, are necessary components of projections used in the decision-making process \parencite{reum2020ensemble}.  While our approach is only a first step towards partitioning observed fish community dynamics among environmental impacts, it may serve as a useful baseline with which to rigorously evaluate various precautionary approaches to fishing in communities burdened by hypoxic waters.

\subsection*{Conflict of Interest Statement}

The authors declare there are no conflicts of interest.

\subsection*{Acknowledgements}

We are grateful to Gustav Delius and Ken Andersen for discussions regarding proper use of \textbf{mizer}.  We would also like to thank Max Lindmark for his guidance on using \textbf{mizer} to investigate environmental effects on food webs.  The entire SPECTRE reading and discussion group encouraged our use and understanding of size spectrum models more generally.  This work was supported by funding for Project Breathless (NSF OCE1923965; to KL, MC and AG).  Additional funding was provided by the Swedish Research Council Formas (grant no. 2018-00775 to MC).

\clearpage

\begingroup
\raggedright
\printbibliography
\end{refsection}
\endgroup
\clearpage

\section*{Figure Captions}

\paragraph{Figure 1.} A conceptual illustration of key components and outcomes of the oxygen-dependent size spectrum food web model.  Quantitative descriptions of food- and size-dependent growth (a), size-based feeding preference (b), density dependent reproduction (c), fisheries size selectivity (d), and size-dependent maturation determine the central processes of growth, reproduction, and mortality.  In this model, metabolism depends on oxygen with a negative exponential function (e), whereas other physiological processes scale with oxygen according to a logistic function.  Together, these processes produce an emergent number and mass of individuals in the population (size spectra) of each of the interacting fish species (g).

\paragraph{Figure 2.} Projections of body growth for cod during the projection period (2001--2019) arising from the best model (BOM i.e. oxygen dependence of benthic food, cod occupancy in the benthic habitat, and mortality), all possible combinations of its components, as well as the model with no oxygen dependence (None).  Data are included as small, black triangles.  Dotted, light blue lines are von Bertalanffy growth estimates from each year.  Note that those lines either with or without oxygen dependence of benthic food availability (B), typically fall nearly beneath or above one another.

\paragraph{Figure 3.} Projections of biomass of mature size classes (SSB) throughout the calibration period (1991--2000; shaded region) and the projection period (2001--2019).  Model abbreviations as in Figure {\ref{fig:cod_growth_eight}}.  Blue triangles are observations (as estimated from stock assessments; see Methods in main text).

\paragraph{Figure 4.} Projections of yield at observed fishing mortality, throughout the calibration period (1991--2000; shaded region) and the projection period (2001--2019).  Model abbreviations as in Figure {\ref{fig:cod_growth_eight}}.  Symbols as in Figure \ref{fig:sim_ssb}.

\paragraph{Figure 5.} Modeled spawning stock biomass (SSB; kt), yield (kt) at mean fishing mortality observed during the calibration period (1991--2000), and size (g) of individuals at maximum observed age for all species (cod, 15; flounder, 26; sprat, 16; herring, 13) under three oxygen scenarios applied in the best model (BOM) and all possible combinations of its components.  Model abbreviations as in Figure {\ref{fig:cod_growth_eight}}.  We also included model results with no oxygen-dependence (None; for which results were identical across oxygen scenarios).

\paragraph{Figure 6.} Changes in proportion of each prey item consumed by cod across body sizes as oxygen-dependent benthic food availability (B) and cod benthic occupancy (O) change with declines in ambient oxygen concentration.  The BOM model is used, but note that mortality (M) does not vary with oxygen level.

\paragraph{Figure 7.} Changes in energy flow among community members as oxygen-dependent benthic food availability (B) and cod occupancy of the benthic habitat (O) change with declines in ambient oxygen concentration.  The BOM model is used, but note that mortality (M) does not vary with oxygen level.  The position of the cod relative to the center black line is calculated from their occupancy of the benthic versus the pelagic habitat, while length and height of the fish drawings reflect declines in body length and weight (g), respectively.  The arrows reflect energy flow.  Arrow thickness is proportional to the natural log of flow in g\(\cdot\)year\textsuperscript{-1}.  Arrow colors represent changes in flow on the linear scale in each scenario relative to those in the 3 mL\(\cdot\)L\textsuperscript{-1} scenario.  On this scale, red shades represent declines, while blues represent increases.

\clearpage

\begin{figure}[ht]
    \centering
    \includegraphics[scale=0.5]{figures/conceptual.jpg}
    \caption{}
    \label{fig:concept}
\end{figure}
\clearpage

\begin{figure}
    \centering
    \includegraphics[scale=0.6]{figures/cod_growth_eight_best.jpg}
    \caption{}
    \label{fig:cod_growth_eight}
\end{figure}
\clearpage

\begin{figure}[ht]
    \centering
    \includegraphics[scale=0.6]{figures/sim_biomass_best.jpg}
    \caption{}
    \label{fig:sim_ssb}
\end{figure}
\clearpage

\begin{figure}[ht]
    \centering
    \includegraphics[scale=0.6]{figures/sim_yield_best.jpg}
    \caption{}
    \label{fig:sim_yield}
\end{figure}
\clearpage

\begin{figure}
    \centering
    \includegraphics[scale=0.8]{figures/table_scenarios.jpg}
    \caption{}
    \label{fig:scenarios}
\end{figure}
\clearpage

\begin{figure}
    \centering
    \includegraphics[scale=0.8]{figures/diet_scenarios.jpg}
    \caption{}
    \label{fig:diet_scenarios}
\end{figure}
\clearpage

\begin{figure}[ht]
    \centering
    \includegraphics[scale=0.6]{figures/foodweb_scenarios.jpg}
    \caption{}
    \label{fig:flow_scenarios}
\end{figure}
\clearpage

\setcounter{equation}{0}
\renewcommand\theequation{S.\arabic{equation}}
\begin{refsection}

\renewcommand\thefigure{S\arabic{figure}}
\setcounter{table}{0}
\setcounter{figure}{0}
\renewcommand{\thetable}{S\arabic{table}}

\singlespacing
\title{Supporting Information: Declining food availability and habitat shifts drive community responses to marine hypoxia}
\maketitle

\tableofcontents
%\doublespacing
\clearpage

\subsection*{Appendix S1 Model structure}
\addcontentsline{toc}{section}{Appendix S1 Model structure}

Our novel model structure builds off the baseline provided in \textbf{mizer} \parencite{mizer}.  We introduced oxygen dependence for several model components, including benthic resource carrying capacity, fish benthic occupancy, natural mortality, and several physiological rates.  We describe each of these components, and the impetus for singling them out, in the sections below.

\subsubsection*{Benthic resources}

Benthic resources -- in our case, composed primarily of benthic invertebrate prey for cod and flounder species -- are typically more tolerant of hypoxia than their benthopelagic predators \parencite{casini2016hypoxic}.  For example, \textit{Saduria entomon}, one of the Baltic cod's primary benthic food sources \parencite{haahtela1990baltic}, does not experience reduced abundance until oxygen falls below 1 mL\(\cdot\)L\textsuperscript{-1}, well below cod's threshold \parencite{gray2002effects}.  Initially, benthic invertebrates may provide an increasingly abundant food source as low oxygen levels encourage escape to the water column \parencite{lowell1999cumulative,eby2005habitat}.  As hypoxia develops, biomass of benthic invertebrates also declines, exacerbating competition as primary production is largely redirected to the microbial loop \parencite{baird2004consequences}.  Invertebrates may reallocate resources away from reproduction and towards survival strategies, such as increasing ventilation, decreasing both turnover rate and carrying capacity \parencite{calder2015mesocosm}.  We chose to scale carrying capacity of the benthic resource spectrum with equation \ref{eq:scaling} (see main text), which we rewrite here for reference.
    \begin{linenomath*}
    \begin{equation} \label{eq:supp_scaling}
        \xi(O_{iwt}) = \frac{1}{1 + \exp{\left(-U_{i}(O_{iwt}-a_{i}\,\textsf{P}_{\textsf{crit},iw})\right)}}
    \end{equation}
    \end{linenomath*}
Though, lacking values of P\textsubscript{crit} for the benthos, we lumped \(a_{i}\,\textsf{P}_{\textsf{crit},iw}\) into a single value that we call \(k_\textsf{res}\) which, along with \(U_\textsf{res}\), reflected sensitivity of benthic resources to declining oxygen.

\subsection*{Occupancy}

As environmental hypoxia develops, fish may alter their spatial distribution within a given system.  This may result in changes in overlap and thus in species interactions, leading to altered diets \parencite{hrycik2017sub}.  Evidence suggests that depth distribution of cod, for example, has a complex relationship with environmental factors \parencite{orio2019spatial}.  They do not completely avoid hypoxic waters, but rather make brief, frequent visits to deeper waters, likely to forage \parencite{limburg2018effect,neuenfeldt2009some}.  As hypoxia increases in frequency, fish mortality may decrease as avoidance behaviors increase, given an alternative habitat is available \parencite{diaz2009hypoxic}.
\paragraph{} We chose to represent avoidance of hypoxia by scaling probability of benthic occupancy, alternatively framed as proportion time spent in benthic habitat (equation \ref{eq:supp_scaling}).  In our case, this would be for cod only, as we consider sprat and herring to be entirely pelagic and flounder to be entirely benthic.  We distinguished between the effects of ambient oxygen exposure on habitat choice from the effects on the scaling of physiological rates to represent the trade-off between exposure and access to feeding grounds.
\paragraph{} As a baseline, we calculated horizontal overlap of each fish species with each other, called \(\theta_{ij}\), from spatially indexed catch per unit effort (CPUE) from bottom trawl surveys for cod and flounder, and acoustic survey estimates for sprat and herring (Appendix S2).  We calculated vertical overlap of species as a function of their occupancy calculated within the model.  The vertical overlap of species \(i\) with species \(j\) can be represented by the probability of any individual of species \(i\) occupying the same habitat as any individual of species \(j\).  If we assume the distribution of fish within benthic and pelagic habitats is homogeneous, and that vertical occupancy of a given species is independent of the occupancy all other species, then we can write the probability of co-occurrence as follows:
    \begin{linenomath*}
    \postdisplaypenalty = 0
    \begin{align*}
        P(i \cap j) &= P((i \in \textsf{benthic} \cap j \in \textsf{benthic}) \cup  (i \in \textsf{pelagic} \cap j \in \textsf{pelagic})) \\
        &= P(i \in \textsf{benthic} \cap j \in \textsf{benthic}) + P(i \in \textsf{benthic} \cap j \in \textsf{benthic}) \\
        &= P(i \in \textsf{benthic}) P(j \in \textsf{benthic}) + P(i \in \textsf{pelagic}) P(j \in \textsf{pelagic}) \\
        &= \psi_{it} \psi_{jt} + (1-\psi_{it})(1-\psi_{jt})
    \end{align*}
    \end{linenomath*}
So, \(\theta_{ijt}\), or the overlap of species \(i\) with species \(j\) at time \(t\), is then:
    \begin{linenomath*}
    \begin{equation} \label{eq:vertical_overlap}
        \theta_{ijt} = \theta_{ij}\left( \psi_{it} \psi_{jt}+(1-\psi_{it})(1-\psi_{jt})\right)
    \end{equation}
    \end{linenomath*}
Given that occupancy is size dependent, it is more accurate to write \(\theta_{i_{w}j_{w_{p}}}\), where \(w\) and \(w_{p}\) are predator and prey weight, respectively.  This product essentially amounts to a three-dimensional representation of overlap between species.  While, realistically, deleterious effects of low oxygen levels may require relatively prolonged episodes of exposure, we calculated exposure, referred to as \(O_{iwt}\) in equation \ref{eq:supp_scaling}, as a simple weighted average of the probability of occupancy within the benthic and pelagic habitats.  This exposure determines the degree of scaling of mortality and physiology.  We fit the parameters describing sensitivity of occupancy to oxygen, referred to as \(U_{hab}\) and \(a_{hab}\), to the ratio of benthic to pelagic CPUE of cod.

\subsection*{Mortality}

Direct mortality due to exposure to hypoxia has been documented through observations of fish kills and experimental exposure \parencite[e.g.][]{diaz2001overview,shimps2005hypoxia,small2014hypoxia}.  We included potential increases in mortality in hypoxic waters via hazard functions of the form:
    \begin{linenomath*}
    \begin{equation} \label{eq:hazard}
        h(t) = b_{0,i}e^{b_{i}O_{iwt}}
    \end{equation} 
    \end{linenomath*}
where \(O_{iwt}\) is, as above, average oxygen level experienced by an individual of species \(i\) of size \(w\) at time \(t\), \(b_{0,i}\) gives mortality at zero oxygen and \(b_{i}\leq0\) reflects sensitivity of mortality to oxygen.  The primary outcome of adjusting natural mortality in the \textbf{mizer} framework is the death of larger individuals, which is typically used to adjust modeled levels of yield.  Therefore, we fit \(b_{0,i}\) and \(b_{i}\) for cod and flounder to annual observations of yield.

\subsection*{Physiology}

Dependence of physiological rates on oxygen consists of both increases in metabolic costs and general decreases of activity with decreasing oxygen levels.  We describe these pieces together as all physiological parameters are fit simultaneously during the calibration procedure.  Scaling of metabolic costs is determined by parameters that we call \(U_{\textsf{met}}\) and \(a_{\textsf{met}}\) of equation \ref{eq:met_scaling} (see main text), again repeated here:
\begin{linenomath}
    \begin{equation} \label{eq:suppmet_scaling}
        \xi(O_{iwt}) = 1 + \exp{(-U_{\textsf{met},i}(O_{iwt}-a_{\textsf{met},i}\,\textsf{P}_{\textsf{crit},iw}))}
    \end{equation}
    \end{linenomath}
All other physiological scaling is determined by \(U_{\textsf{phys}}\) and \(a_{\textsf{phys}}\) of equation \ref{eq:supp_scaling}.  These are all fit to observations of annual growth of all species.

\subsubsection*{Metabolism}

As ambient oxygen levels fall, fish must expend more energy to maintain internal oxygen, whether through increased ventilation, movement, aquatic surface respiration, or other compensatory behaviors \parencite{kramer1987dissolved}.  This may have an indirect effect on the availability of energy for other activities.  When oxygen falls to levels low enough to stimulate the onset of anaerobic glycolysis, individuals may incur additional metabolic costs.  Though hypoxia tolerant fish may suppress metabolic rate at very low oxygen levels prior to the onset of anaerobiosis \parencite{muusze1998hypoxia}, they too will eventually incur these same costs.  Metabolic costs in \textbf{mizer} are a product of the following: critical feeding level \(f_{c}\), or the feeding level below which intake is less than metabolic costs; maximum consumption \(h_{iwt}\); and assimilation efficiency \(\alpha_{iwt}\) \parencite{mizer}.  Scaling of metabolic costs indicate that consequences of exposure to hypoxia increase exponentially as oxygen continues to decline.

\subsubsection*{Feeding behavior}

The \textbf{mizer} framework assumes feeding follows a Type II functional response, which is determined by encounter rates and maximum consumption rates, the former of which is in turn a function of clearance rates and prey size preference \parencite[see][Table 1]{andersen2016theoretical}.  One of the most commonly cited causes of growth rate reductions in response to hypoxia is the loss of appetite \parencite{wang2009effects}.  In fact, fish exposed to hypoxia and fish fed a reduced diet may appear quite similar \parencite{pichavant2001comparative}.  Overall, energy acquisition declines both as intake declines and as specific dynamic action (SDA), or the costs of digestion and assimilation, increase.  Cod, for example, consume large meals, followed by a period of fasting, during which SDA may account for up to 90\% of maximum metabolic rate (MMR) \parencite{wang2009effects}.  Therefore, reductions in feeding may free a significant portion of the aerobic scope.  We down-scaled maximum consumption to reflect a decline in feeding with falling oxygen levels.
\paragraph{} Behavioral responses to vary with the severity of hypoxia, and with the natural character of a given species in normoxia.  Relatively sedentary benthic and benthopelagic species may decrease activity to conserve oxygen, whereas more active and pelagic species may increase activity \parencite{chapman2009behavioral}.  Cod, for example, are active benthopelagic predators, and will initially increase activity in response to developing hypoxia, perhaps as a means of escaping unfavorable conditions \parencite{chapman2009behavioral}.  However, as oxygen continues to decline, activity tends to decrease.  Thus, we also down-scaled search or clearance rate with decreasing oxygen in an attempt to capture changes in feeding activity.
\paragraph{} The \textbf{mizer} framework also includes a single assimilation value for all species at all sizes.  Assimilation efficiency may decline in response to hypoxia, though some studies show no effect \parencite{wang2009effects}.  \textcite{jordan2007effects} showed that exposure to hypoxic conditions increases the amount of time required for digestion and decreases the overall energy acquisition from digestion of meals.  Though evidence is somewhat mixed, we chose to account for increased costs of feeding with declining assimilation efficiency when oxygen is low.

\subsubsection*{Reproduction efficiency}

Hypoxia affects fish reproduction through two major pathways.  First, exposure to low oxygen may disrupt hormones and enzymes which regulate reproduction and development of eggs, leading to egg mortality.  In our study system, cod eggs are neutrally buoyant at salinities of 10-12 psu, and may therefore sink below the halocline, resulting in exposure to hypoxic waters, altered development, and ultimately mortality \parencite{baird2004consequences}.  Second, declines in energy acquisition due to decreased activity may reduce energy available for reproduction \parencite{wu2009effects}.  Declining energy acquisition is addressed by oxygen dependence of feeding behavior.  We included egg mortality by scaling reproductive efficiency (see \textcite{mizer}) as a logistic function of oxygen, which is a measure of egg survival or rather production.

\subsection*{Appendix S2 Parameter estimation}
\addcontentsline{toc}{section}{Appendix S2 Parameter estimation}

We parameterized the model with several data sources (Table \ref{tab:data}).  These parameters describe growth, maturation, selectivity, feeding preferences, and species interactions.  We used the years 1991-2000 as a calibration period.  The ideal calibration period is one which is comprised of several years during which conditions were stable.  In our case, conditions within the Baltic Sea were deteriorating during the years 1991-2000.  However, 1991 is the earliest year for which data are available for all species for nearly all model components, excepting the European and Baltic flounders (\textit{Platichthys flesus} and \textit{Platichthys solemdali}, respectively), for which there are limited data on stomach contents, weight, length, age, and maturity.  For these fish, we used all available data for calibration, which correspond to the mid-2000's through the mid-2010's.  This introduces the implicit assumption that flounder life history was identical throughout this period and in 1991-2000.  Though perhaps unfounded, we are limited by the scope of the data.

\begin{table}[h]
    \centering
    \tiny
    \begin{threeparttable}
    \caption{Model parameters and data sources for each species.  CPUE is catch per unit effort, w/l/a/m are weight/length/age/maturity, and F is fishing mortality.}
    \label{tab:data}
    \begin{tabular}{lccccr}
        \toprule
        \textbf{Parameters} & \textbf{Description} & \textbf{Data} & \textbf{Species} & \textbf{Years} & \textbf{Source} \\
        \midrule
        \(\beta,\sigma\) & Mean and SD of predator-prey mass ratios & stomach & \begin{tabular}{@{}c@{}} \textit{Gadus morhua} \\ \textit{Platichthys flesus} \\ \textit{Clupea harengus} \\ \textit{Sprattus sprattus} \end{tabular} & \begin{tabular}{@{}c@{}} 1963-2014\tnote{a}\,,2015-2018\tnote{b} \\ 2015-2018\tnote{b} \\ 1999-2000\tnote{c} \\ 1999-2000\tnote{c} \end{tabular} & \begin{tabular}{@{}r@{}} a,b \\ b \\ c \\ c \end{tabular} \\
        \midrule
        \(\theta_{ij}\) & Horizontal degree to which species \(i\) and \(j\) overlap &  CPUE/abundance & \begin{tabular}{@{}c@{}} \textit{G. morhua} \\ \textit{P. flesus} \\ \textit{C. harengus} \\ \textit{S. sprattus} \end{tabular} & \begin{tabular}{@{}c@{}} 1991-2020\tnote{d} \\ 1991-2020\tnote{d} \\ 1979-2020\tnote{e} \\ 1979-2020\tnote{e} \\  \end{tabular} & \begin{tabular}{@{}r@{}} d \\ d \\ e \\ e \end{tabular} \\
        \midrule
        \(w_{m}\) & Weight at which probability of maturation is \(0.5\) & w/l/a/m & \begin{tabular}{@{}c@{}} \textit{G. morhua} \\ \textit{P. flesus} \\ \textit{C. harengus} \\ \textit{S. sprattus} \end{tabular} & \begin{tabular}{@{}c@{}} 1991-2020\tnote{d} \\ 2007-2015\tnote{d} \\ 1979-2020\tnote{e} \\ 1979-2020\tnote{e} \\  \end{tabular} & \begin{tabular}{@{}r@{}} d \\ d \\ e \\ e \end{tabular} \\
        \midrule
        \(W_{\infty},k_{vb},t_{0},a,b\) & von Bertalanffy growth and length-weight conversion & w/l/a & \begin{tabular}{@{}c@{}} \textit{G. morhua} \\ \textit{P. flesus} \\ \textit{C. harengus} \\ \textit{S. sprattus} \end{tabular} & \begin{tabular}{@{}c@{}} 1991-2020\tnote{d} \\ 2007-2015\tnote{d} \\ 1979-2020\tnote{e} \\ 1979-2020\tnote{e} \\  \end{tabular} & \begin{tabular}{@{}r@{}} d \\ d \\ e \\ e \end{tabular} \\
        \midrule
        \(L_{25},L_{50}\) & Lengths at 25\% and 50\% selectivity & F & \begin{tabular}{@{}c@{}} \textit{G. morhua} \\ \textit{P. flesus} \\ \textit{C. harengus} \\ \textit{S. sprattus} \end{tabular} & \begin{tabular}{@{}c@{}} 1946-2019\tnote{f} \\ -- \\ 1974-2019\tnote{f} \\ 1974-2019\tnote{f} \\  \end{tabular} & \begin{tabular}{@{}r@{}} f \\ -- \\ f \\ f \end{tabular} \\
        \midrule
        \(L_{k\mbox{-}e}\) & Lengths at 100\% selectivity, or knife-edge & F & \begin{tabular}{@{}c@{}} \textit{G. morhua} \\ \textit{P. flesus} \\ \textit{C. harengus} \\ \textit{S. sprattus} \end{tabular} & \begin{tabular}{@{}c@{}} -- \\ 1955-2004\tnote{g} \\ -- \\ -- \\  \end{tabular} & \begin{tabular}{@{}r@{}} -- \\ g \\ -- \\ -- \end{tabular} \\
        \bottomrule
    \end{tabular}
    \begin{tablenotes}
        \item[a] ICES Year of the Stomach; \textcite{YoTSdata2020}
        \item[b] \textcite{haase2020diet}
        \item[c] \textcite{casini2004feeding}
        \item[d] ICES CPUE from BITS survey database DATRAS \textcite{datrasdata2020}
        \item[e] ICES BIAS acoustic survey estimates; \parencite{wgbifs2020}
        \item[f] \parencite{wgbifs2020}
        \item[g] \textcite{draganik2007status}
    \end{tablenotes}
    \end{threeparttable}
\end{table}

\subsubsection*{Growth}

We estimated body growth parameters for all four species using the inverse of the von Bertalanffy function:  
    \begin{linenomath*}
    \begin{equation} \label{eq:ivb}
        f(L) = -\frac{1}{k_{vb}}\ln{\left(1 - \frac{L}{L_{\infty}}\right)} + t_{0}
    \end{equation}
    \end{linenomath*}
where \(L\) is the length of a given individual, \(k_{vb}\) is the von Bertalanffy growth factor, \(L_{\infty}\) is the asymptotic length, \(t_{0}\) is age at size zero, and \(f(L)\) is estimated age.  We extracted sex-maturity-age-length-key (SMALK) data from \href{https://datras.ices.dk/}{DATRAS} \parencite{datrasdata2020} and used size and age observations for cod and flounder.  For sprat and herring, we used size and age observations from BIAS trawl surveys \parencite{wgbifs2020}.  The data (Table \ref{tab:data}) are given as weight and age of individuals within a given size class.  Data are also given for a relatively wide range of size classes for each species.  Therefore, the inverse von Bertalanffy is ideal \parencite{mackay1990note}.  Fish at age zero were collected roughly half a year post-spawning, thus we added 0.5 to reported age and assumed size at birth, or \(t_{0}\), was equal to zero, or close enough to zero to warrant dropping the parameter.
\paragraph{} We chose to run all models for each species in a Bayesian framework within \textbf{Stan} \parencite{carpenter2017stan}.  We used the following priors:
    \begin{linenomath*}
    \postdisplaypenalty = 0
    \begin{align*}
        k_{vb} &\sim \textsf{Uniform}(0,5) \\
        L_{\infty} &\sim N(\mu_{L_{\infty}},\sigma_{L_{\infty}}^{2}) \, T[L_{\textsf{min}},] \numberthis \\
        \sigma_{\textsf{age}}^{2} &\sim \textsf{Cauchy}(0,1000)
    \end{align*}
    \end{linenomath*}
where \(\mu_{L_{\infty}}\) and \(\sigma_{L_{\infty}}^{2}\) were the mean and variance of the \(L_{\infty}\) prior distribution for any given species, \(L_{\textsf{min}}\) was the minimum possible value for \(L_{\infty}\), and \(\sigma_{\textsf{age}}^{2}\) was a hyperparameter describing variance in estimated ages (see equation \ref{eq:ivb_priors}).  The addition of a \(T[x,]\) or \(T[,x]\) to a distribution indicates that it is truncated from below or above, respectively.  We used the following likelihood:
    \begin{linenomath*}
    \begin{equation} \label{eq:ivb_priors}
        \textsf{age} \sim N\left(-\frac{1}{k_{vb}}\ln{\left(1 - \frac{L}{L_{\infty}}\right)}, \sigma_{\textsf{age}}^{2}\right)
    \end{equation}
    \end{linenomath*}
During implementation, we chose \(\mu_{L_{\infty}}=100\) for all species, which implies that the resulting prior is only moderately informative.  We chose \(\mu_{L_{\infty}}\) as the maximum observed length class for each species, plus 1 mm.  Similarly, we chose \(L_{\textsf{min}}\) as the maximum observed length class.  This prevented failures in the model due to attempting to take the natural log of a negative number.  The \textbf{mizer} framework requires body size in grams, and therefore we estimated length-weight relationships using nonlinear least squares for each species with the following standard equation: 
    \begin{linenomath*}
    \begin{equation} \label{eq:lengthweight}
        W = aL^{b}
    \end{equation}
    \end{linenomath*}
Results were typical, and apparent outliers had little to no observable effect on estimates (Figure \ref{fig:all_lengthweight}). 

\begin{figure}[h]
    \centering
    \includegraphics[scale=0.6]{figures/all_lengthweight.jpeg}
    \caption{Length-weight relationships for each species.  The solid red lines give mean predictions.}
    \label{fig:all_lengthweight}
\end{figure}

\paragraph{} We ran the Bayesian model on size-at-age data for all four species.  For cod and flounder, we partitioned the data sets by ICES subdivision, sex, and year.  For sprat and herring, we partitioned the data sets by ICES subdivision and year.  The former two exhibit sexual dimorphism while the latter do not.  We ran the model separately on each data set, excluding those data sets with fewer than 25 points, which resulted in a total of 193 models across all species.  We ran each model for 2,000 samples as warmup, followed by 20,000 iterations over four chains running in parallel, resulting in 80,000 samples from each model.  Models finished anywhere from 3 to 147 seconds, with a median time of 11 seconds, a mean of 19 seconds, and a total of 62 minutes.  Note that we increased the default maximum tree depth to 20, and increased the target average acceptance probability to \(0.99\) so that all models would finish with no divergent transitions \parencite{carpenter2017stan}.  All models finished with more than 100 effective samples for both the bulk and the tail, indicating adequate sampling across posterior quantiles.  We took the area-weighted median of each estimate to arrive at final estimates for each species (Table \ref{tab:allparams}).  Resulting predictions of length-at-age agree fairly well with observations (Figure \ref{fig:all_agelength}).  Apparent biases are due largely to data partitioning and area weighting.

\begin{table}[h]
    \centering
    \scriptsize
    \caption{All parameter estimates for all four species, when applicable.  See parameter descriptions in Table \ref{tab:data}.}
    \begin{tabular}{llccccc}
        \toprule
        \textbf{Parameter} & \textbf{Description} & \textbf{Unit} & \textbf{Cod} & \textbf{Flounder} & \textbf{Sprat} & \textbf{Herring} \\
        \midrule
        \(\beta\) & Mean preferred predator prey mass ratio & -- & 673 & 937 & 173302 & 702819  \\
        \(\sigma\) & SD preferred predator prey mass ratio & -- & 1.63 & 1.70 & 0.74 & 0.52 \\
        \(w_{m}\) & g & 577 & 64 & 7 & 20 \\
        \(L_{\infty}\) & Maximum length & mm & 1243 & 447 & 157 & 324 \\
        \(W_{\infty}\) & Maximum weight & mm & 17918 & 1239 & 21 & 213 \\
        \(k_{vb}\) & von Bertalanffy growth & y\textsuperscript{-1} & 0.13 & 0.12 & 0.43 & 0.22 \\
        \(a\) & Length-weight coefficient & 10\textsuperscript{-5} & 1.99 & 0.48 & 4.09 & 0.34 \\
        \(b\) & Length-weight exponent & -- & 2.89 & 3.18 & 2.60 & 3.11 \\
        \(L_{25}\) & Length at 25\% selectivity & mm & 402 & -- & 121 & 140 \\
        \(L_{50}\) & Length at 50\% selectivity & mm & 489 & -- & 130 & 169 \\
        \(L_{k\mbox{-}e}\) & Knife-edge selectivity & mm & -- & 180 & -- & -- \\
        \(F_{\textsf{max}}\) & Maximum fishing mortality & y\textsuperscript{-1} & 1.00 & 0.90 & 0.34 & 0.47 \\
        \(F_{k\mbox{-}e}\) & Knife-edge fishing mortality & y\textsuperscript{-1} & -- & 0.90 & -- & -- \\
        \bottomrule
    \end{tabular}
    \label{tab:allparams}
\end{table}

\begin{figure}
    \centering
    \includegraphics[scale=0.6]{figures/all_agelength.jpeg}
    \caption{Length-at-age data for each species.  Solid, red lines give model estimates.  Note that while data are pooled here, estimates arise from data partitioning (see text).}
    \label{fig:all_agelength}
\end{figure}

\subsubsection*{\textbf{Maturation}}

We calculated probabilistic maturation reaction norms \parencite[PMRN;][]{dieckmann2007probabilistic} for all four species.  The PMRN attempts to remove influences of growth and mortality on maturity, thereby isolating variability in size at maturation \parencite{barot2003estimating}.  We included age, length, and Fulton's condition factor (\(W/L^{3}\)) in our model, as in \textcite{vainikka2009two}.  In our case, then, the PMRN is given by:
    \begin{linenomath*}
    \begin{equation} \label{eq:pmrn}
        m(a,s,c) = \frac{o(a,s,c)-o(a-1,s-\Delta s,c-\Delta c)}{1-o(a-1,s-\Delta s,c-\Delta c)}
    \end{equation}
    \end{linenomath*}
where \(\Delta s\) and \(\Delta c\) refer to annual increments in weight and condition factor, respectively. Probability of maturation at a given age, size, and condition factor is thus conditional on those individuals who have already matured in the previous time step, an extension of the relationship described by \parencite{barot2003estimating}.
\paragraph{} We extracted data on maturation from the BITS survey database \href{https://datras.ices.dk/}{DATRAS}.  For cod and flounder, we downloaded sex-maturity-age-length-key (SMALK) data, and used the ICES maturity key to determine whether or not each individual was mature, converting from the Maier scale to the ICES scale when necessary \parencite{grygiel2002sexual,bitsmanual2017}.  On the ICES scale, fish in stage 1 are immature, and the remaining stages are mature.  For sprat and herring, we used BIAS trawl data, and again categorized fish based on the ICES scale.  For all four fish, we used von Bertalanffy growth function parameters to calculate growth increments, and simple linear models to estimate body condition increments.  We then modelled maturity ogives as a logistic function of age, weight, and condition, and all possible interactions.  We performed simple stepwise model selection using AIC as criteria.  The models retained the following covariates: for cod, age, weight, condition, and weight*condition; for flounder, all covariates and possible interactions; for sprat, only age and weight; and for herring, all covariates besides age*weight*condition.
\paragraph{} We used age estimates, weight and condition increments, and ogives to calculate probability of maturation, or PMRN, for each individual of each species as in equation \ref{eq:pmrn}.  We only used individuals in age groups over which each species typically matures: ages 2 to 3 for cod, 1 to 2 for flounder, 1 to 2 for sprat, and age 1 for herring.  Finally, we used a sigmoid function to model PMRN as a function of weight:
    \begin{linenomath*}
    \begin{equation} \label{eq:sigmat}
        \textsf{PMRN} = \left[1 + \left(\frac{L}{L_{50}}\right)^{-10}\right]^{-1}
    \end{equation}
    \end{linenomath*}
where \(L\) is length of an individual, and where \(L_{50}\), the parameter of interest, is the length at which the probability of maturation is \(0.5\).  The exponential value of -10 ensures that the model is not overwhelmed by data concentration at larger sizes, at which fish have a higher probability of maturation.  When we chose to estimate the exponent of equation \ref{eq:sigmat}, the model treated smaller fish as outliers.  This somewhat arbitrary choice asserts our knowledge that this small pool of immature fish are critical data (Figure \ref{fig:all_pmrn}).  We converted length to weight values using each species' length-weight parameters (Table \ref{tab:allparams}) to arrive at weight at maturity.

\begin{figure}[h]
    \centering
    \includegraphics[scale=0.6]{figures/all_pmrn.jpeg}
    \caption{Estimated probability maturation reaction norms (PMRN) for individuals of each species.  We only included individuals of ages 2 to 3 for cod, 1 to 2 for flounder, 1 to 2 for sprat, and age 1 for herring.  Solid red lines are fits to equation \ref{eq:sigmat}.}
    \label{fig:all_pmrn}
\end{figure}

\subsubsection*{\textbf{Selectivity}}

We used estimates of fishing mortality at age from stock assessments \parencite{wgbifs2020} to estimate selectivity parameters for cod, sprat, and herring.  We used the following ogive function:
    \begin{linenomath*}
    \begin{equation} \label{eq:matogive}
        S(L) = \frac{1}{1 + \exp\left({\frac{L_{50}\ln{(3)}}{L_{50}-L_{25}} - \frac{\ln{(3)}}{L_{50}-L_{25}}L}\right)}
    \end{equation}
    \end{linenomath*}
where \(S(L)\) is selectivity -- or normalized fishing mortality -- as a function of length, and \(L_{25}\) and \(L_{50}\) are lengths at 25\% and 50\% selectivity.  We converted age to length using von Bertalanffy parameters (Table \ref{tab:allparams}).  We retained the maximum estimated fishing mortality for each species and treated length-specific fishing mortality as a product of length-specific selectivity and maximum fishing mortality.
\paragraph{} Currently, the ICES stock assessment lacks fishing mortality estimates for flounder.  \textcite{draganik2007status} estimated fishing mortality of Baltic flounder stocks in ICES subdivision 26 using extended survivor analysis (XSA) \parencite{shepherd1999extended}.  Yields in this subdivision tend to be analogous to yields throughout the Baltic \parencite{draganik2007status}.  Therefore, the identified trends may be accurate, though the actual regional value may be less than that reported for subdivision 26, given fishing mortality tends to be lower in subdivisions 24 and 25.  We extracted F\textsubscript{bar} from this study \parencite[see Figure 10 op. cit.]{draganik2007status} using WebPlotDigitizer \parencite{Rohatgi2020}.  These data were available for the years 1955--2004.  For years beyond 2004, we used the terminal-three year mean of F\textsubscript{bar}.  We calculated the mean of F\textsubscript{bar} for the years 1991--2000, and used a knife-edge selection size of 18cm, corresponding to the minimum legal size in Estonia, and in fact the smallest of all minimum legal sizes \parencite{wgbifs2020}.  As above, we treated fishing mortality as a product of selectivity and maximum fishing mortality.  Results indicate that models agree well with observations (Figure \ref{fig:all_selectivity}).

\begin{figure}[h]
    \centering
    \includegraphics[scale=0.6]{figures/all_selectivity.jpeg}
    \caption{Estimated selectivity for all four species.  Data points are normalized estimates of fishing mortality at age, with age converted to length.  Selectivity for flounder is given by a knife-edge, with all individuals at or above 180mm fully selected by the fishery.}
    \label{fig:all_selectivity}
\end{figure}

\subsubsection*{Prey preference}

The \textbf{mizer} framework assumes that prey selection is a function of the ratio of predator and prey weight.  That is, bigger fish eat smaller fish; how much bigger and smaller dependent upon the species in question.  We used a log-normal function to formalize this relationship as a function of a predator of species \(i\) and of size \(w\):    
    \begin{linenomath*}
    \begin{equation} \label{eq:ppmr}
        \phi\left(\frac{w_{\textsf{prey}}}{w}\right) = \exp{\left(\frac{-\ln\left({\beta_{i}(w)\,w_{\textsf{prey}}}-w\right)^{2}}{2\sigma_{i}^{2}}\right)}
    \end{equation}
    \end{linenomath*}
We calculated PPMR from stomach data for each species.  We limited our analyses to prey items in early stages of digestion, and took the ratio of predator size in g to prey size in g.  We calculated the log of these ratios, and extracted the mean (\(\beta\)) and standard deviation (\(\sigma\)) for each species.  While we limited these calculations to data available during the years 1991--2000 for cod, data were only available during 2015--2018 for flounder.  
\paragraph{} For herring and sprat, we used the WebPlotDigitizer \parencite{Rohatgi2020} to extract feeding data from Figure 3 of \textcite{casini2004feeding}.  These data were recorded as percent composition of eight major species or groups of zooplankton, amphipods, and mysids in the diets of several size classes of both herring and sprat.  We used documented wet weights of each prey item to calculate the mean size of prey items appearing in clupeid stomachs \parencite{hernroth1985zoop}.  When wet weights were partitioned among regions, we used data from ``Remaining Baltic Sea''.  For \textit{Temora longicornis}, we used mean wet weight of copepodite stages CIV-CV and CVI in October through December as a typical prey item for sprat, which dominate in the fall \parencite{mollmann2002population}, and the wet weight of stages CI-CIII, CIV-CV, and CVI for herring.  We included mean weight of stages CI-CIII because herring are observed to switch prey sources from larger stages to stage CII when in competition with high densities of sprat \parencite{mollmann2002population}.  For \textit{Bosmina longispina maritima}, \textit{Podon intermedius}, and \textit{Pleopis polyphemoides} (syn. \textit{Podon polyphemoides}) we used the mean wet weight across all size classes for October through December, for lack of more detailed information.  For \textit{Acartia spp.}, we calculated the mean wet weight of all size classes of \textit{Acartia longiremis} from January through March for Herring, and for \textit{Acarctia bifilosa} from April through December for sprat, given that these time frames correspond roughly to seasons during which each species is observed to feed on these copepods \parencite{mollmann1999food}.  The exact choice of time period and species is not critical, as wet weights are similar throughout the year and within \textit{Acartia spp.}.  We used a mean length of 1.75 cm for \textit{Mysis mixta}, based on the size range of 1.5--2.0 cm for mysids in the Baltic Sea \parencite{snoeijs2017patterns}.  For the group labeled ``amphipods'', we used a rough estimate for body length of 0.9cm, based on size range of 0.6--1.2cm \parencite{hill1992seasonal}.  We converted to weight in grams using the relationship: \(\textsf{weight} = 0.01 \cdot \textsf{length}^{3}\).
\paragraph{}  The data extracted from \textcite{casini2004feeding} served as weights with which to calculate the mean of a typical prey item for each size class of herring and sprat.  We converted each size class from length in cm to weight in g using estimated length-weight relationships.  We used the resulting PPMR values from all four species in the models and calculations described above (Figure \ref{fig:all_ppmr} and see Table \ref{tab:allparams}).

\begin{figure}[h]
    \centering
    \includegraphics[scale=0.6]{figures/all_ppmr.jpeg}
    \caption{Frequency plots of PPMR for all four species.  Vertical, red, broken lines give the mean across all predator weights.  The value of \(\sigma\) gives the standard deviation for each species.}
    \label{fig:all_ppmr}
\end{figure}

\subsubsection*{\textbf{Species interactions}}

Species interactions emerge as a function of spatial overlap in both the horizontal and vertical directions.  Vertical overlap is a dynamic model component (see main text), whereas we treated horizontal overlap as parameters.  We input horizontal spatial overlap as an index calculated from data: 
    \begin{linenomath*}
    \begin{equation}
        D(p_{it},p_{jt}) = 1 - \frac{1}{2}\sum_{k}|p_{itk}-p_{jtk}|
    \end{equation} \label{eq:overlap}
    \end{linenomath*} 
where \(p_{itk}\) is the proportion of the total abundance index of species \(i\) in ICES statistical rectangle \(k\) at time t\parencite{blanchard2014evaluating}.  In our case, these abundance indices arose from BITS for cod and flounder, and BIAS acoustic estimates for sprat and herring (Table \ref{tab:data}).  We calculated overlap between each species during all years between 1991 and 2020 for which data were available, and averaged the results.  This resulted in a mean, static interaction matrix.  We multiplied the elements of this matrix by the vertical overlap values \(\theta_{ijt}\) (equation \ref{eq:vertical_overlap}) which are fit to data extracted from \parencite[Figure 6b of][]{casini2019spatio} using Web Plot Digitizer \parencite{Rohatgi2020}.  This gives both vertical and horizontal overlap.

\subsection*{Appendix S3 Oxygen estimates}
\addcontentsline{toc}{section}{Appendix S3 Oxygen estimates}

We extracted oxygen data for the years 1980 through 2020 from the Swedish Meteorological and Hydrological Institute (SMHI; www.smhi.se).  These data arise from hydrological measurements collected during coordinated fish trawl surveys, complemented by regional and national monitoring programs throughout the Baltic Sea \parencite{smhi2020oxygen}.  We used bottle data to construct spatiotemporal depth profiles of oxygen in the ICES statistical rectangles.  We only included rectangles whose maximum depth fell between 50 and 120 m, and observations that fell in the months of April through November.  The former restriction placed an emphasis on benthopelagic habitat suitable for cod and flounder, while the latter included the major growth and spawning months for all four species.
\paragraph{} We used a generalized additive model (GAM) to model oxygen as a function of two-dimensional spatial coordinates, as well as water depth and year.  The observed distribution of oxygen values was positive with non-zero mass at zero oxygen.  Therefore, we used a tweedie distribution in the \textbf{mgcv} package \parencite{wood2016smoothing}.  We modeled oxygen as a function of x-y coordinates with a thin plate spline, and depth and year with a tensor product smoother.  The tensor product allows one to quantify the interaction between two terms on different scales.  All terms were significant (Table \ref{tab:oxygam}).  While the EDF of each smooth term for the default model was close to the  basis dimension, or \textit{k}, we chose not to increase this value, as we wanted to represent very broad trends over time, space, and depth.  Results indicate there were higher values of oxygen in the benthic habitat in the 1980s and early 1990s than in the late 1990s through the present day (Figure \ref{fig:oxy_years}).  We averaged predicted oxygen values for each year over each ICES statistical rectangle from 10 to 40 m for pelagic habitat and from the bottom 10 m for benthic habitat.  The resulting time series was fairly steady at roughly 6 mL\(\cdot\)L\textsuperscript{-1} in the pelagic habitat (Figure \ref{fig:oxy_years}).  Oxygen levels in the benthic habitat varied between about 1 mL\(\cdot\)L\textsuperscript{-1} and 3 mL\(\cdot\)L\textsuperscript{-1}, with the rise in the 1980s reflective of a major Baltic inflow (MBI) event \parencite{schinke1998causes}.

\begin{table}[h]
    \centering
    \scriptsize
    \caption{Results of the generalized additive model (GAM) of oxygen as a two-dimensional smooth function of x-y coordinates, or \textbf{s(x,y)}, and a tensor product smooth of depth and year, or \textbf{te(depth,year)}.  The term \(\sigma_{\mu}\) is standard area and Ref. EDF is reference degrees of freedom.  Bold and starred \textit{p}-values indicate significant portions of the model.  The adjusted \(R^{2}\) was 0.44, with 42.3\% of deviance explained.}
    \begin{tabular}{lcccc}
        \toprule
        \multicolumn{5}{c}{\textbf{Parametric Terms}} \\
        \midrule
        & Coefficient & \(\sigma_{\mu}\) & \textit{T} & \textit{p} \\
        \textbf{Intercept} & 1.74 & 0.001 & 1427 & \textless \textbf{.001*} \\
        \toprule
        \multicolumn{5}{c}{\textbf{Smooth Terms}} \\
        \midrule
        & EDF & Ref. EDF & \textit{F} & \textit{p} \\
        \textbf{s(x,y)} & 23.87 & 24 & 1759 & \textless \textbf{.001*} \\
        \textbf{te(depth,year)} & 28.39 & 29 & 457 & \textless \textbf{.001*} \\
        \bottomrule
    \end{tabular}
    \label{tab:oxygam}
\end{table}

\begin{figure}[h]
    \centering
    \includegraphics[scale=0.6]{figures/oxygen.jpeg}
    \caption{Predicted mean oxygen level in benthic and pelagic habitat from 1980 through 2020.}
    \label{fig:oxy_years}
\end{figure}

\subsection*{Appendix S4 P\textsubscript{crit}}
\addcontentsline{toc}{section}{Appendix S4 P\textsubscript{crit}}

We estimated P\textsubscript{crit} for each species at a given weight using the database provided by \textcite{pcritdatabase2021}.  This database contains several hundred measurements of P\textsubscript{crit} extracted from the literature, along with other experimental conditions such as water temperature and salinity, and individual attributes such as body mass.  As in \textcite{rogers2016new} we performed stepwise linear regression to describe P\textsubscript{crit} as a function of \(\ln\)(body mass), salinity, temperature, and \(\ln\)(resting metabolic rate, RMR).  We included all measurements in the database in order to represent very general patterns arising in whole ecosystems.  The stepwise regression retained all four covariates, and resulted in the following function:
    \begin{linenomath*}
    \begin{equation}
        \label{eq:pcrit}
        \textsf{P}_{\textsf{crit}} = 30.854 + 0.027\,\textsf{S} - 0.111\,\textsf{T} + 0.410\,\ln{(w)} + 1.645\,\ln{(\textsf{RMR})}
    \end{equation}
    \end{linenomath*}
where S is salinity in practical salinity units (psu), T is temperature in degrees C, \(w\) is weight in kg, and RMR is routine metabolic rate in mg \ch{O2}\(\cdot\)kg\textsuperscript{-1}\(\cdot\)h\textsuperscript{-1}.  The \textbf{mizer} package \parencite{mizer} calculates metabolic rate as an exponential function of weight with, in our case, a scaling exponent of 0.89 \parencite{jerde2019strong}.  The result is given in units of g\(\cdot\)year\textsuperscript{-1}.  We assumed this baseline value calculated by \textbf{mizer} was equivalent to routine metabolic rate (RMR) and converted to units of mg \ch{O2}\(\cdot\)kg\textsuperscript{-1}\(\cdot\)h\textsuperscript{-1} with the following formula:
    \begin{linenomath*}
    \begin{equation} \label{eq:rmr}
        \textsf{RMR}_{w} = \frac{1000\cdot1.429\cdot1.2905\cdot \textsf{RMR}_{\textsf{mizer}}}{365.25\cdot24\cdot (w/1000)}
    \end{equation}
    \end{linenomath*}
In the numerator of equation \ref{eq:rmr}, \(1.2905\) converts from energy intake in g to L \ch{O2}.  We assumed this biomass had a composition of 60\% protein, 35\% fat, and 5\% cholesterol, and used these values to calculate a weighted average of L \ch{O2} required for metabolism from values provided in \textcite{nelson2004biological}.  Also in the numerator, \(1.429\) converts from L \ch{O2} to g \ch{O2}, and finally \(1000\) converts from g to mg.  In the denominator, \(w/1000\) gives each size bin in kg, while \(24\) and \(365.25\) convert from years to hours.  Finally, we converted the resulting P\textsubscript{crit} values from kPa to mL\(\cdot\)L\textsuperscript{-1} using Henry's Law, correcting for a reference temperature of 292.15 K with the van't Hoff equation.  In general, the final P\textsubscript{crit} values increase with weight, though the magnitude differs by species (Figure \ref{fig:pcrit_size}).  This is largely dependent upon the different values of RMR, driven by differences in maximum feeding rate, driven in turn by age and weight at maturation and feeding levels \parencite[see][]{mizer}.  Faster growth rates relative to age at maturation result in higher maximum intake rates, and ultimately in greater values of RMR at a given size.  Note that the dependence of the values of P\textsubscript{crit} on RMR -- which in the \textbf{mizer} framework is dependent on maximum consumption rate -- implies that it could decrease as oxygen levels decline, given that fish appetites decline with exposure to hypoxia \parencite{chabot2008environmental,wang2009effects}.  While this sort of acclimation is perhaps more realistic in the long term, we chose to fix P\textsubscript{crit} relative to growth in order to focus on shorter term responses to environmental hypoxia, as well as to express the assumption that RMR is constant when oxygen is at or above P\textsubscript{crit} \parencite{wang2009effects}.

\begin{figure}[h]
    \centering
    \includegraphics[scale=0.6]{figures/pcrit.jpeg}
    \caption{Calculated P\textsubscript{crit} as a function of size for all four species included in the model.}
    \label{fig:pcrit_size}
\end{figure}

\paragraph{} We calculated P\textsubscript{crit} for cod and flounder.  At present, the P\textsubscript{crit} of sprat and herring is irrelevant, though we report the results all the same (Figure \ref{fig:pcrit_size}).  Cod are relatively large, active, benthopelagic fish, and we thus expected to have high values of RMR to support a higher maximum metabolic rate \parencite[MMR;][]{killen2016ecological}.  By contrast, flounder are smaller, less active, and largely benthic, and we expected smaller values of RMR.  The results of the conversion reflect these expectations (Figure \ref{fig:pcrit_size}).

\subsection*{Appendix S5 Calibration}
\addcontentsline{toc}{section}{Appendix S5 Calibration}

\subsubsection*{Procedure}

Each model must be calibrated before it can be used for projection and analysis.  One typically calibrates to a period during which the community is relatively stable, and subsequently projects forwards or backwards in time.  This process involves adjusting several of the parameters in order to match observed growth, biomass, and fisheries yield, if available.  However, given the relatively complex, highly interdependent nature of the various structures within the model, adjusting a parameter for one species may result in a changed outcome for all species.  There are currently no universally accepted calibration procedures for models of this type.  
\paragraph{} We chose to combine stepwise, manual parameter adjustment with automated optimization procedures, as in \textcite{lindmark2021bottom}.  We treated the following data sources as observations with which to compare to model output: benthic occupancy, SSB, yield, and von Bertalanffy growth (Table \ref{tab:cal_params}).  Note that cod were the only species that we allowed to move between the pelagic and benthic habitats.  We verified vertical occupancy of cod with ratios of pelagic and benthic trawl CPUE found in \textcite{casini2019spatio}.  For cod, sprat, and herring, we extracted SSB and yield from stock assessments \parencite{wgbifs2020}.  These data are reported for cod in ICES subdivisions 24-32, for sprat in 22-32, and for herring in 25-29 and 32.  Flounder were not assessed, and we therefore assumed that the ratio of flounder and cod CPUE was equal to the ratio of flounder and cod SSB in a given year.  We divided the rough estimates of flounder SSB by the mean ratio of estimated yield -- the annual product of F\textsubscript{bar} \parencite{draganik2007status} and SSB -- to observed yield during the calibration period. The procedures for estimating von Bertalanffy growth parameters are described in Appendix S2 (Table \ref{tab:allparams}). 

\begin{table}[h]
    \centering
    \scriptsize
    \caption{All fixed (above) and tuning (below) parameters that may be adjusted during the course of model calibration.  Species-specific parameters are marked with the subscript \(i\).}
    \begin{tabular}{lcl}
        \toprule
        \multicolumn{3}{c}{\textbf{Fixed}} \\
        \midrule
        Parameter & Value & Description \\
        \cmidrule(r){1-1} \cmidrule(lr){2-2} \cmidrule(l){3-3}
        \(p\) & 0.89 & Metabolic scaling exponent \\
        \(q\) & 0.89 & Predation or search rate scaling exponent \\
        \toprule
        \multicolumn{3}{c}{\textbf{Tuning}} \\
        \midrule
        Parameter & Value & Description \\
        \cmidrule(r){1-1} \cmidrule(lr){2-2} \cmidrule(l){3-3}
        \(R_{\textsf{max},i}\) & g & Maximum reproduction \\
        \(U_{\textsf{hab}}\) & -- & Cod occupancy oxygen sensitivity \\
        \(a_{\textsf{hab}}\) & -- & Cod occupancy critical oxygen adjustment \\
        \(U_{\textsf{benthic}}\) & -- & Benthic resource oxygen sensitivity \\
        \(k_{\textsf{benthic}}\) & -- & Benthic resource critical oxygen level \\
        \(U_{\textsf{phys},i}\) & -- & Physiological rate oxygen sensitivity \\
        \(a_{\textsf{phys},i}\) & -- & Physiological rate critical oxygen adjustment \\
        \(U_{k_{s},i}\) & -- & Metabolic rate oxygen sensitivity \\
        \(a_{k_{s},i}\) & -- & Metabolic rate critical oxygen adjustment \\
        \(z_{h0,i}\) & y\textsuperscript{-1} & Baseline mortality due to hypoxia exposure \\
        \(b_{i}\) & -- & Mortality oxygen sensitivity \\
        \(\gamma_{i}\) & volume\(\cdot\)y\textsuperscript{-1} & Predation or clearance rate coefficient \\
        \(h_{i}\) & g\(\cdot\)y\textsuperscript{-1} & Size-scaled maximum consumption rate \\
        \(\kappa\) & g & Benthic resource carrying capacity \\
        \(z_{0}\) & g\textsuperscript{1\(-p\)}\(\cdot\)y\textsuperscript{-1} & Background mortality coefficient \\
        \(\varepsilon_{\textsf{repro}}\) & -- & Reproduction efficiency or survival of recruits \\
        \bottomrule
    \end{tabular}
    \label{tab:cal_params}
\end{table}

\paragraph{} Note that benthic and pelagic oxygen (see Section S2) and F served as data input.  For steps 1--11 we used average oxygen and F during the calibration period, and for steps 12--13 we used time series.  The code provided includes all details of implementation (\href{https://github.com/epduskey/hypoxiaSSM}{found here}).  The steps are summarized below.
    \begin{enumerate}
        \item Choose values for carrying capacity \(\kappa\) and scaling parameters of benthos and plankton.  These values should be consistent with the scale of the observed yield and SSB.  We used total biomass, but one may also use biomass per area or volume.  Note that we estimated parameters using data from ICES subdivisions 25 through 29, while some stock assessment estimates of SSB and F cover subdivisions 22 through 32, and subsets therein.  The implicit assumption is that all fish reside in the Central Baltic Sea.  The majority of eastern Baltic cod are indeed concentrated primarily in the Central Baltic Sea, while sprat, herring, and flounder extend into the Gulf of Finland \parencite[ICES subdivision 32;][]{wgbifs2020}.  For sprat and herring, the larger area can be accommodated by using a larger \(\kappa_{\textsf{pelagic}}\), reflecting greater overall food availability.  The relative distribution of flounder biomass is unknown, and therefore we lumped these species together in a single benthic habitat with cod.  Note that the values of \(\kappa\) can have a major impact on the realized diets of the fish.  Higher values generally force greater reliance on the background resource while lower values are reflective of a diet composed of explicitly modeled fish species.  Therefore, while adjusting, check diets.  Scaling parameters may arise from literature regarding the tolerance of major prey organisms in each habitat to hypoxia.  At this stage, only reasonable values are required, as they are fit to time series data in another step.
        \item Choose starting values for each species' maximum reproduction rates \(R_{\textsf{max},i}\) such that model SSB for each species is within an order of magnitude of observed SSB.  These are affected by the choice of \(\kappa\).  In practice, we have found that several combinations of \(R_{\textsf{max}}\) and \(\kappa\) and scaling values may result in matching values of SSB.  This is also affected by growth, but these parameters are calibrated in a later step.  Therefore, at this stage, we recommend focusing on the diet to narrow the range of possible values.
        \item Choose reasonable starting values for oxygen dependence of physiological rates of assimilation, feeding, and reproduction efficiency (equation \ref{eq:supp_scaling}). Generally, reasonable values of \(U_{\textsf{phys},i}\) fall below ten, and will never fall below zero.  Values of \(a_{\textsf{phys},i}>1\) shift the oxygen level at which physiological scaling is \(0.5\) to the right, while \(a_{\textsf{phys},i}<1\) shift this value to the left.  Negative values reflect a decrease in oxygen sensitivity with weight, while positive values reflect an increase.  We allowed both positive and negative values as studies differ in their claims regarding the relationship of hypoxia tolerance and weight \parencite[e.g.][]{nilsson2008does,pan2016hypoxia}.   Declines in performance begin well above \(P_{\textsf{crit},iw}\) \parencite{hrycik2017sub}.  Therefore, \(U_{\textsf{phys},i}\) will often be low, perhaps three or less, depending on the value of \(a_{\textsf{phys},i}\).  These parameters may best be characterized relative to other species.  For example, to represent our community -- and more generally a community in which one species is the superior competitor in an hypoxic environment -- we chose a lower value of \(U_{\textsf{phys},i}\) and a higher value of \(a_{\textsf{phys},i}\) for cod relative to flounder.  Check that modeled biomass is still within an order of magnitude of observed biomass.  Prioritize SSB and coexistence over specific values of \(U_{\textsf{phys},i}\) and \(a_{\textsf{phys},i}\).  Note that these values are calibrated using time series data in a later step.
        \item Choose reasonable starting values for metabolic scaling (equation \ref{eq:suppmet_scaling}).  Values of \(a_{\textsf{met},i}\) shift the point of doubling left or right, as described in the previous step.  We also allowed both positive and negative values of \(a_{\textsf{met},i}\).  In this model, metabolic costs are also a function of maximum consumption and assimilation efficiency, both of which decline with oxygen according to the physiological scaling function described in the previous step.  Therefore, overall changes in metabolic costs are a complex function of oxygen dependence.  We again chose values to characterize species relative to one another, with higher values of \(U_{\textsf{met},i}\) and lower values of \(a_{\textsf{met},i}\) for cod than for flounder.  This also reflects a greater sensitivity to hypoxic conditions in the former than in the latter.  Prioritize SSB and coexistence over specific values of \(U_{\textsf{met},i}\) and \(a_{\textsf{phys},i}\).  Note that these values are also calibrated using time series data in a later step.  Repeat steps 1 through 4 until modeled SSB is within one order of magnitude of observed SSB.
        \item Check density dependence in reproduction for each species.  One can check relative controls on each species by evaluating the ratio of density independent to density dependent recruitment.  If these values are very high, it indicates that the given population is largely affected by density dependence rather than predation or competition with other species.  If they are very low, the population may be immune to sources of mortality and competition.  In general, larger species should experience a higher degree of density dependence.  Recall that increasing density dependence within a species can result in saturating yield with increasing F \parencite{beverton2012dynamics}.  Therefore, observing this kind of relationship of yield with F in modeled species is not necessarily cause for alarm.  If time series of yield at F appear insensitive to F, then consider returning to this step and changing the degree of density dependence.
        \item Choose starting values for the background mortality coefficients and the clearance rate coefficients for each species.  Check coexistence, feeding level, growth, SSB, and yield at F.  The latter three are compared to observations.  Feeding level emerges from realized diets, and is contained in the interval \([0,1]\), with \(0\) being unfed and \(1\) being completely satiated.  Emergent values should be similar among species and above \(0.2\), which is the minimum required for maintenance.  Increasing clearance rate coefficients increases both growth rates and feeding level relative to satiation.  It may also be necessary to adjust maximum consumption in tandem with clearance rates.  Increasing maximum consumption rate (\(h\)) increases growth rates, but it decreases feeding level.  At this stage, prioritize fits to observed yield with precise values of the background mortality coefficients. 
        \item This and all subsequent steps use the \textbf{optimParallel} package.  This package applies the ``L-BFGS-B'' method, a quasi-Newton method with box constraints \parencite{byrd1995limited}, in parallel to minimize a user defined function.  If a given optimization procedure failed to converge, we increased the control argument \texttt{factr} by factors of ten until convergence was achieved.  If this failed, we increased control argument \texttt{pgtol} by factors of ten until convergence was achieved.  This only occurred in one instance, and essentially indicated that our solution space was very flat.  That is, optimization was superfluous.  Using this procedure, we first fit habitat sensitivity parameters to observed benthic occupancy of relevant species -- cod, in this case.  Recall that occupancy scales with equation \ref{eq:supp_scaling}.  For occupancy, we did not allow \(a_{i}\) to fall below zero, suggesting larger fish are more likely to escape to the pelagic habitat at a given oxygen level.
        \item Fit to observed mean yield of al species during the calibration period.  The objective function adjusts the background mortality coefficients, then calculates and returns the sum of squared errors between the natural logarithms of modeled yield at F and mean observed yield during the calibration period.
        \item Fit to observed mean SSB and growth during the calibration period for all species.  The objective function adjusts the base ten logarithm of each species' maximum reproductive rates \(R_{\textsf{max},i}\) and clearance rates \(\gamma_{i}\).  It calculates sum of squared errors between modeled biomass of mature size classes and observed SSB.  The function also predicts the weight as a percentage of \(W_{\infty}\) at fifty evenly spaced ages beginning at zero and ending at the maximum observed age for each species.  It calculates the sum of squared errors between modeled growth and von Bertalanffy growth estimates arising from the procedures in Section S2.  The function returns a weighted sum of the the error in SSB and in growth.  The weights are dependent on the scale of the errors of SSB and growth.  We found that errors in growth were typically two orders of magnitude greater than errors in SSB, and adjusted the former by a factor of 0.01.  Adjust starting values for clearance rates to roughly match von Bertalanffy growth estimates, if necessary.  Note that the order of steps 7 through 9 are in the reverse order of priority or data trustworthiness.  That is, we prioritize fits to growth and SSB above fits to yield and fits to yield above fits to occupancy.  The processes in this model are highly interdependent, and fitting one set of parameters to a subset of the data can negate the fit of another set of parameters to another subset of the data.  Thus, order reflects priority.
        \item Fit to annual growth patterns during the calibration period by adjusting the physiological and scaling parameters.  This includes scaling of assimilation, feeding, and reproductive efficiency in addition to scaling of metabolic costs.  The objective function adjusts \(U_{i}\), \(a_{i}\), \(U_{\textsf{met},i}\), and \(a_{\textsf{phys},i}\) (see equations \ref{eq:supp_scaling} and \ref{eq:suppmet_scaling}).  This procedure calculates the error in growth as described in step 9, and is performed once for each species that experiences any level of exposure to the benthic environment i.e. for cod and flounder.
        \item Fit to annual growth patterns during the calibration period by adjusting benthic resource carrying capacity scaling parameters.  Recall that this scaling function is written as:
            \begin{linenomath*}
            \begin{equation}
                \xi_{\kappa_{\textsf{benthic}}}(O_{\textsf{benthic}})=\frac{1}{1+e^{-U_{\textsf{benthic}}(O_{\textsf{benthic}}-k_{\textsf{benthic}})}}
            \end{equation}
            \end{linenomath*}
        where \(O_{\textsf{benthic}}\) references the oxygen level in the benthic environment only.  The objective function adjusts \(U_{\textsf{benthic}}\) and \(k_{\textsf{benthic}}\), and calculates the error in growth of all species. 
        \item Fit to annual patterns of yield during the calibration period by adjusting the parameters describing additional mortality due to hypoxia exposure.  Recall that this mortality is expressed by the hazard function in equation \ref{eq:hazard} (see main text), again repeated here:
            \begin{linenomath*}
            \begin{equation} \label{eq:supp_hazard}
                h(t) = b_{0,i}e^{b_{i}O_{iwt}}
            \end{equation} 
            \end{linenomath*}
        The objective function adjusts \(b_{0,i}\) and \(b_{i}\) and predicts yield at observed values of fishing mortality (F).  It then calculates the sum of squared differences between the natural logarithm of these values and observed yield.  We perform this step once for each species that experiences any level of exposure to the benthic environment i.e. for cod and flounder.
    \end{enumerate}
The procedure described above is intended for calibration of the full model, which includes oxygen dependence of all four processes: benthic resource carrying capacity (B), avoidance via changes in occupancy (O), mortality due to exposure to hypoxic waters (M), and physiological scaling (P; the so-called BOMP model).  We repeated calibration for all possible combinations of oxygen dependence.  We also calibrated a model with no oxygen dependence.  This resulted in 16 total model calibrations.  For those processes which were not included in a given model, we turned off the associated scaling function by setting the scaling coefficient to one for all species at all sizes, or by setting the result to zero for all species at all sizes in the case of mortality.  We would then skip the associated step.  However, all steps were still performed in the order listed above, and with the same checks, if applicable.

\subsubsection*{Error calculation}

We used the calibrated models to project forward in time from 2001 through 2019 and calculated errors in modeled quantities during both the calibration, or in-sample, period (1991-2000) and the projection, or out-of-sample, period (2001-2019).  Only the out-of-sample or projection errors were used for model selection, as this is generally more robust \parencite{cooke2014out}.  We calculated a weighted sum of squared errors between the natural logarithms of modeled biomass of mature size classes and observed SSB for all species:  
    \begin{linenomath*}
    \begin{equation} \label{eq:ssb_error}
        \varepsilon_{\textsf{SSB}} = \sum_{i=1}^{4} \sum_{j=1}^{19} w_{\textsf{SSB},i}(\ln{\textsf{SSB}_{\textsf{mod},ij}}-\ln{\textsf{SSB}_{\textsf{obs},ij}})^{2}
    \end{equation}
    \end{linenomath*}
where \(w_{\textsf{SSB},i}=\{0.3,0.1,0.3,0.3\}\) for cod, flounder, sprat, and herring, in that order.  The weights reflected our confidence in each data source.  The relatively low confidence in flounder data reflects the ad-hoc method by which we estimated values for flounder SSB.  We used the same method to calculate errors between the natural logarithms of modeled yield at F and observed yield:
    \begin{linenomath*}
    \begin{equation} \label{eq:yield_error}
        \varepsilon_{\textsf{Yield}} = \sum_{i=1}^{4} \sum_{j=1}^{19} w_{\textsf{Yield},i}(\ln{\textsf{Yield}_{\textsf{mod},ij}}-\ln{\textsf{yield}_{\textsf{obs},ij}})^{2}
    \end{equation}
    \end{linenomath*}
where \(w_{\textsf{Yield},i}=\{0.25,0.25,0.25,0.25\}\) for cod, flounder, sprat, and herring, in that order.  Lastly, we calculated annual error in size at age by predicting the weight as a percentage of \(W_{\infty}\) at fifty evenly spaced ages beginning at zero and ending at the maximum observed age for each species and comparing to annual von Bertalanffy estimates:
    \begin{linenomath*}
    \begin{equation} \label{eq:growth_error}
        \varepsilon_{\textsf{Growth}} = \sum_{i=1}^{4} \sum_{j=1}^{19} \sum_{k=1}^{50} w_{\textsf{Growth},i}(\textsf{Size}_{\textsf{mod},ijk}-\textsf{Size}_{\textsf{obs},ijk})^{2}
    \end{equation}
    \end{linenomath*}
where \(w_{\textsf{Yield},i}=\{0.25,0.25,0.25,0.25\}\) for cod, flounder, sprat, and herring, in that order.
\paragraph{} We repeated these calculations for each of the 16 calibrated models to arrive at a set of errors: \(\symbf{\varepsilon}_{\textsf{SSB}}=\{\varepsilon_{\textsf{SSB},1},...,\varepsilon_{\textsf{SSB},16}\}\).  Similarly, we have \(\symbf{\varepsilon}_{\textsf{Yield}}\) and \(\symbf{\varepsilon}_{\textsf{Growth}}\), containing errors in yield and growth, respectively, for all 16 models.  We normalized each set of errors to span across the interval \([0,1]\) with the standard normalization function:
    \begin{linenomath*}
    \begin{equation} \label{eq:norm}
        \symbf{\varepsilon}_{\textsf{norm}} = \frac{\symbf{\varepsilon}-\min(\symbf{\varepsilon})}{\max(\symbf{\varepsilon})-\min(\symbf{\varepsilon})}
    \end{equation}
    \end{linenomath*}
This ensured that each set of errors were compared on the same scale, and allowed us to arrive at a final ranking for each model by computing a weighted sum of normalized errors:
    \begin{linenomath*}
    \begin{equation} \label{eq:final_error}
        \varepsilon_{\textsf{Final}} = 0.3\,\varepsilon_{\textsf{SSB}} + 0.1\,\varepsilon_{\textsf{Yield}} + 0.6\,\varepsilon_{\textsf{Growth}}
    \end{equation}
    \end{linenomath*}
These weights reflect our priority in fitting to each set of observations, which in turn reflects our degree of confidence in the data.  The model with the lowest value of \(\varepsilon_{\textsf{Final}}\) was treated as the best model overall.

\subsubsection*{Results of model calibration}

Estimated scaling parameters reflected both the nature of multispecies size spectrum models and differences in sensitivity of cod and flounder (Table \ref{tab:all_scaling} and Figure \ref{fig:all_scaling}).  For example, estimated benthic resource scaling parameters (\(U_{\textsf{res}}\) and \(k_{\textsf{res}}\)) were nearly identical across models, and did not stray far from initial values of 2 and 0.8, respectively.  However, these scaling values also affect initialization of the \textbf{mizer} model itself.  That is, in choosing values for resource scaling, initial values for abundance across size classes are adjusted.  This is true for all scaling parameters, and thus values may not reflect \textit{actual} oxygen sensitivity of each process, but rather an amalgam of internal \textbf{mizer} setup and oxygen sensitivity.

\begin{table}[h]
    \scriptsize
    \centering
    \caption{All estimated scaling parameters for all 16 models for cod and flounder.  The subscript res refers to benthic resource scaling and is identical within a given model.  The subscript hab refers to benthic occupancy scaling; crit refers to consumption, assimilation efficiency, and reproduction efficiency scaling; and met refers to metabolic costs scaling.  The mortality parameters \(b_{0}\) and \(b_{i}\) are drawn from the hazard function given by equation {\ref{eq:hazard}}.}
    \begin{tabular}{llrrrrrrrrrr}
        \toprule
        Species & Model & \(U_{\textsf{res}}\) & \(k_{\textsf{res}}\) & \(U_{\textsf{hab}}\) & \(a_{\textsf{hab}}\) & \(U_{\textsf{crit}}\) & \(a_{\textsf{crit}}\) & \(U_{\textsf{met}}\) & \(a_{\textsf{met}}\) & \(b_{0}\) & \(b_{i}\) \\
        \midrule
        Cod & BOMP & 2.00 & 0.79 & 0.82 & 0.00 & 3.82 & 0.74 & 1.84 & 0.90 & 0.00 & 1.83 \\ 
        & BOM & 2.04 & 0.79 & 0.82 & 0.00 & -- & -- & -- & -- & 0.00 & 1.98 \\ 
        & BOP & 2.00 & 0.80 & 0.82 & 0.00 & 3.58 & 0.82 & 1.64 & 0.61 & -- & -- \\ 
        & BMP & 2.00 & 0.85 & -- & -- & 2.62 & 0.77 & 1.09 & 0.36 & 0.00 & 1.92 \\ 
        & OMP & -- & -- & 0.82 & 0.00 & 3.34 & 0.82 & 1.49 & 0.33 & 0.00 & 1.79 \\ 
        & BO & 1.97 & 0.80 & 0.82 & 0.00 & -- & -- & -- & -- & -- & -- \\ 
        & BM & 2.03 & 0.80 & -- & -- & -- & -- & -- & -- & 0.00 & 1.97 \\ 
        & BP & 2.00 & 0.84 & -- & -- & 2.44 & 0.74 & 1.34 & 0.42 & -- & -- \\ 
        & OM & -- & -- & 0.82 & 0.00 & -- & -- & -- & -- & 0.00 & 1.98 \\ 
        & OP & -- & -- & 0.82 & 0.00 & 1.00 & 0.21 & 1.00 & 0.20 & -- & -- \\ 
        & MP & -- & -- & -- & -- & 2.29 & 0.73 & 1.40 & 0.32 & 0.00 & 1.87 \\ 
        & B & 2.05 & 0.77 & -- & -- & -- & -- & -- & -- & -- & -- \\ 
        & O & -- & -- & 0.82 & 0.00 & -- & -- & -- & -- & -- & -- \\ 
        & M & -- & -- & -- & -- & -- & -- & -- & -- & 0.00 & 1.98 \\ 
        & P & -- & -- & -- & -- & 2.26 & 0.68 & 1.60 & 0.71 & -- & -- \\ 
        & None & -- & -- & -- & -- & -- & -- & -- & -- & -- & -- \\ 
        \midrule
        Flounder & BOMP & 2.00 & 0.79 & -- & -- & 1.03 & -0.77 & 1.25 & -0.65 & 0.00 & 2.14 \\ 
        & BOM & 2.04 & 0.79 & -- & -- & -- & -- & -- & -- & 0.00 & 2.11 \\ 
        & BOP & 2.00 & 0.80 & -- & -- & 0.82 & -1.23 & 1.98 & -0.95 & -- & -- \\ 
        & BMP & 2.00 & 0.85 & -- & -- & 0.60 & -1.92 & 1.40 & -2.04 & 0.00 & 2.15 \\ 
        & OMP & -- & -- & -- & -- & 0.33 & -4.93 & 0.48 & -3.23 & 0.00 & 2.15 \\ 
        & BO & 1.97 & 0.80 & -- & -- & -- & -- & -- & -- & -- & -- \\ 
        & BM & 2.03 & 0.80 & -- & -- & -- & -- & -- & -- & 0.00 & 2.12 \\ 
        & BP & 2.00 & 0.84 & -- & -- & 0.56 & -2.29 & 0.98 & -1.36 & -- & -- \\ 
        & OM & -- & -- & -- & -- & -- & -- & -- & -- & 0.00 & 2.12 \\ 
        & OP & -- & -- & -- & -- & 0.32 & -4.50 & 1.62 & -2.50 & -- & -- \\ 
        & MP & -- & -- & -- & -- & 0.44 & -3.42 & 0.31 & -4.91 & 0.00 & 2.14 \\ 
        & B & 2.05 & 0.77 & -- & -- & -- & -- & -- & -- & -- & -- \\ 
        & O & -- & -- & -- & -- & -- & -- & -- & -- & -- & -- \\ 
        & M & -- & -- & -- & -- & -- & -- & -- & -- & 0.00 & 2.11 \\ 
        & P & -- & -- & -- & -- & 0.39 & -3.85 & 1.37 & -2.56 & -- & -- \\ 
        & None & -- & -- & -- & -- & -- & -- & -- & -- & -- & -- \\ 
    \bottomrule
    \label{tab:all_scaling}
\end{tabular}
\end{table}

\begin{figure}[h]
    \centering
    \includegraphics[scale=0.5]{figures/all_scaling.jpg}
    \caption{Scaling of benthic resource carrying capacity (Benthos), benthic occupancy of cod (Occupancy), and physiological rates of cod and flounder for all 16 models.  For the latter two plots, lines above 1 are up-scaling of metabolic costs while lines below 1 are down-scaling of assimilation, feeding, and reproduction efficiency.  Note the vast difference in scale between cod and flounder.  Opacity of line coloration declines as weight increases..}
    \label{fig:all_scaling}
\end{figure}

\paragraph{} Cod benthic occupancy parameters were identical across models, and suggested occupancy was identical across all sizes (Table \ref{tab:all_scaling}, \(a_{\textsf{hab}}\) and Figure).  Given the data were interpreted as single probabilities of benthic occupancy for each year, it is unsurprising that there was no differentiation among sizes, and thus no differentiation among models.

\begin{figure}[h]
    \centering
    \includegraphics[scale=0.6]{figures/occupancy.jpg}
    \caption{Results of fitting occupancy parameters for each model to observations from \textcite{casini2019spatio}.  The shaded region indicated the calibration period.  Grayscale lines are models with various combinations of B/O/M/P, with lighter shades indicating the model has less variables.  Lines with zero slope indicate that O was not included in the model.}
    \label{fig:occupancy}
\end{figure}

\paragraph{} Oxygen dependence of natural mortality was always estimated to be zero, no matter the model (Table {\ref{tab:all_scaling}}).  However, it is important to note that the model generally underestimated yield of cod and flounder during the calibration period (see Figure {\ref{fig:sim_yield}} in the main text), and the inclusion of additional mortality during model setup caused some initial values to be up-scaled, thereby increasing yield overall, and decreasing error.  Given natural mortality parameters were fit to yield, models including these components generally performed better than those which did not, but not for any interesting mechanistic reason.
\paragraph{} Physiological scaling parameters varied by species and by model.  Generally speaking, cod exhibited greater sensitivity to declining oxygen than flounder, in accordance with expectations.  Smaller cod were also consistently less sensitive to low oxygen than larger cod.  Conversely, smaller flounder were more sensitive to low oxygen than larger flounder (Figure \ref{fig:all_scaling}).  Furthermore, most models suggested that increases in metabolic costs began at similar oxygen levels to those at which other physiological processes began to decline.  This was true for both cod and flounder, evident in relatively similar values of \(a_{\textsf{crit}}\) and \(a_{\textsf{met}}\).  That said, values of \(a_{\textsf{met}}\) were generally smaller than values of \(a_{\textsf{crit}}\) for cod, and less negative for flounder, indicating a fairly consistent characterization.  This likely compensated for the exponential nature of the curve, which results in steep increases in metabolic costs at zero oxygen, for example, with modest increases in \(a_{\textsf{met}}\).
\paragraph{}  Recall that we only used errors during the projection period for model selection.  These errors in fits to SSB, yield, and growth during both the calibration and the projection period suggest that there are complex interactions between model components.  First, note that errors in cod and flounder model output experienced the largest change in magnitude across models in the projection period (Figure \ref{fig:sim_error}).  This was unsurprising, given that they experienced direct consequences from exposure to hypoxic waters.  Of the two, errors in cod output spanned a wider range of values (Figure \ref{fig:sim_error}).

\begin{figure}[h]
    \centering
    \includegraphics[scale=0.55]{figures/simulation_error.jpg}
    \caption{Weighted error during the projection period (2001--2019) in \(\ln(\textsf{SSB})\), \(\ln(\textsf{yield})\), and size-at-age or growth as a percentage of \(W_{\infty}\) (see Table \ref{tab:allparams}).  Black, inverted triangles are sum totals, other filled symbols correspond to each species.  Models are ordered from best to worst for each component.}
    \label{fig:sim_error}
\end{figure}

\paragraph{} Second, errors in SSB during the projection period were clearly exacerbated by models which included physiological scaling without occupancy scaling (Figure \ref{fig:sim_error}).  That is, physiological rates are oxygen dependent, it is essential that cod be allowed to escape the environment.  This was also true of errors in yield.  For growth, those models which include physiological scaling tend to perform better when occupancy scaling is also present, though it is not entirely consistent.
\paragraph{} We relied upon summarized rankings during the projection period (2001--2019) to provide the best model as a weighted compromise among model components.  Final weighted rankings (equation \ref{eq:final_error}) largely follow patterns evident in growth, given the higher weight based on observations of size-at-age (Figure \ref{fig:final_rank_sim}).  In general, models including benthic resource scaling performed significantly better than those without.  Though, again, models which include physiological scaling must also include occupancy scaling lest they be overwhelmed by error in SSB and yield (Figure \ref{fig:sens_rank_sim}).  Model choice through calibration and projection may best be performed as a check on those components which must appear together, or which introduce egregious error.

\begin{figure}[ht!]
    \centering
    \includegraphics[scale=0.55]{figures/final_error_sim.jpg}
    \caption{Final weighted error rankings during the projection period (2001--2019).  Black filled circles are final rankings, while other filled symbols give weighted rankings of other model components.  Models are ordered from best to worst.}
    \label{fig:final_rank_sim}
\end{figure}

\paragraph{} We did not rely upon errors during the calibration period (1991--2000) for model selection.  Still, we note that good performance during the calibration period did not translate to good performance during the projection period (Figure \ref{fig:final_rank_cal}).  This could possibly be due to a change in the mechanisms driving responses to hypoxia during over time.  Testing this would require extensive re-calibration and hindcasting, which we leave for future studies.

\begin{figure}[ht!]
    \centering
    \includegraphics[scale=0.55]{figures/final_error_cal.jpg}
    \caption{Final weighted error rankings during the calibration period (1991--2000).  Symbols and order are given as in Figure \ref{fig:final_rank_sim}.}
    \label{fig:final_rank_cal}
\end{figure}
\clearpage

\subsection*{Appendix S6 Sensitivity analysis}
\addcontentsline{toc}{section}{Appendix S6 Sensitivity analysis}

We repeated the calibration process detailed in Appendix S5 for all 16 models three times to test the sensitivity of our results to parameter starting values.  Each time, we drew new starting values for all oxygen dependence parameters from a uniform distribution with a minimum and maximum 20\% below and 20\% above the original starting value, respectively.  We set three separate random seeds for these starting values so that results are repeatable, and so that the same set of starting values could be used to calibrate all 16 models within each repetition.  While three repetitions gives a very small sample size, the calibration procedure is both extensive and labor intensive, which limits the number of times it can be repeated within a reasonable amount of time.
\paragraph{} Results of this procedure indicate that while model rankings are sensitive to starting values, benthic food availability and habitat use consistently play vital roles in impacts of hypoxia on fish communities.  Models with the lowest error during the projection period (2001--2019) always include oxygen dependence of benthic food resources (Figure \ref{fig:sens_rank_sim}).  The worst models always include oxygen dependence of physiological rates without an oxygen replete refuge.  These mechanisms, above all others, determine the ability of the models to explain observed trends in spawning stock biomass (SSB), commercial yield, and somatic growth.

\begin{figure}[ht!]
    \centering
    \includegraphics[scale=0.6]{figures/sens_error_sim.jpg}
    \caption{Final weighted error rankings during the projection period (2001--2019) for all three model calibration repetitions completed for the sensitivity analysis.  Symbols and model order are given as in Figure \ref{fig:final_rank_sim}.}
    \label{fig:sens_rank_sim}
\end{figure}

\paragraph{} Error during the calibration period follows a similar pattern, with the same sets of models clustering among those which are best or worst at explaining observed patterns (Figure \ref{fig:sens_rank_cal}).  Thus, our general conclusions given in the main text are still supported.  We only caution against allowing single runs to serve as definitive evidence in favor of some mechanisms and to the exclusion of others.  Rather, it may be best to focus on a collection of models which consistently appear among the highest ranked models during sensitivity analyses, or rather to focus on a collection of models with similarly high rankings.

\begin{figure}[ht!]
    \centering
    \includegraphics[scale=0.6]{figures/sens_error_cal.jpg}
    \caption{Final weighted error rankings during the calibration period (1991--2000) for all three model calibration repetitions completed for the sensitivity analysis.  Symbols and model order are given as in Figure \ref{fig:final_rank_sim}.}
    \label{fig:sens_rank_cal}
\end{figure}
\clearpage
\clearpage

\subsection*{Appendix S7 Tables and Figures}
\addcontentsline{toc}{section}{Appendix S7 Tables and Figures}

\begin{figure}[ht!]
    \centering
    \includegraphics[scale=0.6]{figures/cod_growth_eight.jpg}
    \caption{Projections of body growth for cod during the projection period (2001--2019) arising from all 16 models.  Data are included as small, black triangles.  Dotted, light blue lines are von Bertalanffy growth estimates from each year.}
    \label{fig:cod_growth_eight_supp}
\end{figure}
\clearpage

\begin{figure}[ht]
    \centering
    \includegraphics[scale=0.6]{figures/sim_biomass.jpg}
    \caption{Projections of biomass of mature size classes (SSB) throughout the calibration period (1991--2000; shaded region) and the projection period (2001--2019) using all 16 versions of the oxygen-dependent food web model.  Blue triangles are observations (as estimated from stock assessments; see Methods of main text).}
    \label{fig:sim_ssb_supp}
\end{figure}
\clearpage

\begin{figure}[ht]
    \centering
    \includegraphics[scale=0.6]{figures/sim_yield.jpg}
    \caption{Projections of yield at observed fishing mortality, throughout the calibration period (1991--2000; shaded region) and the projection period (2001--2019), using all 16 versions of the oxygen-dependent food web model.  Symbols as in Figure \ref{fig:sim_ssb_supp}.}
    \label{fig:sim_yield_supp}
\end{figure}
\clearpage

\begin{figure}[ht]
    \centering
    \begin{adjustbox}{addcode={\begin{minipage}{\width}}{\caption{Projections of growth for each species throughout the projection period (2001--2019).  Dotted blue lines are predictions from annual estimates of von Bertalanffy growth parameters.\label{fig:sim_growth}}\end{minipage}},rotate=90,center}
    \includegraphics[scale=0.5]{figures/sim_growth.jpg}
    \end{adjustbox}
\end{figure}
\clearpage

\begin{figure}[ht]
    \centering
    \begin{adjustbox}{addcode={\begin{minipage}{\width}}{\caption{Projections of somatic growth (size-at-age) during the first eight years of life for each species throughout the calibration period (1991--2000) and the projection period (2001--2019), using the oxygen-dependent food web model.  Lines and symbols given as in Figure \ref{fig:sim_growth}.\label{fig:sim_growth_eight}}\end{minipage}},rotate=90,center}
    \includegraphics[scale=0.5]{figures/sim_growth_eight.jpg}
    \end{adjustbox}
\end{figure}
\clearpage

\begin{table}[ht]
    \scriptsize
    \centering
    \caption{Modeled SSB (kt), yield (kt) at mean fishing mortality observed during the calibration period (1991--2000), and size (g) of individuals at maximum observed age for all species under three oxygen scenarios applied in all 16 models.}
    \begin{tabular}{llrrrrrrrrrrrr}
        \toprule
        & & \multicolumn{3}{c}{Cod} & \multicolumn{3}{c}{Flounder} & \multicolumn{3}{c}{Sprat} & \multicolumn{3}{c}{Herring} \\
        \cmidrule(r){3-5} \cmidrule(lr){6-8} \cmidrule(lr){9-11} \cmidrule(l){12-14}
        Model & O\textsubscript{2} & SSB & Yield & Size & SSB & Yield & Size & SSB & Yield & Size & SSB & Yield & Size \\ 
        \midrule
        BOMP & 3 & 87.3 & 62.1 & 14266 & 9.2 & 7.4 & 1125 & 1688.9 & 519.5 & 20 & 570.1 & 243.6 & 172 \\ 
        & 2 & 82.2 & 51.2 & 9746 & 8.2 & 6.5 & 1087 & 1385.8 & 426.6 & 20 & 526.8 & 225.4 & 174 \\ 
        & 1 & 64.3 & 31.8 & 4286 & 2.7 & 2.0 & 795 & 811.1 & 250.2 & 21 & 438.5 & 188.2 & 176 \\ 
        BOM & 3 & 75.8 & 51.7 & 12269 & 10.0 & 8.0 & 1103 & 1726.9 & 531.1 & 20 & 572.2 & 244.3 & 172 \\ 
        & 2 & 82.3 & 51.1 & 12169 & 8.3 & 6.6 & 1086 & 1429.7 & 440.1 & 20 & 533.4 & 228.2 & 173 \\ 
        & 1 & 72.6 & 42.9 & 8777 & 5.7 & 4.3 & 923 & 860.1 & 265.3 & 21 & 452.2 & 194.0 & 176 \\ 
        BOP & 3 & 86.0 & 62.0 & 14562 & 8.9 & 7.1 & 1123 & 1702.7 & 523.7 & 20 & 571.2 & 244.0 & 172 \\ 
        & 2 & 82.3 & 51.1 & 9439 & 8.3 & 6.6 & 1090 & 1429.7 & 440.1 & 20 & 533.4 & 228.2 & 174 \\ 
        & 1 & 66.5 & 33.5 & 4495 & 3.5 & 2.6 & 860 & 886.3 & 273.3 & 21 & 452.0 & 193.9 & 176 \\ 
        BMP & 3 & 78.1 & 54.4 & 13139 & 9.1 & 7.3 & 1123 & 1964.3 & 603.7 & 20 & 608.4 & 259.4 & 170 \\ 
        & 2 & 0.0 & 0.0 & 143 & 14.7 & 11.8 & 1102 & 1964.3 & 603.7 & 20 & 608.4 & 259.4 & 170 \\ 
        & 1 & 0.0 & 0.0 & 0 & 8.3 & 6.2 & 895 & 1964.3 & 603.7 & 20 & 608.4 & 259.4 & 170 \\ 
        OMP & 3 & 86.4 & 61.9 & 14296 & 7.9 & 6.3 & 1107 & 1673.8 & 514.8 & 20 & 566.7 & 242.1 & 173 \\ 
        & 2 & 86.4 & 54.1 & 9589 & 8.2 & 6.6 & 1098 & 1415.3 & 435.7 & 20 & 531.0 & 227.2 & 174 \\ 
        & 1 & 88.4 & 49.9 & 6719 & 7.9 & 6.3 & 1084 & 920.5 & 283.8 & 21 & 458.1 & 196.5 & 176 \\ 
        BO & 3 & 73.2 & 50.0 & 12324 & 9.9 & 7.9 & 1106 & 1710.2 & 526.1 & 20 & 573.1 & 244.8 & 173 \\ 
        & 2 & 77.5 & 52.1 & 12204 & 9.2 & 7.3 & 1087 & 1432.9 & 441.1 & 21 & 532.7 & 227.9 & 175 \\ 
        & 1 & 70.1 & 41.7 & 9020 & 5.5 & 4.2 & 924 & 923.6 & 284.8 & 21 & 457.6 & 196.3 & 177 \\ 
        BM & 3 & 69.2 & 47.2 & 11719 & 10.0 & 8.0 & 1103 & 1863.1 & 572.8 & 20 & 592.7 & 252.8 & 172 \\ 
        & 2 & 65.3 & 43.8 & 11098 & 9.4 & 7.5 & 1085 & 1863.1 & 572.8 & 20 & 592.7 & 252.8 & 172 \\ 
        & 1 & 43.8 & 25.9 & 6696 & 6.2 & 4.7 & 924 & 1863.1 & 572.8 & 20 & 592.7 & 252.8 & 172 \\ 
        BP & 3 & 76.5 & 53.1 & 12996 & 9.2 & 7.4 & 1122 & 1934.9 & 594.7 & 20 & 605.7 & 258.4 & 171 \\ 
        & 2 & 0.0 & 0.0 & 366 & 14.6 & 11.7 & 1103 & 1934.9 & 594.7 & 20 & 605.7 & 258.4 & 171 \\ 
        & 1 & 0.0 & 0.0 & 0 & 8.2 & 6.2 & 902 & 1934.9 & 594.7 & 20 & 605.7 & 258.4 & 171 \\ 
        OM & 3 & 73.3 & 50.1 & 12330 & 9.3 & 7.4 & 1091 & 1711.5 & 526.5 & 20 & 570.2 & 243.4 & 171 \\ 
        & 2 & 80.8 & 55.1 & 12648 & 9.3 & 7.4 & 1091 & 1417.6 & 436.5 & 21 & 529.8 & 226.5 & 172 \\ 
        & 1 & 88.2 & 58.7 & 12259 & 9.4 & 7.5 & 1091 & 868.4 & 267.8 & 21 & 452.0 & 193.9 & 175 \\ 
        OP & 3 & 75.3 & 52.0 & 13345 & 8.7 & 7.0 & 1109 & 1741.4 & 535.6 & 20 & 576.0 & 246.0 & 172 \\ 
        & 2 & 74.1 & 47.9 & 11420 & 8.8 & 7.0 & 1100 & 1487.2 & 457.7 & 20 & 540.7 & 231.2 & 173 \\ 
        & 1 & 78.3 & 48.7 & 10469 & 8.2 & 6.5 & 1084 & 968.7 & 298.7 & 21 & 465.4 & 199.6 & 176 \\ 
        MP & 3 & 76.7 & 52.7 & 12698 & 8.5 & 6.8 & 1108 & 1919.6 & 590.1 & 20 & 600.0 & 255.9 & 171 \\ 
        & 2 & 0.0 & 0.0 & 624 & 14.6 & 11.7 & 1108 & 1919.6 & 590.1 & 20 & 600.0 & 255.9 & 171 \\ 
        & 1 & 0.0 & 0.0 & 0 & 13.4 & 10.7 & 1089 & 1919.6 & 590.1 & 20 & 600.0 & 255.9 & 171 \\ 
        B & 3 & 67.0 & 45.6 & 11678 & 10.2 & 8.2 & 1103 & 1894.0 & 582.2 & 20 & 592.6 & 252.7 & 171 \\ 
        & 2 & 63.5 & 42.6 & 11102 & 9.7 & 7.7 & 1087 & 1894.0 & 582.2 & 20 & 592.6 & 252.7 & 171 \\ 
        & 1 & 43.5 & 25.8 & 6889 & 6.5 & 4.9 & 935 & 1894.0 & 582.2 & 20 & 592.6 & 252.7 & 171 \\ 
        O & 3 & 71.4 & 48.2 & 11929 & 9.5 & 7.6 & 1089 & 1725.6 & 530.8 & 20 & 569.9 & 243.3 & 172 \\ 
        & 2 & 78.7 & 53.1 & 12266 & 9.5 & 7.6 & 1089 & 1435.2 & 441.8 & 20 & 531.8 & 227.4 & 174 \\ 
        & 1 & 86.0 & 56.5 & 11878 & 9.6 & 7.7 & 1089 & 891.8 & 275.0 & 20 & 458.2 & 196.5 & 176 \\ 
        M & 3 & 67.2 & 45.3 & 11311 & 9.6 & 7.7 & 1091 & 1860.2 & 571.9 & 20 & 588.6 & 251.0 & 171 \\ 
        & 2 & 67.2 & 45.3 & 11311 & 9.6 & 7.7 & 1091 & 1860.2 & 571.9 & 20 & 588.6 & 251.0 & 171 \\ 
        & 1 & 67.2 & 45.3 & 11311 & 9.6 & 7.7 & 1091 & 1860.2 & 571.9 & 20 & 588.6 & 251.0 & 171 \\ 
        P & 3 & 73.4 & 50.5 & 12719 & 8.7 & 7.0 & 1107 & 1900.5 & 584.3 & 20 & 600.7 & 256.2 & 171 \\ 
        & 2 & 0.0 & 0.0 & 280 & 14.9 & 12.0 & 1108 & 1900.5 & 584.3 & 20 & 600.7 & 256.2 & 171 \\ 
        & 1 & 0.0 & 0.0 & 0 & 13.9 & 11.1 & 1092 & 1900.5 & 584.3 & 20 & 600.7 & 256.2 & 171 \\ 
        None & 3 & 65.1 & 44.4 & 11767 & 9.3 & 7.4 & 1092 & 1875.8 & 576.7 & 20 & 590.4 & 251.7 & 170 \\ 
        \bottomrule
    \end{tabular}
    \label{tab:supp_scenarios}
\end{table}

\clearpage

\raggedright
\printbibliography
\end{refsection}

\end{document}
