## Supplementary material for "Declining food availability and habitat shifts drive community responses to marine hypoxia": LaTeX files: ssm.bib

@incollection{diaz2009hypoxic,
  title={The {H}ypoxic {E}nvironment},
  author={Diaz, Robert J and Breitburg, Denise L},
  booktitle={Fish Physiology},
  volume={27},
  pages={1--23},
  year={2009},
  publisher={Elsevier},
  doi={10.1016/S1546-5098(08)00001-0}
}

@article{pichavant2001comparative,
  title={Comparative effects of long-term hypoxia on growth, feeding and oxygen consumption in juvenile turbot and {E}uropean sea bass},
  author={Pichavant, Karine and Person-Le-Ruyet, J and Bayon, N Le and Severe, Armelle and Roux, A Le and Boeuf, Gilles},
  journal={Journal of Fish Biology},
  volume={59},
  number={4},
  pages={875--883},
  year={2001},
  publisher={Wiley Online Library},
  doi={10.1111/j.1095-8649.2001.tb00158.x}
}

@incollection{chapman2009behavioral,
  title={Behavioral {R}esponses and {E}cological {C}onsequences},
  author={Chapman, Lauren J and Mckenzie, David J},
  booktitle={Fish Physiology},
  volume={27},
  pages={25--77},
  year={2009},
  publisher={Elsevier},
  doi={10.1016/S1546-5098(08)00002-2}
}

@incollection{wu2009effects,
  title={Effects of {H}ypoxia on {F}ish {R}eproduction and {D}evelopment},
  author={Wu, Rudolf SS},
  booktitle={Fish Physiology},
  volume={27},
  pages={79--141},
  year={2009},
  publisher={Elsevier},
  doi={10.1016/S1546-5098(08)00003-4}
}

@incollection{portner2009oxygen,
  title={Oxygen and {C}apacity {L}imited {T}hermal {T}olerance},
  author={P{\"o}rtner, Hans O and Lannig, Gisela},
  booktitle={Fish Physiology},
  volume={27},
  pages={143--191},
  year={2009},
  publisher={Elsevier},
  doi={10.1016/S1546-5098(08)00004-6}
  
}

@article{rogers2016new,
  title={A new analysis of hypoxia tolerance in fishes using a database of critical oxygen level ({P}\textsubscript{crit})},
  author={Rogers, Nicholas J and Urbina, Mauricio A and Reardon, Erin E and McKenzie, David J and Wilson, Rod W},
  journal={Conservation Physiology},
  volume={4},
  number={1},
  year={2016},
  publisher={Oxford University Press},
  note = {cow012},
  doi={10.1093/conphys/cow012}
}

@incollection{wang2009effects,
  title={The {E}ffects of {H}ypoxia on {G}rowth and {D}igestion},
  author={Wang, Tobias and Lefevre, Sjannie and Huong, Do Thi Thanh and van Cong, Nguyen and Bayley, Mark},
  booktitle={Fish Physiology},
  volume={27},
  pages={361--396},
  year={2009},
  publisher={Elsevier},
  doi={10.1016/S1546-5098(08)00008-3}
}

@article{von2002effects,
  title={The effects of body mass and feeding on metabolic rate in small juvenile {A}tlantic cod},
  author={von Herbing, I Hunt and White, L},
  journal={Journal of Fish Biology},
  volume={61},
  number={4},
  pages={945--958},
  year={2002},
  publisher={Wiley Online Library},
  doi={10.1111/j.1095-8649.2002.tb01854.x}
}

@incollection{farrell2009defining,
  title={{D}efining {H}ypoxia: {A}n {I}ntegrative {S}ynthesis of the {R}esponses of {F}ish to {H}ypoxia},
  author={Farrell, Anthony P and Richards, Jeffrey G},
  booktitle={Fish Physiology},
  volume={27},
  pages={487--503},
  year={2009},
  publisher={Elsevier},
  doi={10.1016/S1546-5098(08)00011-3}
}

@article{mizer,
    author = {Finlay Scott and Julia L. Blanchard and Ken H. Andersen},
    title = {mizer: {a}n {R} package for multispecies, trait-based and community size spectrum ecological modelling},
    journal = {Methods in Ecology and Evolution},
    year = {2014},
    volume = {5},
    number={10},
    pages = {1121-1125},
    doi = {10.1111/2041-210X.12256},
}

@article{silvert1978energy,
  title={Energy flux in the pelagic ecosystem: a time-dependent equation},
  author={Silvert, William and Platt, Trevor},
  journal={Limnology and Oceanography},
  volume={23},
  number={4},
  pages={813--816},
  year={1978},
  publisher={Wiley Online Library},
  doi={10.4319/lo.1978.23.4.0813}
}

@article{andersen2016theoretical,
  title={The theoretical foundations for size spectrum models of fish communities},
  author={Andersen, Ken H and Jacobsen, Nis S and Farnsworth, Keith D},
  journal={Canadian Journal of Fisheries and Aquatic Sciences},
  volume={73},
  number={4},
  pages={575--588},
  year={2016},
  publisher={NRC Research Press},
  doi={10.1139/cjfas-2015-0230}
}

@article{casini2021changes,
  title={Changes in population depth distribution and oxygen stratification are involved in the current low condition of the eastern {B}altic {S}ea cod (\textit{{G}adus morhua})},
  author={Casini, Michele and Hansson, Martin and Orio, Alessandro and Limburg, Karin},
  journal={Biogeosciences},
  volume={18},
  number={4},
  pages={1321--1331},
  year={2021},
  publisher={Copernicus GmbH},
  doi={10.5194/bg-18-1321-2021}
}

@article{blanchard2012potential,
  title={Potential consequences of climate change for primary production and fish production in large marine ecosystems},
  author={Blanchard, Julia L and Jennings, Simon and Holmes, Robert and Harle, James and Merino, Gorka and Allen, J Icarus and Holt, Jason and Dulvy, Nicholas K and Barange, Manuel},
  journal={Philosophical Transactions of the Royal Society B: Biological Sciences},
  volume={367},
  pages={2979--2989},
  year={2012},
  publisher={The Royal Society},
  doi={10.1098/rstb.2012.0231}
}

@article{blanchard2014evaluating,
  title={Evaluating targets and trade-offs among fisheries and conservation objectives using a multispecies size spectrum model},
  author={Blanchard, Julia L and Andersen, Ken H and Scott, Finlay and Hintzen, Niels T and Piet, Gerjan and Jennings, Simon},
  journal={Journal of Applied Ecology},
  volume={51},
  number={3},
  pages={612--622},
  year={2014},
  publisher={Wiley Online Library},
  doi={10.1111/1365-2664.12238}
}

@article{blanchard2017bacteria,
  title={From {B}acteria to {W}hales: using {F}unctional {S}ize {S}pectra to {M}odel {M}arine {E}cosystems},
  author={Blanchard, Julia L and Heneghan, Ryan F and Everett, Jason D and Trebilco, Rowan and Richardson, Anthony J},
  journal={Trends in Ecology \& Evolution},
  volume={32},
  number={3},
  pages={174--186},
  year={2017},
  publisher={Elsevier},
  doi={10.1016/j.tree.2016.12.003}
}

@article{orio2020long,
  title={Long-term changes in spatial overlap between interacting cod and flounder in the {B}altic {S}ea},
  author={Orio, Alessandro and Bergstr{\"o}m, Ulf and Florin, Ann-Britt and {\v{S}}ics, Ivo and Casini, Michele},
  journal={Hydrobiologia},
  volume={847},
  pages={2541--2553},
  year={2020},
  publisher={Springer},
  doi={10.1007/s10750-020-04272-4}
}

@article{koster2017eastern,
  title={Eastern {B}altic cod recruitment revisited—dynamics and impacting factors},
  author={K{\"o}ster, Friedrich W and Huwer, Bastian and Hinrichsen, Hans-Harald and Neumann, Viola and Makarchouk, Andrei and Eero, Margit and Dewitz, Burkhard V and H{\"u}ssy, Karin and Tomkiewicz, Jonna and Margonski, Piotr and Temming, Axel and Hermann, Jens-Peter and Oesterwind, Daniel and Dierking, Jan and Kotterba, Paul and Plikshs, Maris},
  journal={ICES Journal of Marine Science},
  volume={74},
  number={1},
  pages={3--19},
  year={2017},
  publisher={Oxford University Press},
  doi={10.1093/icesjms/fsw172}
}

@article{orio2019spatial,
  title={Spatial contraction of demersal fish populations in a large marine ecosystem},
  author={Orio, Alessandro and Bergstr{\"o}m, Ulf and Florin, Ann-Britt and Lehmann, Andreas and {\v{S}}ics, Ivo and Casini, Michele},
  journal={Journal of Biogeography},
  volume={46},
  number={3},
  pages={633--645},
  year={2019},
  publisher={Wiley Online Library},
  doi={10.1111/jbi.13510}
}

@article{limburg2018effect,
  title={Effect of {M}arine {H}ypoxia on {B}altic {S}ea {C}od \textit{{G}adus morhua}: evidence {F}rom {O}tolith {C}hemical {P}roxies},
  author={Limburg, Karin E and Casini, Michele},
  journal={Frontiers in Marine Science},
  volume={5},
  number={482},
  year={2018},
  publisher={Frontiers},
  doi={10.3389/fmars.2018.00482}
}

@article{limburg2011tracking,
  title={Tracking {B}altic hypoxia and cod migration over millennia with natural tags},
  author={Limburg, Karin E and Olson, Carina and Walther, Yvonne and Dale, Darren and Slomp, Caroline P and H{\o}ie, Hans},
  journal={Proceedings of the National Academy of Sciences},
  volume={108},
  number={22},
  pages={E177--E182},
  year={2011},
  publisher={National Academy of Sciences},
  doi={10.1073/pnas.1100684108}
}

@article{haase2020diet,
  title={Diet of dominant demersal fish species in the {B}altic {S}ea: Is flounder stealing benthic food from cod?},
  author={Haase, Kevin and Orio, Alessandro and Pawlak, Joanna and Pachur, Marzenna and Casini, Michele},
  journal={Marine Ecology Progress Series},
  volume={645},
  pages={159--170},
  year={2020},
  doi={10.3354/meps13360}
}

@article{casini2016hypoxic,
  title={Hypoxic areas, density-dependence and food limitation drive the body condition of a heavily exploited marine fish predator},
  author={Casini, Michele and K{\"a}ll, Filip and Hansson, Martin and Plikshs, Maris and Baranova, Tatjana and Karlsson, Olle and Lundstr{\"o}m, Karl and Neuenfeldt, Stefan and G{\aa}rdmark, Anna and Hjelm, Joakim},
  journal={Royal Society open science},
  volume={3},
  number={160416},
  year={2016},
  publisher={The Royal Society},
  doi={10.1098/rsos.160416}
}

@article{sandblom2016physiological,
  title={Physiological constraints to climate warming in fish follow principles of plastic floors and concrete ceilings},
  author={Sandblom, Erik and Clark, Timothy D and Gr{\"a}ns, Albin and Ekstr{\"o}m, Andreas and Brijs, Jeroen and Sundstr{\"o}m, L Fredrik and Odelstr{\"o}m, Anne and Adill, Anders and Aho, Teija and Jutfelt, Fredrik},
  journal={Nature Communications},
  volume={7},
  number={11447},
  year={2016},
  publisher={Nature Publishing Group},
  doi={10.1038/ncomms11447}
}

@article{baird2004consequences,
  title={{CONSEQUENCES OF HYPOXIA ON ESTUARINE ECOSYSTEM FUNCTION: ENERGY DIVERSION FROM CONSUMERS TO MICROBES}},
  author={Baird, Daniel and Christian, Robert R and Peterson, Charles H and Johnson, Galen A},
  journal={Ecological Applications},
  volume={14},
  number={3},
  pages={805--822},
  year={2004},
  publisher={Wiley Online Library},
  doi={10.1890/02-5094}
}

@article{eby2005habitat,
  title={Habitat degradation from intermittent hypoxia: {i}mpacts on demersal fishes},
  author={Eby, Lisa A and Crowder, Larry B and McClellan, Catherine M and Peterson, Charles H and Powers, Monica J},
  journal={Marine Ecology Progress Series},
  volume={291},
  pages={249--262},
  year={2005},
  doi={10.3354/meps291249}
}

@article{lowell1999cumulative,
  title={Cumulative effects of multiple effluent and low dissolved oxygen stressors on mayflies at cold temperatures},
  author={Lowell, Richard B and Culp, Joseph M},
  journal={Canadian Journal of Fisheries and Aquatic Sciences},
  volume={56},
  number={9},
  pages={1624--1630},
  year={1999},
  publisher={NRC Research Press Ottawa, Canada},
  doi={10.1139/f99-091}
}

@article{wootton2005measurement,
  title={Measurement of interaction strength in nature},
  author={Wootton, J Timothy and Emmerson, Mark},
  journal={Annual Review of Ecology, Evolution, and Systematics},
  volume={36},
  pages={419--444},
  year={2005},
  publisher={Annual Reviews},
  doi={10.1146/annurev.ecolsys.36.091704.175535}
}

@article{audzijonyte2019oxygen,
  title={Is oxygen limitation in warming waters a valid mechanism to explain decreased body sizes in aquatic ectotherms?},
  author={Audzijonyte, Asta and Barneche, Diego R and Baudron, Alan R and Belmaker, Jonathan and Clark, Timothy D and Marshall, C Tara and Morrongiello, John R and van Rijn, Itai},
  journal={Global Ecology and Biogeography},
  volume={28},
  number={2},
  pages={64--77},
  year={2019},
  publisher={Wiley Online Library},
  doi={10.1111/geb.12847}
}

@article{deutsch2020metabolic,
  title={Metabolic trait diversity shapes marine biogeography},
  author={Deutsch, Curtis and Penn, Justin L and Seibel, Brad},
  journal={Nature},
  volume={585},
  pages={557--562},
  year={2020},
  publisher={Nature Publishing Group},
  doi={10.1038/s41586-020-2721-y}
}

@article{ekau2010impacts,
  title={Impacts of hypoxia on the structure and processes in pelagic communities (zooplankton, macro-invertebrates and fish)},
  author={Ekau, Werner and Auel, Holger and P{\"o}rtner, H-O and Gilbert, Denis},
  journal={Biogeosciences},
  volume={7},
  number={5},
  pages={1669--1699},
  year={2010},
  publisher={Copernicus GmbH},
  doi={10.5194/bg-7-1669-2010}
}

@article{jordan2007effects,
  title={Effects of {R}ation {S}ize and {H}ypoxia on {S}pecific {D}ynamic {A}ction in the {C}od},
  author={Jordan, Anders D and Steffensen, John F},
  journal={Physiological and Biochemical Zoology},
  volume={80},
  number={2},
  pages={178--185},
  year={2007},
  publisher={The University of Chicago Press},
  note={PMID: 17252514},
  doi={10.1086/510565}
}

@article{gislason2010size,
  title={Size, growth, temperature and the natural mortality of marine fish},
  author={Gislason, Henrik and Daan, Niels and Rice, Jake C and Pope, John G},
  journal={Fish and Fisheries},
  volume={11},
  number={2},
  pages={149--158},
  year={2010},
  publisher={Wiley Online Library},
  doi={10.1111/j.1467-2979.2009.00350.x}
}

@article{draganik2007status,
  title={Status of exploited {B}altic flounder stocks in the southern {B}altic area ({ICES} {SD} 26)},
  author={Draganik, B and Ivanow, S and Tomczak, Maciej and Maksimov, B and Psuty-Lipska, I},
  journal={Oceanological and Hydrobiological Studies},
  volume={36},
  number={4},
  pages={47--64},
  year={2007},
  publisher={Versita},
  doi={10.2478/v10009-007-0029-y}
}

@article{casini2019spatio,
  title={Spatio-temporal dynamics and behavioural ecology of a ``demersal'' fish population as detected using research survey pelagic trawl catches: {t}he {E}astern {B}altic {S}ea cod (\textit{{G}adus morhua})},
  author={Casini, Michele and Tian, Huidong and Hansson, Martin and Grygiel, W{\l}odzimierz and Strods, Guntars and Statkus, Romas and Sepp, Elor and Gr{\"o}hsler, Tomas and Orio, Alessandro and Larson, Niklas},
  journal={ICES Journal of Marine Science},
  volume={76},
  number={6},
  pages={1591--1600},
  year={2019},
  publisher={Oxford University Press},
  doi={10.1093/icesjms/fsz016}
}

@article{casini2004feeding,
  title={Feeding preferences of herring (\textit{{C}lupea harengus}) and sprat (\textit{{S}prattus sprattus}) in the southern {B}altic {S}ea},
  author={Casini, Michele and Cardinale, Massimiliano and Arrhenius, Fredrik},
  journal={ICES Journal of Marine Science},
  volume={61},
  number={8},
  pages={1267--1277},
  year={2004},
  publisher={Oxford University Press},
  doi={10.1016/j.icesjms.2003.12.011}
}

@article{kramer1987dissolved,
  title={Dissolved oxygen and fish behavior},
  author={Kramer, Donald L},
  journal={Environmental Biology of Fishes},
  volume={18},
  number={2},
  pages={81--92},
  year={1987},
  publisher={Springer},
  doi={10.1007/BF00002597}
}

@article{jerde2019strong,
  title={Strong {E}vidence for an {I}ntraspecific {M}etabolic {S}caling {C}oefficient {N}ear 0.89 in {F}ish},
  author={Jerde, Christopher L and Kraskura, Krista and Eliason, Erika J and Csik, Samantha R and Stier, Adrian C and Taper, Mark L},
  journal={Frontiers in Physiology},
  volume={10},
  number={1166},
  year={2019},
  publisher={Frontiers},
  doi={10.3389/fphys.2019.01166}
}

@article{pan2016hypoxia,
  title={Hypoxia tolerance decreases with body size in red drum \textit{{S}ciaenops ocellatus}},
  author={Pan, YK and Ern, R and Esbaugh, AJ},
  journal={Journal of Fish Biology},
  volume={89},
  number={2},
  pages={1488--1493},
  year={2016},
  publisher={Wiley Online Library},
  doi={10.1111/jfb.13035}
}

@book{nelson2004biological,
  title={Biological {P}hysics},
  author={Nelson, Philip},
  year={2004},
  publisher={WH Freeman New York}
}

@techreport{smhi2020oxygen,
  title={Oxygen Survey in the Baltic Sea 2020 - Extent of Anoxia and Hypoxia, 1960-2020},
  author={Hansson, Martin and Viktorsson, Lena},
  journal={Report Oceanography},
  number={70},
  institution={Swedish Meteorological and Hydrological Institute},
  address={G\"{o}teborg, Sweden},
  year={2020}
}

@article{seibel2020oxygen,
  title={Oxygen supply capacity in animals evolves to meet maximum demand at the current oxygen partial pressure regardless of size or temperature},
  author={Seibel, Brad A and Deutsch, Curtis},
  journal={Journal of Experimental Biology},
  volume={223},
  number={12},
  pages={jeb210492},
  year={2020},
  publisher={The Company of Biologists Ltd},
  doi={10.1242/jeb.210492}
}

@article{brewer2016ocean,
  title={Ocean chemistry, ocean warming, and emerging hypoxia: Commentary},
  author={Brewer, Peter G and Peltzer, Edward T},
  journal={Journal of Geophysical Research: Oceans},
  volume={121},
  number={5},
  pages={3659--3667},
  year={2016},
  publisher={Wiley Online Library},
  doi={10.1002/2016JC011651}
}

@article{rose2009does,
  title={Does hypoxia have population-level effects on coastal fish? Musings from the virtual world},
  author={Rose, Kenneth A and Adamack, Aaron T and Murphy, Cheryl A and Sable, Shaye E and Kolesar, Sarah E and Craig, J Kevin and Breitburg, Denise L and Thomas, Peter and Brouwer, Marius H and Cerco, Carl F and Diamond, Sandra},
  journal={Journal of Experimental Marine Biology and Ecology},
  volume={381},
  pages={S188--S203},
  year={2009},
  publisher={Elsevier},
  doi={10.1016/j.jembe.2009.07.022}
}

@article{luo2001spatially,
  title={A spatially-explicit approach for estimating carrying capacity: an application for the Atlantic menhaden (\textit{{B}revoortia tyrannus}) in {C}hesapeake {B}ay},
  author={Luo, Jiangang and Hartman, Kyle J and Brandt, Stephen B and Cerco, Carl F and Rippetoe, Thomas H},
  journal={Estuaries},
  volume={24},
  number={4},
  pages={545--556},
  year={2001},
  publisher={Springer},
  doi={10.2307/1353256}
}

@article{tuong2018clown,
  title={Clown knifefish (\textit{{C}hitala ornata}) oxygen uptake and its partitioning in present and future temperature environments},
  author={Tuong, Dang Diem and Ngoc, Tran Bao and Huynh, Vo Thi Nhu and Phuong, Nguyen Thanh and Hai, Tran Ngoc and Wang, Tobias and Bayley, Mark},
  journal={Comparative Biochemistry and Physiology Part A: Molecular \& Integrative Physiology},
  volume={216},
  pages={52--59},
  year={2018},
  publisher={Elsevier},
  doi={10.1016/j.cbpa.2017.11.018}
}

@article{alvarez2020growth,
  title={Growth performance, physiological responses to hypoxia and flesh quality of {C}hilean croaker (\textit{{C}ilus gilberti}) stocked at different densities},
  author={{\'A}lvarez, Claudio A and Jerez-Cepa, Ismael and C{\'a}rcamo, Claudia B and Toledo, Pedro and Flores, H{\'e}ctor and Brokordt, Katherina},
  journal={Aquaculture},
  volume={525},
  number={735316},
  year={2020},
  publisher={Elsevier},
  doi={10.1016/j.aquaculture.2020.735316}
}

@article{calder2015mesocosm,
  title={A mesocosm study investigating the effects of hypoxia and population density on respiration and reproductive biology in the brittlestar \textit{{A}mphiura filiformis}},
  author={Calder-Potts, R and Spicer, JI and Calosi, P and Findlay, HS and Widdicombe, S},
  journal={Marine Ecology Progress Series},
  volume={534},
  pages={135--147},
  year={2015},
  doi={10.3354/meps11379}
}

@article{killen2012relationship,
  title={A relationship between metabolic rate and risk-taking behaviour is revealed during hypoxia in juvenile {E}uropean sea bass},
  author={Killen, Shaun S and Marras, Stefano and Ryan, Mary R and Domenici, Paolo and McKenzie, David J},
  journal={Functional Ecology},
  volume={26},
  number={1},
  pages={134--143},
  year={2012},
  publisher={Wiley Online Library},
  doi={10.1111/j.1365-2435.2011.01920.x}
}

@article{maciak2010repeatability,
  title={Repeatability of standard metabolic rate ({SMR}) in a small fish, the spined loach (\textit{{C}obitis taenia})},
  author={Maciak, Sebastian and Konarzewski, Marek},
  journal={Comparative Biochemistry and Physiology Part A: Molecular \& Integrative Physiology},
  volume={157},
  number={2},
  pages={136--141},
  year={2010},
  publisher={Elsevier},
  doi={10.1016/j.cbpa.2010.05.017}
}

@article{muusze1998hypoxia,
  title={Hypoxia tolerance of {A}mazon fish: respirometry and energy metabolism of the cichlid \textit{{A}stronotus ocellatus}},
  author={Muusze, Bie and Marcon, Jaydione and van den Thillart, Guido and Almeida-Val, Vera},
  journal={Comparative Biochemistry and Physiology Part A: Molecular \& Integrative Physiology},
  volume={120},
  number={1},
  pages={151--156},
  year={1998},
  publisher={Elsevier},
  doi={10.1016/S1095-6433(98)10023-5}
}

@online{Rohatgi2020,
  url = {https://automeris.io/WebPlotDigitizer},
  author = {Rohatgi,  Ankit},
  title = {Webplotdigitizer: Version 4.5},
  year = {2021}
}

@manual{hernroth1985zoop,
  title={Recommendations on methods for marine biological studies in the Baltic Sea},
  editor={Hernroth, L.},
  number={10},
  organization={Baltic Marine Biologists:Working Group 9},
  address={Institute of Marine Research, Lysekill, Sweden},
  edition={Working Group 14},
  year={1985},
  issn={0282-8839}
}

@article{mollmann2002population,
  title={Population dynamics of calanoid copepods and the implications of their predation by clupeid fish in the {C}entral {B}altic {S}ea},
  author={M{\"o}llmann, Christian and K{\"o}ster, Friedrich W},
  journal={Journal of Plankton Research},
  volume={24},
  number={10},
  pages={959--978},
  year={2002},
  publisher={Oxford University Press},
  doi={10.1093/plankt/24.10.959}
}

@article{mollmann1999food,
  title={Food consumption by clupeids in the {C}entral {B}altic: evidence for top-down control?},
  author={M{\"o}llmann, Christian and K{\"o}ster, Friedrich},
  journal={ICES Journal of Marine Science},
  volume={56},
  pages={100--113},
  year={1999},
  publisher={ICES},
  doi={10.1006/jmsc.1999.0630}
}

@inbook{snoeijs2017patterns,
  title={Patterns of biodiversity},
  author={Snoeijs-Leijonmalm, Pauline},
  editor={Snoeijs-Leijonmalm, Pauline and Schubert, Hendrik and Radziejewska, Teresa},
  booktitle={Biological Oceanography of the Baltic Sea},
  pages={123--191},
  year={2017},
  publisher={Springer Netherlands},
  doi={10.1007/978-94-007-0668-2_4}
}

@article{hill1992seasonal,
  title={Seasonal changes in lipid content and composition in the benthic amphipods \textit{{M}onoporeia affinis} and \textit{{P}ontoporeia femorata}},
  author={Hill, Cathy and Quigley, Michael A and Cavaletto, Joann F and Gordon, Wendy},
  journal={Limnology and Oceanography},
  volume={37},
  number={6},
  pages={1280--1289},
  year={1992},
  publisher={Wiley Online Library},
  doi={10.4319/lo.1992.37.6.1280}
}

@article{barnes2010global,
  title={Global patterns in predator--prey size relationships reveal size dependency of trophic transfer efficiency},
  author={Barnes, Carolyn and Maxwell, David and Reuman, Daniel C and Jennings, Simon},
  journal={Ecology},
  volume={91},
  number={1},
  pages={222--232},
  year={2010},
  publisher={Wiley Online Library},
  doi={10.1890/08-2061.1}
}

@article{tsai2016predator,
  title={Predator--prey mass ratio revisited: {d}oes preference of relative prey body size depend on individual predator size?},
  author={Tsai, Cheng-Han and Hsieh, Chih-hao and Nakazawa, Takefumi},
  journal={Functional Ecology},
  volume={30},
  number={12},
  pages={1979--1987},
  year={2016},
  publisher={Wiley Online Library},
  doi={10.1111/1365-2435.12680}
}

@article{vainikka2009two,
  title={Two-and three-dimensional maturation reaction norms for the eastern {B}altic cod, \textit{{G}adus morhua}},
  author={Vainikka, Anssi and G{\aa}rdmark, Anna and Bland, Barbara and Hjelm, Joakim},
  journal={ICES Journal of Marine Science},
  volume={66},
  number={2},
  pages={248--257},
  year={2009},
  publisher={Oxford University Press},
  doi={10.1093/icesjms/fsn199}
}

@article{barot2003estimating,
  title={Estimating {R}eaction {N}orms for {A}ge and {S}ize at {M}aturation {W}hen {A}ge at {F}irst {R}eproduction is {U}nknown},
  author={Barot, S{\'e}bastien and Heino, Mikko and O'Brien, Loretta and Dieckmann, Ulf},
  year={2003},
  publisher={IR-03-043},
  url={https://pure.iiasa.ac.at/7040}
}

@manual{bitsmanual2017,
  title={Manual for the {B}altic {I}nternational {T}rawl {S}urveys},
  series={Series of ICES Survey Protocols SISP 7 -- BITS},
  organization={ICES Baltic International Fish Survey Working Group},
  number={Version 2.0},
  year={2017},
  pages={95},
  issn={2304-6252},
  DOI={10.17895/ices.pub.2883}
}

@article{grygiel2002sexual,
  title={Sexual maturation of the southern {B}altic herring and sprat (1980-2001)},
  author={Grygiel, W{\l}odzimierz and Wyszy{\'n}ski, Miros{\l}aw},
  journal={International Council Meeting},
  year={2002}
}

@article{mackay1990note,
  title={A note on the inverse function of the von Bertalanffy growth function},
  author={Mackay, D and Moreau, J},
  year={1990},
  volume={8},
  number={1},
  pages={28--31},
  publisher={ICLARM},
  url={https://hdl.handle.net/20.500.12348/3147}
}

@article{gavin2019levenberg,
  title={The Levenberg-Marquardt algorithm for nonlinear least squares curve-fitting problems},
  author={Gavin, Henri P},
  journal={Department of Civil and Environmental Engineering, Duke University},
  pages={1--19},
  year={2019}
}

@article{carpenter2017stan,
  title={Stan: A probabilistic programming language},
  author={Carpenter, Bob and Gelman, Andrew and Hoffman, Matthew D and Lee, Daniel and Goodrich, Ben and Betancourt, Michael and Brubaker, Marcus and Guo, Jiqiang and Li, Peter and Riddell, Allen},
  journal={Journal of Statistical Software},
  volume={76},
  number={1},
  pages={1--32},
  year={2017},
  publisher={Columbia University, New York, NY (United States); Harvard University, Cambridge, MA (United State)},
  doi={10.18637/jss.v076.i01}
}

@Manual{cmdstanr,
    title = {cmdstanr: R Interface to 'CmdStan'},
    author = {Jonah Gabry and Rok Cešnovar},
    year = {2021},
    url = {https://mc-stan.org/cmdstanr},
}

@techreport{wgbifs2020,
  title={Baltic {F}isheries {A}ssessment {W}orking {G}roup ({WGBFAS})},
  author={ICES},
  institution={ICES Scientific Reports},
  volume={2},
  number={45},
  pages={643},
  year={2020},
  doi={10.17895/ices.pub.6024}
}

@article{shepherd1999extended,
  title={Extended survivors analysis: an improved method for the analysis of catch-at-age data and abundance indices},
  author={Shepherd, JG},
  journal={ICES Journal of Marine Science},
  volume={56},
  number={5},
  pages={584--591},
  year={1999},
  publisher={Oxford University Press},
  doi={10.1006/jmsc.1999.0498}
}

@misc{pcritdatabase2021,
  title={Database of critical oxygen level ({P}\textsubscript{crit}) in freshwater and marine fishes 1974--2015},
  author={Rogers, Nicholas J and Urbina, Mauricio A and Reardon, Erin E and McKenzie, David J and Birchenough, Silvana NR and Wilson, Rod W},
  address={Cefas, UK},
  version={1},
  year={2021},
  doi={10.14466/CefasDataHub.121}
}

@article{lindmark2021bottom,
  author = {Lindmark, Max and Audzijonyte, Asta and Blanchard, Julia L. and Gårdmark, Anna},
  title = {Temperature impacts on fish physiology and resource abundance lead to faster growth but smaller fish sizes and yields under warming},
  journal = {Global Change Biology},
  volume = {28},
  number = {21},
  pages = {6239--6253},
  doi = {10.1111/gcb.16341},
  year = {2022}
}

@article{byrd1995limited,
  title={A limited memory algorithm for bound constrained optimization},
  author={Byrd, Richard H and Lu, Peihuang and Nocedal, Jorge and Zhu, Ciyou},
  journal={SIAM Journal on Scientific Computing},
  volume={16},
  number={5},
  pages={1190--1208},
  year={1995},
  publisher={SIAM},
  doi={10.1137/0916069}
}

@article{hrycik2017sub,
  title={Sub-lethal effects on fish provide insight into a biologically-relevant threshold of hypoxia},
  author={Hrycik, Allison R and Almeida, L Zoe and H{\"o}{\"o}k, Tomas O},
  journal={Oikos},
  volume={126},
  number={3},
  pages={307--317},
  year={2017},
  publisher={Wiley Online Library},
  doi={10.1111/oik.03678}
}

@article{killen2016ecological,
  title={Ecological {I}nfluences and {M}orphological {C}orrelates of {R}esting and {M}aximal {M}etabolic {R}ates across {T}eleost {F}ish {S}pecies},
  author={Killen, Shaun S and Glazier, Douglas S and Rezende, Enrico L and Clark, Timothy D and Atkinson, David and Willener, Astrid ST and Halsey, Lewis G},
  journal={The American Naturalist},
  volume={187},
  number={5},
  pages={592--606},
  year={2016},
  publisher={University of Chicago Press Chicago, IL},
  doi={10.1086/685893}
}

@article{wood2016smoothing,
  title={Smoothing {P}arameter and {M}odel {S}election for {G}eneral {S}mooth {M}odels},
  author={Wood, Simon N and Pya, Natalya and S{\"a}fken, Benjamin},
  journal={Journal of the American Statistical Association},
  volume={111},
  number={516},
  pages={1548--1563},
  year={2016},
  publisher={Taylor \& Francis},
  doi={10.1080/01621459.2016.1180986}
}

@article{nilsson2008does,
  title={Does size matter for hypoxia tolerance in fish?},
  author={Nilsson, G{\"o}ran E and {\"O}stlund-Nilsson, Sara},
  journal={Biological Reviews},
  volume={83},
  number={2},
  pages={173--189},
  year={2008},
  publisher={Wiley Online Library},
  doi={10.1111/j.1469-185X.2008.00038.x}
}

@book{beverton2012dynamics,
  title={On the {D}ynamics of {E}xploited {F}ish {P}opulations},
  author={Beverton, Raymond JH and Holt, Sidney J},
  volume={11},
  year={2012},
  publisher={Springer Science \& Business Media}
}

@Manual{rmanual,
    title = {R: A Language and Environment for Statistical Computing},
    author = {{R Core Team}},
    organization = {R Foundation for Statistical Computing},
    address = {Vienna, Austria},
    year = {2021},
    url = {https://www.R-project.org/},
    note = {v. 4.2.2}
  }
  
@article{karim2003model,
  title={A model of fish preference and mortality under hypoxic water in the coastal environment},
  author={Karim, Md Rezaul and Sekine, Masahiko and Ukita, Masao},
  journal={Marine Pollution Bulletin},
  volume={47},
  number={1},
  pages={25--29},
  year={2003},
  publisher={Elsevier},
  doi={10.1016/S0025-326X(02)00409-5},
  note = {Environmental Management of Enclosed Coastal Seas},
}

@article{shimps2005hypoxia,
  title={Hypoxia tolerance in two juvenile estuary-dependent fishes},
  author={Shimps, Elizabeth L and Rice, James A and Osborne, Jason A},
  journal={Journal of Experimental Marine Biology and Ecology},
  volume={325},
  number={2},
  pages={146--162},
  year={2005},
  publisher={Elsevier},
  doi={10.1016/j.jembe.2005.04.026}
}

@article{small2014hypoxia,
  title={Hypoxia, {B}lackwater and {F}ish {K}ills: experimental {L}ethal {O}xygen {T}hresholds in {J}uvenile {P}redatory {L}owland {R}iver {F}ishes},
  author={Small, Kade and Kopf, R Keller and Watts, Robyn J and Howitt, Julia},
  journal={PLoS ONE},
  volume={9},
  pages={1--10},
  year={2014},
  publisher={Public Library of Science San Francisco, USA},
  doi={10.1371/journal.pone.0094524}
}

@article{diaz2001overview,
  title={Overview of {H}ypoxia around the {W}orld},
  author={Diaz, Robert J},
  journal={Journal of Environmental Quality},
  volume={30},
  number={2},
  pages={275--281},
  year={2001},
  publisher={Wiley Online Library},
  doi={10.2134/jeq2001.302275x}
}

@article{gray2002effects,
  title={Effects of hypoxia and organic enrichment on the coastal marine environment},
  author={Gray, John S and Wu, Rudolf Shiu-sun and Or, Ying Ying},
  journal={Marine Ecology Progress Series},
  volume={238},
  pages={249--279},
  year={2002},
  doi={10.3354/meps238249}
}

@article{neuenfeldt2020feeding,
  title={Feeding and growth of {A}tlantic cod (\textit{{G}adus morhua} L.) in the eastern {B}altic {S}ea under environmental change},
  author={Neuenfeldt, Stefan and Bartolino, Valerio and Orio, Alessandro and Andersen, Ken H and Andersen, Niels G and Niiranen, Susa and Bergstr{\"o}m, Ulf and Ustups, Didzis and Kulatska, Nataliia and Casini, Michele},
  journal={ICES Journal of Marine Science},
  volume={77},
  number={2},
  pages={624--632},
  year={2020},
  publisher={Oxford University Press},
  doi={10.1093/icesjms/fsz248}
}

@article{neuenfeldt2006environmentally,
  title={Environmentally driven predator--prey overlaps determine the aggregate diet of the cod \textit{{G}adus morhua} in the {B}altic {S}ea},
  author={Neuenfeldt, Stefan and Beyer, Jan E},
  journal={Marine Ecology Progress Series},
  volume={310},
  pages={151--163},
  year={2006},
  doi={10.3354/meps310151}
}

@incollection{casini2013spatio,
  title={Spatio-temporal ecosystem shifts in the Baltic Sea: top-down control and reversibility potential},
  author={Casini, Michele},
  booktitle={Advances in Environmental Research},
  volume={28},
  pages={149--167},
  year={2013},
  publisher={Nova Science Publishers},
  address={New York},
  editor={Daniels, Justin A}
}

@article{zhu2013strategies,
  title={Strategies for hypoxia adaptation in fish species: {a} review},
  author={Zhu, Chang-Dong and Wang, Zhen-Hua and Yan, Biao},
  journal={Journal of Comparative Physiology B},
  volume={183},
  number={8},
  pages={1005--1013},
  year={2013},
  publisher={Springer},
  doi={10.1007/s00360-013-0762-3}
}

@article{urbina2013relationship,
  title={Relationship between {F}ish {S}ize and {M}etabolic {R}ate in the {O}xyconforming {I}nanga \textit{{G}alaxias maculatus} {R}eveals {S}ize-{D}ependent {S}trategies to {W}ithstand {H}ypoxia},
  author={Urbina, Mauricio A and Glover, Chris N},
  journal={Physiological and Biochemical Zoology},
  volume={86},
  number={6},
  pages={740--749},
  year={2013},
  publisher={University of Chicago Press Chicago, IL},
  doi={10.1086/673727}
}

@incollection{farrell2009defining,
  title={Defining {H}ypoxia: an {I}ntegrative {S}ynthesis of the {R}esponses of {F}ish to {H}ypoxia},
  author={Farrell, Anthony P and Richards, Jeffrey G},
  booktitle={Fish Physiology},
  volume={27},
  pages={487--503},
  year={2009},
  publisher={Elsevier},
  doi={10.1016/S1546-5098(08)00011-3}
}

@article{carstensen2019baltic,
  title={Baltic {S}ea {H}ypoxia {T}akes {M}any {S}hapes and {S}izes},
  author={Carstensen, Jacob and Conley, Daniel J},
  journal={Limnology and Oceanography Bulletin},
  volume={28},
  number={4},
  pages={125--129},
  year={2019},
  publisher={Wiley Online Library},
  doi={10.1002/lob.10350}
}

@article{pollock2007effects,
  title={The effects of hypoxia on fishes: {f}rom ecological relevance to physiological effects},
  author={Pollock, MS and Clarke, LMJ and Dub{\'e}, MG},
  journal={Environmental Reviews},
  volume={15},
  number={NA},
  pages={1--14},
  year={2007},
  dio={10.1139/a06-006}
}

@article{orio2022deoxygenation,
  title={Deoxygenation impacts on {B}altic {S}ea cod: Dramatic declines in ecosystem services of an iconic keystone predator},
  author={Orio, Alessandro and Heimbrand, Yvette and Limburg, Karin},
  journal={Ambio},
  volume={51},
  number={3},
  pages={626--637},
  year={2022},
  publisher={Springer},
  doi={10.1007/s13280-021-01572-4}
}

@article{froehlich2017does,
  title={When does hypoxia affect management performance of a fishery? {A} management strategy evaluation of {D}ungeness crab (\textit{{M}etacarcinus magister}) fisheries in {H}ood {C}anal, {W}ashington, {USA}},
  author={Froehlich, Halley E and Essington, Timothy E and McDonald, P Sean},
  journal={Canadian Journal of Fisheries and Aquatic Sciences},
  volume={74},
  number={6},
  pages={922--932},
  year={2017},
  publisher={NRC Research Press},
  doi={10.1139/cjfas-2016-0269}
}

@article{zhang2014effects,
  title={Effects of hypoxia on habitat quality of pelagic planktivorous fishes in the northern {G}ulf of {M}exico},
  author={Zhang, Hongyan and Mason, Doran M and Stow, Craig A and Adamack, Aaron T and Brandt, Stephen B and Zhang, Xinsheng and Kimmel, David G and Roman, Michael R and Boicourt, William C and Ludsin, Stuart A},
  journal={Marine Ecology Progress Series},
  volume={505},
  pages={209--226},
  year={2014},
  doi={10.3354/meps10768}
}

@article{chu2017ecophysiological,
  title={Ecophysiological limits to aerobic metabolism in hypoxia determine epibenthic distributions and energy sequestration in the northeast {P}acific ocean},
  author={Chu, Jackson WF and Gale, Katie SP},
  journal={Limnology and Oceanography},
  volume={62},
  number={1},
  pages={59--74},
  year={2017},
  publisher={Wiley Online Library},
  doi={10.1002/lno.10370}
}

@article{slater2020fewer,
  title={Fewer {C}opepods, {F}ewer {A}nchovies, and {M}ore {J}ellyfish: How {D}oes {H}ypoxia {I}mpact the {C}hesapeake {B}ay {Z}ooplankton {C}ommunity?},
  author={Slater, Wencheng L and Pierson, James J and Decker, Mary Beth and Houde, Edward D and Lozano, Carlos and Seuberling, James},
  journal={Diversity},
  volume={12},
  number={1},
  pages={35},
  year={2020},
  publisher={Multidisciplinary Digital Publishing Institute},
  doi={10.3390/d12010035}
}

@article{koenigstein2016modelling,
  title={Modelling climate change impacts on marine fish populations: {p}rocess-based integration of ocean warming, acidification and other environmental drivers},
  author={Koenigstein, Stefan and Mark, Felix C and G{\"o}{\ss}ling-Reisemann, Stefan and Reuter, Hauke and Poertner, Hans-Otto},
  journal={Fish and Fisheries},
  volume={17},
  number={4},
  pages={972--1004},
  year={2016},
  publisher={Wiley Online Library},
  doi={10.1111/faf.12155}
}

@article{riedel2014effect,
  title={Effect of hypoxia and anoxia on invertebrate behaviour: {e}cological perspectives from species to community level},
  author={Riedel, Bettina and Pados, Theodora and Pretterebner, Katrin and Schiemer, L and Steckbauer, Alexandra and Haselmair, Alexandra and Zuschin, Martin and Stachowitsch, Michael},
  journal={Biogeosciences},
  volume={11},
  number={6},
  pages={1491--1518},
  year={2014},
  publisher={Copernicus GmbH},
  doi={10.5194/bg-11-1491-2014}
}

@article{pollock2007effects,
  title={The effects of hypoxia on fishes: {f}rom ecological relevance to physiological effects},
  author={Pollock, MS and Clarke, LMJ and Dub{\'e}, MG},
  journal={Environmental Reviews},
  volume={15},
  number={NA},
  pages={1--14},
  year={2007},
  doi={10.1139/a06-006}
}

@article{carstensen2014deoxygenation,
  title={Deoxygenation of the {B}altic {S}ea during the last century},
  author={Carstensen, Jacob and Andersen, Jesper H and Gustafsson, Bo G and Conley, Daniel J},
  journal={Proceedings of the National Academy of Sciences},
  volume={111},
  number={15},
  pages={5628--5633},
  year={2014},
  publisher={National Academy of Sciences},
  doi = {10.1073/pnas.1323156111}
}

@article{brown2004toward,
  title={{TOWARD A METABOLIC THEORY OF ECOLOGY}},
  author={Brown, James H and Gillooly, James F and Allen, Andrew P and Savage, Van M and West, Geoffrey B},
  journal={Ecology},
  volume={85},
  number={7},
  pages={1771--1789},
  year={2004},
  publisher={Wiley Online Library},
  doi={10.1890/03-9000}
}

@article{ultsch2019utility,
  title={The utility and determination of {P}\textsubscript{crit} in fishes},
  author={Ultsch, Gordon R and Regan, Matthew D},
  journal={Journal of Experimental Biology},
  volume={222},
  number={22},
  note={jeb203646},
  year={2019},
  publisher={The Company of Biologists Ltd},
  doi = {10.1242/jeb.203646}
}

@inproceedings{haahtela1990baltic,
  title={What do {B}altic studies tell us about the isopod \textit{{S}aduria entomon} (L.)?},
  author={Haahtela, Ilpo},
  booktitle={Annales Zoologici Fennici},
  volume={27},
  number={3},
  pages={269--278},
  year={1990},
  organization={JSTOR},
  url={https://www.jstor.org/stable/23736048}
}

@article{oschlies2018drivers,
  title={Drivers and mechanisms of ocean deoxygenation},
  author={Oschlies, Andreas and Brandt, Peter and Stramma, Lothar and Schmidtko, Sunke},
  journal={Nature Geoscience},
  volume={11},
  number={7},
  pages={467--473},
  year={2018},
  publisher={Nature Publishing Group},
  doi={10.1038/s41561-018-0152-2}
}

@article{pachur2013food,
  title={{FOOD COMPOSITION AND PREY SELECTION OF COD, \textit{GADUS MORHUA} (ACTINOPTERYGII: GADIFORMES: GADIDAE), IN THE SOUTHERN BALTIC SEA}},
  author={Pachur, Marzenna E and Horbowy, Jan},
  journal={Acta Ichthyologica et Piscatoria},
  volume={43},
  number={2},
  year={2013},
  pages={109--118},
  doi={10.3750/AIP2013.43.2.03}
}

@article{eero2012spatial,
  title={Spatial management of marine resources can enhance the recovery of predators and avoid local depletion of forage fish},
  author={Eero, Margit and Vinther, Morten and Haslob, Holger and Huwer, Bastian and Casini, Michele and Storr-Paulsen, Marie and K{\"o}ster, Friedrich W},
  journal={Conservation Letters},
  volume={5},
  number={6},
  pages={486--492},
  year={2012},
  publisher={Wiley Online Library},
  doi={10.1111/j.1755-263X.2012.00266.x}
}

@article{schmidtko2017decline,
  title={Decline in global oceanic oxygen content during the past five decades},
  author={Schmidtko, Sunke and Stramma, Lothar and Visbeck, Martin},
  journal={Nature},
  volume={542},
  number={7641},
  pages={335--339},
  year={2017},
  publisher={Nature Publishing Group},
  doi={10.1038/nature21399}
}

@online{sdg,
  author={{United Nations}},
  title={Department of Economic and Social Affairs, Sustainable Development},
  year={2022},
  url={https://sdgs.un.org/goals},
  urldate={2022-6-24}
}

@article{breitburg2003pattern,
  title={The pattern and influence of low dissolved oxygen in the {P}atuxent {R}iver, a seasonally hypoxic estuary},
  author={Breitburg, Denise L and Adamack, Aaron and Rose, Kenneth A and Kolesar, Sarah E and Decker, Beth and Purcell, Jennifer E and Keister, Julie E and Cowan, James H},
  journal={Estuaries},
  volume={26},
  number={2},
  pages={280--297},
  year={2003},
  publisher={Springer},
  doi={10.1007/BF02695967}
}

@article{buchheister2013patterns,
  title={Patterns and drivers of the demersal fish community of {C}hesapeake {B}ay},
  author={Buchheister, Andre and Bonzek, Christopher F and Gartland, James and Latour, Robert J},
  journal={Marine Ecology Progress Series},
  volume={481},
  pages={161--180},
  year={2013},
  doi={10.3354/meps10253}
}

@article{bergman2019scaling,
  title={Scaling from individual physiological measures to population-level demographic change: case studies and future directions for conservation management},
  author={Bergman, Jordanna N and Bennett, Joseph R and Binley, Allison D and Cooke, Steven J and Fyson, Vincent and Hlina, Benjamin L and Reid, Connor H and Vala, Michelle A and Madliger, Christine L},
  journal={Biological Conservation},
  volume={238},
  pages={108242},
  year={2019},
  publisher={Elsevier},
  doi={10.1016/j.biocon.2019.108242}
}

@article{breitburg1997varying,
  title={VARYING EFFECTS OF LOW DISSOLVED OXYGEN ON TROPHIC INTERACTIONS IN AN ESTUARINE FOOD WEB},
  author={Breitburg, Denise L and Loher, Timothy and Pacey, Carol A and Gerstein, Adam},
  journal={Ecological Monographs},
  volume={67},
  number={4},
  pages={489--507},
  year={1997},
  publisher={Wiley Online Library},
  doi={10.1890/0012-9615(1997)067[0489:VEOLDO]2.0.CO;2}
}

@article{breitburg1999linking,
  title={Linking water quality to larval survival: {p}redation mortality of fish larvae in an oxygen-stratified water column},
  author={Breitburg, Denise L and Rose, Kenneth A and Cowan Jr, James H},
  journal={Marine Ecology Progress Series},
  volume={178},
  pages={39--54},
  year={1999},
  doi={10.3354/meps178039}
}

@article{dwyer2014morphology,
  title={Morphology, metabolism and behaviour: {r}esponses of three fishes with different lifestyles to acute hypoxia},
  author={Dwyer, Georgia K and Stoffels, Rick J and Pridmore, Peter A},
  journal={Freshwater Biology},
  volume={59},
  number={4},
  pages={819--831},
  year={2014},
  publisher={Wiley Online Library},
  doi={10.1111/fwb.12306}
}

@incollection{snoeijs2017baltic,
  title={Why is the {B}altic {S}ea so special to live in?},
  author={Snoeijs-Leijonmalm, Pauline and Andr{\'e}n, Elinor},
  booktitle={Biological oceanography of the Baltic Sea},
  editor={Snoeijs-Leijonmalm, Pauline and Schubert, Hendrik and Radziejewska, Teresa},
  pages={23--84},
  year={2017},
  publisher={Springer},
  doi={10.1007/978-94-007-0668-2_2}
}

@article{mohrholz2018major,
  title={Major {B}altic {I}nflow {S}tatistics--{R}evised},
  author={Mohrholz, Volker},
  journal={Frontiers in Marine Science},
  volume={5},
  pages={384},
  year={2018},
  publisher={Frontiers Media SA},
  doi={10.3389/fmars.2018.00384}
}

@book{peters1986ecological,
  title={The ecological implications of body size},
  author={Peters, Robert Henry},
  volume={2},
  year={1986},
  publisher={Cambridge University Press}
}

@article{chabot2008environmental,
  title={Environmental hypoxia as a metabolic constraint on fish: the case of {A}tlantic cod, \textit{{G}adus morhua}},
  author={Chabot, D and Claireaux, G},
  journal={Marine Pollution Bulletin},
  volume={57},
  number={6},
  pages={287--294},
  year={2008},
  publisher={Elsevier},
  doi={10.1016/j.marpolbul.2008.04.001}
}

@incollection{richards2009metabolic,
  title={Metabolic and {M}olecular {R}esponses of {F}ish to {H}ypoxia},
  author={Richards, Jeffrey G},
  booktitle={Fish Physiology},
  volume={27},
  pages={443--485},
  year={2009},
  publisher={Elsevier},
  doi={10.1016/S1546-5098(08)00010-1}
}

@article{essington2001bertalanffy,
  title={The von {B}ertalanffy growth function, bioenergetics, and the consumption rates of fish},
  author={Essington, Timothy E and Kitchell, James F and Walters, Carl J},
  journal={Canadian Journal of Fisheries and Aquatic Sciences},
  volume={58},
  number={11},
  pages={2129--2138},
  year={2001},
  publisher={NRC Research Press Ottawa, Canada},
  doi={10.1139/f01-151}
}

@misc{froese2010fishbase,
  title={FishBase},
  editor={Froese, Rainer and Pauly, Daniel},
  year={2022},
  howpublished={www.fishbase.org},
}

@article{hagerman1997oxygen,
  title={Oxygen binding characteristics of haemocyanin in the {B}altic isopod \textit{{S}aduria entomon}},
  author={Hagerman, Lars and Vismann, Bent},
  journal={Ophelia},
  volume={46},
  number={2},
  pages={141--151},
  year={1997},
  publisher={Taylor \& Francis},
  doi={10.1080/00785326.1997.10432580}

}

@article{svedang2017historic,
  title={Historic changes in length distributions of three {B}altic cod (\textit{{G}adus morhua}) stocks: Evidence of growth retardation},
  author={Sved{\"a}ng, Henrik and Hornborg, Sara},
  journal={Ecology and evolution},
  volume={7},
  number={16},
  pages={6089--6102},
  year={2017},
  publisher={Wiley Online Library},
  doi={10.1002/ece3.3173}
}

@article{mollmann2004feeding,
  title={Feeding ecology of central {B}altic {S}ea herring and sprat},
  author={M{\"o}llmann, Christian and Kornilovs, Georgs and Fetter, M and K{\"o}ster, FW},
  journal={Journal of Fish Biology},
  volume={65},
  number={6},
  pages={1563--1581},
  year={2004},
  publisher={Wiley Online Library},
  doi={10.1111/j.0022-1112.2004.00566.x}
}

@article{tunney2014effects,
  title={Effects of differential habitat warming on complex communities},
  author={Tunney, Tyler D and McCann, Kevin S and Lester, Nigel P and Shuter, Brian J},
  journal={Proceedings of the National Academy of Sciences},
  volume={111},
  number={22},
  pages={8077--8082},
  year={2014},
  publisher={National Academy of Sciences},
  doi = {10.1073/pnas.1319618111}
}

@article{ludsin2009hypoxia,
  title={Hypoxia-avoidance by planktivorous fish in {C}hesapeake {B}ay: implications for food web interactions and fish recruitment},
  author={Ludsin, Stuart A and Zhang, Xinsheng and Brandt, Stephen B and Roman, Michael R and Boicourt, William C and Mason, Doran M and Costantini, Marco},
  journal={Journal of Experimental Marine Biology and Ecology},
  volume={381},
  pages={S121--S131},
  year={2009},
  publisher={Elsevier},
  doi={10.1016/j.jembe.2009.07.016}
}

@article{rose2019impacts,
  title={Impacts of ocean deoxygenation on fisheries},
  author={Rose, Kenneth A and Guti{\'e}rrez, Dimitri and Breitburg, Denise and Conley, Daniel and Craig, Kevin J and Froehlich, Halley E and Jeyabaskaran, R and Kripa, V and Mbaye, Baye Cheikh and Mohamed, KS and Padua, Shelton and Prema, D},
  year={2019},
  publisher={International Union for Conservation of Nature and Natural Resources (IUCN)}
}

@incollection{rose2019impacts,
  title={Impacts of ocean deoxygenation on fisheries},
  author={Rose, Kenneth A and Guti{\'e}rrez, Dimitri and Breitburg, Denise and Conley, Daniel and Craig, Kevin J and Froehlich, Halley E and Jeyabaskaran, R and Kripa, V and Mbaye, Baye Cheikh and Mohamed, KS and Padua, Shelton and Prema, D},
  booktitle={Ocean deoxygenation: Everyone’s problem Causes, impacts, consequences and solutions},
  year={2019},
  pages={519-544},
  publisher={International Union for Conservation of Nature and Natural Resources (IUCN)},
  address={Gland, Switzerland}
}

@article{limburg2019otolith,
  title={Otolith chemistry indicates recent worsened {B}altic cod condition is linked to hypoxia exposure},
  author={Limburg, Karin E and Casini, Michele},
  journal={Biology Letters},
  volume={15},
  number={12},
  pages={20190352},
  year={2019},
  publisher={The Royal Society},
  doi={10.1098/rsbl.2019.0352}
}

@article{breitburg2018declining,
  title={Declining oxygen in the global ocean and coastal waters},
  author={Breitburg, Denise and Levin, Lisa A and Oschlies, Andreas and Gr{\'e}goire, Marilaure and Chavez, Francisco P and Conley, Daniel J and Gar{\c{c}}on, V{\'e}ronique and Gilbert, Denis and Guti{\'e}rrez, Dimitri and Isensee, Kirsten and Jacinto, Gil S and Limburg, Karin E and Montes, Ivonne and Naqvi, SWA and Pitcher, Grant C and Rabalais, Nancy N and Roman, Michael R and Rose, Kenneth A and Seibel, Brad A and Telszewski, Maciej and Yasuhara, Moriaki and Zhang, Jing},
  journal={Science},
  volume={359},
  number={6371},
  pages = {eaam7240},
  year = {2018},
  doi = {10.1126/science.aam7240},
  publisher={American Association for the Advancement of Science}
}

@article{casini2011spatial,
  title={Spatial and temporal density dependence regulates the condition of central {B}altic {S}ea clupeids: {c}ompelling evidence using an extensive international acoustic survey},
  author={Casini, Michele and Kornilovs, Georgs and Cardinale, Massimiliano and M{\"o}llmann, Christian and Grygiel, Wlodzimierz and Jonsson, Patrik and Raid, Tiit and Flinkman, Juha and Feldman, Valeriy},
  journal={Population Ecology},
  volume={53},
  number={4},
  pages={511--523},
  year={2011},
  publisher={Springer}
}

@incollection{domenici2013effect,
  title={The {E}ffect of {H}ypoxia on {F}ish {S}wimming {P}erformance and {B}ehaviour},
  author={Domenici, P and Herbert, NA and Lefran{\c{c}}ois, C and Steffensen, JF and McKenzie, DJ},
  editor={Palstra, Arjan P. and Planas, Josep V.},
  booktitle={Swimming Physiology of Fish},
  publisher={Springer Berlin Heidelberg},
  address={Berlin, Heidelberg},
  pages={129--159},
  year={2013},
  publisher={Springer},
  doi={10.1007/978-3-642-31049-2_6}
}

@article{andersen2017long,
  title={Long-term temporal and spatial trends in eutrophication status of the {B}altic {S}ea},
  author={Andersen, Jesper H and Carstensen, Jacob and Conley, Daniel J and Dromph, Karsten and Fleming-Lehtinen, Vivi and Gustafsson, Bo G and Josefson, Alf B and Norkko, Alf and Villn{\"a}s, Anna and Murray, Ciar{\'a}n},
  journal={Biological Reviews},
  volume={92},
  number={1},
  pages={135--149},
  year={2017},
  publisher={Wiley Online Library},
  doi={10.1111/brv.12221}
}

@article{koster2005baltic,
  title={Baltic cod recruitment--the impact of climate variability on key processes},
  author={K{\"o}ster, Friedrich W and M{\"o}llmann, Christian and Hinrichsen, Hans-Harald and Wieland, Kai and Tomkiewicz, Jonna and Kraus, Gerd and Voss, R{\"u}diger and Makarchouk, Andrei and MacKenzie, Brian R and St. John, Michael A and Schnack, Dietrich and Rohlf, Norbert and Linkowski, Tomasz and Beyer, Jan E},
  journal={ICES Journal of Marine Science},
  volume={62},
  number={7},
  pages={1408--1425},
  year={2005},
  publisher={Oxford University Press},
  doi={10.1016/j.icesjms.2005.05.004}
}

@article{powers2005effects,
  title={Effects of eutrophication on bottom habitat and prey resources of demersal fishes},
  author={Powers, Sean P and Peterson, Charles H and Christian, Robert R and Sullivan, Elizabeth and Powers, Monica J and Bishop, Melanie J and Buzzelli, Christopher P},
  journal={Marine Ecology Progress Series},
  volume={302},
  pages={233--243},
  year={2005},
  doi={10.3354/meps302233}
}

@article{roberts2009effects,
  title={Effects of hypolimnetic hypoxia on foraging and distributions of {L}ake {E}rie yellow perch},
  author={Roberts, James J and H{\"o}{\"o}k, Tomas O and Ludsin, Stuart A and Pothoven, Steven A and Vanderploeg, Henry A and Brandt, Stephen B},
  journal={Journal of Experimental Marine Biology and Ecology},
  volume={381},
  pages={S132--S142},
  year={2009},
  note = {Ecological Impacts of Hypoxia on Living Resources},
  publisher={Elsevier},
  doi={10.1016/j.jembe.2009.07.017}
}

@article{bell2005species,
  title={Species-specific avoidance responses by blue crabs and fish to chronic and episodic hypoxia},
  author={Bell, GW and Eggleston, DB},
  journal={Marine Biology},
  volume={146},
  number={4},
  pages={761--770},
  year={2005},
  publisher={Springer},
  doi={10.1007/s00227-004-1483-7}
}

@article{kraus2015dynamic,
  title={Dynamic hypoxic zones in {L}ake {E}rie compress fish habitat, altering vulnerability to fishing gears},
  author={Kraus, Richard T and Knight, Carey T and Farmer, Troy M and Gorman, Ann Marie and Collingsworth, Paris D and Warren, Glenn J and Kocovsky, Patrick M and Conroy, Joseph D},
  journal={Canadian Journal of Fisheries and Aquatic Sciences},
  volume={72},
  number={6},
  pages={797--806},
  year={2015},
  publisher={NRC Research Press},
  doi={10.1139/cjfas-2014-0517}
}

@article{chamberlin2020hypoxia,
  title={Hypoxia augments edge effects of water column stratification on fish distribution},
  author={Chamberlin, Derek W and Knight, Carey T and Kraus, Richard T and Gorman, Ann Marie and Xu, Wenzhao and Collingsworth, Paris D},
  journal={Fisheries Research},
  volume={231},
  pages={105684},
  year={2020},
  publisher={Elsevier},
  doi={10.1016/j.fishres.2020.105684}
}

@article{pihl1992hypoxia,
  title={Hypoxia-induced structural changes in the diet of bottom-feeding fish and {C}rustacea},
  author={Pihl, Leif and Baden, Susanne P and Diaz, Robert J and Schaffner, Linda C},
  journal={Marine Biology},
  volume={112},
  number={3},
  pages={349--361},
  year={1992},
  publisher={Springer},
  doi={10.1007/BF00356279}
}

@article{wang2016hypoxia,
  title={Hypoxia causes transgenerational impairments in reproduction of fish},
  author={Wang, Simon Yuan and Lau, Karen and Lai, Keng-Po and Zhang, Jiang-Wen and Tse, Anna Chung-Kwan and Li, Jing-Woei and Tong, Yin and Chan, Ting-Fung and Wong, Chris Kong-Chu and Chiu, Jill Man-Ying and , Wai-Ting Au, Doris and Sze-Tsai Wong, Alice and Yuen-Chong Kong, Richard  and Shiu-Sun Wu, Rudolf},
  journal={Nature communications},
  volume={7},
  number={12114},
  pages={1--9},
  year={2016},
  publisher={Nature Publishing Group},
  doi={10.1038/ncomms12114}
}

@article{stramma2012expansion,
  title={Expansion of oxygen minimum zones may reduce available habitat for tropical pelagic fishes},
  author={Stramma, Lothar and Prince, Eric D and Schmidtko, Sunke and Luo, Jiangang and Hoolihan, John P and Visbeck, Martin and Wallace, Douglas WR and Brandt, Peter and K{\"o}rtzinger, Arne},
  journal={Nature Climate Change},
  volume={2},
  pages={33--37},
  year={2012},
  publisher={Nature Publishing Group},
  doi={10.1038/nclimate1304}
}

@article{karlson2020linking,
  title={Linking consumer physiological status to food-web structure and prey food value in the {B}altic {S}ea},
  author={Karlson, Agnes ML and Gorokhova, Elena and G{\aa}rdmark, Anna and Pekcan-Hekim, Zeynep and Casini, Michele and Albertsson, Jan and Sundelin, Brita and Karlsson, Olle and Bergstr{\"o}m, Lena},
  journal={Ambio},
  volume={49},
  number={2},
  pages={391--406},
  year={2020},
  publisher={Springer},
  doi={10.1007/s13280-019-01201-1}
}

@article{pihl1994changes,
  title={Changes in the {D}iet of {D}emersal {F}ish due to {E}utrophication-{I}nduced {H}ypoxia in the {K}attegat, {S}weden},
  author={Pihl, Leif},
  journal={Canadian Journal of Fisheries and Aquatic Sciences},
  volume={51},
  number={2},
  pages={321--336},
  year={1994},
  publisher={NRC Research Press Ottawa, Canada},
  doi={10.1139/f94-033}
}

@article{griffiths2017importance,
  title={The importance of benthic--pelagic coupling for marine ecosystem functioning in a changing world},
  author={Griffiths, Jennifer R and Kadin, Martina and Nascimento, Francisco JA and Tamelander, Tobias and T{\"o}rnroos, Anna and Bonaglia, Stefano and Bonsdorff, Erik and Br{\"u}chert, Volker and G{\aa}rdmark, Anna and J{\"a}rnstr{\"o}m, Marie and Jonne Kotta, MARTIN Lindegren, Marie C. Nordstr{\"o}m and Norkko, Alf and Olsson, Jens and Weigel, Benjamin and \v{Z}ydelis, Ramunas and Blenckner, Thorsten and Niiranen, Susa and Winder, Monika},
  journal={Global change biology},
  volume={23},
  number={6},
  pages={2179--2196},
  year={2017},
  publisher={Wiley Online Library},
  doi={10.1111/gcb.13642}
}

@article{mohan2016out,
  title={Out of breath and hungry: {n}atural tags reveal trophic resilience of {A}tlantic croaker to hypoxia exposure},
  author={Mohan, John and Walther, Benjamin},
  journal={Marine Ecology Progress Series},
  volume={560},
  pages={207--221},
  year={2016},
  doi={10.3354/meps11934}
}

@article{steube2021distributive,
  title={Distributive stress: {i}ndividually variable responses to hypoxia expand trophic niches in fish},
  author={Steube, Tyler R and Altenritter, Matthew E and Walther, Benjamin D},
  journal={Ecology},
  volume={102},
  number={6},
  pages={e03356},
  year={2021},
  publisher={Wiley Online Library},
  doi={10.1002/ecy.3356}
}

@article{roman2019interactive,
  title={Interactive {E}ffects of {H}ypoxia and {T}emperature on {C}oastal {P}elagic {Z}ooplankton and {F}ish},
  author={Roman, Michael R and Brandt, Stephen B and Houde, Edward D and Pierson, James J},
  journal={Frontiers in Marine Science},
  volume={6},
  pages={139},
  year={2019},
  publisher={Frontiers Media SA},
  doi={10.3389/fmars.2019.00139}
}

@article{mcbryan2013responses,
  title={Responses to {T}emperature and {H}ypoxia as {I}nteracting {S}tressors in {F}ish: implications for {A}daptation to {E}nvironmental {C}hange},
  author={McBryan, TL and Anttila, K and Healy, TM and Schulte, PM},
  journal={Integrative and Comparative Biology},
  volume={53},
  number={4},
  pages={648--659},
  year={2013},
  doi={10.1093/icb/ict066},
  publisher={Oxford University Press},
  doi={10.1093/icb/ict066}
}

@article{clark1999effects,
  title={Effects of an erroneous natural mortality rate on a simple age-structured stock assessment},
  author={Clark, William G},
  journal={Canadian Journal of Fisheries and Aquatic Sciences},
  volume={56},
  number={10},
  pages={1721--1731},
  year={1999},
  publisher={NRC Research Press Ottawa, Canada},
  doi={10.1139/f99-085}
}

@article{reum2020ensemble,
  title={Ensemble {P}rojections of {F}uture {C}limate {C}hange {I}mpacts on the {E}astern {B}ering {S}ea {F}ood {W}eb {U}sing a {M}ultispecies {S}ize {S}pectrum {M}odel},
  author={Reum, Jonathan CP and Blanchard, Julia L and Holsman, Kirstin K and Aydin, Kerim and Hollowed, Anne B and Hermann, Albert J and Cheng, Wei and Faig, Amanda and Haynie, Alan C and Punt, Andr{\'e} E},
  journal={Frontiers in Marine Science},
  volume={7},
  number={124},
  year={2020},
  publisher={Frontiers Media SA},
  doi={10.3389/fmars.2020.00124}
}

@article{cooke2014out,
  title={Out-of-{S}ample {V}alidation for {S}tructured {E}xpert {J}udgment of {A}sian {C}arp {E}stablishment in {L}ake {E}rie},
  author={Cooke, Roger M and Wittmann, Marion E and Lodge, David M and Rothlisberger, John D and Rutherford, Edward S and Zhang, Hongyan and Mason, Doran M},
  journal={Integrated Environmental Assessment and Management},
  volume={10},
  number={4},
  pages={522--528},
  year={2014},
  publisher={Wiley Online Library},
  doi={10.1002/ieam.1559}
}

@article{ekstrom2021cardiorespiratory,
  title={Cardiorespiratory adjustments to chronic environmental warming improve hypoxia tolerance in {E}uropean perch (\textit{{P}erca fluviatilis})},
  author={Ekstr{\"o}m, Andreas and Sundell, Erika and Morgenroth, Daniel and McArley, Tristan and G{\aa}rdmark, Anna and Huss, Magnus and Sandblom, Erik},
  journal={Journal of Experimental Biology},
  volume={224},
  number={6},
  note={jeb241554},
  year={2021},
  publisher={The Company of Biologists Ltd},
  doi={10.1242/jeb.241554}
}

@article{behrens2012effects,
  title={Effects of {H}ypoxic {E}xposure {D}uring {F}eeding on {SDA} and {P}ostprandial {C}ardiovascular {P}hysiology in the {A}tlantic cod, \textit{{G}adus morhua}},
  author={Behrens, Jane W and Axelsson, Michael and Neuenfeldt, Stefan and Seth, Henrik},
  year={2012},
  journal={PLoS ONE},
  volume={7},
  number={9},
  pages={e46227},
  publisher={Public Library of Science San Francisco, USA},
  doi={10.1371/journal.pone.0046227}
}

@misc{datrasdata2020,
  title={ICES Database of Trawl Surveys (DATRAS)},
  year={2020},
  note={Extracted 9 November 2020 from Baltic International Trawl Survey (BITS)},
  author={{ICES}},
  address={Copenhagen},
  howpublished={https://www.ices.dk/data/data-portals/Pages/DATRAS.aspx}
}

@misc{icesgis2020,
  title={ICES Spatial Facility},
  year={2020},
  note={Extracted 13 July 2020},
  author={{ICES}}, 
  address={Copenhagen},
  howpublished={https://gis.ices.dk/sf/index.html?widget=StatRec#}
}

@article{mccrackin2018,
  author = {McCrackin, Michelle L. and Muller-Karulis, Bärbel and Gustafsson, Bo G. and Howarth, Robert W. and Humborg, Christoph and Svanbäck, Annika and Swaney, Dennis P.},
  title = {A Century of Legacy Phosphorus Dynamics in a Large Drainage Basin},
  journal = {Global Biogeochemical Cycles},
  volume = {32},
  number = {7},
  pages = {1107-1122},
  doi = {10.1029/2018GB005914},
  year = {2018}
}
