## Supplementary figures and images for "Declining food availability and habitat shifts drive community responses to marine hypoxia"

### all_agelength.jpeg

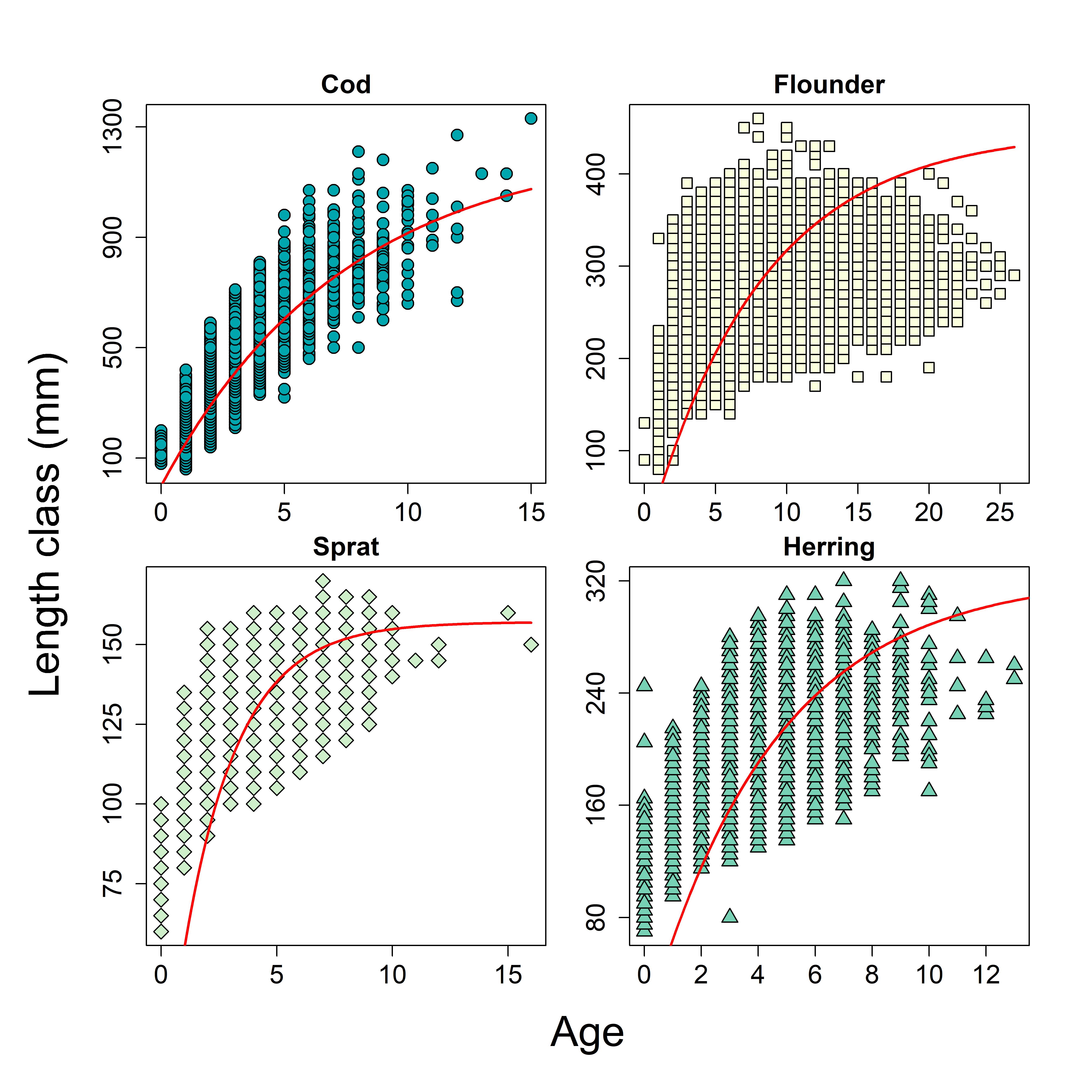

### all_lengthweight.jpeg

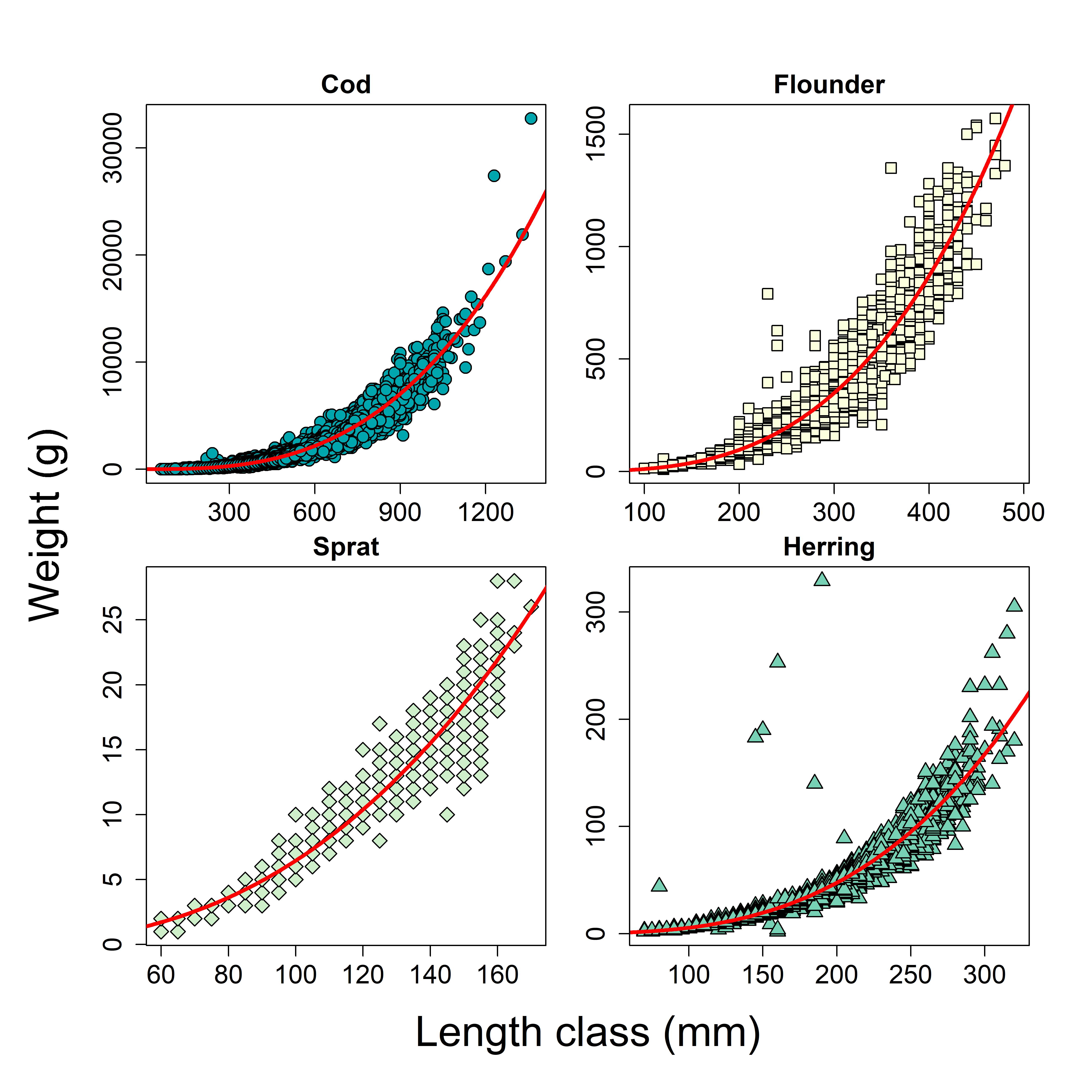

### all_pmrn.jpeg

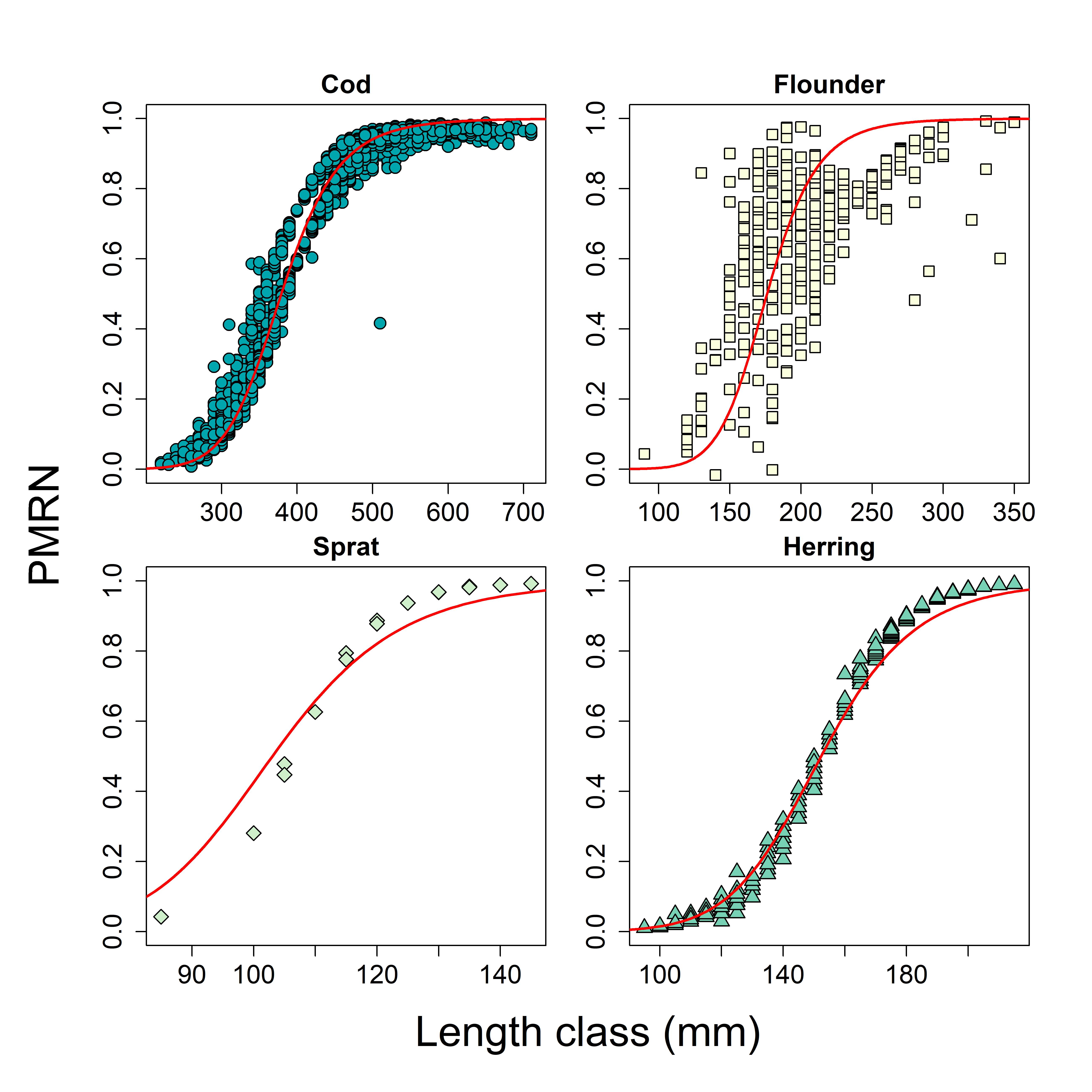

### all_ppmr.jpeg

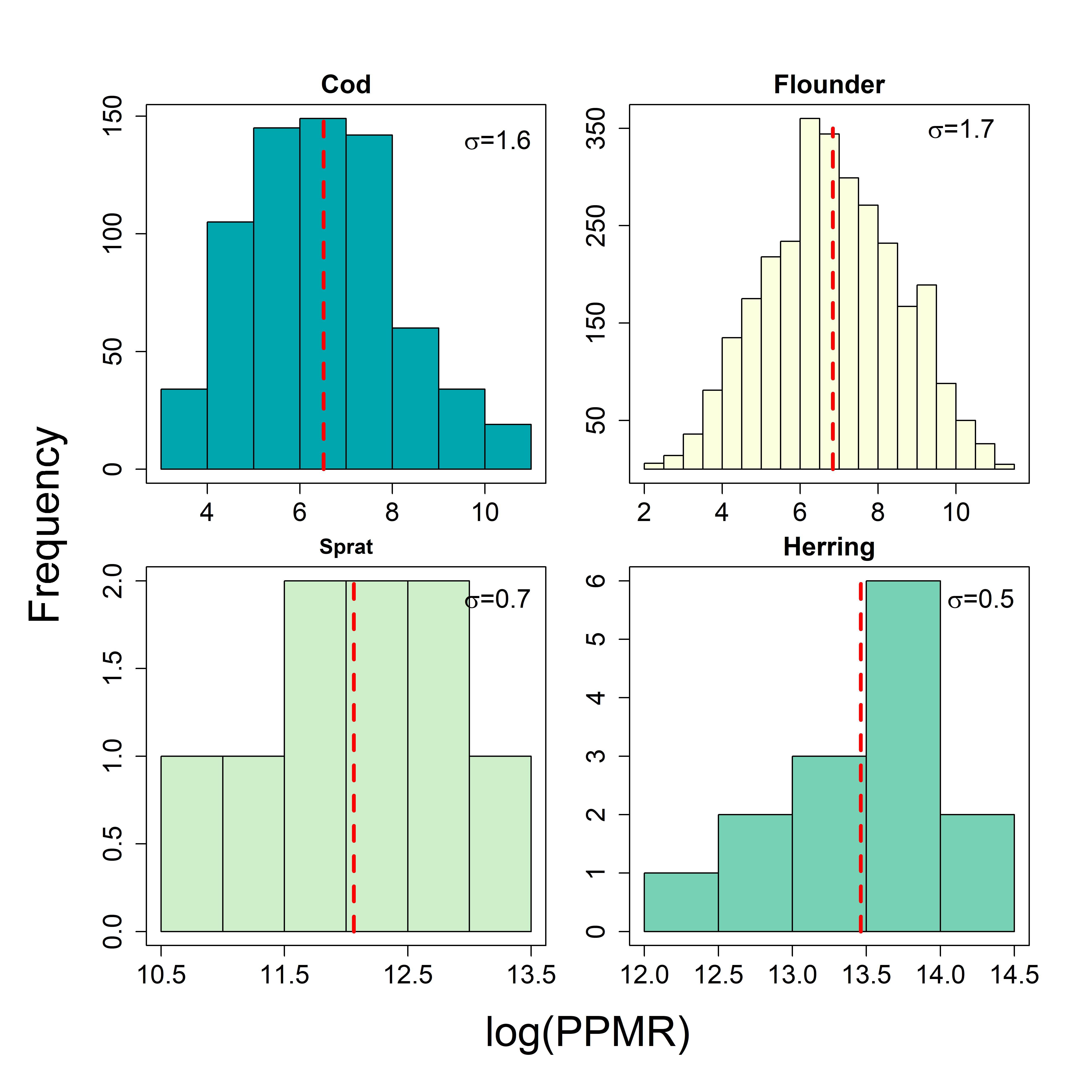

### all_scaling.jpg

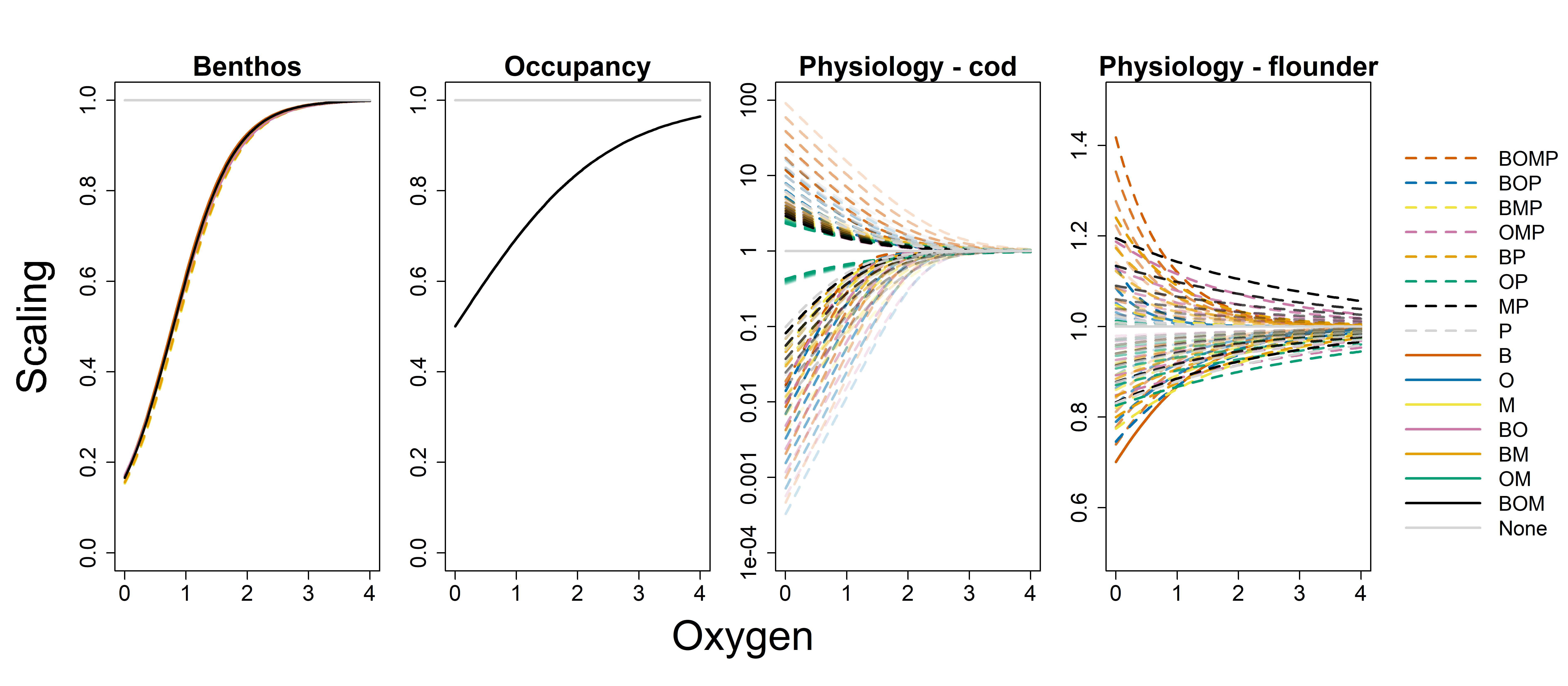

### all_selectivity.jpeg

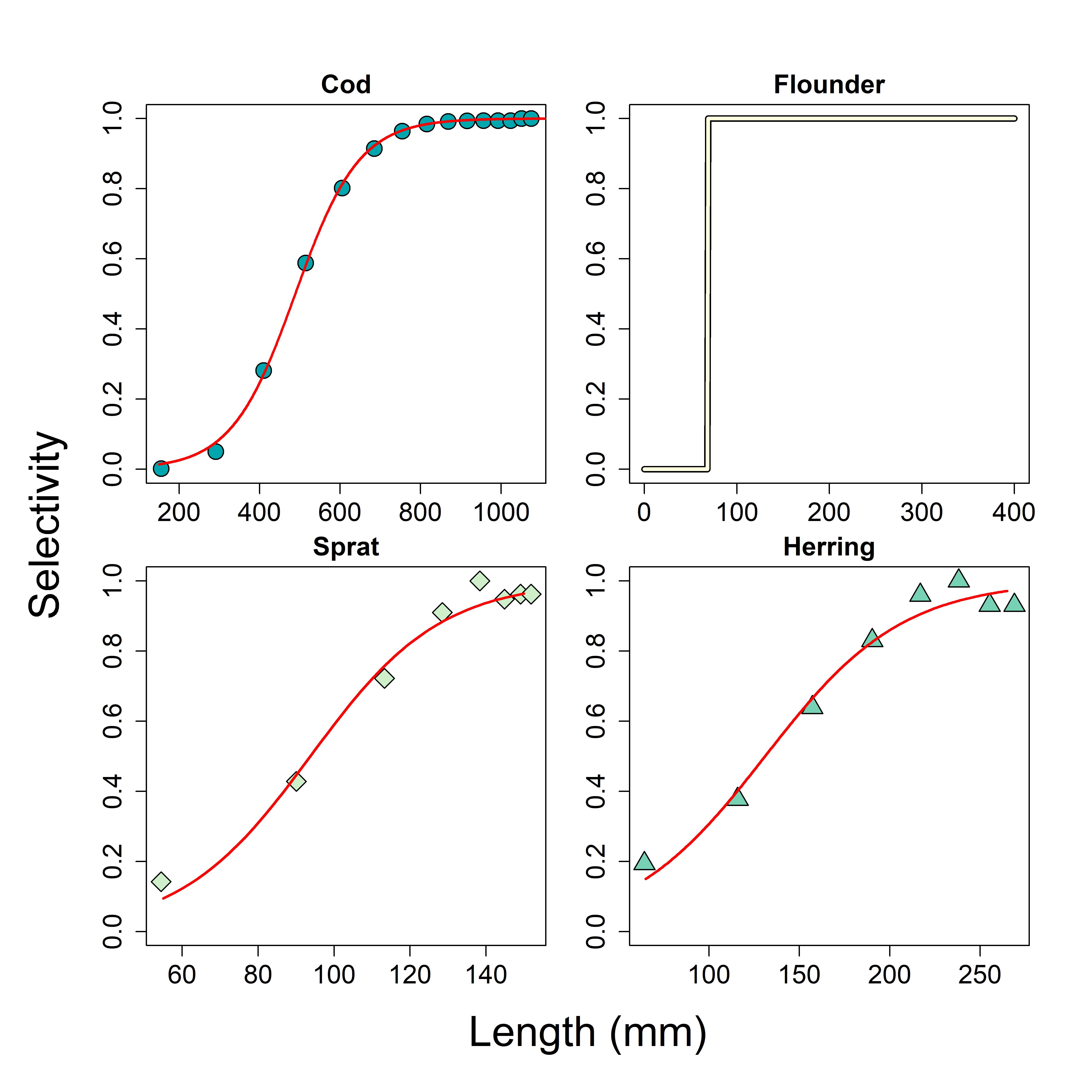

### cal_growth.jpg

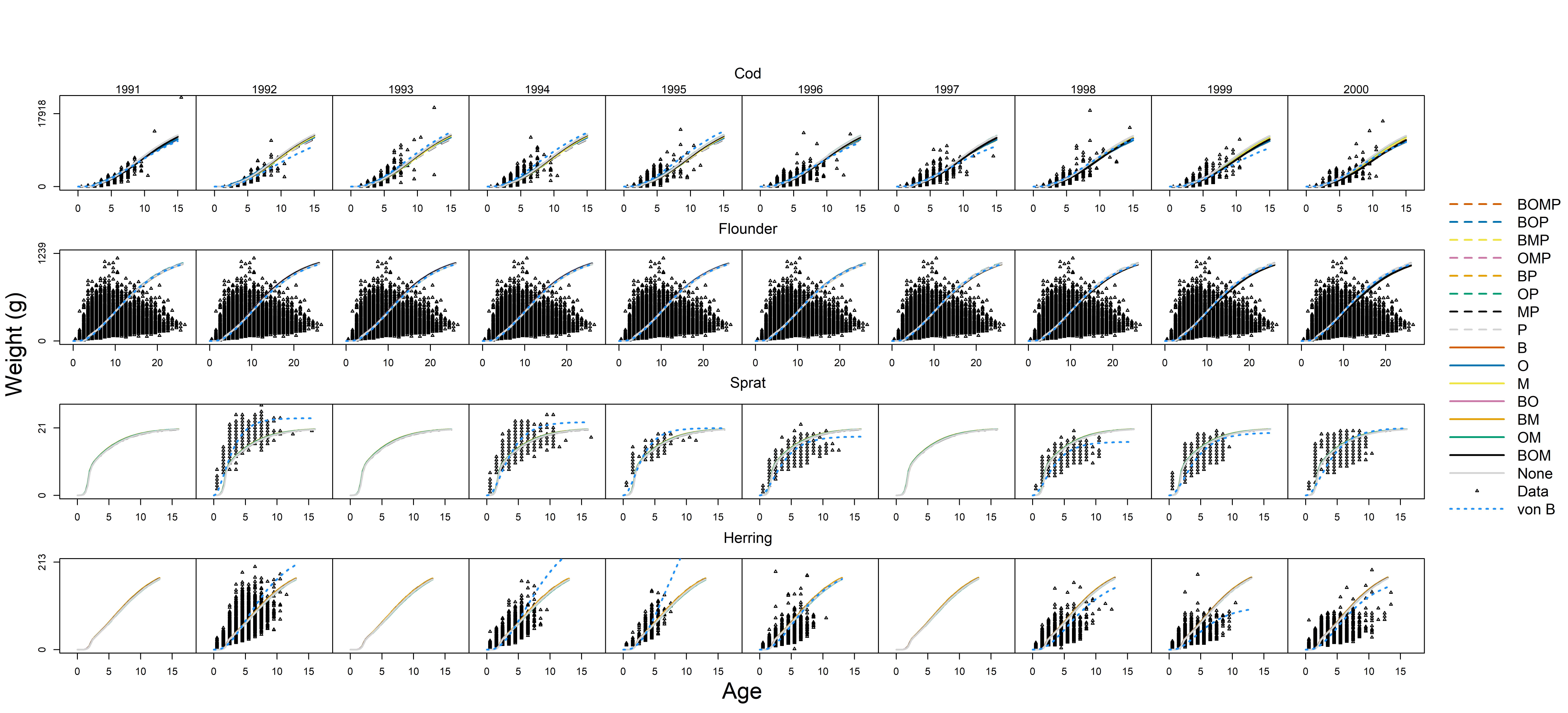

### calibration_error.jpg

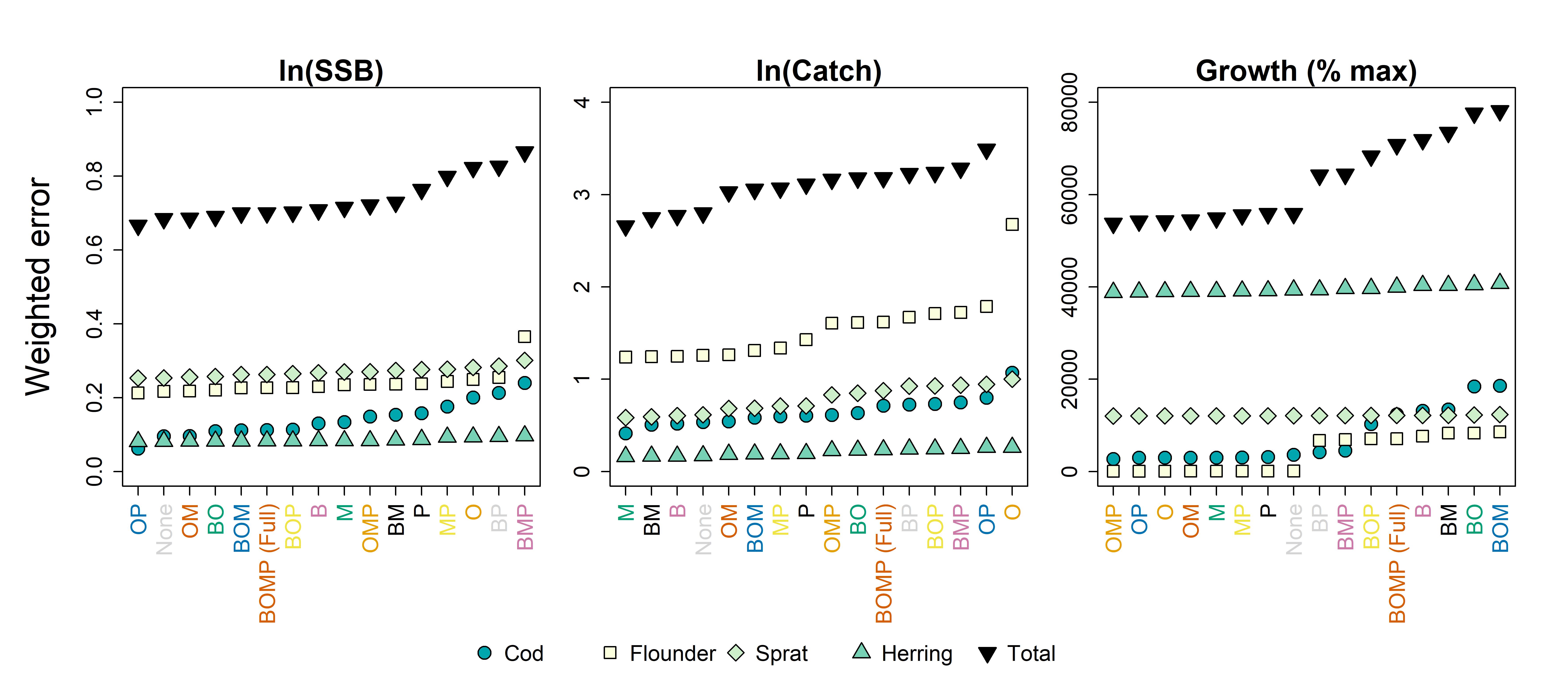

### calibration_error_rank.jpg

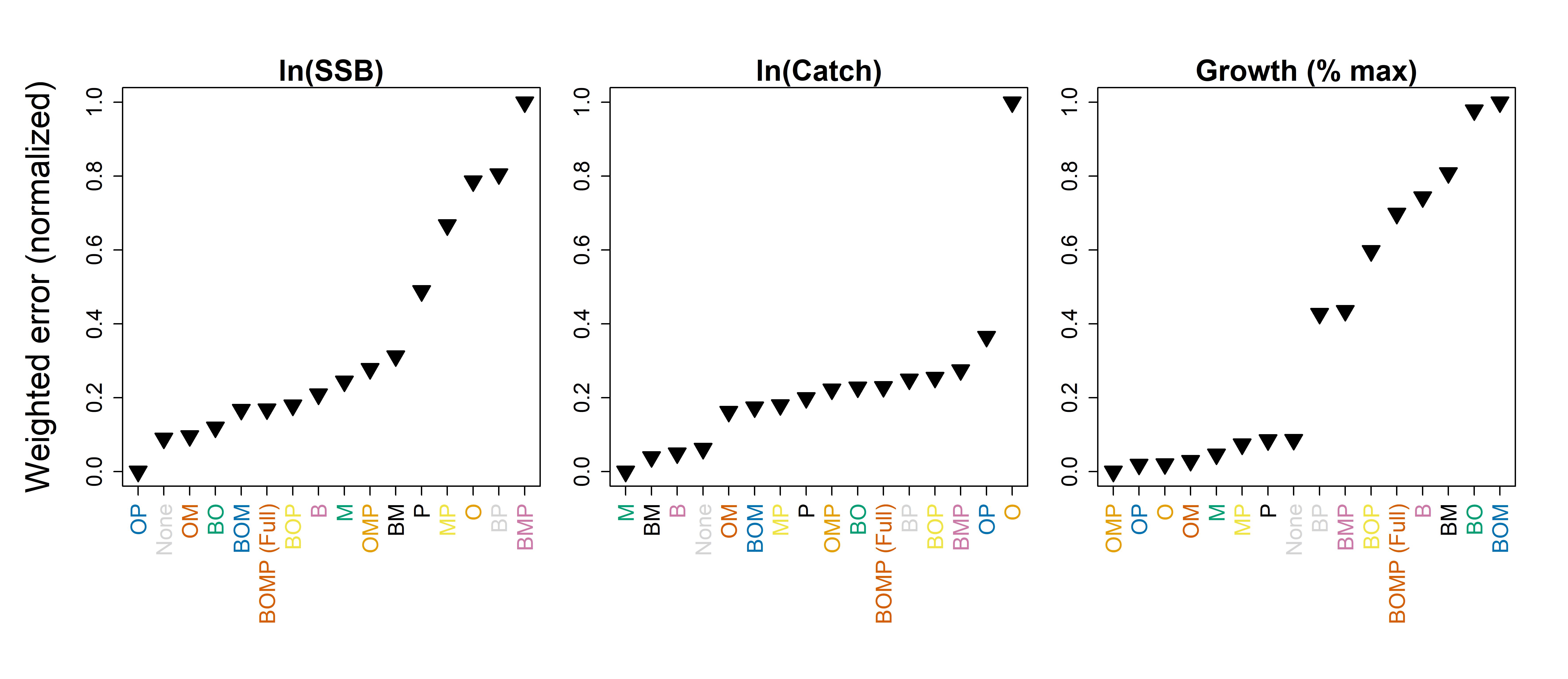

### cod_growth_eight.jpg

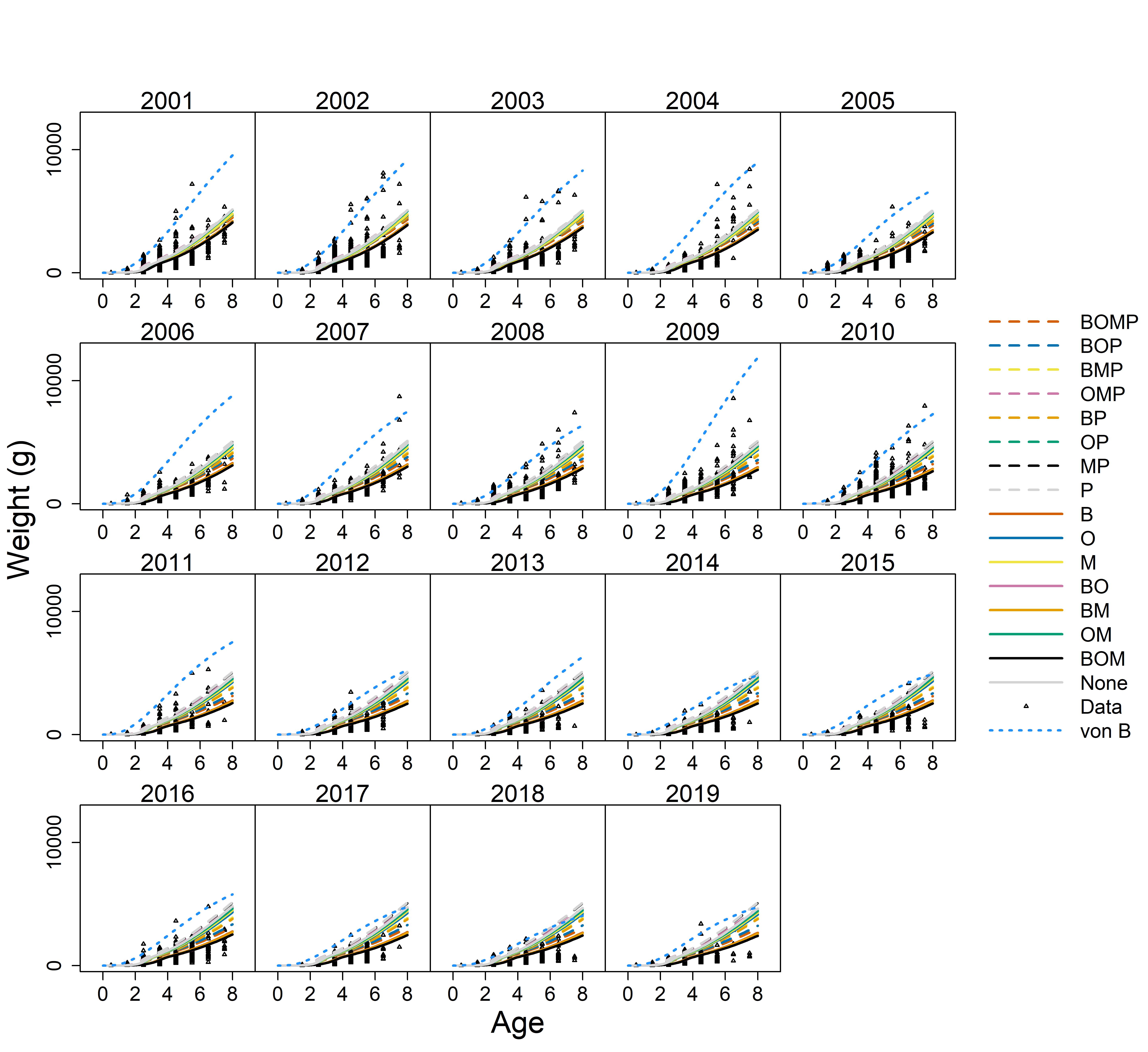

### cod_growth_eight_best.jpg

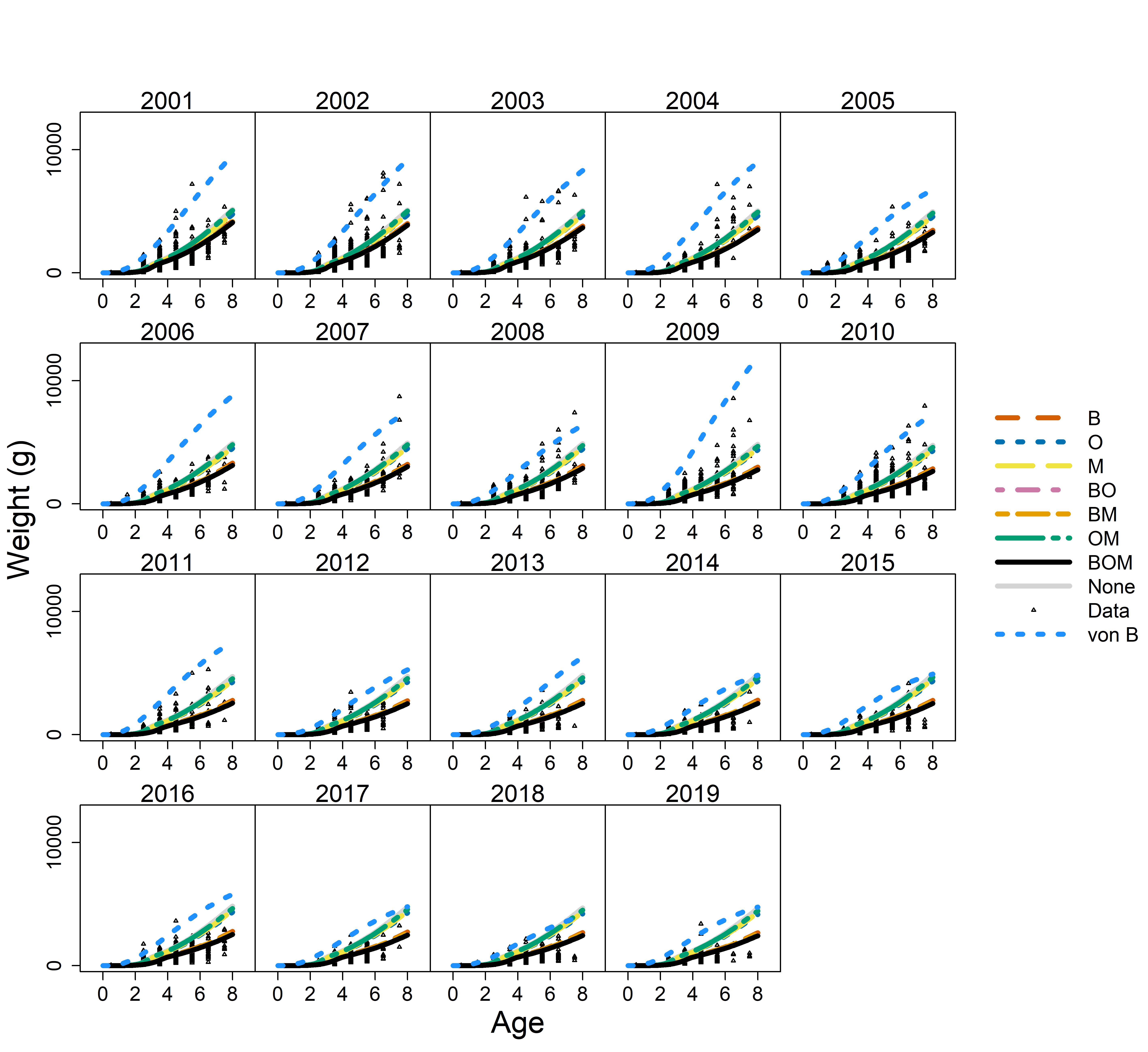

### conceptual.jpg

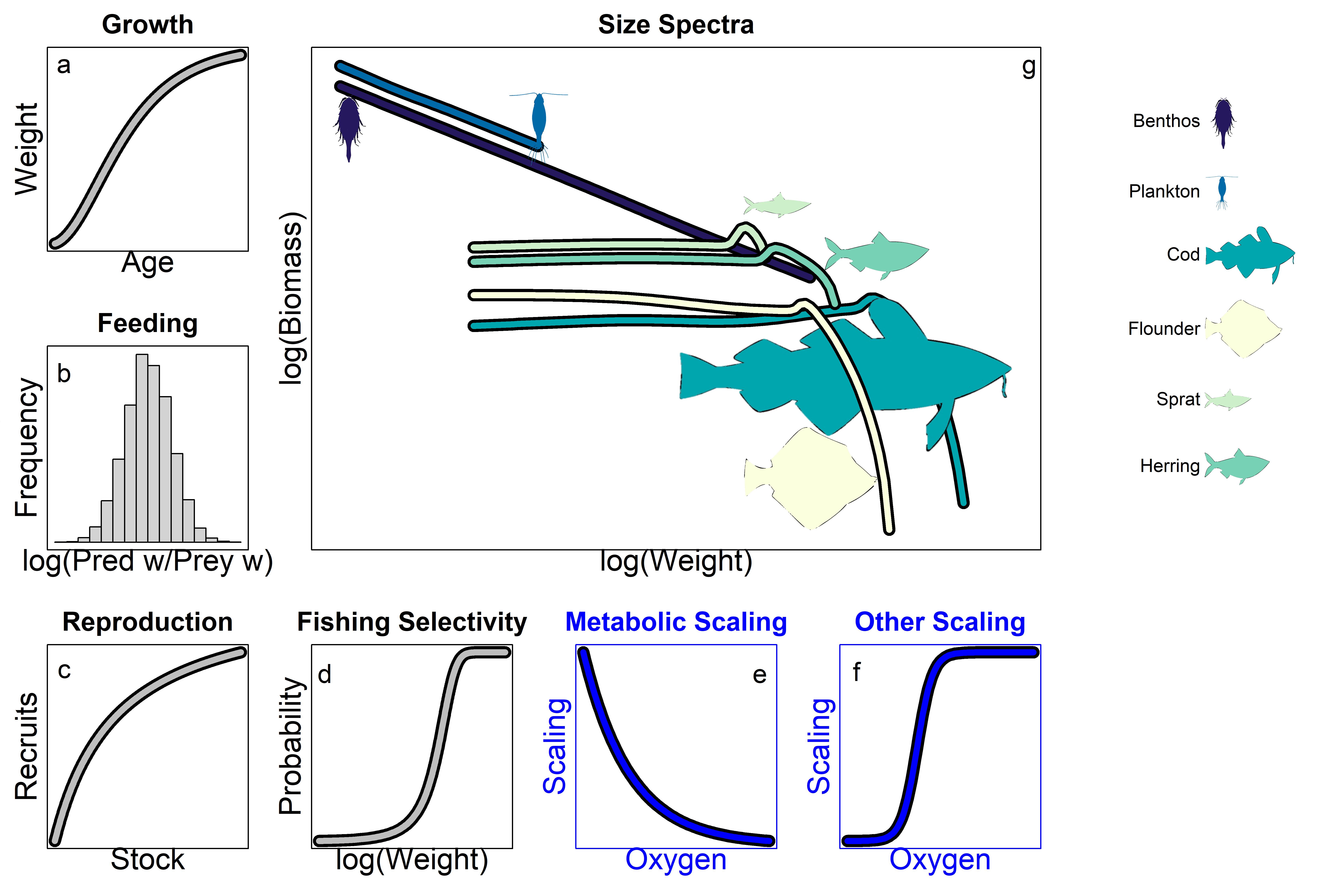

### diet_scenarios.jpg

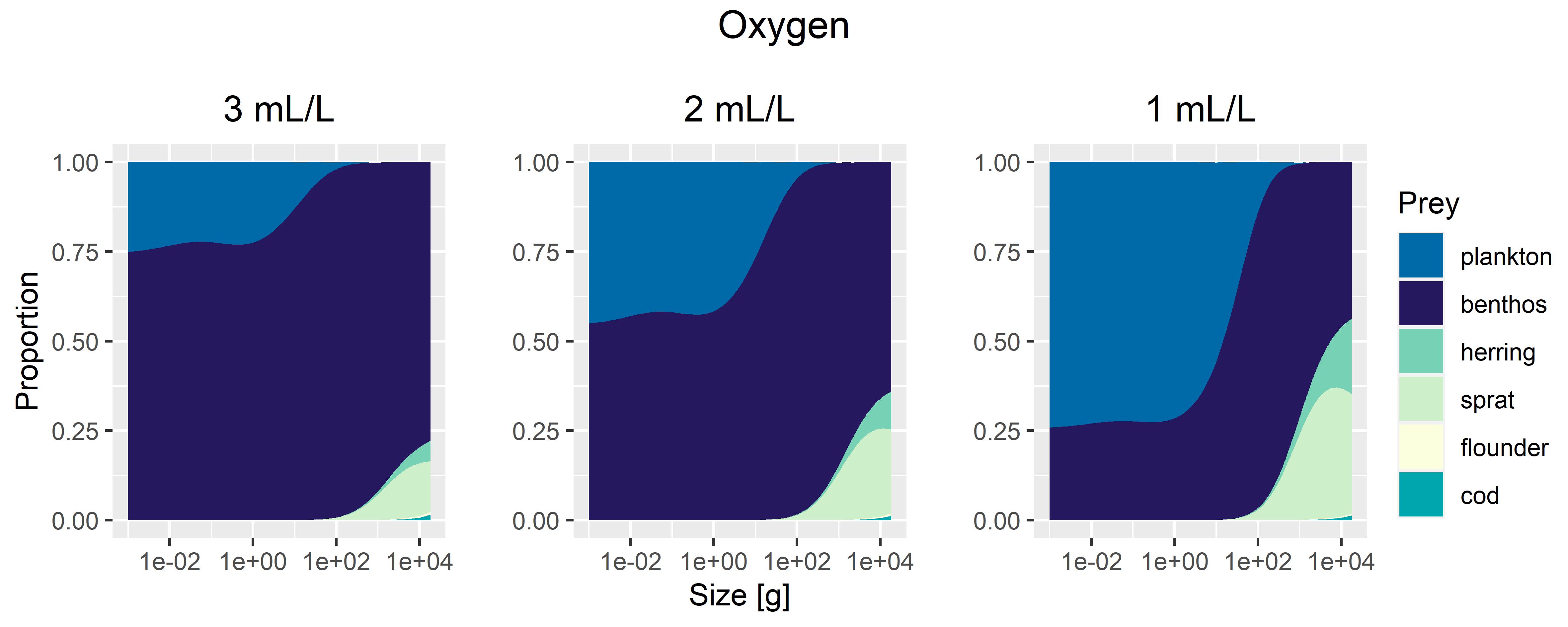

### final_error_cal.jpg

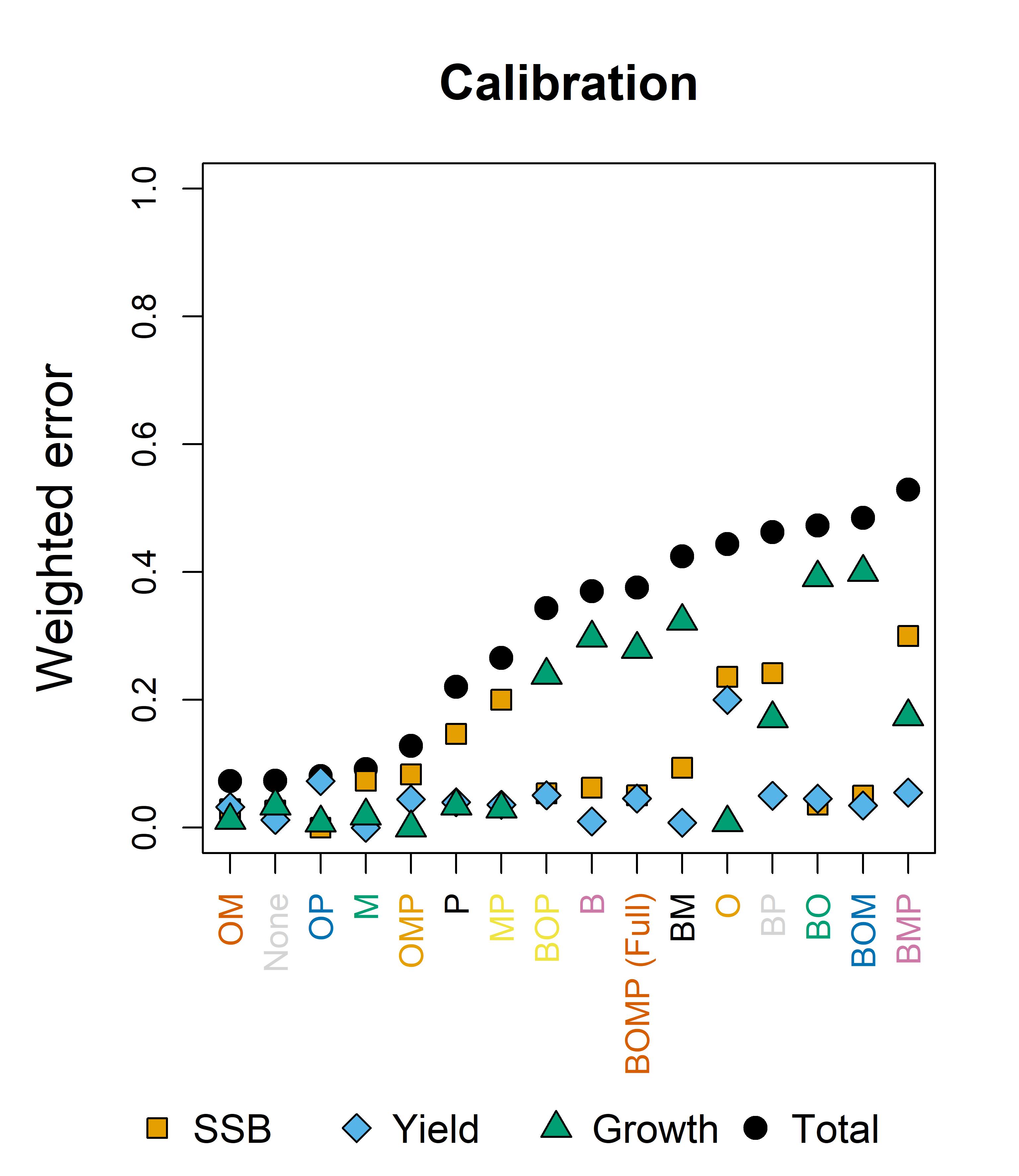

### final_error_sim.jpg

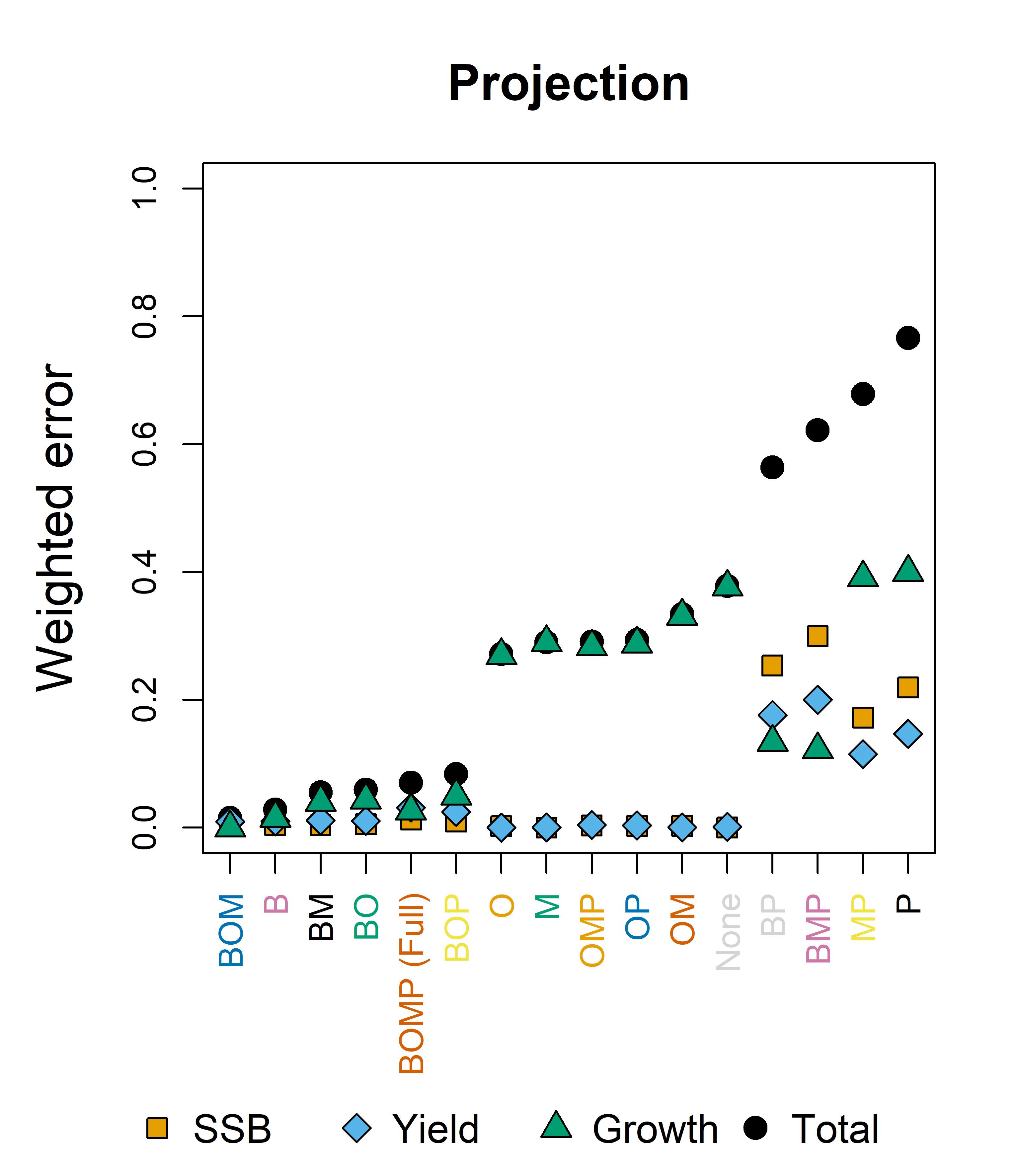

### flounder_obs_diet.jpg

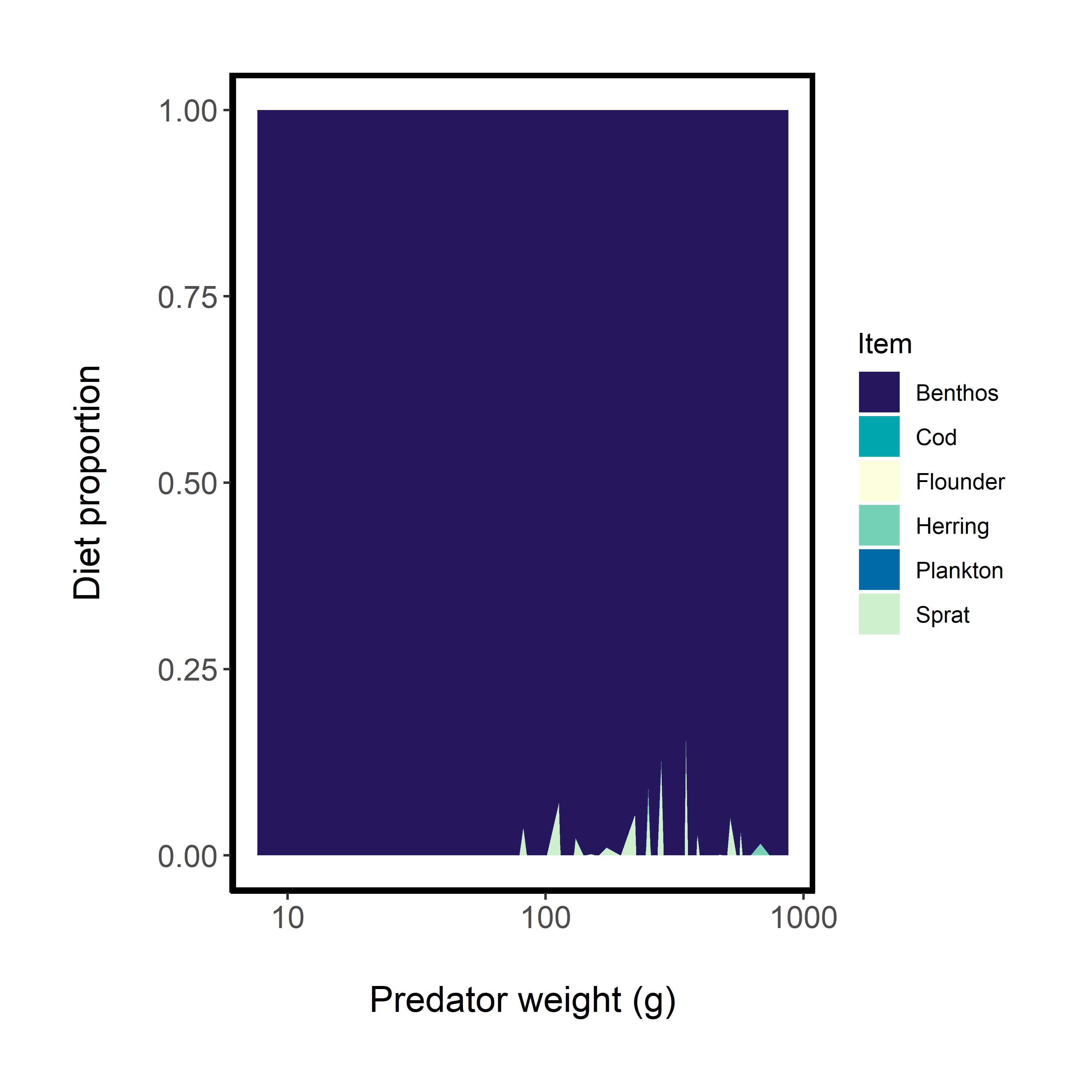

### foodweb_scenarios.jpg

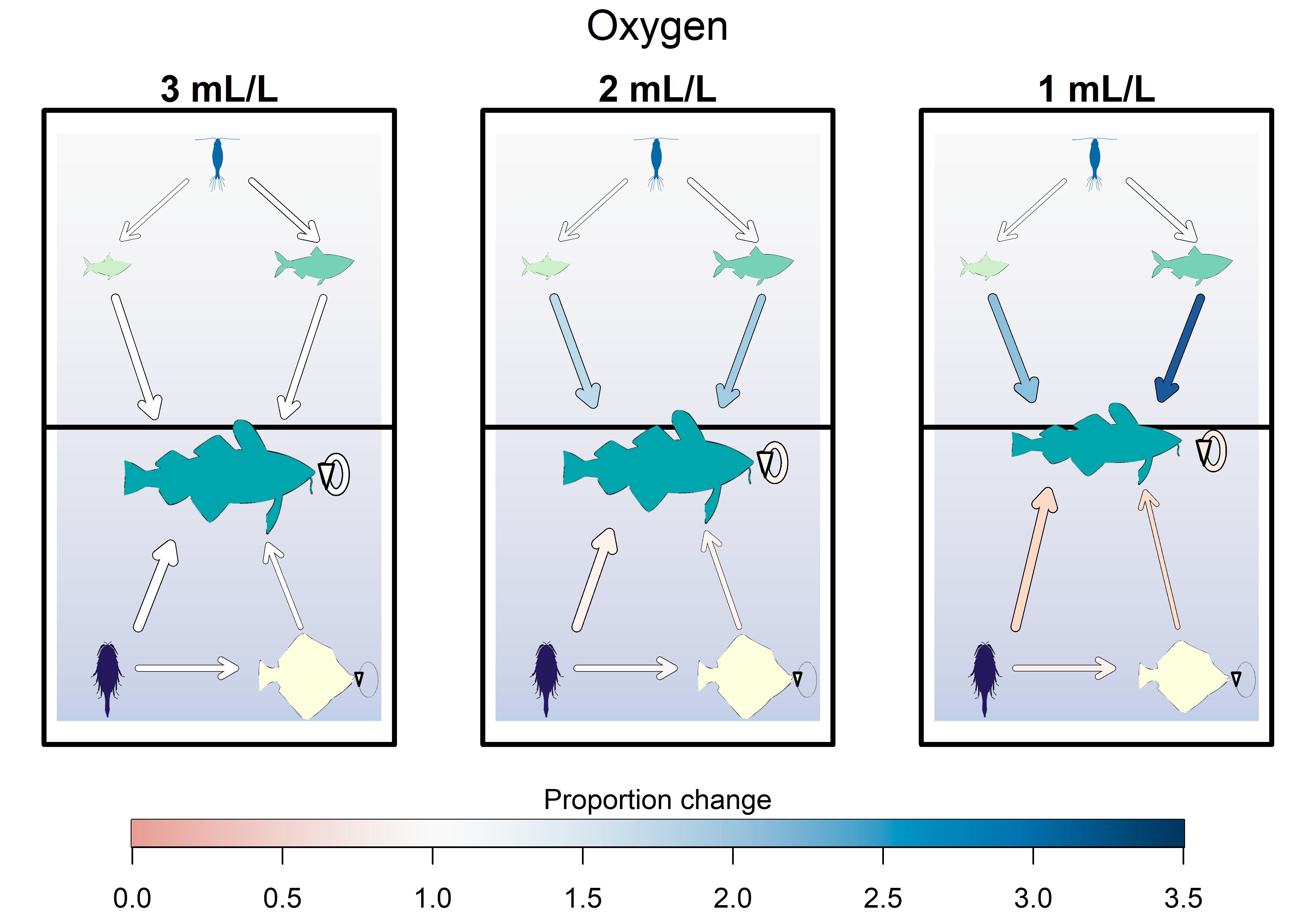

### occupancy.jpg

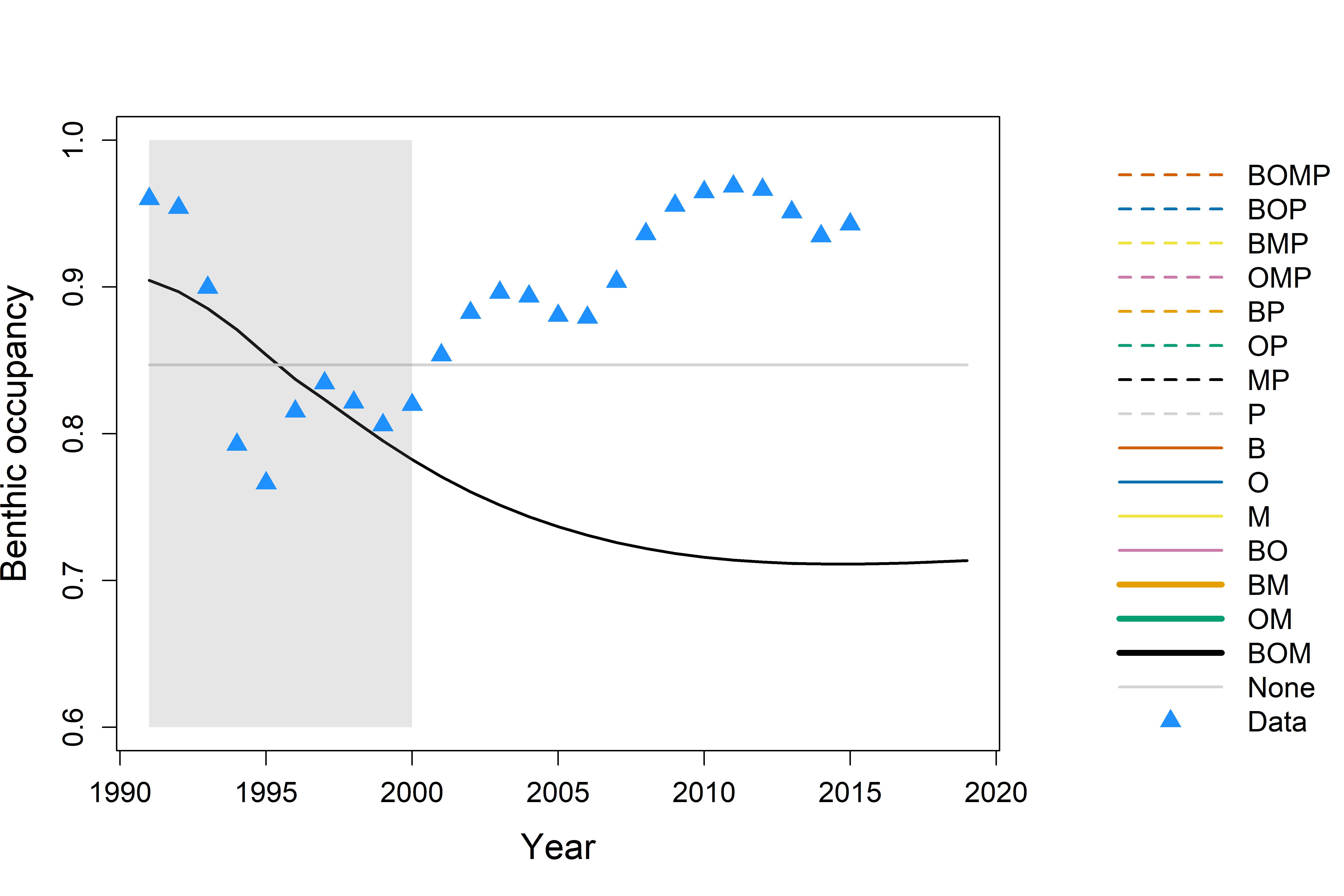

### oxygen.jpeg

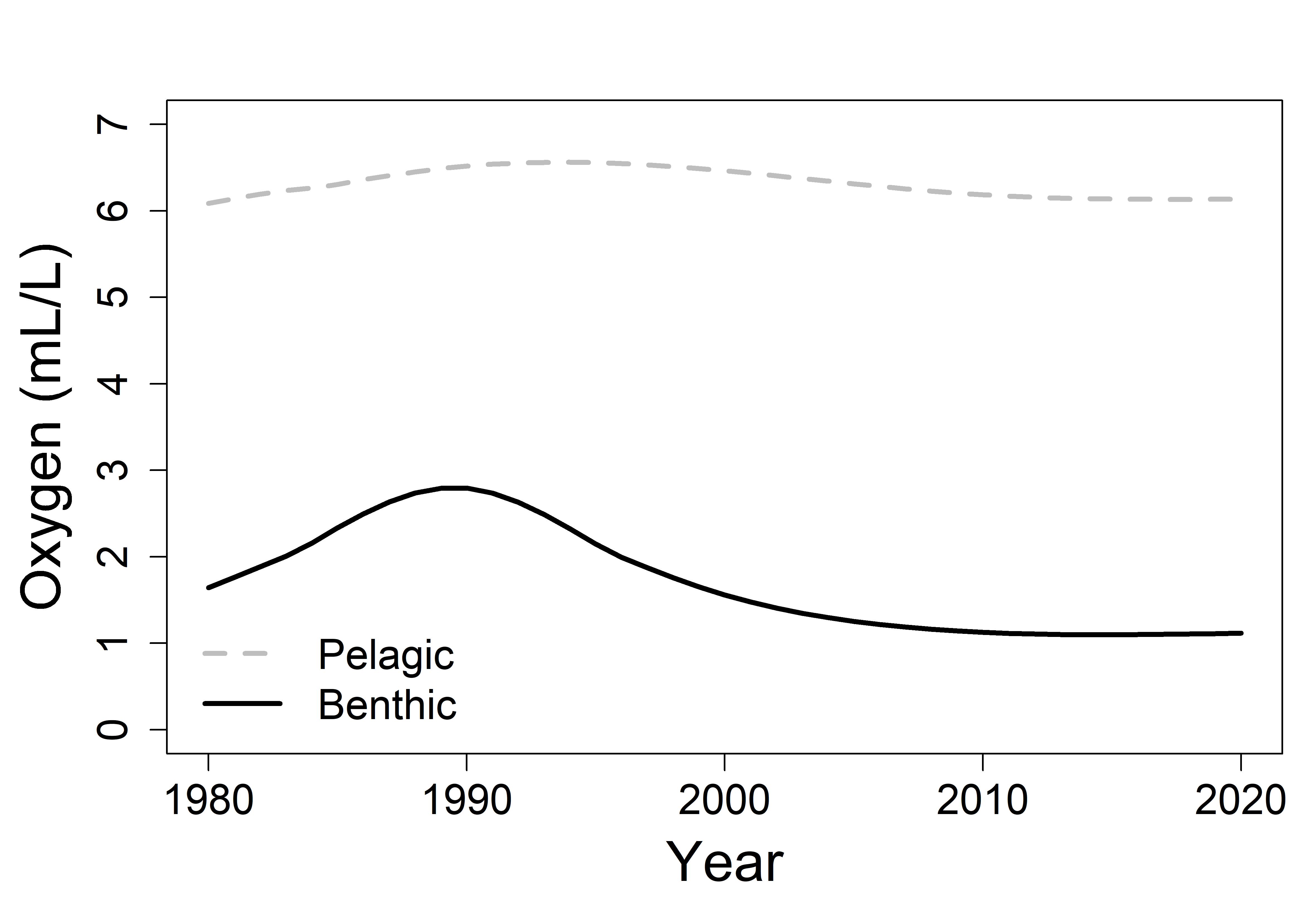

### oxygen.jpg

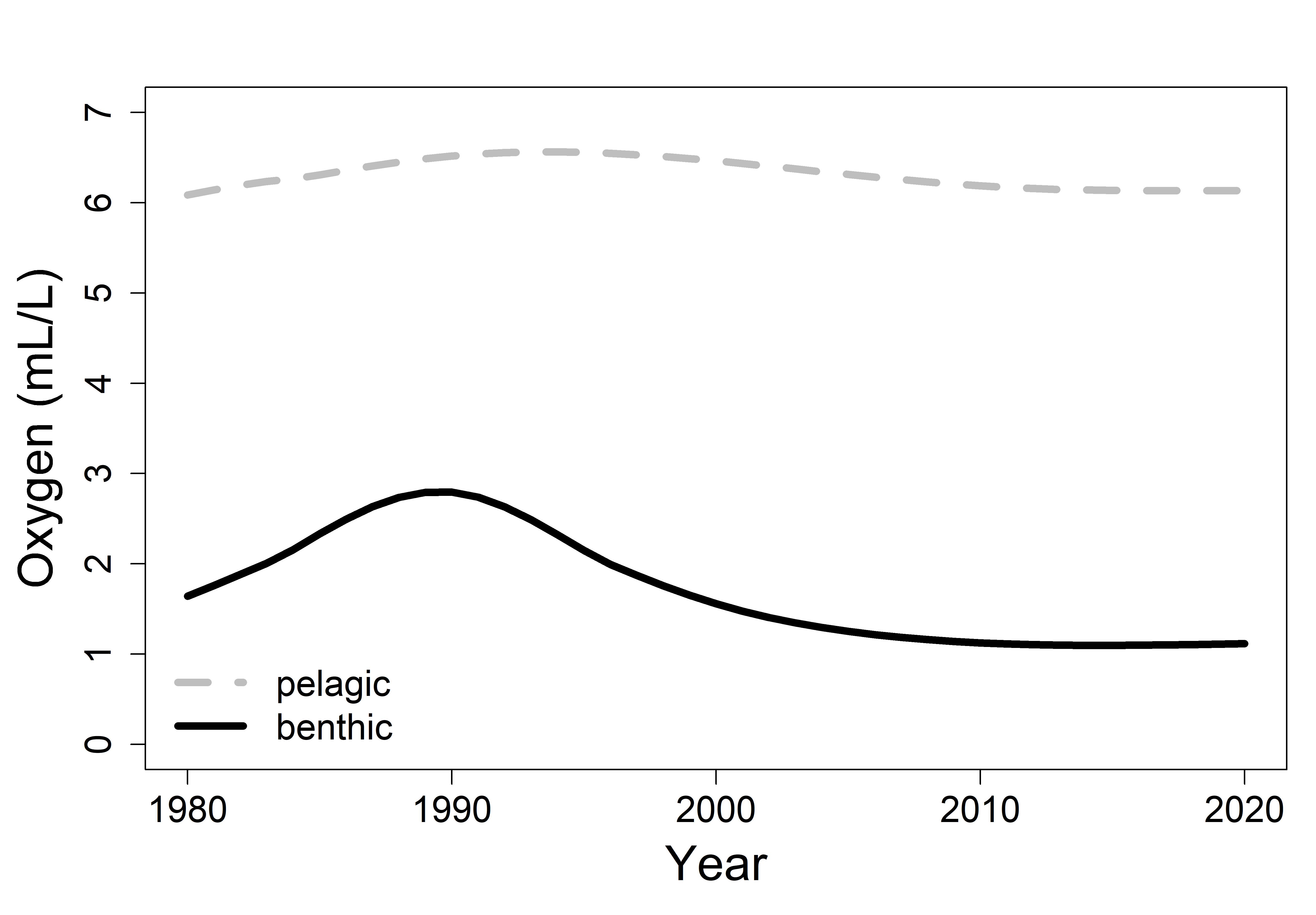

### pcrit.jpeg

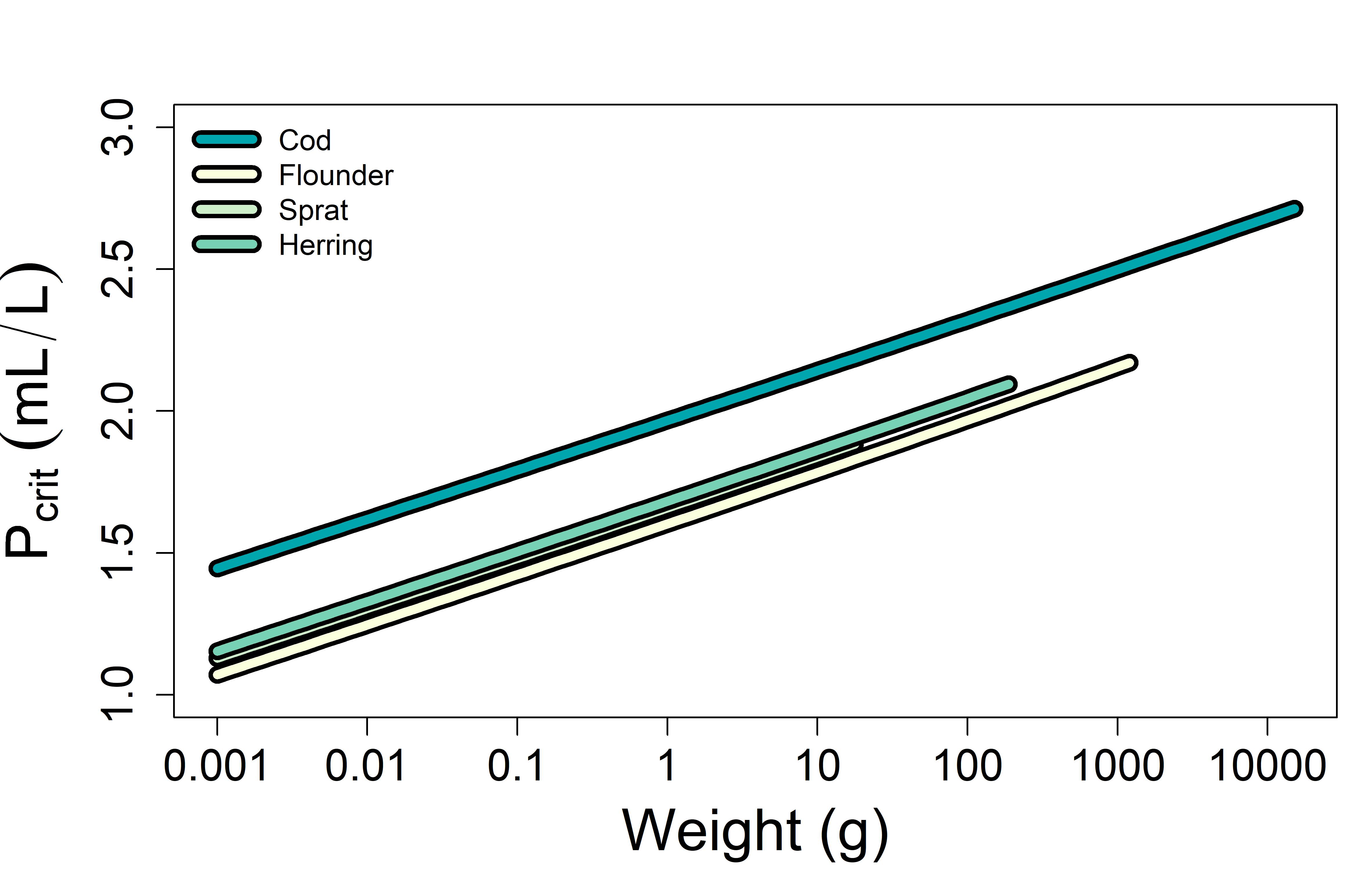

### sens_error_cal.jpg

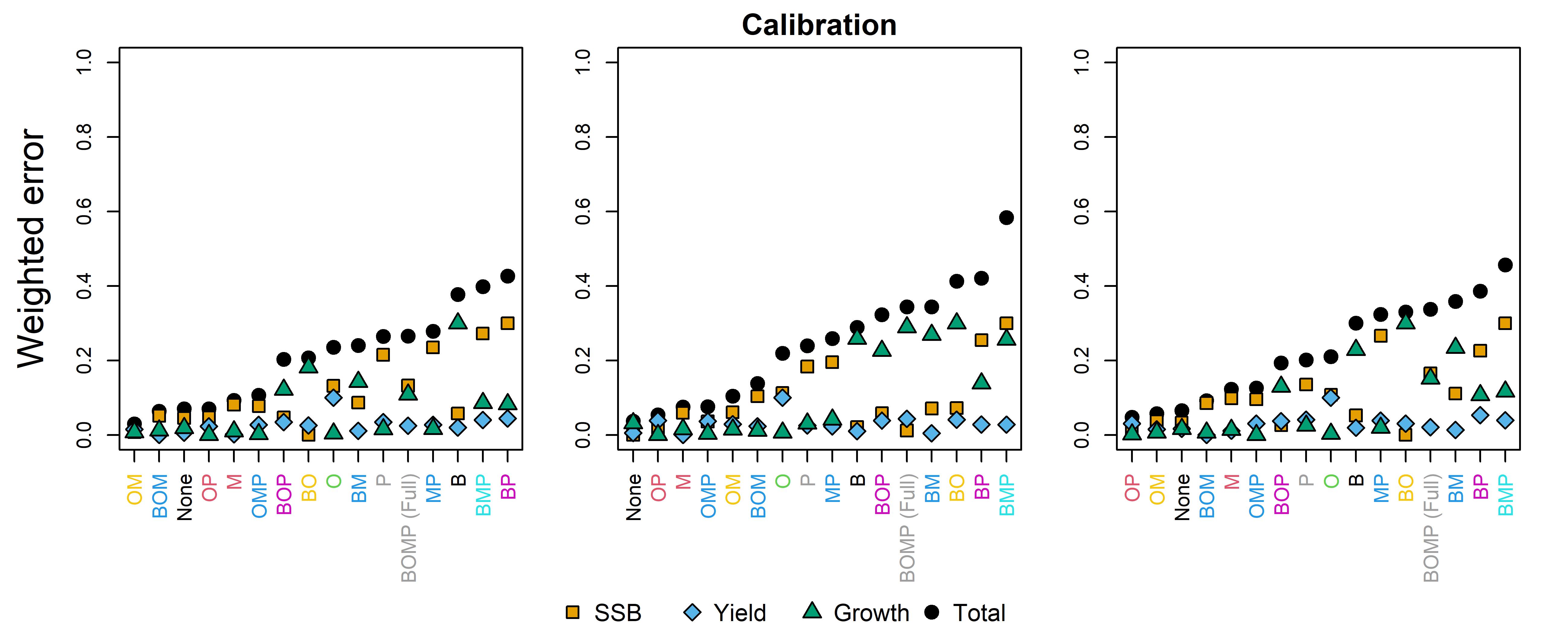

### sens_error_sim.jpg

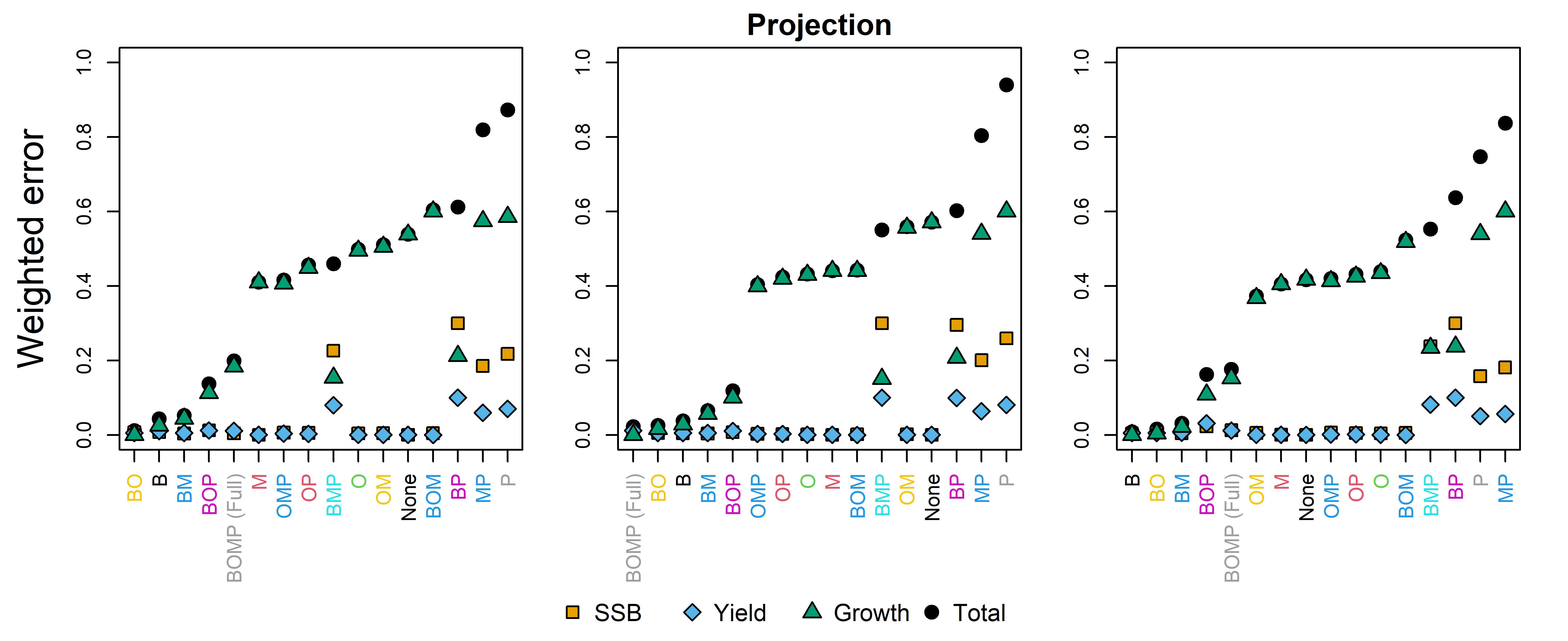

### sim_biomass.jpg

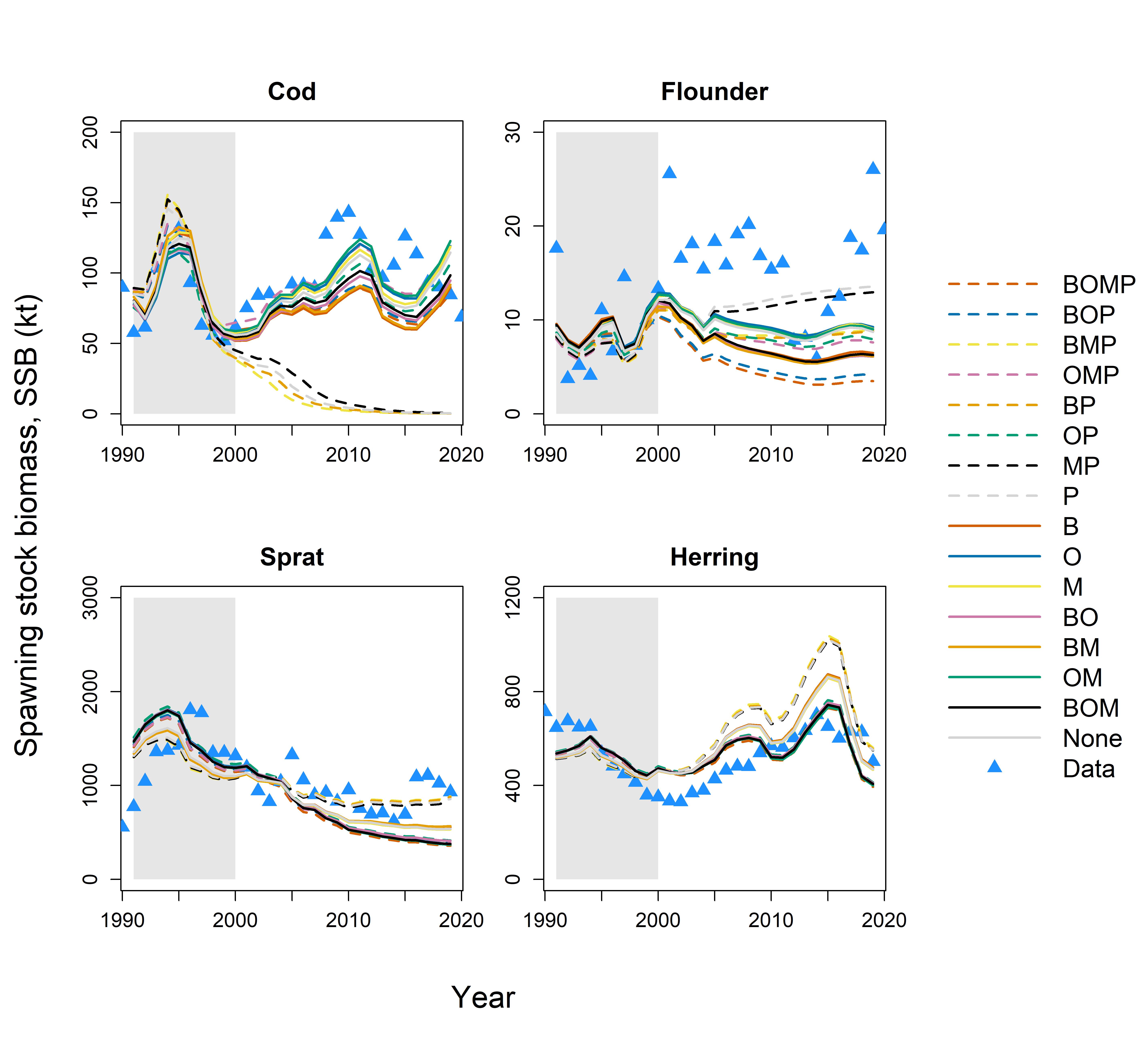

### sim_biomass_best.jpg

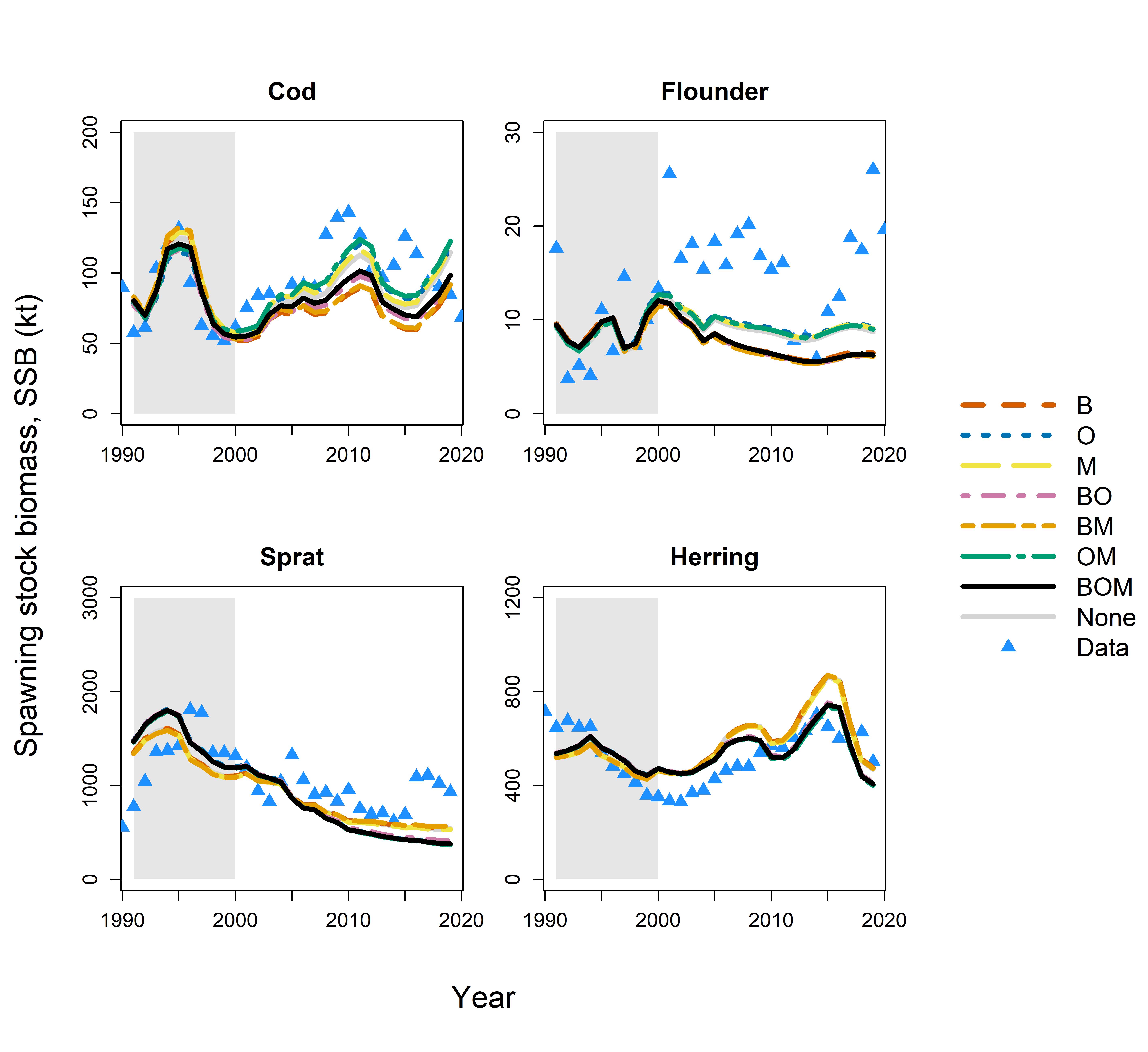

### sim_growth.jpg

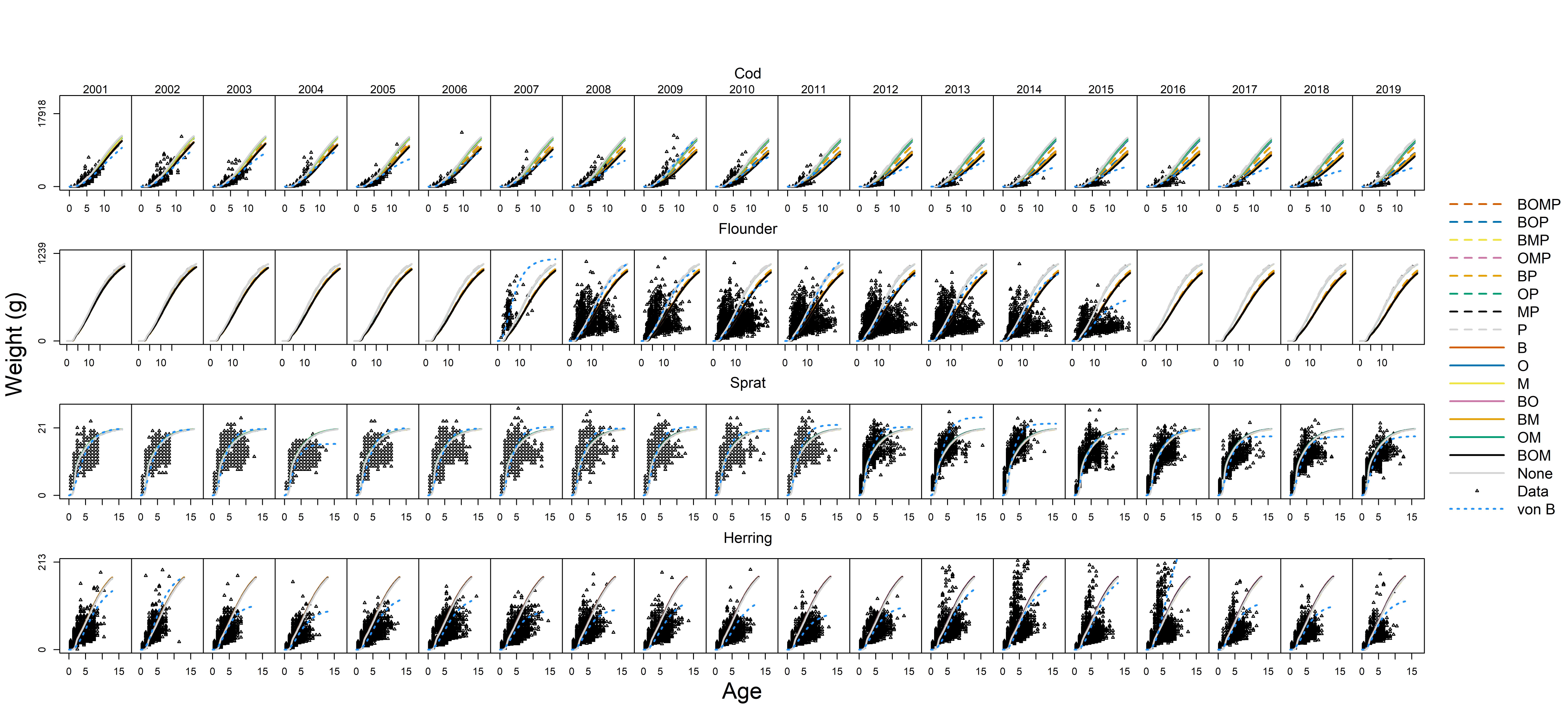

### sim_growth_eight.jpg

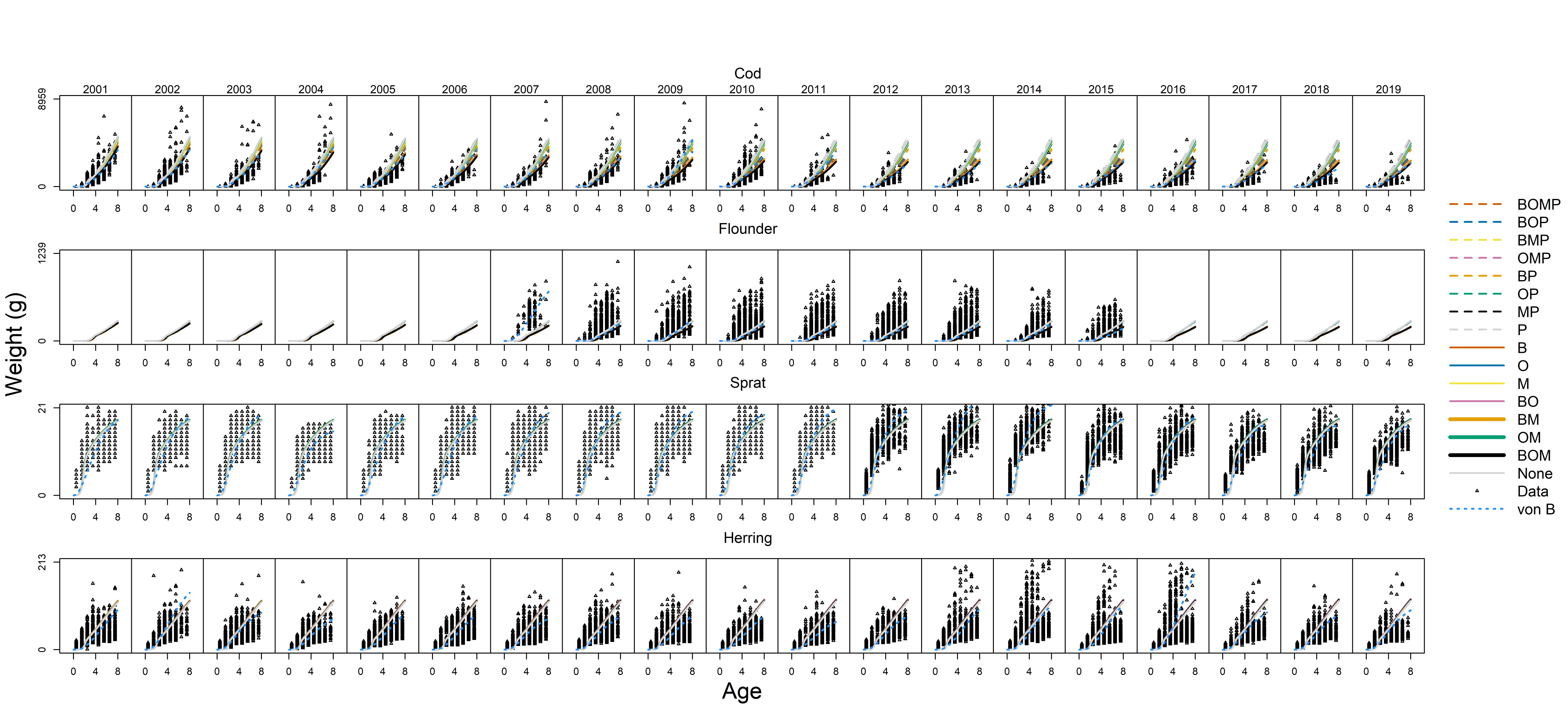

### sim_yield.jpg

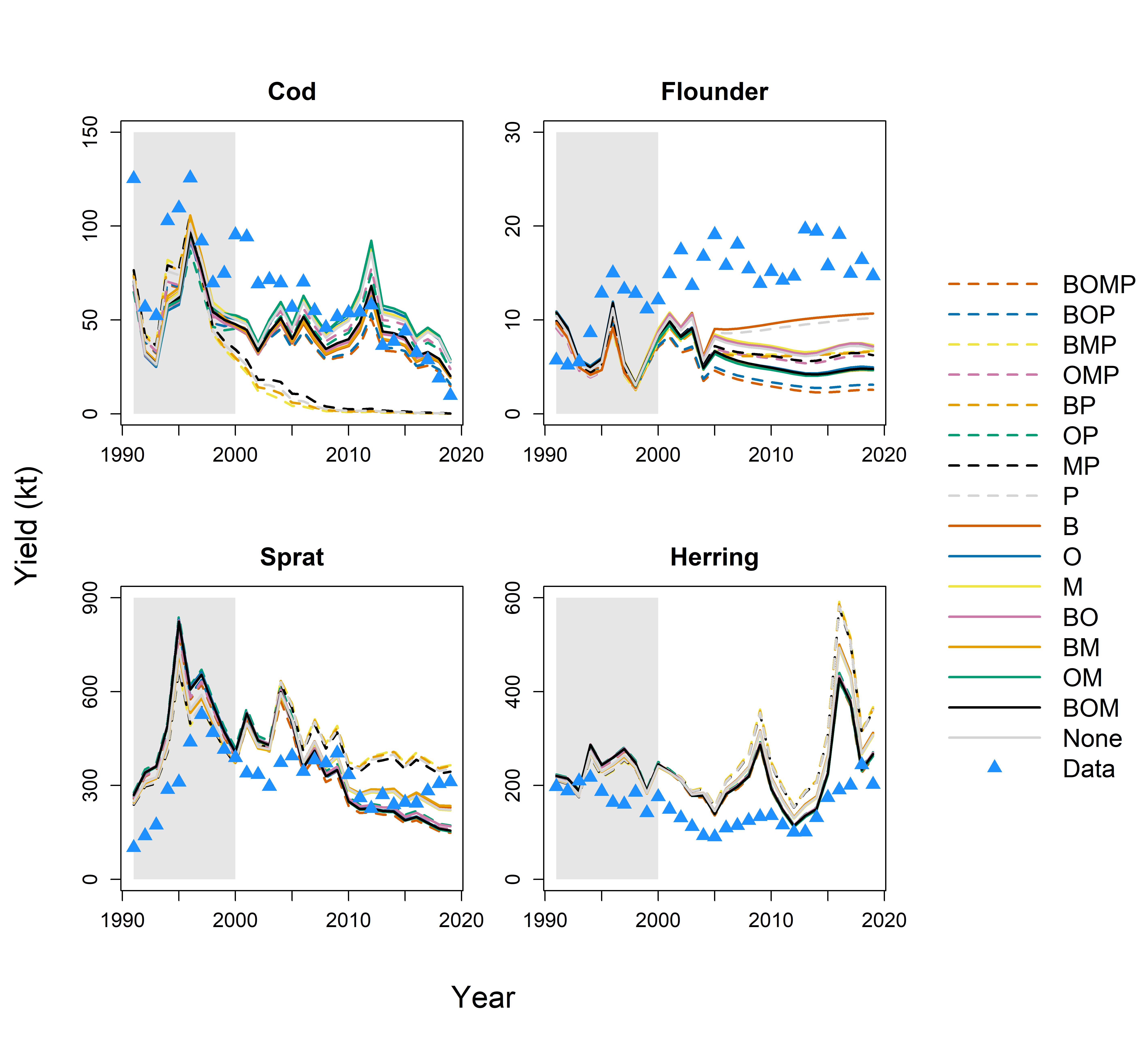

### sim_yield_best.jpg

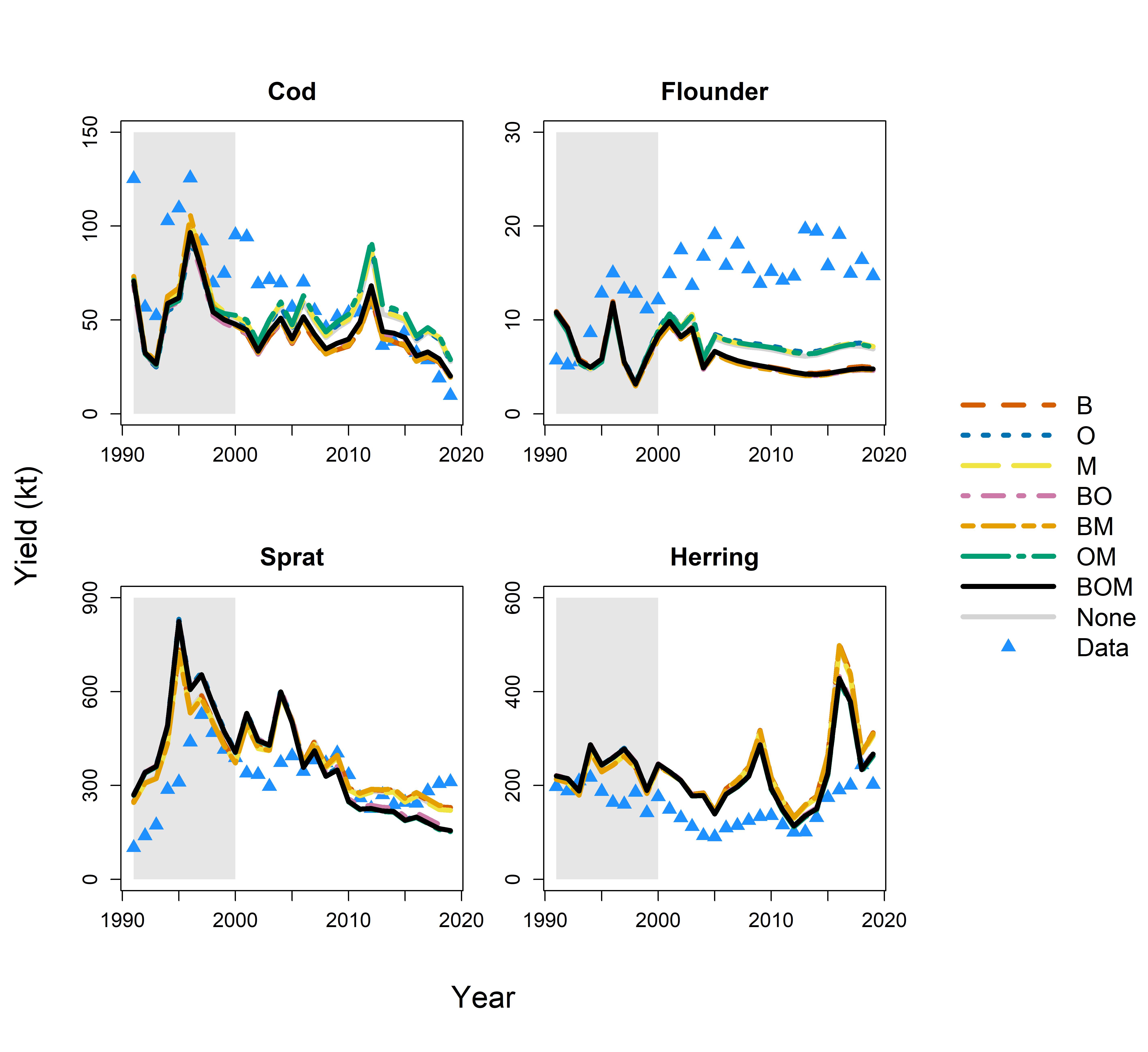

### simulation_error.jpg

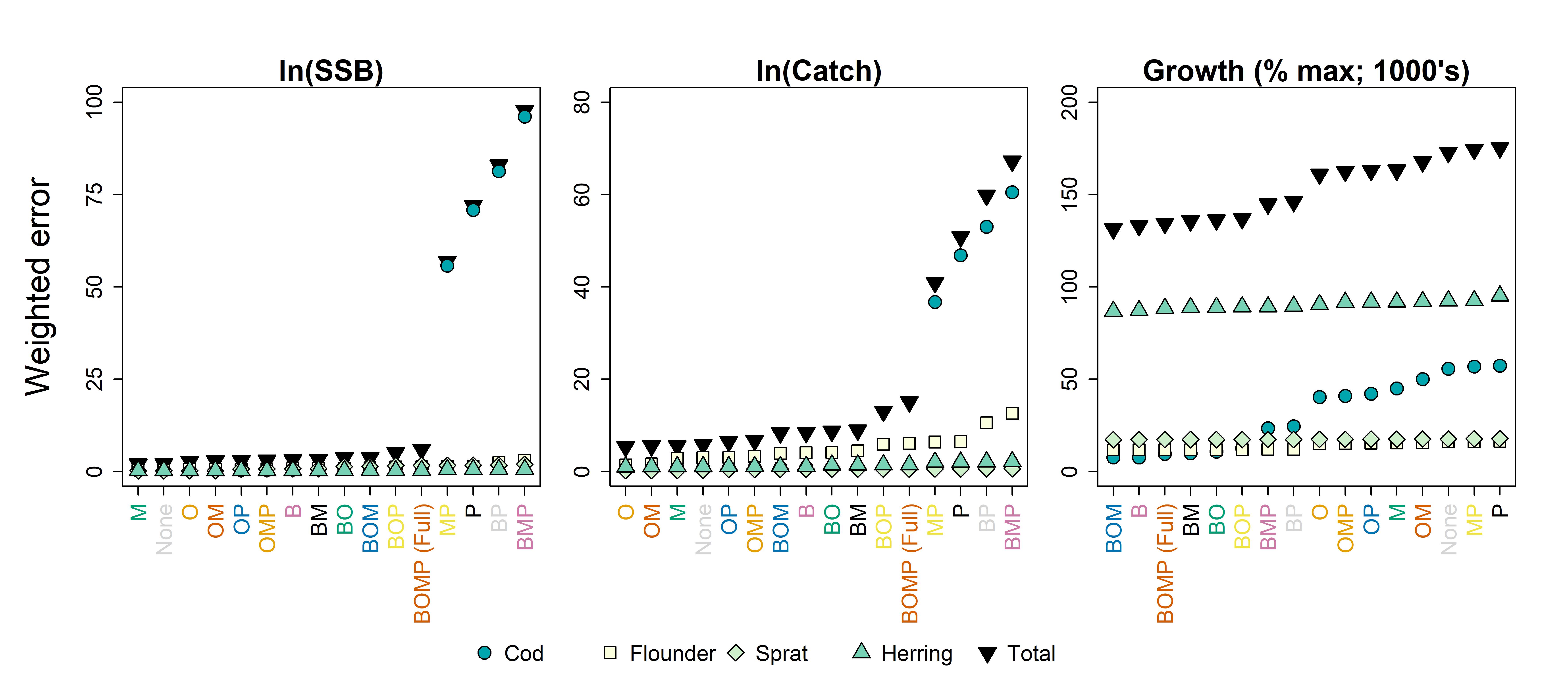

### simulation_error_rank.jpg

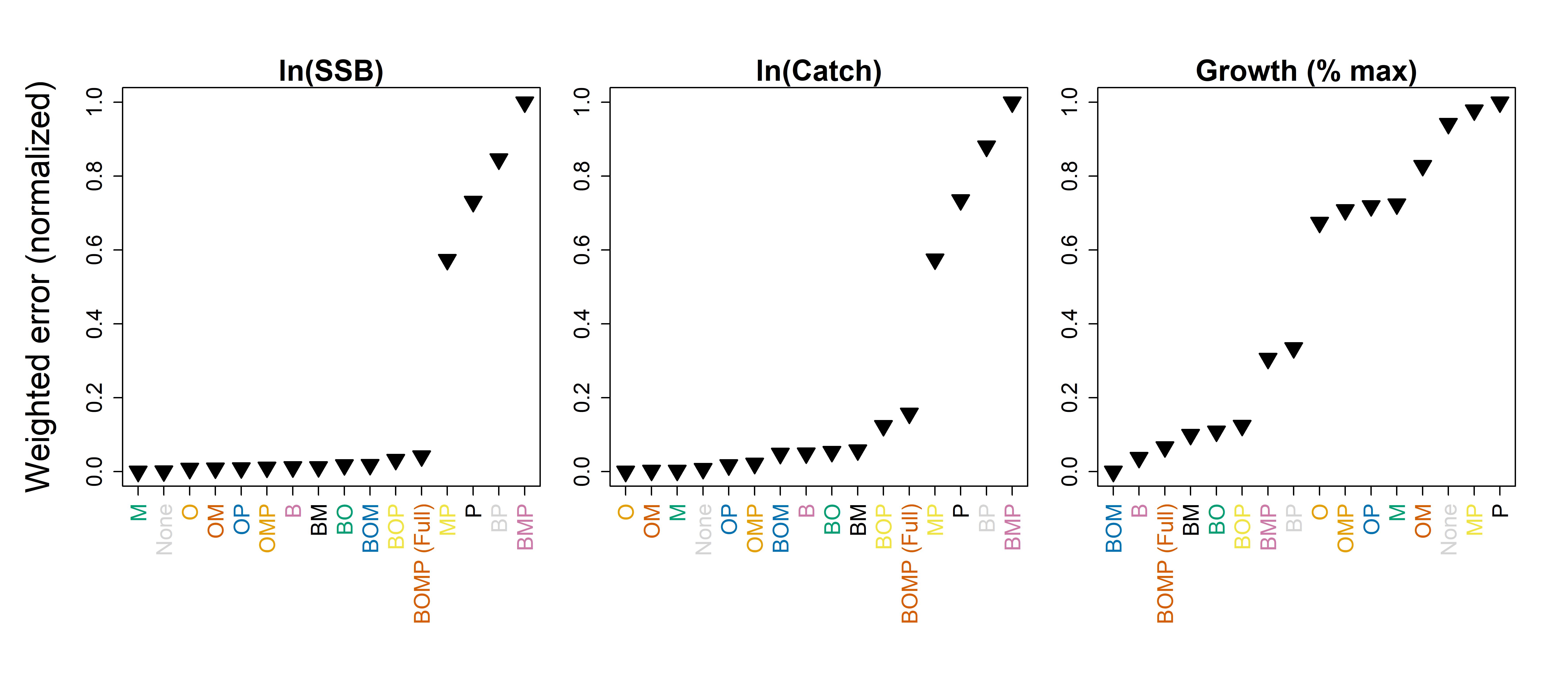

### table_scenarios.jpg

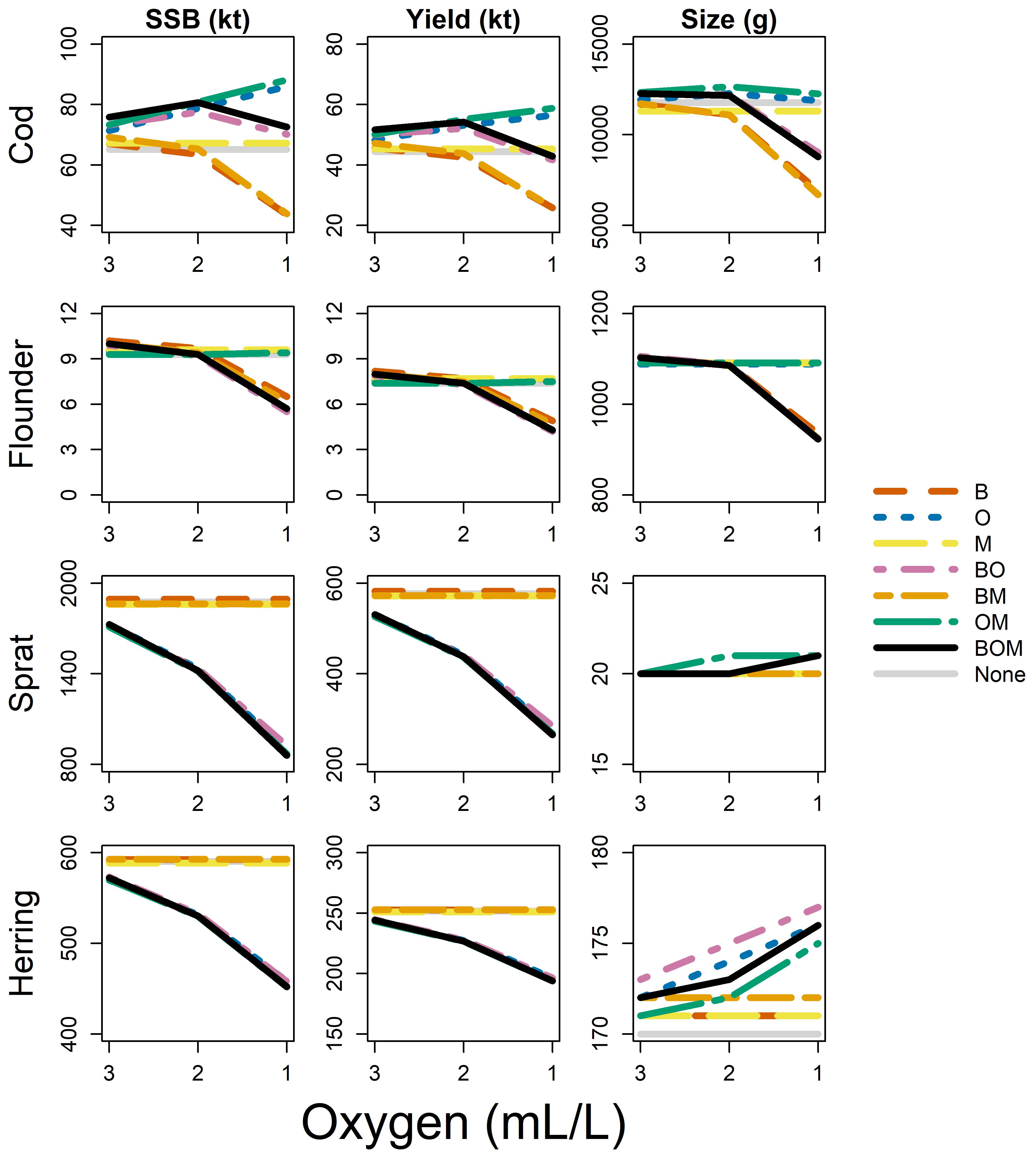
