## Supporting Information for "Declining food availability and habitat shifts drive community responses to marine hypoxia"

#### Contents

**Appendix S1 Model structure** **45**

**Appendix S2 Parameter estimation** **48**

**Appendix S3 Oxygen estimates** **57**

**Appendix S4 $P_{crit}$** **58**

**Appendix S5 Calibration** **60**

**Appendix S6 Sensitivity analysis** **71**

**Appendix S7 Tables and Figures** **73**

### Appendix S1 Model structure

Our novel model structure builds off the baseline provided in **mizer** (Scott et al. 2014). We introduced oxygen dependence for several model components, including benthic resource carrying capacity, fish benthic occupancy, natural mortality, and several physiological rates. We describe each of these components, and the impetus for singling them out, in the sections below.

##### Benthic resources

Benthic resources – in our case, composed primarily of benthic invertebrate prey for cod and flounder species – are typically more tolerant of hypoxia than their benthopelagic predators (Casini et al. 2016). For example, *Saduria entomon*, one of the Baltic cod's primary benthic food sources (Haahtela 1990), does not experience reduced abundance until oxygen falls below 1 $mL \cdot L^{-1}$, well below cod's threshold (Gray et al. 2002). Initially, benthic invertebrates may provide an increasingly abundant food source as low oxygen levels encourage escape to the water column (Lowell and Culp 1999; Eby et al. 2005). As hypoxia develops, biomass of benthic invertebrates also declines, exacerbating competition as primary production is largely redirected to the microbial loop (Baird et al. 2004). Invertebrates may reallocate resources away from reproduction and towards survival strategies, such as increasing ventilation, decreasing both turnover rate and carrying capacity (Calder-Potts et al. 2015). We chose to scale carrying capacity of the benthic resource spectrum with equation 3 (see main text), which we rewrite here for reference.

$$\xi(O_{iwt}) = \frac{1}{1 + \exp\left(-U_i(O_{iwt} - a_i\,\mathsf{P}_{\mathsf{crit},iw})\right)} \tag{S.1}$$

Though, lacking values of $\mathsf{P}_{\mathsf{crit}}$ for the benthos, we lumped $a_i\,\mathsf{P}_{\mathsf{crit},iw}$ into a single value that we call $k_{\mathsf{res}}$ which, along with $U_{\mathsf{res}}$, reflected sensitivity of benthic resources to declining oxygen.

#### Occupancy

As environmental hypoxia develops, fish may alter their spatial distribution within a given system. This may result in changes in overlap and thus in species interactions, leading to altered diets (Hrycik et al. 2017). Evidence suggests that depth distribution of cod, for example, has a complex relationship with environmental factors (Orio et al. 2019). They do not completely avoid hypoxic waters, but rather make brief, frequent visits to deeper waters, likely to forage (Limburg and Casini 2018; Neuenfeldt et al. 2009). As hypoxia increases in frequency, fish mortality may decrease as avoidance behaviors increase, given an alternative habitat is available (Diaz and Breitburg 2009).

We chose to represent avoidance of hypoxia by scaling probability of benthic occupancy, alternatively framed as proportion time spent in benthic habitat (equation S.1). In our case, this would be for cod only, as we consider sprat and herring to be entirely pelagic and flounder to be entirely benthic. We distinguished between the effects of ambient oxygen exposure on habitat choice from the effects on the scaling of physiological rates to represent the trade-off between exposure and access to feeding grounds.

As a baseline, we calculated horizontal overlap of each fish species with each other, called $\theta_{ij}$, from spatially indexed catch per unit effort (CPUE) from bottom trawl surveys for cod and flounder, and acoustic survey estimates for sprat and herring (Appendix S2). We calculated vertical overlap of species as a function of their occupancy calculated within the model. The vertical overlap of species $i$ with species $j$ can be represented by the probability of any individual of species $i$ occupying the same habitat as any individual of species $j$. If we assume the distribution of fish within benthic and pelagic habitats is homogeneous, and that vertical occupancy of a given species is independent of the occupancy all other species, then we can write the probability of co-occurrence as follows:

$$
\begin{aligned}
P(i \cap j) &= P((i \in \text{benthic} \cap j \in \text{benthic}) \cup (i \in \text{pelagic} \cap j \in \text{pelagic})) \\
&= P(i \in \text{benthic} \cap j \in \text{benthic}) + P(i \in \text{benthic} \cap j \in \text{benthic}) \\
&= P(i \in \text{benthic})P(j \in \text{benthic}) + P(i \in \text{pelagic})P(j \in \text{pelagic}) \\
&= \psi_{it}\psi_{jt} + (1 - \psi_{it})(1 - \psi_{jt})
\end{aligned}
$$

So, $\theta_{ijt}$, or the overlap of species $i$ with species $j$ at time $t$, is then:

$$\theta_{ijt} = \theta_{ij}\left(\psi_{it}\psi_{jt} + (1 - \psi_{it})(1 - \psi_{jt})\right) \tag{S.2}$$

Given that occupancy is size dependent, it is more accurate to write $\theta_{i_w j_{w_p}}$, where $w$ and $w_p$ are predator and prey weight, respectively. This product essentially amounts to a three-dimensional representation of overlap between species. While, realistically, deleterious effects of low oxygen levels may require relatively prolonged episodes of exposure, we calculated exposure, referred to as $O_{iwt}$ in equation S.1, as a simple weighted average of the probability of occupancy within the benthic and pelagic habitats. This exposure determines the degree of scaling of mortality and physiology. We fit the parameters describing sensitivity of occupancy to oxygen, referred to as $U_{hab}$ and $a_{hab}$, to the ratio of benthic to pelagic CPUE of cod.

#### Mortality

Direct mortality due to exposure to hypoxia has been documented through observations of fish kills and experimental exposure (e.g. Diaz 2001; Shimps et al. 2005; Small et al. 2014). We included potential increases in mortality in hypoxic waters via hazard functions of the form:

$$h(t) = b_{0,i} e^{b_i O_{iwt}} \tag{S.3}$$

where $O_{iwt}$ is, as above, average oxygen level experienced by an individual of species $i$ of size $w$ at time $t$, $b_{0,i}$ gives mortality at zero oxygen and $b_i \leq 0$ reflects sensitivity of mortality to oxygen. The primary outcome of adjusting natural mortality in the **mizer** framework is the death of larger individuals, which is typically used to adjust modeled levels of yield. Therefore, we fit $b_{0,i}$ and $b_i$ for cod and flounder to annual observations of yield.

#### Physiology

Dependence of physiological rates on oxygen consists of both increases in metabolic costs and general decreases of activity with decreasing oxygen levels. We describe these pieces together as all physiological parameters are fit simultaneously during the calibration procedure. Scaling of metabolic costs is determined by parameters that we call $U_{\mathsf{met}}$ and $a_{\mathsf{met}}$ of equation 4 (see main text), again repeated here:

$$\xi(O_{iwt}) = 1 + \exp\left(-U_{\mathsf{met},i}(O_{iwt} - a_{\mathsf{met},i}\,\mathsf{P}_{\mathsf{crit},iw})\right) \tag{S.4}$$

All other physiological scaling is determined by $U_{\mathsf{phys}}$ and $a_{\mathsf{phys}}$ of equation S.1. These are all fit to observations of annual growth of all species.

##### Metabolism

As ambient oxygen levels fall, fish must expend more energy to maintain internal oxygen, whether through increased ventilation, movement, aquatic surface respiration, or other compensatory behaviors (Kramer 1987). This may have an indirect effect on the availability of energy for other activities. When oxygen falls to levels low enough to stimulate the onset of anaerobic glycolysis, individuals may incur additional metabolic costs. Though hypoxia tolerant fish may suppress metabolic rate at very low oxygen levels prior to the onset of anaerobiosis (Muusze et al. 1998), they too will eventually incur these same costs. Metabolic costs in **mizer** are a product of the following: critical feeding level $f_c$, or the feeding level below which intake is less than metabolic costs; maximum consumption $h_{iwt}$; and assimilation efficiency $\alpha_{iwt}$ (Scott et al. 2014). Scaling of metabolic costs indicate that consequences of exposure to hypoxia increase exponentially as oxygen continues to decline.

##### Feeding behavior

The **mizer** framework assumes feeding follows a Type II functional response, which is determined by encounter rates and maximum consumption rates, the former of which is in turn a function of clearance rates and prey size preference (see Andersen et al. 2016, Table 1). One of the most commonly cited causes of growth rate reductions in response to hypoxia is the loss of appetite (Wang et al. 2009). In fact, fish exposed to hypoxia and fish fed a reduced diet may appear quite similar (Pichavant et al. 2001). Overall, energy acquisition declines both as intake declines and as specific dynamic action (SDA), or the costs of digestion and assimilation, increase. Cod, for example, consume large meals, followed by a period of fasting, during which SDA may account for up to 90% of maximum metabolic rate (MMR) (Wang et al. 2009). Therefore, reductions in feeding may free a significant portion of the aerobic scope. We down-scaled maximum consumption to reflect a decline in feeding with falling oxygen levels.

Behavioral responses to vary with the severity of hypoxia, and with the natural character of a given species in normoxia. Relatively sedentary benthic and benthopelagic species may decrease activity to conserve oxygen, whereas more active and pelagic species may increase activity (Chapman and Mckenzie 2009). Cod, for example, are active benthopelagic predators, and will initially increase activity in response to developing hypoxia, perhaps as a means of escaping unfavorable conditions (Chapman and Mckenzie 2009). However, as oxygen continues to decline, activity tends to decrease. Thus, we also down-scaled search or clearance rate with decreasing oxygen in an attempt to capture changes in feeding activity.

The **mizer** framework also includes a single assimilation value for all species at all sizes. Assimilation efficiency may decline in response to hypoxia, though some studies show no effect (Wang et al. 2009). Jordan and Steffensen (2007) showed that exposure to hypoxic conditions increases the amount of time required for digestion and decreases the overall energy acquisition from digestion of meals. Though evidence is somewhat mixed, we chose to account for increased costs of feeding with declining assimilation efficiency when oxygen is low.

##### Reproduction efficiency

Hypoxia affects fish reproduction through two major pathways. First, exposure to low oxygen may disrupt hormones and enzymes which regulate reproduction and development of eggs, leading to egg mortality. In our study system, cod eggs are neutrally buoyant at salinities of 10-12 psu, and may therefore sink below the halocline, resulting in exposure to hypoxic waters, altered development, and ultimately mortality (Baird et al. 2004). Second, declines in energy acquisition due to decreased activity may reduce energy available for reproduction (Wu 2009). Declining energy acquisition is addressed by oxygen dependence of feeding behavior. We included egg mortality by scaling reproductive efficiency (see Scott et al. (2014)) as a logistic function of oxygen, which is a measure of egg survival or rather production.

#### Appendix S2 Parameter estimation

We parameterized the model with several data sources (Table S1). These parameters describe growth, maturation, selectivity, feeding preferences, and species interactions. We used the years 1991-2000 as a calibration period. The ideal calibration period is one which is comprised of several years during which conditions were stable. In our case, conditions within the Baltic Sea were deteriorating during the years 1991-2000. However, 1991 is the earliest year for which data are available for all species for nearly all model components, excepting the European and Baltic flounders (*Platichthys flesus* and *Platichthys solemdali*, respectively), for which there are limited data on stomach contents, weight, length, age, and maturity. For these fish, we used all available data for calibration, which correspond to the mid-2000's through the mid-2010's. This introduces the implicit assumption that flounder life history was identical throughout this period and in 1991-2000. Though perhaps unfounded, we are limited by the scope of the data.

##### Growth

We estimated body growth parameters for all four species using the inverse of the von Bertalanffy function:

(S.5) $$f(L) = -\frac{1}{k_{vb}} \ln\left(1 - \frac{L}{L_\infty}\right) + t_0$$

Table S1: Model parameters and data sources for each species. CPUE is catch per unit effort, w/l/a/m are weight/length/age/maturity, and F is fishing mortality.

| Parameters | Description | Data | Species | Years | Source |
|---|---|---|---|---|---|
| $\beta, \sigma$ | Mean and SD of predator-prey mass ratios | stomach | *Gadus morhua* | 1963-2014[a],2015-2018[b] | a,b |
| | | | *Platichthys flesus* | 2015-2018[b] | b |
| | | | *Clupea harengus* | 1999-2000[c] | c |
| | | | *Sprattus sprattus* | 1999-2000[c] | c |
| $\theta_{ij}$ | Horizontal degree to which species $i$ and $j$ overlap | CPUE/abundance | *G. morhua* | 1991-2020[d] | d |
| | | | *P. flesus* | 1991-2020[d] | d |
| | | | *C. harengus* | 1979-2020[e] | e |
| | | | *S. spratus* | 1979-2020[e] | e |
| $w_m$ | Weight at which probability of maturation is $0.5$ | w/l/a/m | *G. morhua* | 1991-2020[d] | d |
| | | | *P. flesus* | 2007-2015[d] | d |
| | | | *C. harengus* | 1979-2020[e] | e |
| | | | *S. spratus* | 1979-2020[e] | e |
| $W_\infty, k_{vb}, t_0, a, b$ | von Bertalanffy growth and length-weight conversion | w/l/a | *G. morhua* | 1991-2020[d] | d |
| | | | *P. flesus* | 2007-2015[d] | d |
| | | | *C. harengus* | 1979-2020[e] | e |
| | | | *S. spratus* | 1979-2020[e] | e |
| $L_{25}, L_{50}$ | Lengths at 25% and 50% selectivity | F | *G. morhua* | 1946-2019[f] | f |
| | | | *P. flesus* | – | – |
| | | | *C. harengus* | 1974-2019[f] | f |
| | | | *S. spratus* | 1974-2019[f] | f |
| $L_{k\text{-}e}$ | Lengths at 100% selectivity, or knife-edge | F | *G. morhua* | – | – |
| | | | *P. flesus* | 1955-2004[g] | g |
| | | | *C. harengus* | – | – |
| | | | *S. spratus* | – | – |

[a] ICES Year of the Stomach; ICES (2020c)
[b] Haase et al. (2020)
[c] Casini et al. (2004)
[d] ICES CPUE from BITS survey database DATRAS ICES (2020b)
[e] ICES BIAS acoustic survey estimates; (ICES 2020a)
[f] (ICES 2020a)
[g] Draganik et al. (2007)

where $L$ is the length of a given individual, $k_{vb}$ is the von Bertalanffy growth factor, $L_\infty$ is the asymptotic length, $t_0$ is age at size zero, and $f(L)$ is estimated age. We extracted sex-maturity-age-length-key (SMALK) data from DATRAS (ICES 2020b) and used size and age observations for cod and flounder. For sprat and herring, we used size and age observations from BIAS trawl surveys (ICES 2020a). The data (Table S1) are given as weight and age of individuals within a given size class. Data are also given for a relatively wide range of size classes for each species. Therefore, the inverse von Bertalanffy is ideal (Mackay and Moreau 1990). Fish at age zero were collected roughly half a year post-spawning, thus we added 0.5 to reported age and assumed size at birth, or $t_0$, was equal to zero, or close enough to zero to warrant dropping the parameter.

We chose to run all models for each species in a Bayesian framework within **Stan** (Carpenter et al. 2017). We used the following priors:

$$
\begin{aligned}
k_{vb} &\sim \mathsf{Uniform}(0, 5) \\
L_\infty &\sim N(\mu_{L_\infty}, \sigma^2_{L_\infty})\, T[L_{\mathsf{min}},] \qquad \text{(S.6)} \\
\sigma^2_{\mathsf{age}} &\sim \mathsf{Cauchy}(0, 1000)
\end{aligned}
$$

where $\mu_{L_\infty}$ and $\sigma^2_{L_\infty}$ were the mean and variance of the $L_\infty$ prior distribution for any given species, $L_{\mathsf{min}}$ was the minimum possible value for $L_\infty$, and $\sigma^2_{\mathsf{age}}$ was a hyperparameter describing variance in estimated ages (see equation S.7). The addition of a $T[x,]$ or $T[,x]$ to a distribution indicates that it is truncated from below or above, respectively. We used the following likelihood:

$$\text{(S.7)} \qquad \text{age} \sim N\left(-\frac{1}{k_{vb}}\ln\left(1-\frac{L}{L_\infty}\right), \sigma^2_{\text{age}}\right)$$

During implementation, we chose $\mu_{L_\infty} = 100$ for all species, which implies that the resulting prior is only moderately informative. We chose $\mu_{L_\infty}$ as the maximum observed length class for each species, plus 1 mm. Similarly, we chose $L_{\text{min}}$ as the maximum observed length class. This prevented failures in the model due to attempting to take the natural log of a negative number. The **mizer** framework requires body size in grams, and therefore we estimated length-weight relationships using nonlinear least squares for each species with the following standard equation:

$$\text{(S.8)} \qquad W = aL^b$$

Results were typical, and apparent outliers had little to no observable effect on estimates (Figure S1).

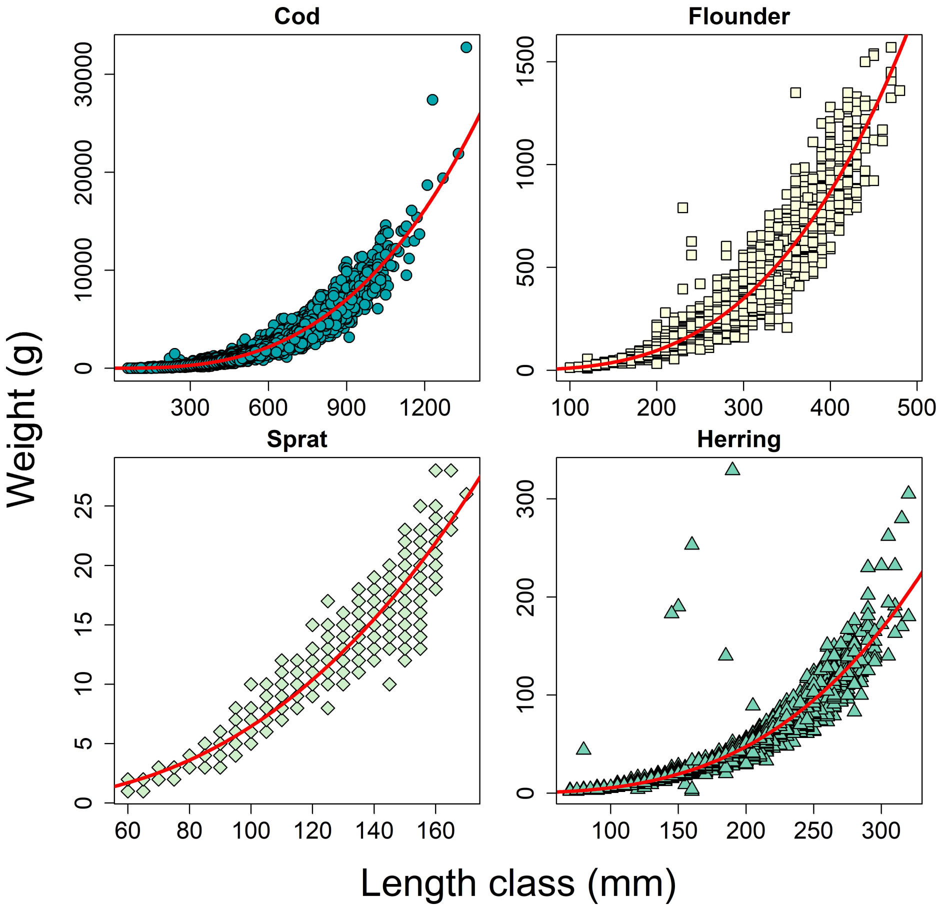


Figure S1: Length-weight relationships for each species. The solid red lines give mean predictions.

We ran the Bayesian model on size-at-age data for all four species. For cod and flounder, we partitioned the data sets by ICES subdivision, sex, and year. For sprat and herring, we partitioned the data sets by ICES subdivision and year. The former two exhibit sexual dimorphism while the latter do not. We ran the model separately on each data set, excluding those data sets with fewer than 25 points, which resulted in a total of 193 models across all species. We ran each model for 2,000 samples as warmup, followed by 20,000 iterations over four chains running in parallel, resulting in 80,000 samples from each model. Models finished anywhere from 3 to 147 seconds, with a median time of 11 seconds, a mean of 19 seconds, and a total of 62 minutes. Note that we increased the default maximum tree depth to 20, and increased the target average acceptance probability to 0.99 so that all models would finish with no divergent transitions (Carpenter et al. 2017). All models finished with more than 100 effective samples for both the bulk and the tail, indicating adequate sampling across posterior quantiles. We took the area-weighted median of each estimate to arrive at final estimates for each species (Table S2). Resulting predictions of length-at-age agree fairly well with observations (Figure S2). Apparent biases are due largely to data partitioning and area weighting.

Table S2: All parameter estimates for all four species, when applicable. See parameter descriptions in Table S1.

| Parameter | Description | Unit | Cod | Flounder | Sprat | Herring |
|---|---|---|---|---|---|---|
| $\beta$ | Mean preferred predator prey mass ratio | – | 673 | 937 | 173302 | 702819 |
| $\sigma$ | SD preferred predator prey mass ratio | – | 1.63 | 1.70 | 0.74 | 0.52 |
| $w_m$ | g | 577 | 64 | 7 | 20 | |
| $L_\infty$ | Maximum length | mm | 1243 | 447 | 157 | 324 |
| $W_\infty$ | Maximum weight | mm | 17918 | 1239 | 21 | 213 |
| $k_{vb}$ | von Bertalanffy growth | $y^{-1}$ | 0.13 | 0.12 | 0.43 | 0.22 |
| $a$ | Length-weight coefficient | $10^{-5}$ | 1.99 | 0.48 | 4.09 | 0.34 |
| $b$ | Length-weight exponent | – | 2.89 | 3.18 | 2.60 | 3.11 |
| $L_{25}$ | Length at 25% selectivity | mm | 402 | – | 121 | 140 |
| $L_{50}$ | Length at 50% selectivity | mm | 489 | – | 130 | 169 |
| $L_{k\text{-}e}$ | Knife-edge selectivity | mm | – | 180 | – | – |
| $F_{\text{max}}$ | Maximum fishing mortality | $y^{-1}$ | 1.00 | 0.90 | 0.34 | 0.47 |
| $F_{k\text{-}e}$ | Knife-edge fishing mortality | $y^{-1}$ | – | 0.90 | – | – |

#### Maturation

We calculated probabilistic maturation reaction norms (PMRN; Dieckmann and Heino 2007) for all four species. The PMRN attempts to remove influences of growth and mortality on maturity, thereby isolating variability in size at maturation (Barot et al. 2003). We included age, length, and Fulton's condition factor ($W/L^3$) in our model, as in Vainikka et al. (2009). In our case, then, the PMRN is given by:

(S.9)
$$m(a,s,c) = \frac{o(a,s,c) - o(a-1, s-\Delta s, c-\Delta c)}{1 - o(a-1, s-\Delta s, c-\Delta c)}$$

where $\Delta s$ and $\Delta c$ refer to annual increments in weight and condition factor, respectively. Probability of maturation at a given age, size, and condition factor is thus conditional on those individuals who have already matured in the previous time step, an extension of the relationship described by (Barot et al. 2003).

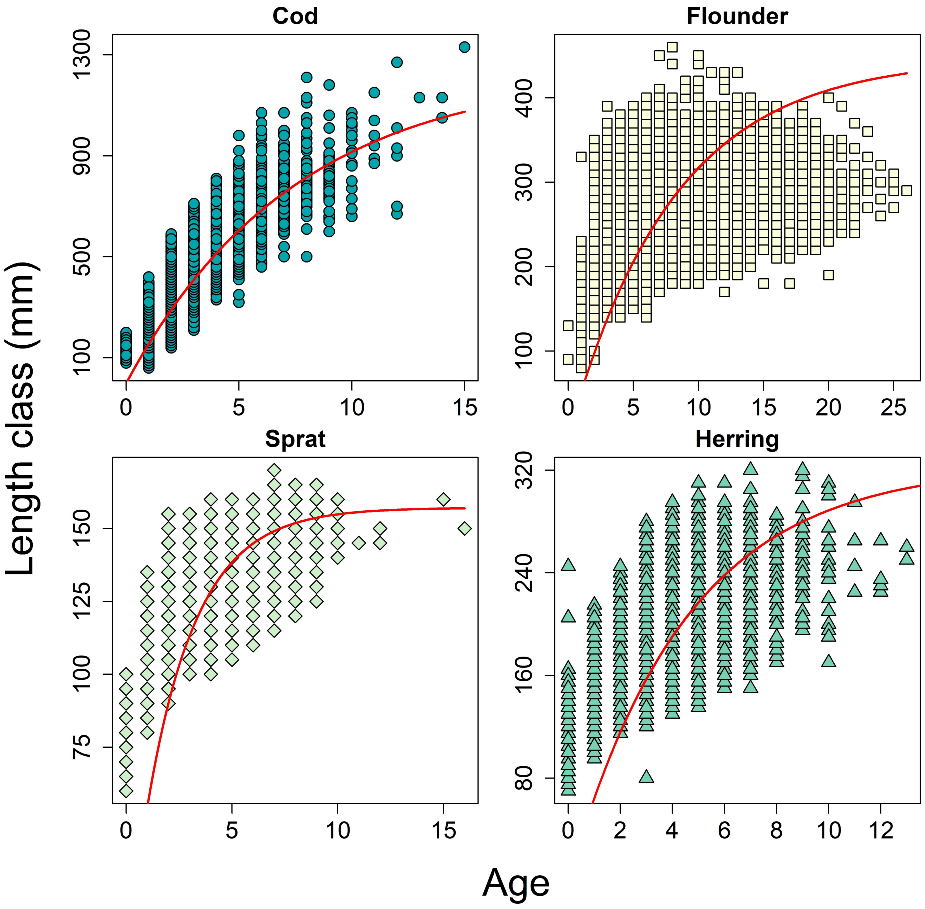


Figure S2: Length-at-age data for each species. Solid, red lines give model estimates. Note that while data are pooled here, estimates arise from data partitioning (see text).

We extracted data on maturation from the BITS survey database DATRAS. For cod and flounder, we downloaded sex-maturity-age-length-key (SMALK) data, and used the ICES maturity key to determine whether or not each individual was mature, converting from the Maier scale to the ICES scale when necessary (Grygiel and Wyszyński 2002; *Manual for the Baltic International Trawl Surveys* 2017). On the ICES scale, fish in stage 1 are immature, and the remaining stages are mature. For sprat and herring, we used BIAS trawl data, and again categorized fish based on the ICES scale. For all four fish, we used von Bertalanffy growth function parameters to calculate growth increments, and simple linear models to estimate body condition increments. We then modelled maturity ogives as a logistic function of age, weight, and condition, and all possible interactions. We performed simple stepwise model selection using AIC as criteria. The models retained the following covariates: for cod, age, weight, condition, and weight*condition; for flounder, all covariates and possible interactions; for sprat, only age and weight; and for herring, all covariates besides age*weight*condition.

We used age estimates, weight and condition increments, and ogives to calculate probability of maturation, or PMRN, for each individual of each species as in equation S.9. We only used individuals in age groups over which each species typically matures: ages 2 to 3 for cod, 1 to 2 for flounder, 1 to 2 for sprat, and age 1 for herring. Finally, we used a sigmoid function to model PMRN as a function of weight:

(S.10)
$$\mathsf{PMRN} = \left[1 + \left(\frac{L}{L_{50}}\right)^{-10}\right]^{-1}$$

where $L$ is length of an individual, and where $L_{50}$, the parameter of interest, is the length at which the probability of maturation is $0.5$. The exponential value of -10 ensures that the model is not overwhelmed by data concentration at larger sizes, at which fish have a higher probability of maturation. When we chose to estimate the exponent of equation S.10, the model treated smaller fish as outliers. This somewhat arbitrary choice asserts our knowledge that this small pool of immature fish are critical data (Figure S3). We converted length to weight values using each species' length-weight parameters (Table S2) to arrive at weight at maturity.

#### Selectivity

We used estimates of fishing mortality at age from stock assessments (ICES 2020a) to estimate selectivity parameters for cod, sprat, and herring. We used the following ogive function:

(S.11)
$$S(L) = \frac{1}{1 + \exp\left(\frac{L_{50}\ln(3)}{L_{50}-L_{25}} - \frac{\ln(3)}{L_{50}-L_{25}}L\right)}$$

where $S(L)$ is selectivity – or normalized fishing mortality – as a function of length, and $L_{25}$ and $L_{50}$ are lengths at 25% and 50% selectivity. We converted age to length using von Bertalanffy parameters (Table S2). We retained the maximum estimated fishing mortality for each species and treated length-specific fishing mortality as a product of length-specific selectivity and maximum fishing mortality.

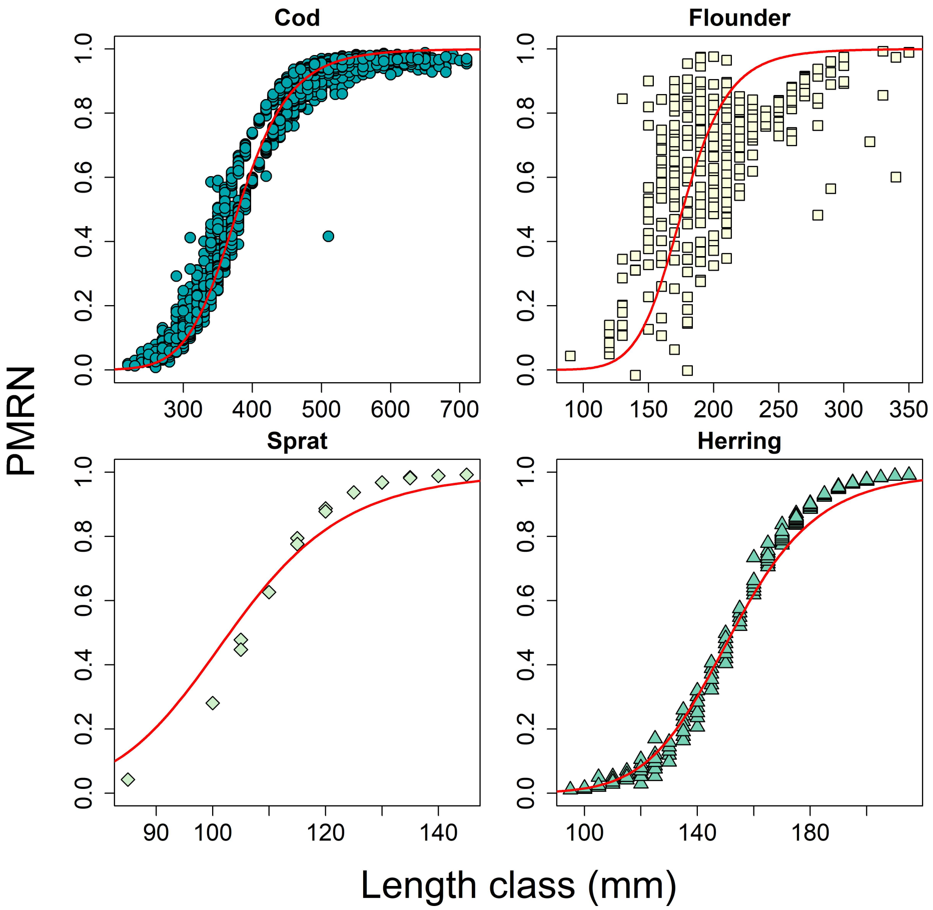


Figure S3: Estimated probability maturation reaction norms (PMRN) for individuals of each species. We only included individuals of ages 2 to 3 for cod, 1 to 2 for flounder, 1 to 2 for sprat, and age 1 for herring. Solid red lines are fits to equation S.10.

Currently, the ICES stock assessment lacks fishing mortality estimates for flounder. Draganik et al. (2007) estimated fishing mortality of Baltic flounder stocks in ICES subdivision 26 using extended survivor analysis (XSA) (Shepherd 1999). Yields in this subdivision tend to be analogous to yields throughout the Baltic (Draganik et al. 2007). Therefore, the identified trends may be accurate, though the actual regional value may be less than that reported for subdivision 26, given fishing mortality tends to be lower in subdivisions 24 and 25. We extracted $F_{bar}$ from this study (Draganik et al. 2007, see Figure 10 op. cit.) using WebPlotDigitizer (Rohatgi 2021). These data were available for the years 1955–2004. For years beyond 2004, we used the terminal-three year mean of $F_{bar}$. We calculated the mean of $F_{bar}$ for the years 1991–2000, and used a knife-edge selection size of 18cm, corresponding to the minimum legal size in Estonia, and in fact the smallest of all minimum legal sizes (ICES 2020a). As above, we treated fishing mortality as a product of selectivity and maximum fishing mortality. Results indicate that models agree well with observations (Figure S4).

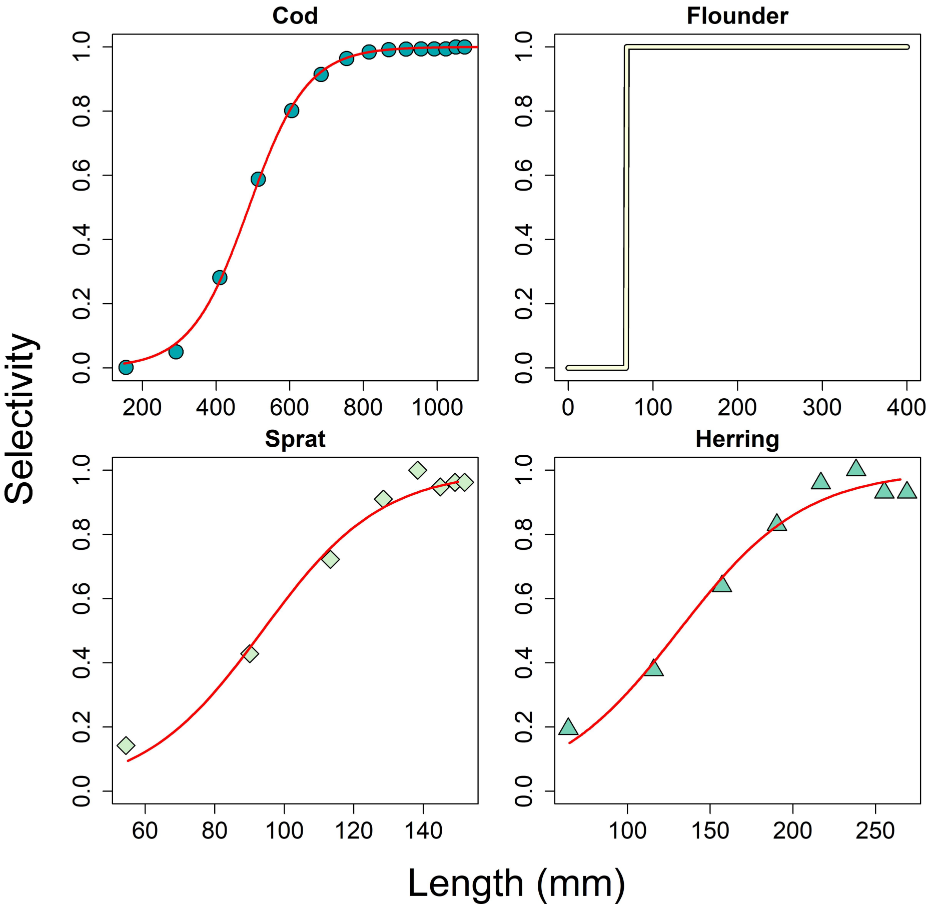


Figure S4: Estimated selectivity for all four species. Data points are normalized estimates of fishing mortality at age, with age converted to length. Selectivity for flounder is given by a knife-edge, with all individuals at or above 180mm fully selected by the fishery.

#### Prey preference

The **mizer** framework assumes that prey selection is a function of the ratio of predator and prey weight. That is, bigger fish eat smaller fish; how much bigger and smaller dependent upon the species in question. We used a log-normal function to formalize this relationship as a function of a predator of species $i$ and of size $w$:

$$\phi\left(\frac{w_{\text{prey}}}{w}\right) = \exp\left(\frac{-\ln\left(\beta_i(w)\, w_{\text{prey}} - w\right)^2}{2\sigma_i^2}\right) \quad \text{(S.12)}$$

We calculated PPMR from stomach data for each species. We limited our analyses to prey items in early stages of digestion, and took the ratio of predator size in g to prey size in g. We calculated the log of these ratios, and extracted the mean ($\beta$) and standard deviation ($\sigma$) for each species. While we limited these calculations to data available during the years 1991–2000 for cod, data were only available during 2015–2018 for flounder.

For herring and sprat, we used the WebPlotDigitizer (Rohatgi 2021) to extract feeding data from Figure 3 of Casini et al. (2004). These data were recorded as percent composition of eight major species or groups of zooplankton, amphipods, and mysids in the diets of several size classes of both herring and sprat. We used documented wet weights of each prey item to calculate the mean size of prey items appearing in clupeid stomachs (Hernroth 1985). When wet weights were partitioned among regions, we used data from "Remaining Baltic Sea". For *Temora longicornis*, we used mean wet weight of copepodite stages CIV-CV and CVI in October through December as a typical prey item for sprat, which dominate in the fall (Möllmann and Köster 2002), and the wet weight of stages CI-CIII, CIV-CV, and CVI for herring. We included mean weight of stages CI-CIII because herring are observed to switch prey sources from larger stages to stage CII when in competition with high densities of sprat (Möllmann and Köster 2002). For *Bosmina longispina maritima*, *Podon intermedius*, and *Pleopis polyphemoides* (syn. *Podon polyphemoides*) we used the mean wet weight across all size classes for October through December, for lack of more detailed information. For *Acartia spp.*, we calculated the mean wet weight of all size classes of *Acartia longiremis* from January through March for Herring, and for *Acarctia bifilosa* from April through December for sprat, given that these time frames correspond roughly to seasons during which each species is observed to feed on these copepods (Möllmann and Köster 1999). The exact choice of time period and species is not critical, as wet weights are similar throughout the year and within *Acartia spp.*. We used a mean length of 1.75 cm for *Mysis mixta*, based on the size range of 1.5–2.0 cm for mysids in the Baltic Sea (Snoeijs-Leijonmalm 2017). For the group labeled "amphipods", we used a rough estimate for body length of 0.9cm, based on size range of 0.6–1.2cm (Hill et al. 1992). We converted to weight in grams using the relationship: $\text{weight} = 0.01 \cdot \text{length}^3$.

The data extracted from Casini et al. (2004) served as weights with which to calculate the mean of a typical prey item for each size class of herring and sprat. We converted each size class from length in cm to weight in g using estimated length-weight relationships. We used the resulting PPMR values from all four species in the models and calculations described above (Figure S5 and see Table S2).

**Species interactions**

Species interactions emerge as a function of spatial overlap in both the horizontal and vertical directions. Vertical overlap is a dynamic model component (see main text), whereas we treated horizontal overlap as parameters. We input horizontal spatial overlap as an index calculated from data:

$$D(p_{it}, p_{jt}) = 1 - \frac{1}{2} \sum_k |p_{itk} - p_{jtk}| \tag{S.13}$$

where $p_{itk}$ is the proportion of the total abundance index of species $i$ in ICES statistical rectangle $k$ at time t(Blanchard et al. 2014). In our case, these abundance indices arose from BITS for cod and flounder, and BIAS acoustic estimates for sprat and herring (Table S1). We calculated

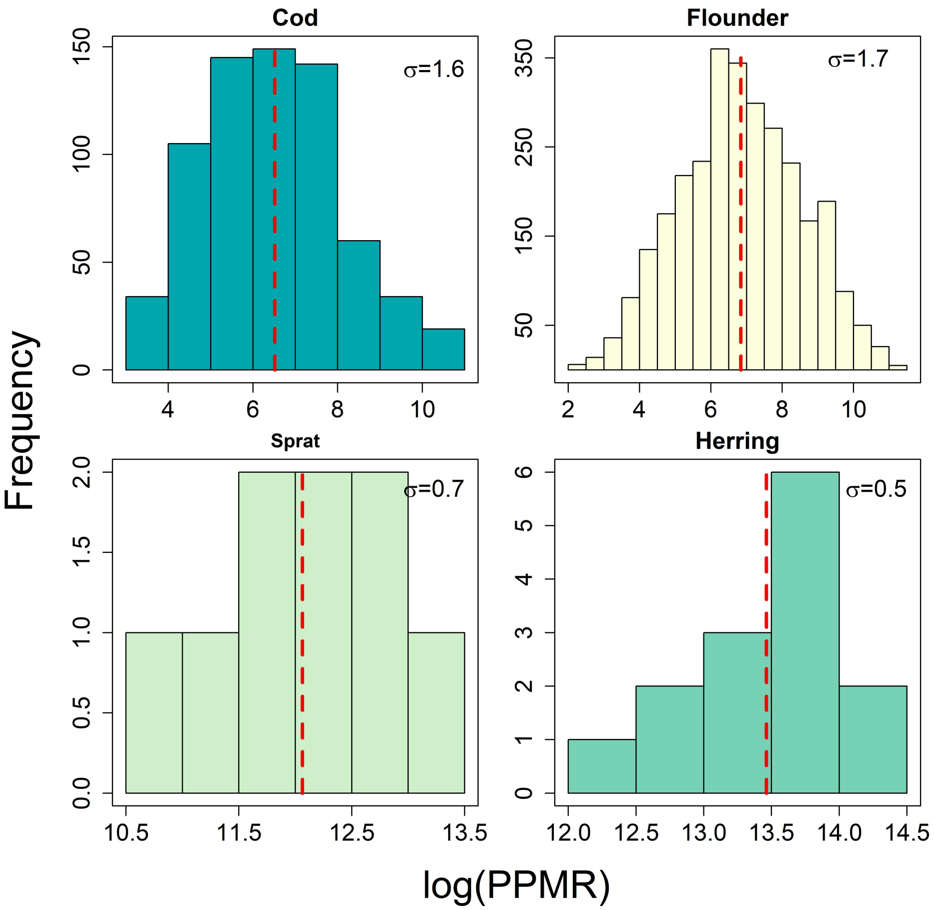


Figure S5: Frequency plots of PPMR for all four species. Vertical, red, broken lines give the mean across all predator weights. The value of $\sigma$ gives the standard deviation for each species.

overlap between each species during all years between 1991 and 2020 for which data were available, and averaged the results. This resulted in a mean, static interaction matrix. We multiplied the elements of this matrix by the vertical overlap values $\theta_{ijt}$ (equation S.2) which are fit to data extracted from (Figure 6b of Casini et al. 2019) using Web Plot Digitizer (Rohatgi 2021). This gives both vertical and horizontal overlap.

#### Appendix S3 Oxygen estimates

We extracted oxygen data for the years 1980 through 2020 from the Swedish Meteorological and Hydrological Institute (SMHI; www.smhi.se). These data arise from hydrological measurements collected during coordinated fish trawl surveys, complemented by regional and national monitoring programs throughout the Baltic Sea (Hansson and Viktorsson 2020). We used bottle data to construct spatiotemporal depth profiles of oxygen in the ICES statistical rectangles. We only included rectangles whose maximum depth fell between 50 and 120 m, and observations that fell in the months of April through November. The former restriction placed an emphasis on benthopelagic habitat suitable for cod and flounder, while the latter included the major growth and spawning months for all four species.

We used a generalized additive model (GAM) to model oxygen as a function of two-dimensional spatial coordinates, as well as water depth and year. The observed distribution of oxygen values was positive with non-zero mass at zero oxygen. Therefore, we used a tweedie distribution in the **mgcv** package (Wood et al. 2016). We modeled oxygen as a function of x-y coordinates with a thin plate spline, and depth and year with a tensor product smoother. The tensor product allows one to quantify the interaction between two terms on different scales. All terms were significant (Table S3). While the EDF of each smooth term for the default model was close to the basis dimension, or *k*, we chose not to increase this value, as we wanted to represent very broad trends over time, space, and depth. Results indicate there were higher values of oxygen in the benthic habitat in the 1980s and early 1990s than in the late 1990s through the present day (Figure S6). We averaged predicted oxygen values for each year over each ICES statistical rectangle from 10 to 40 m for pelagic habitat and from the bottom 10 m for benthic habitat. The resulting time series was fairly steady at roughly 6 mL·L$^{-1}$ in the pelagic habitat (Figure S6). Oxygen levels in the benthic habitat varied between about 1 mL·L$^{-1}$ and 3 mL·L$^{-1}$, with the rise in the 1980s reflective of a major Baltic inflow (MBI) event (Schinke and Matthäus 1998).

Table S3: Results of the generalized additive model (GAM) of oxygen as a two-dimensional smooth function of x-y coordinates, or **s(x,y)**, and a tensor product smooth of depth and year, or **te(depth,year)**. The term $\sigma_\mu$ is standard area and Ref. EDF is reference degrees of freedom. Bold and starred $p$-values indicate significant portions of the model. The adjusted $R^2$ was 0.44, with 42.3% of deviance explained.

| **Parametric Terms** | | | | |
|---|---|---|---|---|
| | Coefficient | $\sigma_\mu$ | $T$ | $p$ |
| **Intercept** | 1.74 | 0.001 | 1427 | **<.001*** |
| **Smooth Terms** | | | | |
| | EDF | Ref. EDF | $F$ | $p$ |
| **s(x,y)** | 23.87 | 24 | 1759 | **<.001*** |
| **te(depth,year)** | 28.39 | 29 | 457 | **<.001*** |

### Appendix S4 $P_{crit}$

We estimated $P_{crit}$ for each species at a given weight using the database provided by Rogers et al. (2021). This database contains several hundred measurements of $P_{crit}$ extracted from the literature, along with other experimental conditions such as water temperature and salinity, and individual attributes such as body mass. As in Rogers et al. (2016) we performed stepwise linear regression to describe $P_{crit}$ as a function of ln(body mass), salinity, temperature, and ln(resting metabolic rate, RMR). We included all measurements in the database in order to represent very general patterns arising in whole ecosystems. The stepwise regression retained all four covariates, and resulted in the following function:

$$\mathsf{P}_{\mathsf{crit}} = 30.854 + 0.027\,\mathsf{S} - 0.111\,\mathsf{T} + 0.410\ \ln(w) + 1.645\ \ln(\mathsf{RMR}) \quad \text{(S.14)}$$

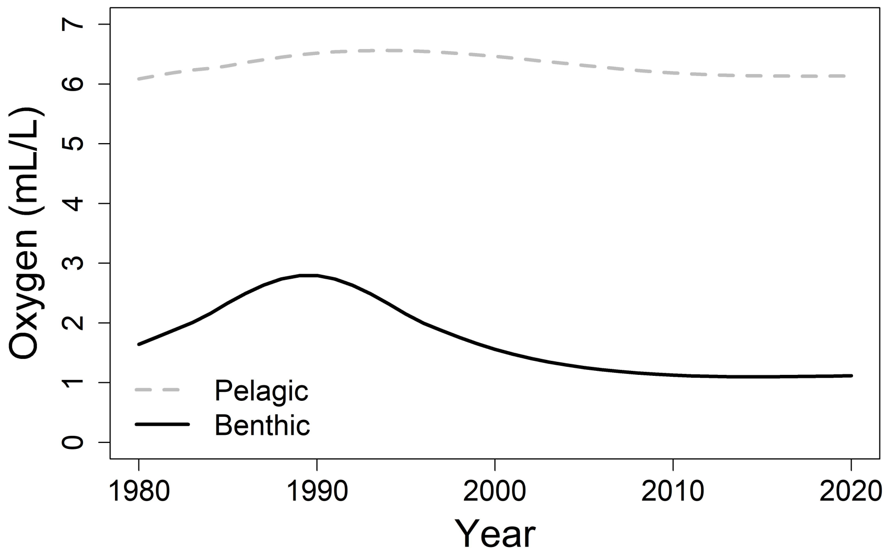


Figure S6: Predicted mean oxygen level in benthic and pelagic habitat from 1980 through 2020.

where S is salinity in practical salinity units (psu), T is temperature in degrees C, $w$ is weight in kg, and RMR is routine metabolic rate in mg $O_2 \cdot kg^{-1} \cdot h^{-1}$. The **mizer** package (Scott et al. 2014) calculates metabolic rate as an exponential function of weight with, in our case, a scaling exponent of 0.89 (Jerde et al. 2019). The result is given in units of $g \cdot year^{-1}$. We assumed this baseline value calculated by **mizer** was equivalent to routine metabolic rate (RMR) and converted to units of mg $O_2 \cdot kg^{-1} \cdot h^{-1}$ with the following formula:

$$\text{(S.15)} \qquad \mathsf{RMR}_w = \frac{1000 \cdot 1.429 \cdot 1.2905 \cdot \mathsf{RMR}_{\mathsf{mizer}}}{365.25 \cdot 24 \cdot (w/1000)}$$

In the numerator of equation S.15, 1.2905 converts from energy intake in g to L $O_2$. We assumed this biomass had a composition of 60% protein, 35% fat, and 5% cholesterol, and used these values to calculate a weighted average of L $O_2$ required for metabolism from values provided in Nelson (2004). Also in the numerator, 1.429 converts from L $O_2$ to g $O_2$, and finally 1000 converts from g to mg. In the denominator, $w/1000$ gives each size bin in kg, while 24 and 365.25 convert from years to hours. Finally, we converted the resulting $P_{crit}$ values from kPa to $mL \cdot L^{-1}$ using Henry's Law, correcting for a reference temperature of 292.15 K with the van't Hoff equation. In general, the final $P_{crit}$ values increase with weight, though the magnitude differs by species (Figure S7). This is largely dependent upon the different values of RMR, driven by differences in maximum feeding rate, driven in turn by age and weight at maturation and feeding levels (see Scott et al. 2014). Faster growth rates relative to age at maturation result in higher maximum intake rates, and ultimately in greater values of RMR at a given size. Note that the dependence of the values of $P_{crit}$ on RMR – which in the **mizer** framework is dependent on maximum consumption rate – implies that it could decrease as oxygen levels decline, given that fish appetites decline with exposure to hypoxia (Chabot and Claireaux 2008; Wang et al. 2009). While this sort of acclimation is perhaps more realistic in the long term, we chose to fix $P_{crit}$ relative to growth in order to focus on shorter term responses to environmental hypoxia, as well as to express the assumption that RMR is constant when oxygen is at or above $P_{crit}$ (Wang et al. 2009).

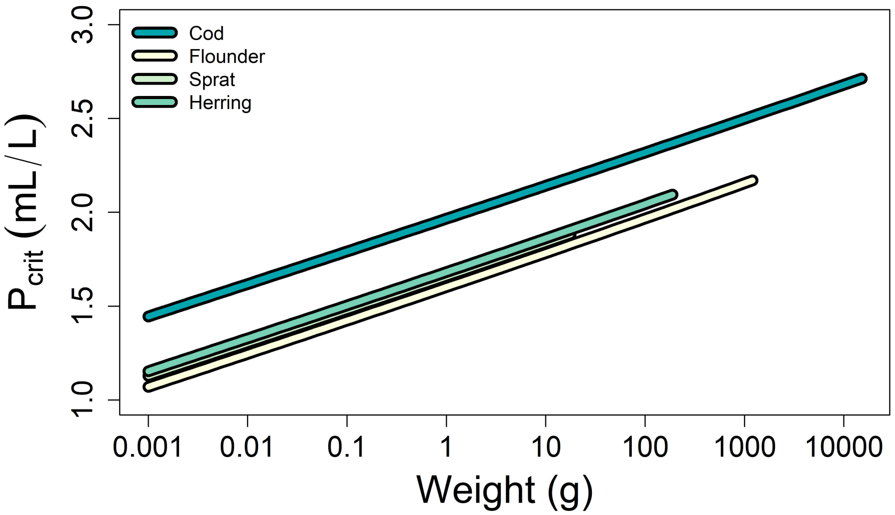


Figure S7: Calculated $P_{crit}$ as a function of size for all four species included in the model.

We calculated $P_{crit}$ for cod and flounder. At present, the $P_{crit}$ of sprat and herring is irrelevant, though we report the results all the same (Figure S7). Cod are relatively large, active, benthopelagic fish, and we thus expected to have high values of RMR to support a higher maximum metabolic rate (MMR; Killen et al. 2016). By contrast, flounder are smaller, less active, and largely benthic, and we expected smaller values of RMR. The results of the conversion reflect these expectations (Figure S7).

### Appendix S5 Calibration

#### Procedure

Each model must be calibrated before it can be used for projection and analysis. One typically calibrates to a period during which the community is relatively stable, and subsequently projects forwards or backwards in time. This process involves adjusting several of the parameters in order to match observed growth, biomass, and fisheries yield, if available. However, given the relatively complex, highly interdependent nature of the various structures within the model, adjusting a parameter for one species may result in a changed outcome for all species. There are currently no universally accepted calibration procedures for models of this type.

We chose to combine stepwise, manual parameter adjustment with automated optimization procedures, as in Lindmark et al. (2022). We treated the following data sources as observations with which to compare to model output: benthic occupancy, SSB, yield, and von Bertalanffy growth (Table S4). Note that cod were the only species that we allowed to move between the pelagic and benthic habitats. We verified vertical occupancy of cod with ratios of pelagic and benthic trawl CPUE found in Casini et al. (2019). For cod, sprat, and herring, we extracted SSB and yield from stock assessments (ICES 2020a). These data are reported for cod in ICES subdivisions 24-32, for sprat in 22-32, and for herring in 25-29 and 32. Flounder were not assessed, and we therefore assumed that the ratio of flounder and cod CPUE was equal to the ratio of flounder and cod SSB in a given year. We divided the rough estimates of flounder SSB by the mean ratio of estimated yield – the annual product of $F_{\text{bar}}$ (Draganik et al. 2007) and SSB – to observed yield during the calibration period. The procedures for estimating von Bertalanffy growth parameters are described in Appendix S2 (Table S2).

Table S4: All fixed (above) and tuning (below) parameters that may be adjusted during the course of model calibration. Species-specific parameters are marked with the subscript $i$.

| **Fixed** | | |
|---|---|---|
| Parameter | Value | Description |
| $p$ | 0.89 | Metabolic scaling exponent |
| $q$ | 0.89 | Predation or search rate scaling exponent |

| **Tuning** | | |
|---|---|---|
| Parameter | Value | Description |
| $R_{\text{max},i}$ | g | Maximum reproduction |
| $U_{\text{hab}}$ | – | Cod occupancy oxygen sensitivity |
| $a_{\text{hab}}$ | – | Cod occupancy critical oxygen adjustment |
| $U_{\text{benthic}}$ | – | Benthic resource oxygen sensitivity |
| $k_{\text{benthic}}$ | – | Benthic resource critical oxygen level |
| $U_{\text{phys},i}$ | – | Physiological rate oxygen sensitivity |
| $a_{\text{phys},i}$ | – | Physiological rate critical oxygen adjustment |
| $U_{k_s,i}$ | – | Metabolic rate oxygen sensitivity |
| $a_{k_s,i}$ | – | Metabolic rate critical oxygen adjustment |
| $z_{h0,i}$ | $y^{-1}$ | Baseline mortality due to hypoxia exposure |
| $b_i$ | – | Mortality oxygen sensitivity |
| $\gamma_i$ | volume·$y^{-1}$ | Predation or clearance rate coefficient |
| $h_i$ | g·$y^{-1}$ | Size-scaled maximum consumption rate |
| $\kappa$ | g | Benthic resource carrying capacity |
| $z_0$ | $g^{1-p}$·$y^{-1}$ | Background mortality coefficient |
| $\varepsilon_{\text{repro}}$ | – | Reproduction efficiency or survival of recruits |

Note that benthic and pelagic oxygen (see Section S2) and F served as data input. For steps 1–11 we used average oxygen and F during the calibration period, and for steps 12–13 we used time series. The code provided includes all details of implementation (found here). The steps are summarized below.

1. Choose values for carrying capacity $\kappa$ and scaling parameters of benthos and plankton. These values should be consistent with the scale of the observed yield and SSB. We used total biomass, but one may also use biomass per area or volume. Note that we estimated parameters using data from ICES subdivisions 25 through 29, while some stock assessment estimates of SSB and F cover subdivisions 22 through 32, and subsets therein. The implicit assumption is that all fish reside in the Central Baltic Sea. The majority of eastern Baltic cod are indeed concentrated primarily in the Central Baltic Sea, while sprat, herring, and flounder extend into the Gulf of Finland (ICES subdivision 32; ICES 2020a). For

sprat and herring, the larger area can be accommodated by using a larger $\kappa_{\text{pelagic}}$, reflecting greater overall food availability. The relative distribution of flounder biomass is unknown, and therefore we lumped these species together in a single benthic habitat with cod. Note that the values of $\kappa$ can have a major impact on the realized diets of the fish. Higher values generally force greater reliance on the background resource while lower values are reflective of a diet composed of explicitly modeled fish species. Therefore, while adjusting, check diets. Scaling parameters may arise from literature regarding the tolerance of major prey organisms in each habitat to hypoxia. At this stage, only reasonable values are required, as they are fit to time series data in another step.

2. Choose starting values for each species' maximum reproduction rates $R_{\text{max},i}$ such that model SSB for each species is within an order of magnitude of observed SSB. These are affected by the choice of $\kappa$. In practice, we have found that several combinations of $R_{\text{max}}$ and $\kappa$ and scaling values may result in matching values of SSB. This is also affected by growth, but these parameters are calibrated in a later step. Therefore, at this stage, we recommend focusing on the diet to narrow the range of possible values.

3. Choose reasonable starting values for oxygen dependence of physiological rates of assimilation, feeding, and reproduction efficiency (equation S.1). Generally, reasonable values of $U_{\text{phys},i}$ fall below ten, and will never fall below zero. Values of $a_{\text{phys},i} > 1$ shift the oxygen level at which physiological scaling is $0.5$ to the right, while $a_{\text{phys},i} < 1$ shift this value to the left. Negative values reflect a decrease in oxygen sensitivity with weight, while positive values reflect an increase. We allowed both positive and negative values as studies differ in their claims regarding the relationship of hypoxia tolerance and weight (e.g. Nilsson and Östlund-Nilsson 2008; Pan et al. 2016). Declines in performance begin well above $P_{\text{crit},iw}$ (Hrycik et al. 2017). Therefore, $U_{\text{phys},i}$ will often be low, perhaps three or less, depending on the value of $a_{\text{phys},i}$. These parameters may best be characterized relative to other species. For example, to represent our community – and more generally a community in which one species is the superior competitor in an hypoxic environment – we chose a lower value of $U_{\text{phys},i}$ and a higher value of $a_{\text{phys},i}$ for cod relative to flounder. Check that modeled biomass is still within an order of magnitude of observed biomass. Prioritize SSB and coexistence over specific values of $U_{\text{phys},i}$ and $a_{\text{phys},i}$. Note that these values are calibrated using time series data in a later step.

4. Choose reasonable starting values for metabolic scaling (equation S.4). Values of $a_{\text{met},i}$ shift the point of doubling left or right, as described in the previous step. We also allowed both positive and negative values of $a_{\text{met},i}$. In this model, metabolic costs are also a function of maximum consumption and assimilation efficiency, both of which decline with oxygen according to the physiological scaling function described in the previous step. Therefore, overall changes in metabolic costs are a complex function of oxygen dependence. We again chose values to characterize species relative to one another, with higher values of $U_{\text{met},i}$ and lower values of $a_{\text{met},i}$ for cod than for flounder. This also reflects a greater sensitivity to hypoxic conditions in the former than in the latter. Prioritize SSB and coexistence over specific values of $U_{\text{met},i}$ and $a_{\text{phys},i}$. Note that these values are also calibrated using time series data in a later step. Repeat steps 1 through 4 until modeled SSB is within one order of magnitude of observed SSB.

5. Check density dependence in reproduction for each species. One can check relative controls on each species by evaluating the ratio of density independent to density dependent recruitment. If these values are very high, it indicates that the given population is largely affected by density dependence rather than predation or competition with other species. If they are very low, the population may be immune to sources of mortality and competition. In general, larger species should experience a higher degree of density dependence. Recall that increasing density dependence within a species can result in saturating yield with increasing F (Beverton and Holt 2012). Therefore, observing this kind of relationship of yield with F in modeled species is not necessarily cause for alarm. If time series of yield at F appear insensitive to F, then consider returning to this step and changing the degree of density dependence.

6. Choose starting values for the background mortality coefficients and the clearance rate coefficients for each species. Check coexistence, feeding level, growth, SSB, and yield at F. The latter three are compared to observations. Feeding level emerges from realized diets, and is contained in the interval $[0, 1]$, with $0$ being unfed and $1$ being completely satiated. Emergent values should be similar among species and above $0.2$, which is the minimum required for maintenance. Increasing clearance rate coefficients increases both growth rates and feeding level relative to satiation. It may also be necessary to adjust maximum consumption in tandem with clearance rates. Increasing maximum consumption rate ($h$) increases growth rates, but it decreases feeding level. At this stage, prioritize fits to observed yield with precise values of the background mortality coefficients.

7. This and all subsequent steps use the **optimParallel** package. This package applies the "L-BFGS-B" method, a quasi-Newton method with box constraints (Byrd et al. 1995), in parallel to minimize a user defined function. If a given optimization procedure failed to converge, we increased the control argument `factr` by factors of ten until convergence was achieved. If this failed, we increased control argument `pgtol` by factors of ten until convergence was achieved. This only occurred in one instance, and essentially indicated that our solution space was very flat. That is, optimization was superfluous. Using this procedure, we first fit habitat sensitivity parameters to observed benthic occupancy of relevant species – cod, in this case. Recall that occupancy scales with equation S.1. For occupancy, we did not allow $a_i$ to fall below zero, suggesting larger fish are more likely to escape to the pelagic habitat at a given oxygen level.

8. Fit to observed mean yield of al species during the calibration period. The objective function adjusts the background mortality coefficients, then calculates and returns the sum of squared errors between the natural logarithms of modeled yield at F and mean observed yield during the calibration period.

9. Fit to observed mean SSB and growth during the calibration period for all species. The objective function adjusts the base ten logarithm of each species' maximum reproductive rates $R_{\mathsf{max},i}$ and clearance rates $\gamma_i$. It calculates sum of squared errors between modeled biomass of mature size classes and observed SSB. The function also predicts the weight as a percentage of $W_\infty$ at fifty evenly spaced ages beginning at zero and ending at the maximum observed age for each species. It calculates the sum of squared errors between

modeled growth and von Bertalanffy growth estimates arising from the procedures in Section S2. The function returns a weighted sum of the the error in SSB and in growth. The weights are dependent on the scale of the errors of SSB and growth. We found that errors in growth were typically two orders of magnitude greater than errors in SSB, and adjusted the former by a factor of 0.01. Adjust starting values for clearance rates to roughly match von Bertalanffy growth estimates, if necessary. Note that the order of steps 7 through 9 are in the reverse order of priority or data trustworthiness. That is, we prioritize fits to growth and SSB above fits to yield and fits to yield above fits to occupancy. The processes in this model are highly interdependent, and fitting one set of parameters to a subset of the data can negate the fit of another set of parameters to another subset of the data. Thus, order reflects priority.

10. Fit to annual growth patterns during the calibration period by adjusting the physiological and scaling parameters. This includes scaling of assimilation, feeding, and reproductive efficiency in addition to scaling of metabolic costs. The objective function adjusts $U_i$, $a_i$, $U_{\mathsf{met},i}$, and $a_{\mathsf{phys},i}$ (see equations S.1 and S.4). This procedure calculates the error in growth as described in step 9, and is performed once for each species that experiences any level of exposure to the benthic environment i.e. for cod and flounder.

11. Fit to annual growth patterns during the calibration period by adjusting benthic resource carrying capacity scaling parameters. Recall that this scaling function is written as:

$$\xi_{\kappa_{\mathsf{benthic}}}(O_{\mathsf{benthic}}) = \frac{1}{1 + e^{-U_{\mathsf{benthic}}(O_{\mathsf{benthic}} - k_{\mathsf{benthic}})}} \tag{S.16}$$

where $O_{\mathsf{benthic}}$ references the oxygen level in the benthic environment only. The objective function adjusts $U_{\mathsf{benthic}}$ and $k_{\mathsf{benthic}}$, and calculates the error in growth of all species.

12. Fit to annual patterns of yield during the calibration period by adjusting the parameters describing additional mortality due to hypoxia exposure. Recall that this mortality is expressed by the hazard function in equation S.3 (see main text), again repeated here:

$$h(t) = b_{0,i} e^{b_i O_{iwt}} \tag{S.17}$$

The objective function adjusts $b_{0,i}$ and $b_i$ and predicts yield at observed values of fishing mortality (F). It then calculates the sum of squared differences between the natural logarithm of these values and observed yield. We perform this step once for each species that experiences any level of exposure to the benthic environment i.e. for cod and flounder.

The procedure described above is intended for calibration of the full model, which includes oxygen dependence of all four processes: benthic resource carrying capacity (B), avoidance via changes in occupancy (O), mortality due to exposure to hypoxic waters (M), and physiological scaling (P; the so-called BOMP model). We repeated calibration for all possible combinations of oxygen dependence. We also calibrated a model with no oxygen dependence. This resulted in 16 total model calibrations. For those processes which were not included in a given model, we turned off the associated scaling function by setting the scaling coefficient to one for all species at all sizes, or by setting the result to zero for all species at all sizes in the case of mortality. We would then skip the associated step. However, all steps were still performed in the order listed above, and with the same checks, if applicable.

#### Error calculation

We used the calibrated models to project forward in time from 2001 through 2019 and calculated errors in modeled quantities during both the calibration, or in-sample, period (1991-2000) and the projection, or out-of-sample, period (2001-2019). Only the out-of-sample or projection errors were used for model selection, as this is generally more robust (Cooke et al. 2014). We calculated a weighted sum of squared errors between the natural logarithms of modeled biomass of mature size classes and observed SSB for all species:

$$\varepsilon_{\mathsf{SSB}} = \sum_{i=1}^{4}\sum_{j=1}^{19} w_{\mathsf{SSB},i}(\ln \mathsf{SSB}_{\mathsf{mod},ij} - \ln \mathsf{SSB}_{\mathsf{obs},ij})^2 \tag{S.18}$$

where $w_{\mathsf{SSB},i} = \{0.3, 0.1, 0.3, 0.3\}$ for cod, flounder, sprat, and herring, in that order. The weights reflected our confidence in each data source. The relatively low confidence in flounder data reflects the ad-hoc method by which we estimated values for flounder SSB. We used the same method to calculate errors between the natural logarithms of modeled yield at F and observed yield:

$$\varepsilon_{\mathsf{Yield}} = \sum_{i=1}^{4}\sum_{j=1}^{19} w_{\mathsf{Yield},i}(\ln \mathsf{Yield}_{\mathsf{mod},ij} - \ln \mathsf{yield}_{\mathsf{obs},ij})^2 \tag{S.19}$$

where $w_{\mathsf{Yield},i} = \{0.25, 0.25, 0.25, 0.25\}$ for cod, flounder, sprat, and herring, in that order. Lastly, we calculated annual error in size at age by predicting the weight as a percentage of $W_\infty$ at fifty evenly spaced ages beginning at zero and ending at the maximum observed age for each species and comparing to annual von Bertalanffy estimates:

$$\varepsilon_{\mathsf{Growth}} = \sum_{i=1}^{4}\sum_{j=1}^{19}\sum_{k=1}^{50} w_{\mathsf{Growth},i}(\mathsf{Size}_{\mathsf{mod},ijk} - \mathsf{Size}_{\mathsf{obs},ijk})^2 \tag{S.20}$$

where $w_{\mathsf{Yield},i} = \{0.25, 0.25, 0.25, 0.25\}$ for cod, flounder, sprat, and herring, in that order.

We repeated these calculations for each of the 16 calibrated models to arrive at a set of errors: $\varepsilon_{\mathsf{SSB}} = \{\varepsilon_{\mathsf{SSB},1}, ..., \varepsilon_{\mathsf{SSB},16}\}$. Similarly, we have $\varepsilon_{\mathsf{Yield}}$ and $\varepsilon_{\mathsf{Growth}}$, containing errors in yield and growth, respectively, for all 16 models. We normalized each set of errors to span across the interval $[0, 1]$ with the standard normalization function:

$$\varepsilon_{\mathsf{norm}} = \frac{\varepsilon - \min(\varepsilon)}{\max(\varepsilon) - \min(\varepsilon)} \tag{S.21}$$

This ensured that each set of errors were compared on the same scale, and allowed us to arrive at a final ranking for each model by computing a weighted sum of normalized errors:

$$\varepsilon_{\mathsf{Final}} = 0.3\,\varepsilon_{\mathsf{SSB}} + 0.1\,\varepsilon_{\mathsf{Yield}} + 0.6\,\varepsilon_{\mathsf{Growth}} \tag{S.22}$$

These weights reflect our priority in fitting to each set of observations, which in turn reflects our degree of confidence in the data. The model with the lowest value of $\varepsilon_{\mathsf{Final}}$ was treated as the best model overall.

#### Results of model calibration

Estimated scaling parameters reflected both the nature of multispecies size spectrum models and differences in sensitivity of cod and flounder (Table S5 and Figure S8). For example, estimated benthic resource scaling parameters ($U_{\mathsf{res}}$ and $k_{\mathsf{res}}$) were nearly identical across models, and did not stray far from initial values of 2 and 0.8, respectively. However, these scaling values also affect initialization of the **mizer** model itself. That is, in choosing values for resource scaling, initial values for abundance across size classes are adjusted. This is true for all scaling parameters, and thus values may not reflect *actual* oxygen sensitivity of each process, but rather an amalgam of internal **mizer** setup and oxygen sensitivity.

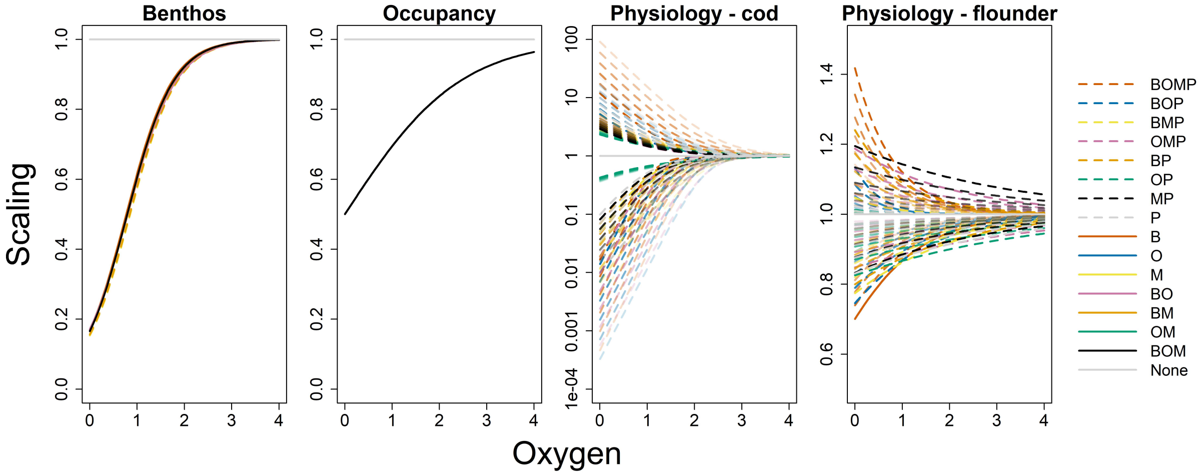


Figure S8: Scaling of benthic resource carrying capacity (Benthos), benthic occupancy of cod (Occupancy), and physiological rates of cod and flounder for all 16 models. For the latter two plots, lines above 1 are up-scaling of metabolic costs while lines below 1 are down-scaling of assimilation, feeding, and reproduction efficiency. Note the vast difference in scale between cod and flounder. Opacity of line coloration declines as weight increases..

Cod benthic occupancy parameters were identical across models, and suggested occupancy was identical across all sizes (Table S5, $a_{\mathsf{hab}}$ and Figure). Given the data were interpreted as single probabilities of benthic occupancy for each year, it is unsurprising that there was no differentiation among sizes, and thus no differentiation among models.

Table S5: All estimated scaling parameters for all 16 models for cod and flounder. The subscript res refers to benthic resource scaling and is identical within a given model. The subscript hab refers to benthic occupancy scaling; crit refers to consumption, assimilation efficiency, and reproduction efficiency scaling; and met refers to metabolic costs scaling. The mortality parameters $b_0$ and $b_i$ are drawn from the hazard function given by equation S.3.

| Species | Model | $U_{\mathrm{res}}$ | $k_{\mathrm{res}}$ | $U_{\mathrm{hab}}$ | $a_{\mathrm{hab}}$ | $U_{\mathrm{crit}}$ | $a_{\mathrm{crit}}$ | $U_{\mathrm{met}}$ | $a_{\mathrm{met}}$ | $b_0$ | $b_i$ |
|---|---|---|---|---|---|---|---|---|---|---|---|
| Cod | BOMP | 2.00 | 0.79 | 0.82 | 0.00 | 3.82 | 0.74 | 1.84 | 0.90 | 0.00 | 1.83 |
| | BOM | 2.04 | 0.79 | 0.82 | 0.00 | – | – | – | – | 0.00 | 1.98 |
| | BOP | 2.00 | 0.80 | 0.82 | 0.00 | 3.58 | 0.82 | 1.64 | 0.61 | – | – |
| | BMP | 2.00 | 0.85 | – | – | 2.62 | 0.77 | 1.09 | 0.36 | 0.00 | 1.92 |
| | OMP | – | – | 0.82 | 0.00 | 3.34 | 0.82 | 1.49 | 0.33 | 0.00 | 1.79 |
| | BO | 1.97 | 0.80 | 0.82 | 0.00 | – | – | – | – | – | – |
| | BM | 2.03 | 0.80 | – | – | – | – | – | – | 0.00 | 1.97 |
| | BP | 2.00 | 0.84 | – | – | 2.44 | 0.74 | 1.34 | 0.42 | – | – |
| | OM | – | – | 0.82 | 0.00 | – | – | – | – | 0.00 | 1.98 |
| | OP | – | – | 0.82 | 0.00 | 1.00 | 0.21 | 1.00 | 0.20 | – | – |
| | MP | – | – | – | – | 2.29 | 0.73 | 1.40 | 0.32 | 0.00 | 1.87 |
| | B | 2.05 | 0.77 | – | – | – | – | – | – | – | – |
| | O | – | – | 0.82 | 0.00 | – | – | – | – | – | – |
| | M | – | – | – | – | – | – | – | – | 0.00 | 1.98 |
| | P | – | – | – | – | 2.26 | 0.68 | 1.60 | 0.71 | – | – |
| | None | – | – | – | – | – | – | – | – | – | – |
| Flounder | BOMP | 2.00 | 0.79 | – | – | 1.03 | -0.77 | 1.25 | -0.65 | 0.00 | 2.14 |
| | BOM | 2.04 | 0.79 | – | – | – | – | – | – | 0.00 | 2.11 |
| | BOP | 2.00 | 0.80 | – | – | 0.82 | -1.23 | 1.98 | -0.95 | – | – |
| | BMP | 2.00 | 0.85 | – | – | 0.60 | -1.92 | 1.40 | -2.04 | 0.00 | 2.15 |
| | OMP | – | – | – | – | 0.33 | -4.93 | 0.48 | -3.23 | 0.00 | 2.15 |
| | BO | 1.97 | 0.80 | – | – | – | – | – | – | – | – |
| | BM | 2.03 | 0.80 | – | – | – | – | – | – | 0.00 | 2.12 |
| | BP | 2.00 | 0.84 | – | – | 0.56 | -2.29 | 0.98 | -1.36 | – | – |
| | OM | – | – | – | – | – | – | – | – | 0.00 | 2.12 |
| | OP | – | – | – | – | 0.32 | -4.50 | 1.62 | -2.50 | – | – |
| | MP | – | – | – | – | 0.44 | -3.42 | 0.31 | -4.91 | 0.00 | 2.14 |
| | B | 2.05 | 0.77 | – | – | – | – | – | – | – | – |
| | O | – | – | – | – | – | – | – | – | – | – |
| | M | – | – | – | – | – | – | – | – | 0.00 | 2.11 |
| | P | – | – | – | – | 0.39 | -3.85 | 1.37 | -2.56 | – | – |
| | None | – | – | – | – | – | – | – | – | – | – |

Oxygen dependence of natural mortality was always estimated to be zero, no matter the model (Table S5). However, it is important to note that the model generally underestimated yield of cod and flounder during the calibration period (see Figure 4 in the main text), and the inclusion of additional mortality during model setup caused some initial values to be up-scaled, thereby increasing yield overall, and decreasing error. Given natural mortality parameters were fit to yield, models including these components generally performed better than those which did not, but not for any interesting mechanistic reason.

Physiological scaling parameters varied by species and by model. Generally speaking, cod exhibited greater sensitivity to declining oxygen than flounder, in accordance with expectations. Smaller cod were also consistently less sensitive to low oxygen than larger cod. Conversely, smaller flounder were more sensitive to low oxygen than larger flounder (Figure S8). Furthermore, most models suggested that increases in metabolic costs began at similar oxygen levels to those at which other physiological processes began to decline. This was true for both cod and

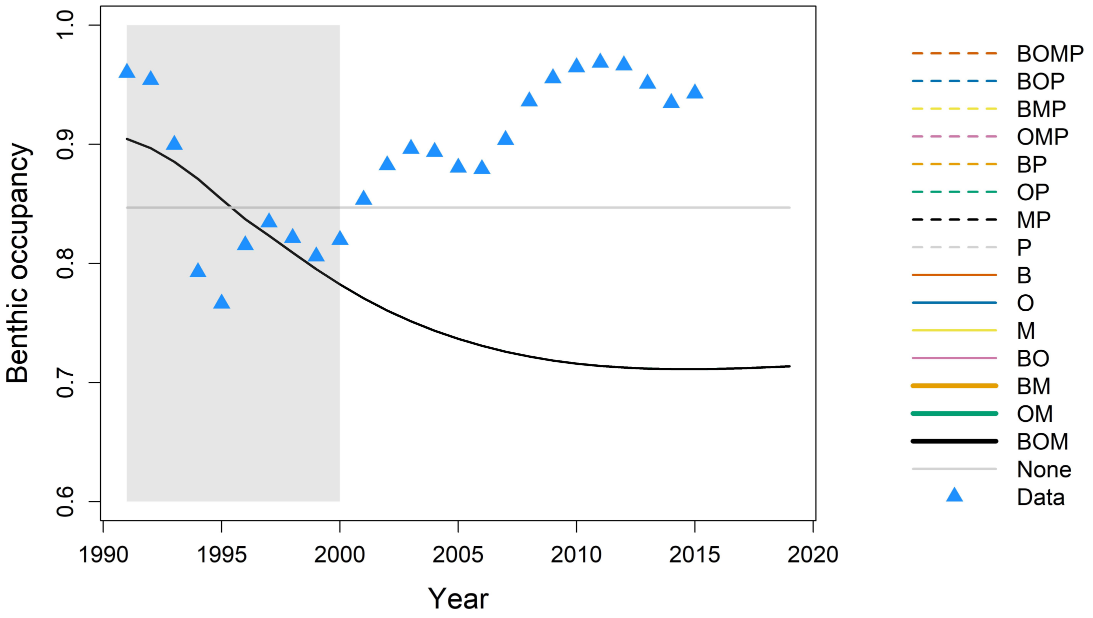


Figure S9: Results of fitting occupancy parameters for each model to observations from Casini et al. (2019). The shaded region indicated the calibration period. Grayscale lines are models with various combinations of B/O/M/P, with lighter shades indicating the model has less variables. Lines with zero slope indicate that O was not included in the model.

flounder, evident in relatively similar values of $a_{\text{crit}}$ and $a_{\text{met}}$. That said, values of $a_{\text{met}}$ were generally smaller than values of $a_{\text{crit}}$ for cod, and less negative for flounder, indicating a fairly consistent characterization. This likely compensated for the exponential nature of the curve, which results in steep increases in metabolic costs at zero oxygen, for example, with modest increases in $a_{\text{met}}$.

Recall that we only used errors during the projection period for model selection. These errors in fits to SSB, yield, and growth during both the calibration and the projection period suggest that there are complex interactions between model components. First, note that errors in cod and flounder model output experienced the largest change in magnitude across models in the projection period (Figure S10). This was unsurprising, given that they experienced direct consequences from exposure to hypoxic waters. Of the two, errors in cod output spanned a wider range of values (Figure S10).

Second, errors in SSB during the projection period were clearly exacerbated by models which included physiological scaling without occupancy scaling (Figure S10). That is, physiological rates are oxygen dependent, it is essential that cod be allowed to escape the environment. This was also true of errors in yield. For growth, those models which include physiological scaling tend to perform better when occupancy scaling is also present, though it is not entirely consistent.

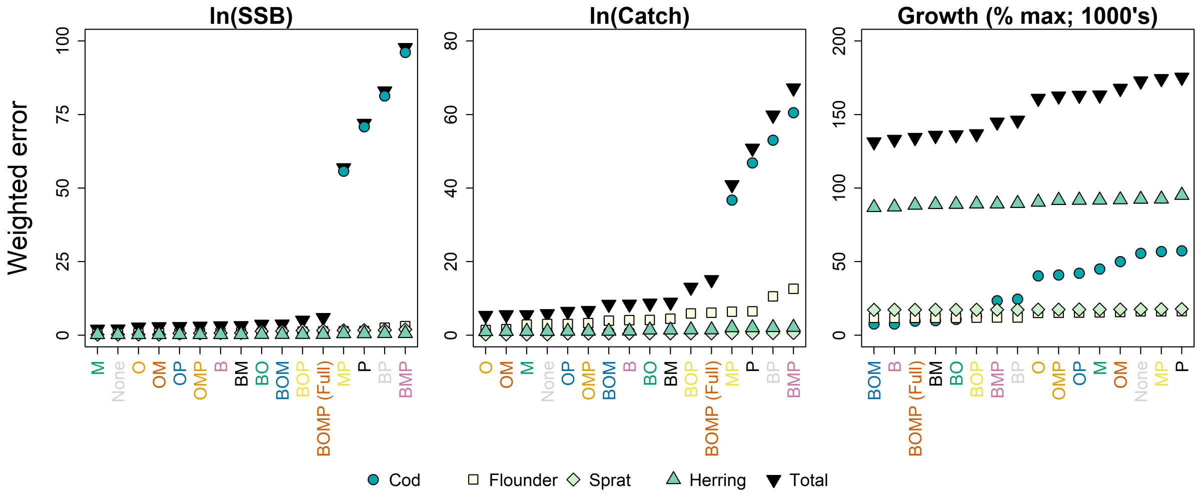


Figure S10: Weighted error during the projection period (2001–2019) in ln(SSB), ln(yield), and size-at-age or growth as a percentage of $W_\infty$ (see Table S2). Black, inverted triangles are sum totals, other filled symbols correspond to each species. Models are ordered from best to worst for each component.

We relied upon summarized rankings during the projection period (2001–2019) to provide the best model as a weighted compromise among model components. Final weighted rankings (equation S.22) largely follow patterns evident in growth, given the higher weight based on observations of size-at-age (Figure S11). In general, models including benthic resource scaling performed significantly better than those without. Though, again, models which include physiological scaling must also include occupancy scaling lest they be overwhelmed by error in SSB and yield (Figure S13). Model choice through calibration and projection may best be performed as a check on those components which must appear together, or which introduce egregious error.

We did not rely upon errors during the calibration period (1991–2000) for model selection. Still, we note that good performance during the calibration period did not translate to good performance during the projection period (Figure S12). This could possibly be due to a change in the mechanisms driving responses to hypoxia during over time. Testing this would require extensive re-calibration and hindcasting, which we leave for future studies.

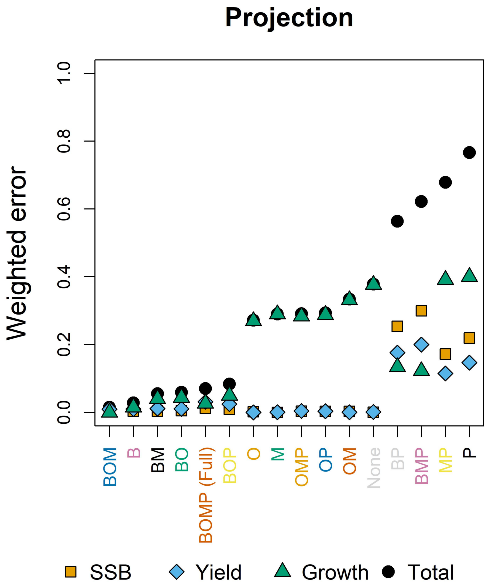


Figure S11: Final weighted error rankings during the projection period (2001–2019). Black filled circles are final rankings, while other filled symbols give weighted rankings of other model components. Models are ordered from best to worst.

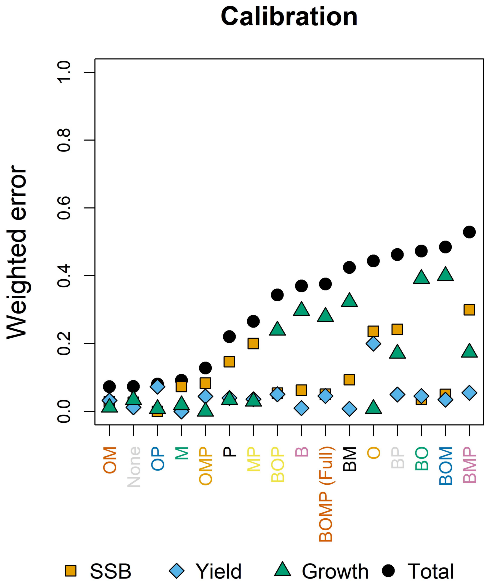


Figure S12: Final weighted error rankings during the calibration period (1991–2000). Symbols and order are given as in Figure S11.

### Appendix S6 Sensitivity analysis

We repeated the calibration process detailed in Appendix S5 for all 16 models three times to test the sensitivity of our results to parameter starting values. Each time, we drew new starting values for all oxygen dependence parameters from a uniform distribution with a minimum and maximum 20% below and 20% above the original starting value, respectively. We set three separate random seeds for these starting values so that results are repeatable, and so that the same set of starting values could be used to calibrate all 16 models within each repetition. While three repetitions gives a very small sample size, the calibration procedure is both extensive and labor intensive, which limits the number of times it can be repeated within a reasonable amount of time.

Results of this procedure indicate that while model rankings are sensitive to starting values, benthic food availability and habitat use consistently play vital roles in impacts of hypoxia on fish communities. Models with the lowest error during the projection period (2001–2019) always include oxygen dependence of benthic food resources (Figure S13). The worst models always include oxygen dependence of physiological rates without an oxygen replete refuge. These mechanisms, above all others, determine the ability of the models to explain observed trends in spawning stock biomass (SSB), commercial yield, and somatic growth.

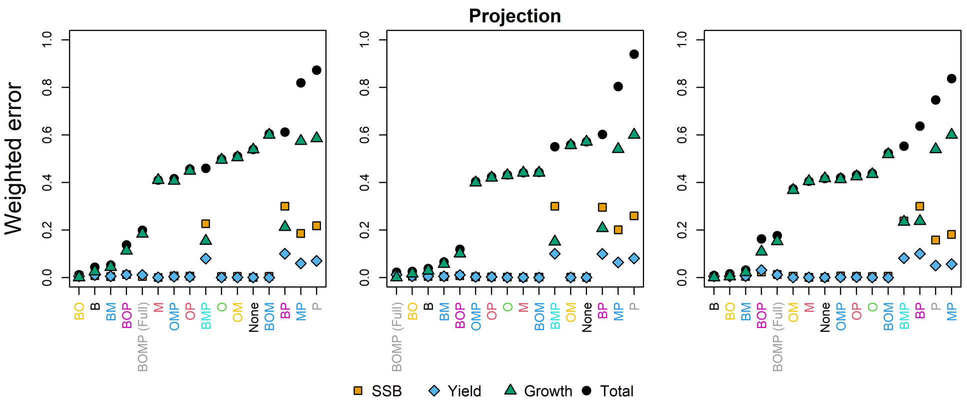


Figure S13: Final weighted error rankings during the projection period (2001–2019) for all three model calibration repetitions completed for the sensitivity analysis. Symbols and model order are given as in Figure S11.

Error during the calibration period follows a similar pattern, with the same sets of models clustering among those which are best or worst at explaining observed patterns (Figure S14). Thus, our general conclusions given in the main text are still supported. We only caution against allowing single runs to serve as definitive evidence in favor of some mechanisms and to the exclusion of others. Rather, it may be best to focus on a collection of models which consistently appear among the highest ranked models during sensitivity analyses, or rather to focus on a collection of models with similarly high rankings.

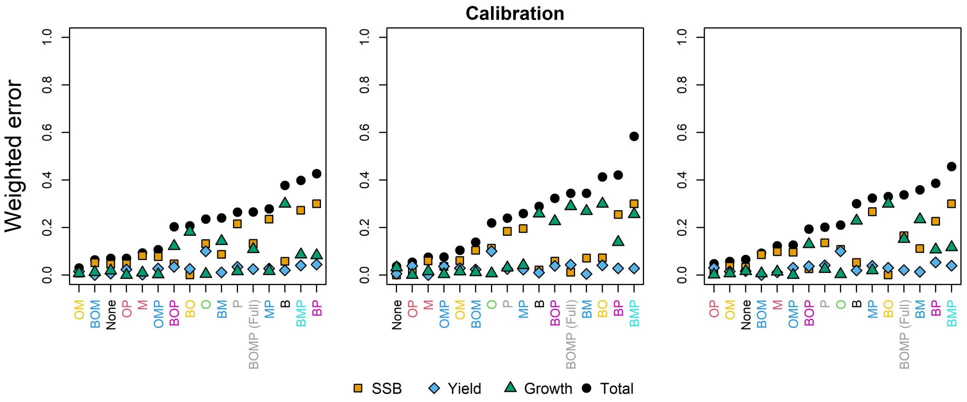


Figure S14: Final weighted error rankings during the calibration period (1991–2000) for all three model calibration repetitions completed for the sensitivity analysis. Symbols and model order are given as in Figure S11.

### Appendix S7 Tables and Figures

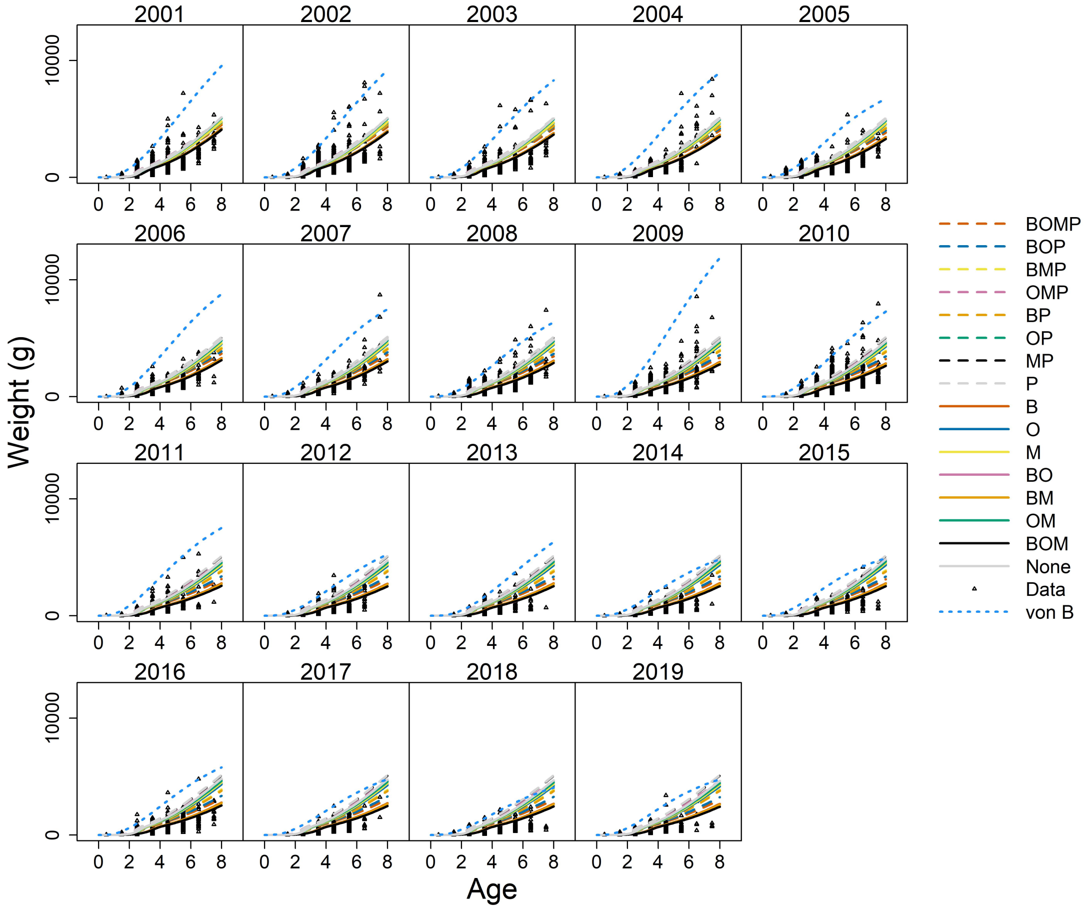


Figure S15: Projections of body growth for cod during the projection period (2001–2019) arising from all 16 models. Data are included as small, black triangles. Dotted, light blue lines are von Bertalanffy growth estimates from each year.

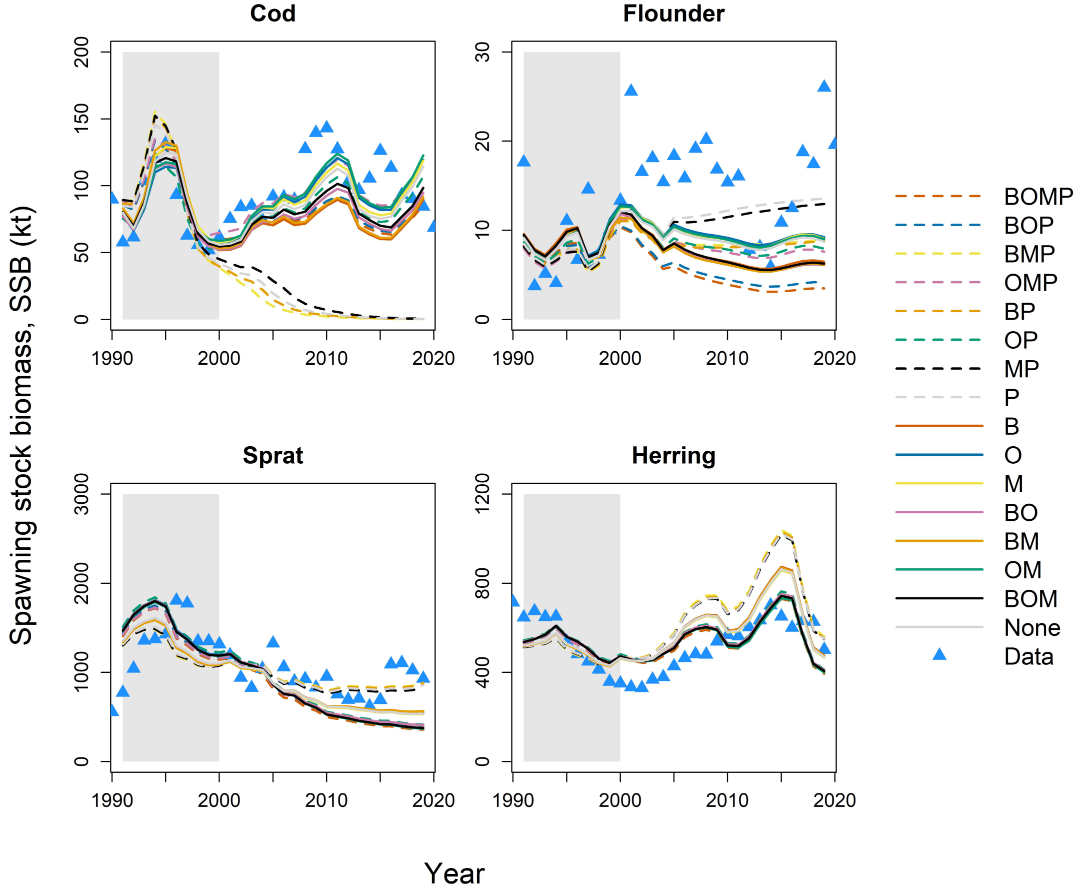


Figure S16: Projections of biomass of mature size classes (SSB) throughout the calibration period (1991–2000; shaded region) and the projection period (2001–2019) using all 16 versions of the oxygen-dependent food web model. Blue triangles are observations (as estimated from stock assessments; see Methods of main text).

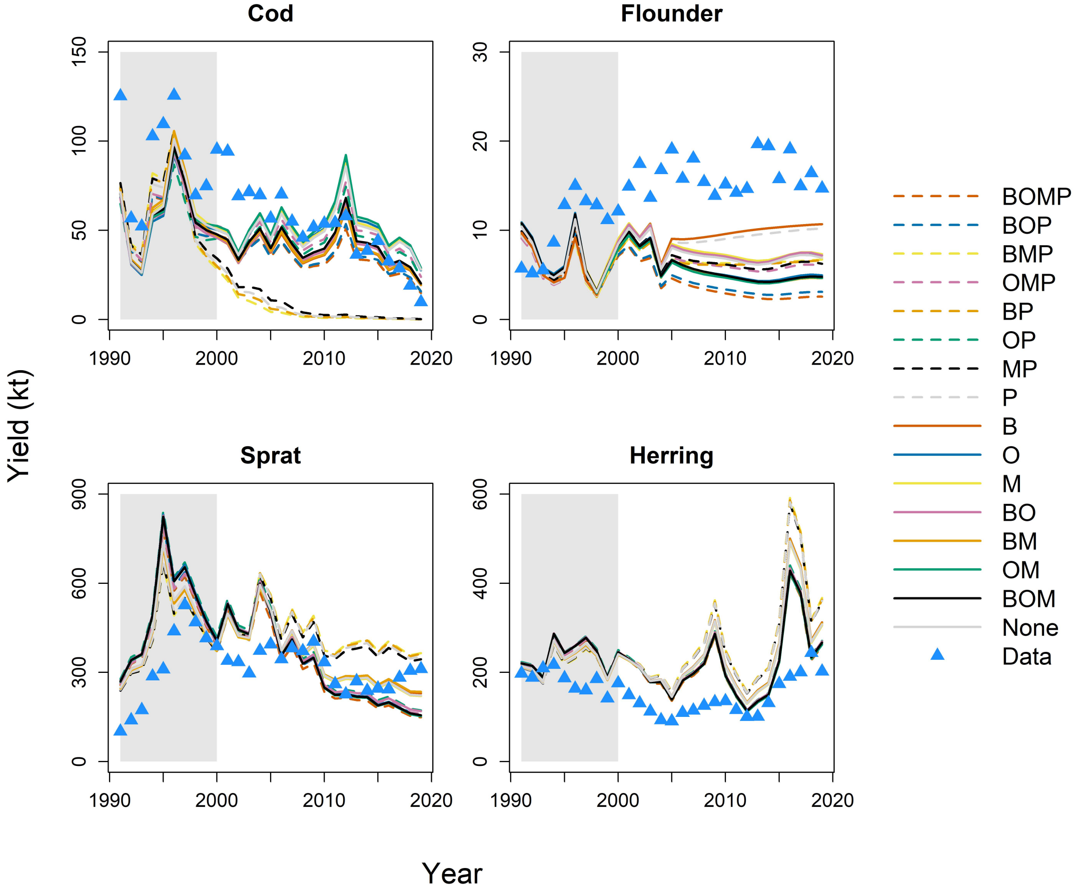


Figure S17: Projections of yield at observed fishing mortality, throughout the calibration period (1991–2000; shaded region) and the projection period (2001–2019), using all 16 versions of the oxygen-dependent food web model. Symbols as in Figure S16.

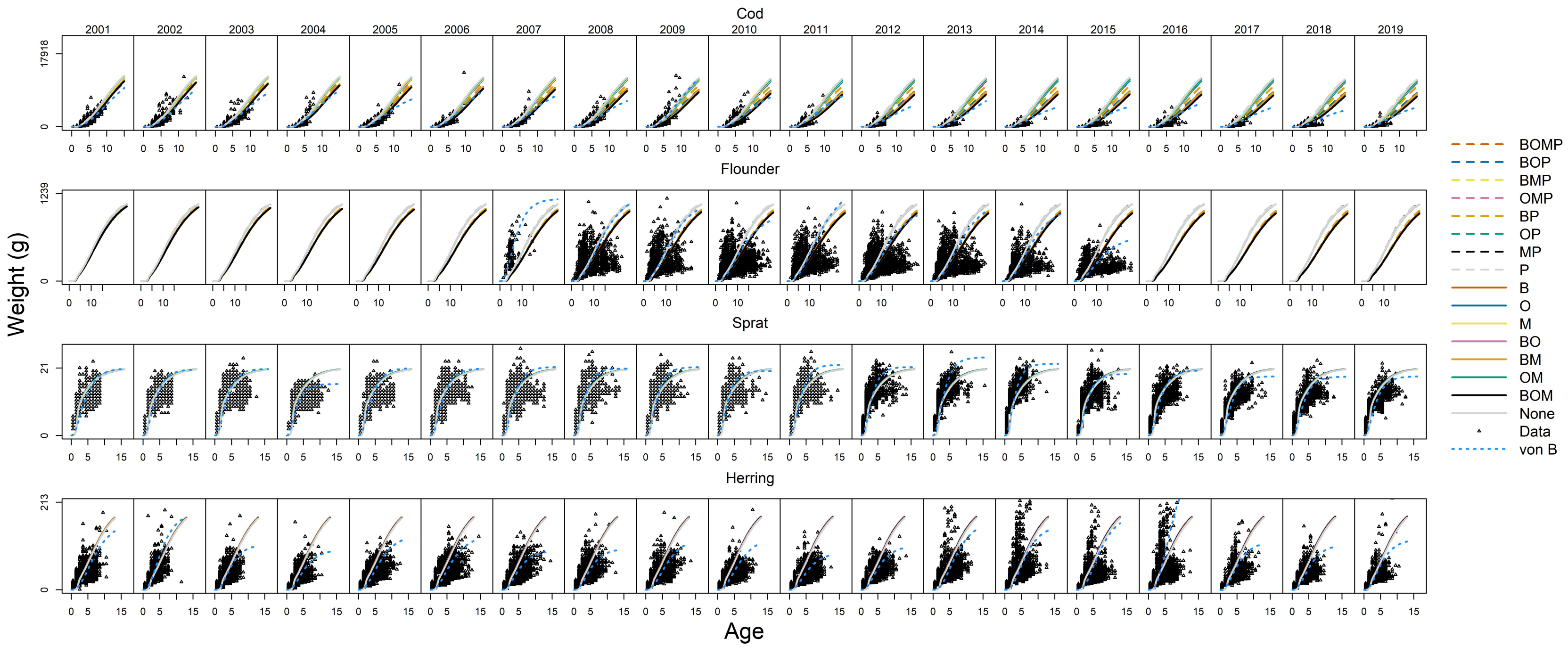


Figure S18: Projections of growth for each species throughout the projection period (2001–2019). Dotted blue lines are predictions from annual estimates of von Bertalanffy growth parameters.

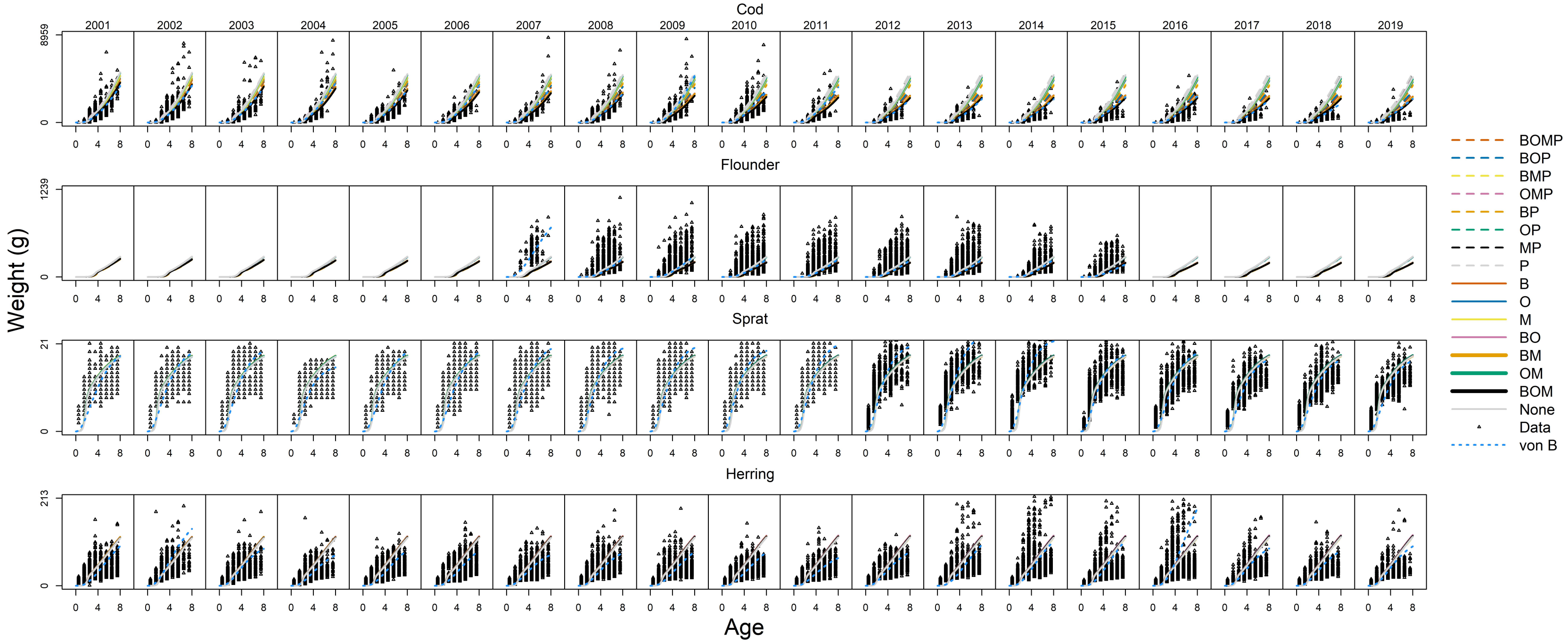


Figure S19: Projections of somatic growth (size-at-age) during the first eight years of life for each species throughout the calibration period (1991–2000) and the projection period (2001–2019), using the oxygen-dependent food web model. Lines and symbols given as in Figure S18.

Table S6: Modeled SSB (kt), yield (kt) at mean fishing mortality observed during the calibration period (1991–2000), and size (g) of individuals at maximum observed age for all species under three oxygen scenarios applied in all 16 models.

| | | Cod | | | Flounder | | | Sprat | | | Herring | | |
|---|---|---|---|---|---|---|---|---|---|---|---|---|---|
| Model | $O_2$ | SSB | Yield | Size | SSB | Yield | Size | SSB | Yield | Size | SSB | Yield | Size |
| BOMP | 3 | 87.3 | 62.1 | 14266 | 9.2 | 7.4 | 1125 | 1688.9 | 519.5 | 20 | 570.1 | 243.6 | 172 |
| | 2 | 82.2 | 51.2 | 9746 | 8.2 | 6.5 | 1087 | 1385.8 | 426.6 | 20 | 526.8 | 225.4 | 174 |
| | 1 | 64.3 | 31.8 | 4286 | 2.7 | 2.0 | 795 | 811.1 | 250.2 | 21 | 438.5 | 188.2 | 176 |
| BOM | 3 | 75.8 | 51.7 | 12269 | 10.0 | 8.0 | 1103 | 1726.9 | 531.1 | 20 | 572.2 | 244.3 | 172 |
| | 2 | 82.3 | 51.1 | 12169 | 8.3 | 6.6 | 1086 | 1429.7 | 440.1 | 20 | 533.4 | 228.2 | 173 |
| | 1 | 72.6 | 42.9 | 8777 | 5.7 | 4.3 | 923 | 860.1 | 265.3 | 21 | 452.2 | 194.0 | 176 |
| BOP | 3 | 86.0 | 62.0 | 14562 | 8.9 | 7.1 | 1123 | 1702.7 | 523.7 | 20 | 571.2 | 244.0 | 172 |
| | 2 | 82.3 | 51.1 | 9439 | 8.3 | 6.6 | 1090 | 1429.7 | 440.1 | 20 | 533.4 | 228.2 | 174 |
| | 1 | 66.5 | 33.5 | 4495 | 3.5 | 2.6 | 860 | 886.3 | 273.3 | 21 | 452.0 | 193.9 | 176 |
| BMP | 3 | 78.1 | 54.4 | 13139 | 9.1 | 7.3 | 1123 | 1964.3 | 603.7 | 20 | 608.4 | 259.4 | 170 |
| | 2 | 0.0 | 0.0 | 143 | 14.7 | 11.8 | 1102 | 1964.3 | 603.7 | 20 | 608.4 | 259.4 | 170 |
| | 1 | 0.0 | 0.0 | 0 | 8.3 | 6.2 | 895 | 1964.3 | 603.7 | 20 | 608.4 | 259.4 | 170 |
| OMP | 3 | 86.4 | 61.9 | 14296 | 7.9 | 6.3 | 1107 | 1673.8 | 514.8 | 20 | 566.7 | 242.1 | 173 |
| | 2 | 86.4 | 54.1 | 9589 | 8.2 | 6.6 | 1098 | 1415.3 | 435.7 | 20 | 531.0 | 227.2 | 174 |
| | 1 | 88.4 | 49.9 | 6719 | 7.9 | 6.3 | 1084 | 920.5 | 283.8 | 21 | 458.1 | 196.5 | 176 |
| BO | 3 | 73.2 | 50.0 | 12324 | 9.9 | 7.9 | 1106 | 1710.2 | 526.1 | 20 | 573.1 | 244.8 | 173 |
| | 2 | 77.5 | 52.1 | 12204 | 9.2 | 7.3 | 1087 | 1432.9 | 441.1 | 21 | 532.7 | 227.9 | 175 |
| | 1 | 70.1 | 41.7 | 9020 | 5.5 | 4.2 | 924 | 923.6 | 284.8 | 21 | 457.6 | 196.3 | 177 |
| BM | 3 | 69.2 | 47.2 | 11719 | 10.0 | 8.0 | 1103 | 1863.1 | 572.8 | 20 | 592.7 | 252.8 | 172 |
| | 2 | 65.3 | 43.8 | 11098 | 9.4 | 7.5 | 1085 | 1863.1 | 572.8 | 20 | 592.7 | 252.8 | 172 |
| | 1 | 43.8 | 25.9 | 6696 | 6.2 | 4.7 | 924 | 1863.1 | 572.8 | 20 | 592.7 | 252.8 | 172 |
| BP | 3 | 76.5 | 53.1 | 12996 | 9.2 | 7.4 | 1122 | 1934.9 | 594.7 | 20 | 605.7 | 258.4 | 171 |
| | 2 | 0.0 | 0.0 | 366 | 14.6 | 11.7 | 1103 | 1934.9 | 594.7 | 20 | 605.7 | 258.4 | 171 |
| | 1 | 0.0 | 0.0 | 0 | 8.2 | 6.2 | 902 | 1934.9 | 594.7 | 20 | 605.7 | 258.4 | 171 |
| OM | 3 | 73.3 | 50.1 | 12330 | 9.3 | 7.4 | 1091 | 1711.5 | 526.5 | 20 | 570.2 | 243.4 | 171 |
| | 2 | 80.8 | 55.1 | 12648 | 9.3 | 7.4 | 1091 | 1417.6 | 436.5 | 21 | 529.8 | 226.5 | 172 |
| | 1 | 88.2 | 58.7 | 12259 | 9.4 | 7.5 | 1091 | 868.4 | 267.8 | 21 | 452.0 | 193.9 | 175 |
| OP | 3 | 75.3 | 52.0 | 13345 | 8.7 | 7.0 | 1109 | 1741.4 | 535.6 | 20 | 576.0 | 246.0 | 172 |
| | 2 | 74.1 | 47.9 | 11420 | 8.8 | 7.0 | 1100 | 1487.2 | 457.7 | 20 | 540.7 | 231.2 | 173 |
| | 1 | 78.3 | 48.7 | 10469 | 8.2 | 6.5 | 1084 | 968.7 | 298.7 | 21 | 465.4 | 199.6 | 176 |
| MP | 3 | 76.7 | 52.7 | 12698 | 8.5 | 6.8 | 1108 | 1919.6 | 590.1 | 20 | 600.0 | 255.9 | 171 |
| | 2 | 0.0 | 0.0 | 624 | 14.6 | 11.7 | 1108 | 1919.6 | 590.1 | 20 | 600.0 | 255.9 | 171 |
| | 1 | 0.0 | 0.0 | 0 | 13.4 | 10.7 | 1089 | 1919.6 | 590.1 | 20 | 600.0 | 255.9 | 171 |
| B | 3 | 67.0 | 45.6 | 11678 | 10.2 | 8.2 | 1103 | 1894.0 | 582.2 | 20 | 592.6 | 252.7 | 171 |
| | 2 | 63.5 | 42.6 | 11102 | 9.7 | 7.7 | 1087 | 1894.0 | 582.2 | 20 | 592.6 | 252.7 | 171 |
| | 1 | 43.5 | 25.8 | 6889 | 6.5 | 4.9 | 935 | 1894.0 | 582.2 | 20 | 592.6 | 252.7 | 171 |
| O | 3 | 71.4 | 48.2 | 11929 | 9.5 | 7.6 | 1089 | 1725.6 | 530.8 | 20 | 569.9 | 243.3 | 172 |
| | 2 | 78.7 | 53.1 | 12266 | 9.5 | 7.6 | 1089 | 1435.2 | 441.8 | 20 | 531.8 | 227.4 | 174 |
| | 1 | 86.0 | 56.5 | 11878 | 9.6 | 7.7 | 1089 | 891.8 | 275.0 | 20 | 458.2 | 196.5 | 176 |
| M | 3 | 67.2 | 45.3 | 11311 | 9.6 | 7.7 | 1091 | 1860.2 | 571.9 | 20 | 588.6 | 251.0 | 171 |
| | 2 | 67.2 | 45.3 | 11311 | 9.6 | 7.7 | 1091 | 1860.2 | 571.9 | 20 | 588.6 | 251.0 | 171 |
| | 1 | 67.2 | 45.3 | 11311 | 9.6 | 7.7 | 1091 | 1860.2 | 571.9 | 20 | 588.6 | 251.0 | 171 |
| P | 3 | 73.4 | 50.5 | 12719 | 8.7 | 7.0 | 1107 | 1900.5 | 584.3 | 20 | 600.7 | 256.2 | 171 |
| | 2 | 0.0 | 0.0 | 280 | 14.9 | 12.0 | 1108 | 1900.5 | 584.3 | 20 | 600.7 | 256.2 | 171 |
| | 1 | 0.0 | 0.0 | 0 | 13.9 | 11.1 | 1092 | 1900.5 | 584.3 | 20 | 600.7 | 256.2 | 171 |
| None | 3 | 65.1 | 44.4 | 11767 | 9.3 | 7.4 | 1092 | 1875.8 | 576.7 | 20 | 590.4 | 251.7 | 170 |

### References

Andersen, K. H., Jacobsen, N. S., and Farnsworth, K. D. (2016). "The theoretical foundations for size spectrum models of fish communities". In: *Canadian Journal of Fisheries and Aquatic Sciences* 73.4, pp. 575–588. DOI: 10.1139/cjfas-2015-0230.

Baird, D., Christian, R. R., Peterson, C. H., and Johnson, G. A. (2004). "CONSEQUENCES OF HYPOXIA ON ESTUARINE ECOSYSTEM FUNCTION: ENERGY DIVERSION FROM CONSUMERS TO MICROBES". In: *Ecological Applications* 14.3, pp. 805–822. DOI: 10.1890/02-5094.

Barot, S., Heino, M., O'Brien, L., and Dieckmann, U. (2003). "Estimating Reaction Norms for Age and Size at Maturation When Age at First Reproduction is Unknown". In: URL: https://pure.iiasa.ac.at/7040.

Beverton, R. J. and Holt, S. J. (2012). *On the Dynamics of Exploited Fish Populations*. Vol. 11. Springer Science & Business Media.

Blanchard, J. L., Andersen, K. H., Scott, F., Hintzen, N. T., Piet, G., and Jennings, S. (2014). "Evaluating targets and trade-offs among fisheries and conservation objectives using a multispecies size spectrum model". In: *Journal of Applied Ecology* 51.3, pp. 612–622. DOI: 10.1111/1365-2664.12238.

Byrd, R. H., Lu, P., Nocedal, J., and Zhu, C. (1995). "A limited memory algorithm for bound constrained optimization". In: *SIAM Journal on Scientific Computing* 16.5, pp. 1190–1208. DOI: 10.1137/0916069.

Calder-Potts, R., Spicer, J., Calosi, P., Findlay, H., and Widdicombe, S. (2015). "A mesocosm study investigating the effects of hypoxia and population density on respiration and reproductive biology in the brittlestar *Amphiura filiformis*". In: *Marine Ecology Progress Series* 534, pp. 135–147. DOI: 10.3354/meps11379.

Carpenter, B., Gelman, A., Hoffman, M. D., Lee, D., Goodrich, B., Betancourt, M., Brubaker, M., Guo, J., Li, P., and Riddell, A. (2017). "Stan: A probabilistic programming language". In: *Journal of Statistical Software* 76.1, pp. 1–32. DOI: 10.18637/jss.v076.i01.

Casini, M., Cardinale, M., and Arrhenius, F. (2004). "Feeding preferences of herring (*Clupea harengus*) and sprat (*Sprattus sprattus*) in the southern Baltic Sea". In: *ICES Journal of Marine Science* 61.8, pp. 1267–1277. DOI: 10.1016/j.icesjms.2003.12.011.

Casini, M., Käll, F., Hansson, M., Plikshs, M., Baranova, T., Karlsson, O., Lundström, K., Neuenfeldt, S., Gårdmark, A., and Hjelm, J. (2016). "Hypoxic areas, density-dependence and food limitation drive the body condition of a heavily exploited marine fish predator". In: *Royal Society open science* 3.160416. DOI: 10.1098/rsos.160416.

Casini, M., Tian, H., Hansson, M., Grygiel, W., Strods, G., Statkus, R., Sepp, E., Gröhsler, T., Orio, A., and Larson, N. (2019). "Spatio-temporal dynamics and behavioural ecology of a "demersal" fish population as detected using research survey pelagic trawl catches: the Eastern Baltic Sea cod (*Gadus morhua*)". In: *ICES Journal of Marine Science* 76.6, pp. 1591–1600. DOI: 10.1093/icesjms/fsz016.

Chabot, D. and Claireaux, G. (2008). "Environmental hypoxia as a metabolic constraint on fish: the case of Atlantic cod, *Gadus morhua*". In: *Marine Pollution Bulletin* 57.6, pp. 287–294. DOI: 10.1016/j.marpolbul.2008.04.001.

Chapman, L. J. and Mckenzie, D. J. (2009). “Behavioral Responses and Ecological Consequences”. In: *Fish Physiology*. Vol. 27. Elsevier, pp. 25–77. DOI: 10.1016/S1546-5098(08)00002-2.

Cooke, R. M., Wittmann, M. E., Lodge, D. M., Rothlisberger, J. D., Rutherford, E. S., Zhang, H., and Mason, D. M. (2014). “Out-of-Sample Validation for Structured Expert Judgment of Asian Carp Establishment in Lake Erie”. In: *Integrated Environmental Assessment and Management* 10.4, pp. 522–528. DOI: 10.1002/ieam.1559.

Diaz, R. J. (2001). “Overview of Hypoxia around the World”. In: *Journal of Environmental Quality* 30.2, pp. 275–281. DOI: 10.2134/jeq2001.302275x.

Diaz, R. J. and Breitburg, D. L. (2009). “The Hypoxic Environment”. In: *Fish Physiology*. Vol. 27. Elsevier, pp. 1–23. DOI: 10.1016/S1546-5098(08)00001-0.

Dieckmann, U. and Heino, M. (2007). “Probabilistic maturation reaction norms: their history, strengths, and limitations”. In: *Marine Ecology Progress Series* 335, pp. 253–269. DOI: 10.3354/meps335253.

Draganik, B., Ivanow, S., Tomczak, M., Maksimov, B., and Psuty-Lipska, I. (2007). “Status of exploited Baltic flounder stocks in the southern Baltic area (ICES SD 26)”. In: *Oceanological and Hydrobiological Studies* 36.4, pp. 47–64. DOI: 10.2478/v10009-007-0029-y.

Eby, L. A., Crowder, L. B., McClellan, C. M., Peterson, C. H., and Powers, M. J. (2005). “Habitat degradation from intermittent hypoxia: impacts on demersal fishes”. In: *Marine Ecology Progress Series* 291, pp. 249–262. DOI: 10.3354/meps291249.

Gray, J. S., Wu, R. S.-s., and Or, Y. Y. (2002). “Effects of hypoxia and organic enrichment on the coastal marine environment”. In: *Marine Ecology Progress Series* 238, pp. 249–279. DOI: 10.3354/meps238249.

Grygiel, W. and Wyszyński, M. (2002). “Sexual maturation of the southern Baltic herring and sprat (1980-2001)”. In: *International Council Meeting*.

Haahtela, I. (1990). “What do Baltic studies tell us about the isopod *Saduria entomon* (L.)?” In: *Annales Zoologici Fennici*. Vol. 27. 3. JSTOR, pp. 269–278. URL: https://www.jstor.org/stable/23736048.

Haase, K., Orio, A., Pawlak, J., Pachur, M., and Casini, M. (2020). “Diet of dominant demersal fish species in the Baltic Sea: Is flounder stealing benthic food from cod?” In: *Marine Ecology Progress Series* 645, pp. 159–170. DOI: 10.3354/meps13360.

Hansson, M. and Viktorsson, L. (2020). *Oxygen Survey in the Baltic Sea 2020 - Extent of Anoxia and Hypoxia, 1960-2020*. Tech. rep. 70. Göteborg, Sweden: Swedish Meteorological and Hydrological Institute.

Hernroth, L., ed. (1985). *Recommendations on methods for marine biological studies in the Baltic Sea*. Working Group 14. 10. Baltic Marine Biologists:Working Group 9. Institute of Marine Research, Lysekill, Sweden.

Hill, C., Quigley, M. A., Cavaletto, J. F., and Gordon, W. (1992). “Seasonal changes in lipid content and composition in the benthic amphipods *Monoporeia affinis* and *Pontoporeia femorata*”. In: *Limnology and Oceanography* 37.6, pp. 1280–1289. DOI: 10.4319/lo.1992.37.6.1280.

Hrycik, A. R., Almeida, L. Z., and Höök, T. O. (2017). “Sub-lethal effects on fish provide insight into a biologically-relevant threshold of hypoxia”. In: *Oikos* 126.3, pp. 307–317. DOI: 10.1111/oik.03678.

ICES (2020a). *Baltic Fisheries Assessment Working Group (WGBFAS)*. Tech. rep. 45. ICES Scientific Reports, p. 643. DOI: 10.17895/ices.pub.6024.

— (2020b). *ICES Database of Trawl Surveys (DATRAS)*. https://www.ices.dk/data/data-portals/Pages/DATRAS.aspx. Extracted 9 November 2020 from Baltic International Trawl Survey (BITS). Copenhagen.

— (2020c). *ICES Year of the Stomach*. https://www.ices.dk/data/data-portals/Pages/Fish-stomach.aspx. Extracted 26 June 2020 from ICES Year of the Stomach. Copenhagen.

Jerde, C. L., Kraskura, K., Eliason, E. J., Csik, S. R., Stier, A. C., and Taper, M. L. (2019). "Strong Evidence for an Intraspecific Metabolic Scaling Coefficient Near 0.89 in Fish". In: *Frontiers in Physiology* 10.1166. DOI: 10.3389/fphys.2019.01166.

Jordan, A. D. and Steffensen, J. F. (2007). "Effects of Ration Size and Hypoxia on Specific Dynamic Action in the Cod". In: *Physiological and Biochemical Zoology* 80.2. PMID: 17252514, pp. 178–185. DOI: 10.1086/510565.

Killen, S. S., Glazier, D. S., Rezende, E. L., Clark, T. D., Atkinson, D., Willener, A. S., and Halsey, L. G. (2016). "Ecological Influences and Morphological Correlates of Resting and Maximal Metabolic Rates across Teleost Fish Species". In: *The American Naturalist* 187.5, pp. 592–606. DOI: 10.1086/685893.

Kramer, D. L. (1987). "Dissolved oxygen and fish behavior". In: *Environmental Biology of Fishes* 18.2, pp. 81–92. DOI: 10.1007/BF00002597.

Limburg, K. E. and Casini, M. (2018). "Effect of Marine Hypoxia on Baltic Sea Cod *Gadus morhua*: evidence From Otolith Chemical Proxies". In: *Frontiers in Marine Science* 5.482. DOI: 10.3389/fmars.2018.00482.

Lindmark, M., Audzijonyte, A., Blanchard, J. L., and Gårdmark, A. (2022). "Temperature impacts on fish physiology and resource abundance lead to faster growth but smaller fish sizes and yields under warming". In: *Global Change Biology* 28.21, pp. 6239–6253. DOI: 10.1111/gcb.16341.

Lowell, R. B. and Culp, J. M. (1999). "Cumulative effects of multiple effluent and low dissolved oxygen stressors on mayflies at cold temperatures". In: *Canadian Journal of Fisheries and Aquatic Sciences* 56.9, pp. 1624–1630. DOI: 10.1139/f99-091.

Mackay, D. and Moreau, J. (1990). "A note on the inverse function of the von Bertalanffy growth function". In: 8.1, pp. 28–31. URL: https://hdl.handle.net/20.500.12348/3147.

*Manual for the Baltic International Trawl Surveys* (2017). Series of ICES Survey Protocols SISP 7 – BITS Version 2.0. ICES Baltic International Fish Survey Working Group, p. 95. DOI: 10.17895/ices.pub.2883.

Möllmann, C. and Köster, F. (1999). "Food consumption by clupeids in the Central Baltic: evidence for top-down control?" In: *ICES Journal of Marine Science* 56, pp. 100–113. DOI: 10.1006/jmsc.1999.0630.

Möllmann, C. and Köster, F. W. (2002). "Population dynamics of calanoid copepods and the implications of their predation by clupeid fish in the Central Baltic Sea". In: *Journal of Plankton Research* 24.10, pp. 959–978. DOI: 10.1093/plankt/24.10.959.

Muusze, B., Marcon, J., Thillart, G. van den, and Almeida-Val, V. (1998). "Hypoxia tolerance of Amazon fish: respirometry and energy metabolism of the cichlid *Astronotus ocellatus*". In: *Comparative Biochemistry and Physiology Part A: Molecular & Integrative Physiology* 120.1, pp. 151–156. DOI: 10.1016/S1095-6433(98)10023-5.

Nelson, P. (2004). *Biological Physics*. WH Freeman New York.
Neuenfeldt, S., Andersen, K. H., and Hinrichsen, H.-H. (2009). "Some Atlantic cod *Gadus morhua* in the Baltic Sea visit hypoxic water briefly but often". In: *Journal of Fish Biology* 75, pp. 290–294. DOI: 10.1111/j.1095-8649.2009.02281.x.
Nilsson, G. E. and Östlund-Nilsson, S. (2008). "Does size matter for hypoxia tolerance in fish?" In: *Biological Reviews* 83.2, pp. 173–189. DOI: 10.1111/j.1469-185X.2008.00038.x.
Orio, A., Bergström, U., Florin, A.-B., Lehmann, A., Šics, I., and Casini, M. (2019). "Spatial contraction of demersal fish populations in a large marine ecosystem". In: *Journal of Biogeography* 46.3, pp. 633–645. DOI: 10.1111/jbi.13510.
Pan, Y., Ern, R., and Esbaugh, A. (2016). "Hypoxia tolerance decreases with body size in red drum *Sciaenops ocellatus*". In: *Journal of Fish Biology* 89.2, pp. 1488–1493. DOI: 10.1111/jfb.13035.
Pichavant, K., Person-Le-Ruyet, J., Bayon, N. L., Severe, A., Roux, A. L., and Boeuf, G. (2001). "Comparative effects of long-term hypoxia on growth, feeding and oxygen consumption in juvenile turbot and European sea bass". In: *Journal of Fish Biology* 59.4, pp. 875–883. DOI: 10.1111/j.1095-8649.2001.tb00158.x.
Rogers, N. J., Urbina, M. A., Reardon, E. E., McKenzie, D. J., Birchenough, S. N., and Wilson, R. W. (2021). *Database of critical oxygen level ($P_{crit}$) in freshwater and marine fishes 1974–2015*. Version 1. Cefas, UK. DOI: 10.14466/CefasDataHub.121.
Rogers, N. J., Urbina, M. A., Reardon, E. E., McKenzie, D. J., and Wilson, R. W. (2016). "A new analysis of hypoxia tolerance in fishes using a database of critical oxygen level ($\mathrm{P_{crit}}$)". In: *Conservation Physiology* 4.1. cow012. DOI: 10.1093/conphys/cow012.
Rohatgi, A. (2021). *Webplotdigitizer: Version 4.5*. URL: https://automeris.io/WebPlotDigitizer.
Schinke, H. and Matthäus, W. (1998). "On the causes of major Baltic inflows–an analysis of long time series". In: *Continental Shelf Research* 18.1, pp. 67–97. DOI: 10.1016/S0278-4343(97)00071-X.
Scott, F., Blanchard, J. L., and Andersen, K. H. (2014). "mizer: an R package for multispecies, trait-based and community size spectrum ecological modelling". In: *Methods in Ecology and Evolution* 5.10, pp. 1121–1125. DOI: 10.1111/2041-210X.12256.
Shepherd, J. (1999). "Extended survivors analysis: an improved method for the analysis of catch-at-age data and abundance indices". In: *ICES Journal of Marine Science* 56.5, pp. 584–591. DOI: 10.1006/jmsc.1999.0498.
Shimps, E. L., Rice, J. A., and Osborne, J. A. (2005). "Hypoxia tolerance in two juvenile estuary-dependent fishes". In: *Journal of Experimental Marine Biology and Ecology* 325.2, pp. 146–162. DOI: 10.1016/j.jembe.2005.04.026.
Small, K., Kopf, R. K., Watts, R. J., and Howitt, J. (2014). "Hypoxia, Blackwater and Fish Kills: experimental Lethal Oxygen Thresholds in Juvenile Predatory Lowland River Fishes". In: *PLoS ONE* 9, pp. 1–10. DOI: 10.1371/journal.pone.0094524.
Snoeijs-Leijonmalm, P. (2017). "Patterns of biodiversity". In: *Biological Oceanography of the Baltic Sea*. Ed. by P. Snoeijs-Leijonmalm, H. Schubert, and T. Radziejewska. Springer Netherlands, pp. 123–191. DOI: 10.1007/978-94-007-0668-2_4.
Vainikka, A., Gårdmark, A., Bland, B., and Hjelm, J. (2009). "Two-and three-dimensional maturation reaction norms for the eastern Baltic cod, *Gadus morhua*". In: *ICES Journal of Marine Science* 66.2, pp. 248–257. DOI: 10.1093/icesjms/fsn199.

Wang, T., Lefevre, S., Huong, D. T. T., Cong, N. van, and Bayley, M. (2009). “The Effects of Hypoxia on Growth and Digestion”. In: *Fish Physiology*. Vol. 27. Elsevier, pp. 361–396. DOI: 10.1016/S1546-5098(08)00008-3.

Wood, S. N., Pya, N., and Säfken, B. (2016). “Smoothing Parameter and Model Selection for General Smooth Models”. In: *Journal of the American Statistical Association* 111.516, pp. 1548–1563. DOI: 10.1080/01621459.2016.1180986.

Wu, R. S. (2009). “Effects of Hypoxia on Fish Reproduction and Development”. In: *Fish Physiology*. Vol. 27. Elsevier, pp. 79–141. DOI: 10.1016/S1546-5098(08)00003-4.
